# Hyper-Truncated Glycans Augment the Activity of Neutrophil Granule Myeloperoxidase

**DOI:** 10.1101/2020.07.24.219956

**Authors:** Harry C. Tjondro, Julian Ugonotti, Rebeca Kawahara, Sayantani Chatterjee, Ian Loke, Siyun Chen, Fabian Soltermann, Hannes Hinneburg, Benjamin L. Parker, Vignesh Venkatakrishnan, Regis Dieckmann, Oliver C. Grant, Johan Bylund, Alison Rodger, Robert J. Woods, Anna Karlsson-Bengtsson, Weston B. Struwe, Morten Thaysen-Andersen

**Author notes:** Contributed equally (Harry C. Tjondro, Julian Ugonotti, Rebeca Kawahara, Sayantani Chatterjee). **Corresponding author:** Dr Morten Thaysen-Andersen, PhD, Department of Molecular Sciences, Macquarie University, NSW-2109, Sydney, Australia.

## Abstract

Myeloperoxidase (MPO) plays essential roles in neutrophil-mediated immunity via the generation of reactive oxidation products. Complex carbohydrates decorate MPO at discrete sites, but their functional relevance remain elusive. To this end, we have characterised the structure-biosynthesis-activity relationship of neutrophil MPO (nMPO). Mass spectrometry demonstrated that nMPO carries both characteristic under-processed and hyper-truncated glycans. Occlusion of the Asn355/Asn391-glycosylation sites and the Asn323-/Asn483-glycans, located in the MPO dimerisation zone, was found to affect the local glycan processing, thereby providing a molecular basis of the site-specific nMPO glycosylation. Native mass spectrometry, mass photometry, and glycopeptide profiling revealed significant molecular complexity of diprotomeric nMPO arising from heterogeneous glycosylation, oxidation, chlorination and polypeptide truncation variants, and a previously unreported low-abundance monomer. Longitudinal profiling of maturing, mature, granule-separated, and pathogen-stimulated neutrophils demonstrated that nMPO is dynamically expressed during granulopoiesis, unevenly distributed across granules and degranulated upon activation. We also show that proMPO-to-MPO maturation occurs during early/mid-stage granulopoiesis. While similar global MPO glycosylation was observed across conditions, the conserved Asn355-/Asn391-sites displayed elevated glycan hyper-truncation, which correlated with higher enzyme activities of MPO in distinct granule populations. Enzymatic trimming of the Asn355-/Asn391-glycans recapitulated the activity gain and showed that nMPO carrying hyper-truncated glycans at these positions exhibits increased thermal stability, polypeptide accessibility, and ceruloplasmin-mediated inhibition potential relative to native nMPO. Finally, structural modelling revealed that hyper-truncated Asn355-glycans positioned in the MPO-ceruloplasmin interface are critical for uninterrupted inhibition. Here, through an innovative and comprehensive approach, we report novel functional roles of MPO glycans, providing new insight into neutrophil-mediated immunity.

**Significance:** Myeloperoxidase (MPO) is an important microbicidal glycoprotein critical for fighting pathogens. We report, for the first time, the intriguingly complex relationship between glycobiology and MPO immune function by demonstrating that uncommon and strategically positioned hyper-truncated glycans both elevate the activity and the inhibition potential of this pathogen-combating enzyme. We have used a multifaceted approach employing integrated biomolecular analytics to generate new insights into the sugar code of MPO. The findings described in this study improve our understanding of key innate immune processes and may guide future glycoengineering efforts aiming to generate therapeutically relevant recombinant MPO products with tuneable activity and inhibition potential tailored to biomedical applications involving persisting and severe pathogen infections.

## Introduction

The myeloid lineage-specific myeloperoxidase (MPO) is an essential component of the innate immune system associated with many pathologies including cardiovascular diseases (Cheng et al., 2019; Kim et al., 2018), rheumatoid arthritis (Stamp et al., 2012), and multiple sclerosis (Gray et al., 2008). Mutations in human *MPO* and genetic ablation in mice have repeatedly been linked to enhanced pathogen infection (Aratani et al., 2006; Lehrer and Cline, 1969; Nauseef et al., 1998; Takeuchi et al., 2012).

While our knowledge of MPO has improved considerably over the past decades (Agner, 1958; Harrison and Schultz, 1976; Klebanoff, 1968), new fascinating facets of MPO biology continue to emerge (Delporte et al., 2018; Nauseef, 2018; Vanhamme et al., 2018). Facilitated by its peroxidase activity, MPO is known to catalyse the formation of reactive oxidation products including hypochlorous acid (HOCl) from chloride ions (Cl^-^) and hydrogen peroxide (H_2_O_2_), substrates found in the maturing phagosomes (Davies et al., 2008; Klebanoff, 2005). From the principal residence within the azurophilic (Az) granules, and to a lesser extent within the specific (Sp) and gelatinase (Ge) granules and in secretory vesicles and the plasma membrane (Se/Pl) of neutrophils (Borregaard and Cowland, 1997; Rørvig et al., 2013), MPO is emptied into phagosomes or secreted through degranulation upon neutrophil activation (Bjornsdottir et al., 2016; Borregaard et al., 2007). Within the phagosome, MPO generates highly reactive hypohalous acids and nitrogen dioxide, which readily react to form diverse reactive oxygen species, key microbicidal and immune-regulatory products of the neutrophil MPO-halide system.

The complex biogenesis and maturation of MPO have been intensely studied (Grishkovskaya et al., 2017; Nauseef, 2018). Briefly, human MPO is translated as a single 80 kDa signal peptide-containing polypeptide chain. This preproMPO form undergoes extensive proteolytic processing initiated by the removal of the signal peptide to form apoproMPO. The enzymatically active proMPO form is rapidly generated via heme acquisition in the endoplasmic reticulum (ER) (Nauseef et al., 1992). Protease-driven polypeptide processing then removes the propeptide and an internal hexapeptide, which separates the light (α, 12.5 kDa) and heavy (β, 60–65 kDa) chains that remain covalently linked via the catalytic heme. His261 and Arg405 (UniProtKB numbering) are key catalytic residues of the active site that is formed around the heme moiety positioned between the α- and β-chains. By mechanisms that remain unclear, but presumably occurring within the Golgi, the α- and β-chains are further processed and extensively post-translationally modified before and/or after the formation of the mature MPO diprotomer (ααββ, ∼150 kDa) that is connected by a Cys319-Cys319 bridge (Nauseef, 2018).

The high-resolution crystal structures of neutrophil-derived MPO (nMPO) with and without cognate and artificial ligands have improved our knowledge of the structure of MPO, the position of the heme and other hetero-atoms essential for enzymatic activity (e.g. Ca^2+^), and the interactions to the endogenous inhibitor ceruloplasmin (Blair-Johnson et al., 2001; Carpena et al., 2009; Chapman et al., 2013; Fiedler et al., 2000; Grishkovskaya et al., 2017; Samygina et al., 2013). Thirty years ago Nauseef and colleagues reported on the existence of five sequons for asparagine-linked (*N*-linked) glycosylation (Asn323, Asn355, Asn391, Asn483, Asn729) all located within the mature MPO β-chain (Nauseef, 1986, 1987). Most MPO crystal structures harbour remnants of *N*-glycan moieties, but our structural knowledge of the MPO glycosylation remains immature since complex carbohydrates are usually incompletely resolved with X-ray crystallography.

Three studies have reported on nMPO glycosylation (Ravnsborg et al., 2010; Reiding et al., 2019; Van Antwerpen et al., 2010). In those studies, mass spectrometry-based glycopeptide analyses detailed the site-specific monosaccharide compositions decorating nMPO. Compositions corresponding to uncommon chitobiose core- and paucimannosidic-type *N*-glycans as well as more conventional oligomannosidic- and complex-type *N*-glycans were reported, but neither the glycan fine structures and occupancy at each site, nor the biosynthesis and functional relevance of the *N*-glycans carried by nMPO distributed across the neutrophil granules, were addressed. In fact, the molecular-level knowledge of the roles of nMPO glycosylation is critically missing despite recent reports suggesting that MPO glycans regulate E-selectin binding (Silvescu and Sackstein, 2014), antigenicity (Wang et al., 2018; Yu et al., 2017), and enzyme activity (Van Antwerpen et al., 2010).

Here, we address this fundamental knowledge gap by characterising the structure-biosynthesis-activity relationship of nMPO and by profiling the dynamic expression, protein processing and site-specific glycosylation of MPO from maturing, mature, granule-separated and activated neutrophils. Integration of mass spectrometry, computational and biochemical assays were used to provide new insight, from atomic to macromolecular detail, into the intriguingly complex MPO sugar code and its related fundamental neutrophil glycobiology and MPO-mediated immune processes.

## Materials and Methods

### Donors, Neutrophil Handling, Granule Fractionation, and Neutrophil-Derived MPO

Buffy coats of healthy individuals (Donor a–i) were collected at The Blood Center, Sahlgrenska University Hospital, Gothenburg, Sweden, see **Supplementary Table S1** for sample overview. Resting neutrophils (>99%/95% viability/purity) were isolated (Clemmensen et al., 2014; Nauseef, 2014a; Rørvig et al., 2013). See **Extended Methods** (SI) for details of all experiments.

Neutrophil granules were separated following nitrogen cavitation as described (Clemmensen et al., 2014). Briefly, three-layered Percoll separated the Az, Sp, and Ge granules and the Se/Pl fraction (Donor c–f) while two-layered Percoll separated Az from Sp/Ge granules and Se/Pl fractions (Donor a–b) as validated using granule markers (Feuk-Lagerstedt et al., 2007). Granules were lysed and protein extracts collected.

Resting neutrophils (Donor g–i) were inoculated with *Staphylococcus aureus* (LS1, multiplicity-of-infection (MOI) 1:5, bacteria:neutrophils), 37°C, 0–120 min. Resting neutrophils without *S. aureus* inoculation and with cytochalasin B/ionomycin (CytB/I) and Triton-X 100 stimulation served as activation and a cell death control, respectively. Degranulated MPO (Dg-MPO) and cell death were monitored longitudinally in supernatants using ELISA (ICL LAB) and lactate dehydrogenase (LDH) release using the Cytotoxicity Detection KitPLUS (Sigma).

Protein and transcript profile data of maturing neutrophils including promyelocytes (PMs), metamyelocytes (MMs), band neutrophils (BNs), neutrophils with segmented nuclei (SNs), and circulating polymorphonuclear cells (PMNs, resting neutrophils, included as a control) were obtained by data re-interrogation (Hoogendijk et al., 2019).

Human neutrophil-derived MPO (nMPO, UniProtKB, P05164, >95% purity) was from pooled donor blood (Lee BioSolutions). The purity, concentration, structural integrity and enzyme activity of nMPO was confirmed prior to analysis (see below).

### Glycan Profiling

*N*-glycans were released from nMPO using *Elizabethkingia miricola* peptide-*N*-glycosidase F (Promega) (Jensen et al., 2012). Reduced *N*-glycans were profiled in technical triplicates using porous graphitised carbon liquid chromatography-tandem mass spectrometry (LC-MS/MS) in negative ion polarity on an LTQ Velos Pro ion trap mass spectrometer (Thermo Scientific) (Hinneburg et al., 2019). Glycan fine structures were manually elucidated (Ashwood et al., 2018). RawMeat v2.1 (Vast Scientific) and GlycoMod (Expasy) aided the process. *N*-glycans were quantified from area-under-the-curve (AUC) measurements of extracted ion chromatograms (EICs) using Skyline v20.1.0.76 (Ashwood et al., 2018).

### Glycopeptide Profiling

Glycopeptides and peptides were profiled from i) nMPO, ii) mono- (αβ) and diprotomeric (ααββ)-separated nMPO, iii) endoglycosidase H- (Endo H-) treated and untreated nMPO, iv) granule-separated MPO, and v) Dg-MPO released from pathogen-activated neutrophils. For i) Reduced and carbamidomethylated nMPO was digested in technical triplicates using sequencing-grade porcine trypsin (Promega), ii) Mono- and diprotomeric nMPO were separated using cooled non-reductive SDS-PAGE. Protein bands were in-gel trypsin digested, iii) Endo H-treated and untreated nMPO (see below) were applied separately to SDS-PAGE. The β-chains (53–58 kDa) were in-gel trypsin digested, iv) Isolated granule fractions were briefly introduced into SDS-PAGE gels. Bands containing all granule proteins were in-gel trypsin digested, and v) Released proteins were acetone precipitated, reduced, alkylated, and in-solution trypsin digested. All peptide mixtures were desalted before LC-MS/MS.

Peptides were separated using C18 chromatography and detected using a Q-Exactive HF-X Hybrid Quadrupole-Orbitrap mass spectrometer (Thermo Scientific) in positive ion polarity. LC-MS/MS data were searched against the canonical human MPO (P05164) and/or the human proteome (reviewed UniProtKB entries) using Byonic v3.6.0 (Protein Metrics) and MaxQuant v1.6, see **Supplementary Table S2** for overview. Variable modifications including Met and Trp mono-/di-oxidation, Tyr mono-/di-chlorination and *N*-glycan libraries were included in the searches. Glycopeptides with Byonic PEP-2D scores < 0.001 were considered and manually validated (Kawahara et al., 2018). Non-glycan modified peptides were filtered to peptide-to-spectral matches and protein false discovery rates < 0.01–0.03 (MaxQuant) or PEP-2D scores < 0.001 (Byonic). Glycopeptides were profiled based on AUCs of monoisotopic EICs of glycopeptide precursors using Skyline v20.1.0.76 or Xcalibur v2.2 (Thermo Scientific). Non-glycosylated peptides and proteins were quantified based on precursor intensities using MaxQuant (Cox and Mann, 2008).

### Intact nMPO Analysis using Native MS and Mass Photometry

Top down/native MS was performed of i) the intact α-chain of nMPO and ii) intact nMPO. For i) nMPO was reduced, desalted and injected on a C4 LC column connected to an Agilent 6538 quadrupole-time-of-flight mass spectrometer operating in high-resolution positive polarity mode. Mass spectra were deconvoluted using MassHunter vB.06 (Agilent Technologies). Assignments were guided by the LC-MS/MS peptide data, see **Supplementary Table S3A**, and ii) Intact nMPO was infused into a modified Q-Exactive (Thermo Scientific) operating in positive ion polarity via nano-ESI using custom-made gold-coated capillaries (Gault et al., 2016). Data were processed with Xcalibur v2.2 (Thermo Scientific), spectra deconvoluted with UniDec (Marty et al., 2015), and annotated using in-house software.

Intact nMPO was analysed using single-molecule mass photometry as described (Soltermann et al., 2020). Coverslips were assembled for sample delivery using silicone CultureWell gaskets (Grace Bio-Labs). Data were acquired, processed and analysed using in-house software (Young et al., 2018).

### Visualisation, Modelling, Molecular Dynamics, Solvent Accessibility, and Sequence Alignments

Human diprotomeric MPO (PDBID, 1D2V) was used for visualisation and modelling. Signature *N*-glycans were added *in silico* using the Carbohydrate and Glycoprotein builders within GLYCAM-Web (http://glycam.org) to mimic nMPO (WT), Endo H-treated nMPO (P1), the hyper-truncated Asn355-/Asn391-glycophenotype elevated in Se/Pl-MPO (P2) and an MPO glycoform with semi-truncated glycans at Asn355 and Asn391 (P3), see **Supplementary Table S4**. The per-residue solvent accessibilities, root mean squared deviation/fluctuation (RMSD/RMSF) and secondary structures were calculated using Cpptraj (AmberTools 18), plotted with Gnuplot 5.2 and visualised using VMD 1.9.3. Snapshots from the molecular dynamics (MD) simulations of WT- and P1-MPO were structurally aligned to the ceruloplasmin-MPO complex (4EJX) via the MPO protein backbone.

Five crystal structures of MPO (PDBID, 1D2V, 1CXP, 1DNU, 1DNW, 5FIW) were used to assess the relative solvent accessibilities to the Asn residue of all sequons of monomeric MPO and to the β1,2-GlcNAc of FA1-glycans at Asn323, Asn483 and Asn729 of mono- and diprotomeric MPO using NACCESS (5 Å radii probe) (Hubber and Thornton, 1993).

Sequence alignments of human MPO (P05164) to i) the human peroxidase family including eosinophil peroxidase (P11678), lactoperoxidase (P22079) and thyroid peroxidase (P07202), and ii) MPO from key mammalian species including mouse MPO (P11247), macaque MPO (F7BAA9), porcine MPO (K7GRV6) and bovine MPO (A6QPT4, all downloaded July 2020) were performed using T-Coffee (http://tcoffee.crg.cat/apps/tcoffee) and Boxshade (http://www.ch.embnet.org/software/BOX_form).

### Endoglycosidase H-Treatment of nMPO

Intact nMPO was incubated with or without *Streptomyces plicatus* endoglycosidase H (Endo H, Promega) under native conditions, 37°C, 16 h. All samples including controls containing only Endo H were used immediately for activity and inhibition profiling and structural characterisation.

### Chlorination and Oxidation Activity and Ceruloplasmin-Mediated Inhibition of MPO

Chlorination activities of various MPO glycoforms and controls were determined by the formation of HOCl captured via taurine per time using HOCl standard curves (activity assay 1) (Dypbukt et al., 2005). Reactions were initiated by sequential addition of taurine and H_2_O_2_ (Sigma), stopped by catalase (Sigma) and measured at 650 nm after addition of 3,3′,5,5′-tetramethylbenzidine (TMB, Sigma). Relative oxidation activities of various MPO glycoforms and controls were determined using a TMB assay (activity assay 2) and an *o*-phenylenediamine assay (activity assay 3). Reactions were initiated by the addition of TMB or *o*-phenylenediamine (Sigma), quenched after incubation with sulphuric acid (Sigma) and the colour intensity measured at 450 nm or 492 nm, respectively. Readings were adjusted based on water and Endo H controls, **Supplementary Table S5**.

Ceruloplasmin-mediated inhibition of the MPO enzyme activity was determined using activity assay 1–3. Endo H-treated and untreated nMPO and controls were incubated with and without human serum-derived ceruloplasmin (P00450, Lee BioSolutions) in technical triplicates prior to activity measurements. Readings were adjusted based on water, Endo H and ceruloplasmin controls.

### Circular Dichroism Profiling and Temperature Stability

Circular dichroism (CD) data of nMPO, Endo H-treated nMPO and controls were collected in technical duplicates using 1 mm-pathlength cuvettes (Starna Scientific) in a Jasco J-1500 spectropolarimeter. CD spectra were recorded using 260–190 nm scans at pre-melting temperatures (20–50°C). Thermal stability was determined by monitoring the 208 nm signal over a temperature range. Readings were baseline corrected based on water and Endo H controls.

### Data Representation and Statistics

Significance was tested using one-/two-tailed paired/unpaired Student’s t-tests. Confidence was designated by \**p* < 0.05, \*\**p* < 0.01, \*\*\**p* < 0.005, \*\*\*\**p* < 0.001, \*\*\*\*\**p* < 0.0005. ns, non-significant test (*p* ≥ 0.05). Biological and technical replicates have been stated. Data were plotted as the mean, while error bars represent their standard deviation (SD).

## Results and Discussion

### Comprehensive Characterisation of Neutrophil-Derived Myeloperoxidase

We first sought to unravel the molecular complexity of the heme-containing MPO (ααββ), which adopts a complex diprotomeric structure comprising two identical αβ protomers, **Figure 1A**. Each β-chain harbours five sequons (Asn323, Asn355, Asn391, Asn483, Asn729) enabling extensive *N*-glycosylation of the protein surface. We applied our established glycomics platforms to fully define the molecular structure of all *N*-glycans present on nMPO, including identification of isomeric glycans, **Figure 1B**. Documentation for all reported structures have been provided, **Supplementary Figure S1, Supplementary Table S6** and **Supplementary Data S1**. Uncommon monoantennary complex-type (FA1G1S1a), under-processed oligomannosidic (M5–M6), and hyper-truncated paucimannosidic (M2Fa-M3F) and chitobiose core-type (GlcNAc1–GlcNAc1F) structures were found to be characteristic *N*-glycans of nMPO. These *N*-glycans are congruent with structures residing in the neutrophil granules (Venkatakrishnan et al., 2020) and those carried by other granule-resident glycoproteins including neutrophil cathepsin G and elastase (Loke et al., 2017; Loke et al., 2015; Thaysen-Andersen et al., 2015). No *O*-glycans were detected in the glycomics datasets.

**Figure 1.**
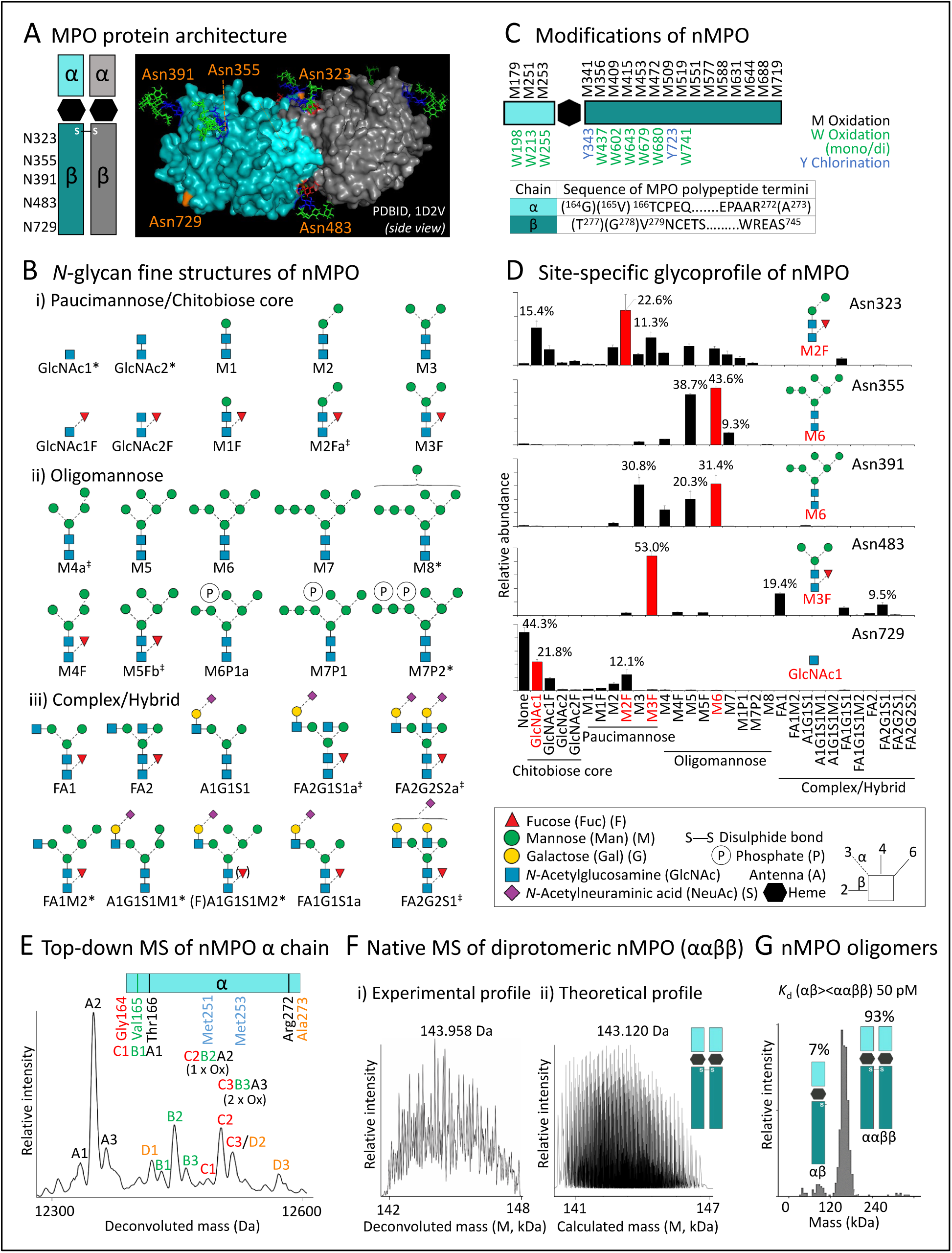
Comprehensive characterisation of neutrophil-derived myeloperoxidase (nMPO). **A**) Protein architecture of the heme-containing diprotomeric MPO (ααββ) (left). Each β-chain harbours five *N*-glycan sequons (orange) mapped on diprotomeric MPO (PDBID, 1D2V). Asn729 was left unoccupied while characteristic glycans were added *in silico* to other sites (see D). **B**) Fine structures and short-hand nomenclature of the nMPO *N*-glycans (**Supplementary Figure S1, Supplementary Table S6** and **Supplementary Data S1**). *Several hyper-truncated and/or trace *N*-glycans were only observed at the peptide level. ^‡^Multiple glycan isomers (designated a, b..) were identified. See key for glycan symbol nomenclature and linkage representation (Varki et al., 2015). **C**) Non-glycan modifications of nMPO including Met (black) and Trp (green) mono- and di-oxidation, Tyr (blue) mono-chlorination and polypeptide truncation variants (**Supplementary Figure S2** and **Supplementary Data S2A-B**). **D**) Site-specific *N*-glycoprofile of nMPO (**Supplementary Figure S3, Supplementary Table S7** and **Supplementary Data S3A-E**). Prominent *N*-glycans at each site are in red. Data plotted as mean ± SD, n = 3, technical replicates. **E**) Intact mass analysis revealed limited α-chain heterogeneity arising from polypeptide truncation and Met251 and Met253 oxidation variants as supported by peptide data (**Supplementary Table S3A**). **F**) i–ii) Native MS revealed significant complexity of diprotomeric nMPO (ααββ) complexes that matched the expected profile generated from approximately 300,000 predicted proteoforms (**Supplementary Table S3B**). **G**) Mass photometry revealed the existence of a previously unreported low-abundance monoprotomer (7%) and a high-abundance diprotomer (93%) of nMPO.

Many non-glycan nMPO modifications including a total of 18 Met and nine Trp mono-/di-oxidations, two sites for Tyr mono-chlorination and several polypeptide truncation variants of both the α- and β-chains were identified using sensitive peptide profiling facilitating a near-complete sequence coverage of nMPO (α-/β-chain, 100.0%/99.8%), **Figure 1C, Supplementary Figure S2** and **Supplementary Data S2A-B**. The observed polypeptide hyper-oxidation, which is known to be mediated by the reactive oxidising agents produced by MPO (Ravnsborg et al., 2010), plays recognised roles in neutrophil function (Winterbourn et al., 2016).

Further, site-specific glycoprofiling revealed an extensive micro- and macro-heterogeneity of all five *N*-glycosylation sites of nMPO, **Figure 1D, Supplementary Figure S3, Supplementary Table S7** and **Supplementary Data S3A-E**. In agreement with studies reporting on the site-specific monosaccharide compositions of nMPO (Ravnsborg et al., 2010; Reiding et al., 2019; Van Antwerpen et al., 2010), we found that Asn323 and Asn483 predominantly carry paucimannosidic-type *N*-glycans (M2F–M3F), Asn355 and Asn391 carry oligomannosidic-type *N*-glycans (M5–M6) while Asn729 is largely unoccupied or carries chitobiose core-type (GlcNAc1–GlcNAc1F) *N*-glycans. We identified low levels of mannose-6-phoshorylation on nMPO relative to a recent study (Reiding et al., 2019). Such discrepancies may arise from differences in the purification and/or analysis of the glycoprotein. Notably, we observed very similar glycosylation of all five sites of nMPO arising from different sources including from unperturbed neutrophil extracts (see detailed comparisons below) and acquired using different techniques supporting an unbiased isolation and characterisation of nMPO.

Intact mass analysis revealed limited α-chain heterogeneity (12,350–12,550 Da) arising from relatively few polypeptide truncation and oxidation variants, presumably Met251 and Met253, as supported by peptide data, **Figure 1E** and **Supplementary Table S3A**. The variable protein modifications identified at the glycopeptide and peptide level were however reflected in our native MS analysis as demonstrated by significant molecular complexity of diprotomeric nMPO (ααββ, 141.5–148.0 kDa, apex 143,958 Da), which matched a published lower-resolution profile of intact nMPO (apex 144,180 Da) (Reiding et al., 2019) as well as a theoretical profile generated from ∼300,000 proteoforms predicted based on quantitative peptide data (140.5–147.0 kDa), **Figure 1F** and **Supplementary Table S3B**. Native MS also indicated the existence of an nMPO monomer with a slightly lower-than-expected molecular mass (αβ, 70–73 kDa, data not shown). Cooled non-reductive SDS-PAGE followed by peptide profiling supported the presence of a maturely processed low-abundance nMPO monomer, **Supplementary Figure S4**. Single-molecule mass photometry, a method capable of quantifying the assembly, binding affinities and kinetics of protein complexes (Haussermann et al., 2019; Soltermann et al., 2020; Young et al., 2018) confirmed that nMPO exists as a low-abundance monoprotomer (αβ, 7%) and high-abundance diprotomer (ααββ, 93%) with an apparent mono-/diprotomer (αβ/ααββ) *K*_d_ of ∼50 pM, **Figure 1G**. The biological role(s) of the maturely processed monoprotomeric MPO, which differs from the secreted monoprotomeric proMPO reportedly elevated in blood in cardiovascular disease (Gorudko et al., 2018; Khalilova et al., 2018), remains unknown.

### Tertiary and Quaternary Structural Features Explain the Site-Specific N-Glycosylation of MPO

The presence of vastly different glycan structures across glycosylation sites on a given protein is interesting because the protein experience the same ensemble of glycan-processing enzymes as it traffics the ER-Golgi pathways. We aimed to identify the biochemical basis for this heterogeneity by first investigating the early-stage glycoprotein processing where MPO enters the *cis*-Golgi as a fully folded monomer (Nauseef, 2018). As we have observed for other mammalian glycoproteins (Thaysen-Andersen and Packer, 2012), strong associations between the Asn accessibility on the MPO surface and the degree of *N*-glycan type processing and core fucosylation were identified, **Figure 2A, Supplementary Table S7** and **Supplementary Table S8A**. The relatively occluded Asn355- and Asn391-glycosylation sites carried under-processed oligomannosidic-type and afucosylated *N*-glycans as opposed to the surface exposed sites (Asn323, Asn483, Asn729) (*p* = 3.5 × 10^−13^ and 5.7 × 10^−13^, respectively) carrying significantly more processed *N*-glycans (*p* = 4.7 × 10^−4^ and 7.6 × 10^−8^) and core fucosylation (both *p* = 1.3 × 10^−7^).

**Figure 2.**
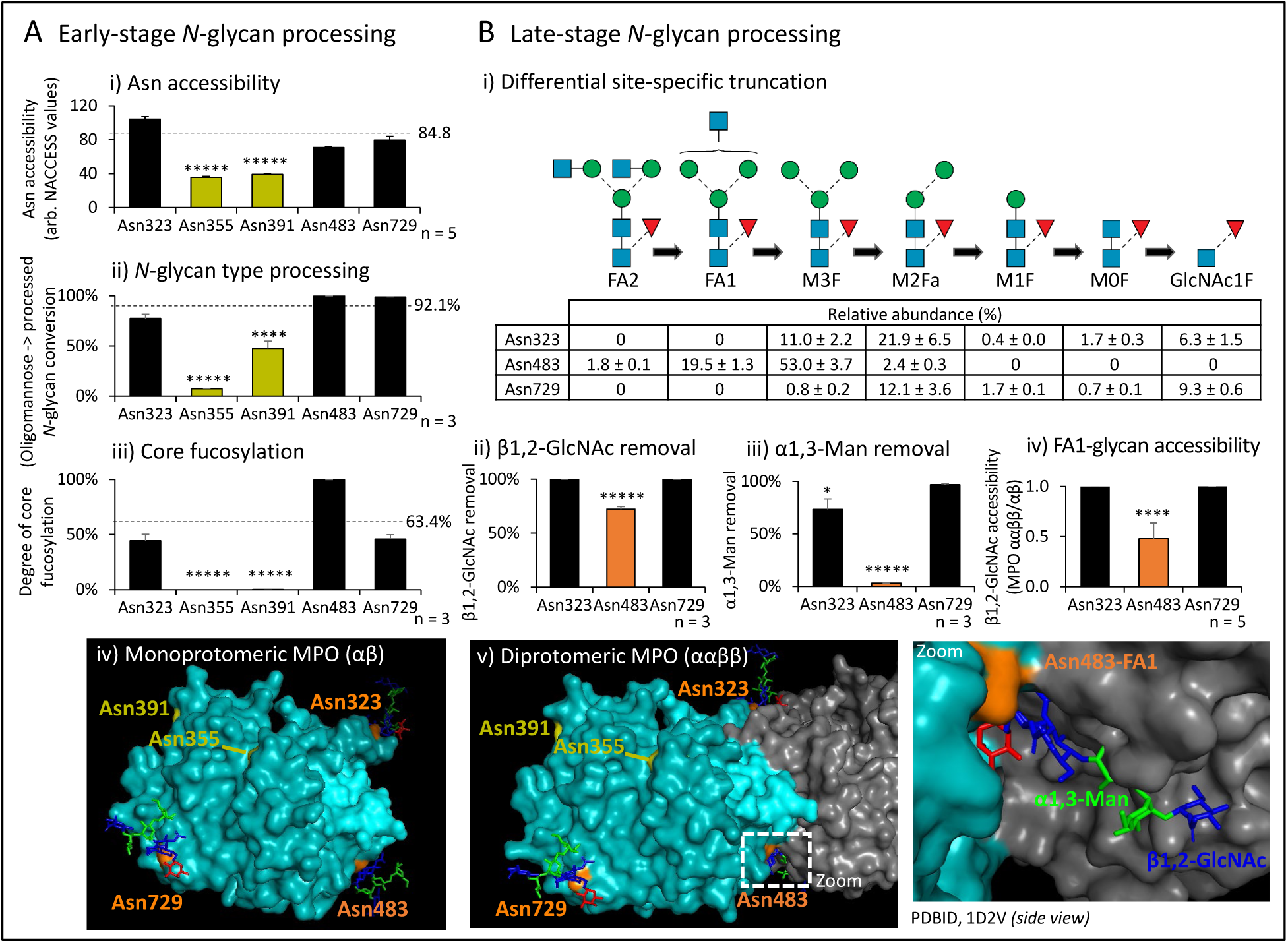
Molecular basis for the site-specific *N*-glycosylation of MPO. **A**) Investigation of maturely folded monoprotomeric MPO (as it appears in the early-stage processing) revealed correlations between i) the Asn accessibility and the degree of ii) *N*-glycan type processing plotted as the oligomannose-to-processed *N*-glycan conversion, and iii) core fucosylation determined from possible FUT8 substrates based on nMPO glycopeptide profiling and solvent accessibility data (**Figure 1D, Supplementary Table S7** and **Supplementary Table S8A**). The occluded Asn355 and Asn391 (green bars) carried mostly under-processed oligomannosidic *N*-glycans and afucosylated *N*-glycans. Mean values for the accessible sites (Asn323, Asn483, Asn729) (broken lines) were statistically compared to Asn355 and Asn391. iv) Occluded Asn355 and Asn391 (green, depicted without glycans for clarity) and accessible Asn323, Asn729 and Asn483 (orange) were mapped on monoprotomeric MPO (PDBID, 1D2V). **B**) i) Exploration of the late-stage glycan processing of maturely folded MPO involving the *N*-glycan truncation pathway (Tjondro et al., 2019; Ugonotti et al., 2020) demonstrated less truncation of the Asn483-glycans and partly Asn323-glycans as measured by the lower removal efficiency of terminal residues including ii) β1,2-GlcNAc and iii) α1,3-Man residues relative to Asn729-glycans based on nMPO glycopeptide data and iv) a reduced β1,2-GlcNAc accessibility of FA1-conjugated Asn483 (orange) upon MPO dimerisation (**Supplementary Table S8B**). Statistical comparisons were made to Asn729. v) Illustration of the dimerisation-dependent occlusion of the Asn483-glycan on diprotomeric MPO (1D2V). Contact between (and hence masking of) the Asn483-FA1 glycan and the protein surface of the other αβ-protomer was observed (see zoom). The Asn323-glycan positioned near the dimer interface was partially affected by MPO dimerisation. For all panels, data plotted as mean ± SD, n = 3 (glycopeptide profiling), n = 5 (solvent accessibility), \**p* < 0.05, \*\**p* < 0.01, \*\*\**p* < 0.005, \*\*\*\**p* < 0.001, \*\*\*\*\**p* < 0.00005. See **Figure 1** for key.

Exploration of the late-stage MPO processing within the *N*-glycan truncation pathway, a glycan-processing pathway highly active in neutrophils (Loke et al., 2015; Tjondro et al., 2019; Ugonotti et al., 2020), demonstrated that the Asn483-glycans (and partly Asn323-glycans) undergo less truncation relative to Asn729-glycans, **Figure 2Bi**. The lower removal efficiency of the outer β1,2-GlcNAc and α1,3-Man residues of Asn483-glycans relative to Asn729-glycans (*p* = 3.5 × 10^−5^ and 1.1 × 10^−8^, respectively) correlated with a reduced solvent accessibility of the β1,2-GlcNAc of Asn483-FA1 glycans upon MPO dimerisation (*p* = 7.6 × 10^−5^), **Figure 2Bii-iv** and **Supplementary Table S8B**. The dimerisation-dependent occlusion of the Asn483-glycans could be observed by the contact between (and masking of) the Asn483-FA1 glycan and the surface of the other αβ-protomer, **Figure 2Bv**. Intuitively, occlusion of the Asn483-glycans restricts the *N*-acetyl-β-hexosaminidase (P06865/P07686) and lysosomal α-mannosidase (O00754), key glycoside hydrolases of the truncation pathway (Tjondro et al., 2019; Ugonotti et al., 2020), to access their glycan substrates and, in turn, less efficiently catalyse β1,2-GlcNAc and α1,3-Man removal of Asn483-FA1 glycans. The Asn323-glycan positioned near the dimer interface was partially affected by MPO dimerisation as indicated by a modestly impaired α1,3-Man removal of Asn323-glycans (*p* = 0.014, Asn323 vs Asn729). The dimerisation-dependent protection of the Asn323- and Asn483-glycans from hydrolase-mediated truncation was supported by glycopeptide profiling of gel-separated mono- and diprotomeric nMPO, **Supplementary Figure S4A**. This analysis confirmed that the outer β1,2-GlcNAc of the Asn323- and Asn483-glycans was more efficiently removed of mono-rather than diprotomeric nMPO (*p* = 0.013 and 0.032, respectively), **Supplementary Table S9A-B**. Similar protective effects could be observed for the Asn323-glycans of diprotomeric MPO exhibiting a lower α1,3-Man removal efficiency relative to monoprotomeric MPO (*p* = 0.008). Importantly, similar processing of the Asn729-glycans, distal to the dimerisation interface, were observed on mono- and diprotomeric nMPO (*p* ≥ 0.05). Collectively, and in line with current literature (Nauseef, 2018), our data show that dimerisation takes place immediately before or quickly after MPO arrives in the neutrophil granules, a maturation step that affects the processing of the Asn323- and Asn483-glycans positioned at the MPO dimer interface.

### Dynamics, Distribution, Processing and Glycosylation of MPO Across Neutrophil Life Stages

We next investigated the possible spatiotemporal expression of nMPO and compartment-specific glycosylation during neutrophil formation and activation. Dynamic expression of MPO mRNA and protein during granulopoiesis was demonstrated by re-interrogation of proteomics data obtained from maturing neutrophils (Hoogendijk et al., 2019), **Figure 3A**. Consistent with a previous protein profiling study of neutrophil granules (Rørvig et al., 2013) and with the targeting-by-timing model that describes the timely packaging of proteins in granules during granulopoiesis (Cowland and Borregaard, 2016), we show MPO predominantly resides in azurophilic (Az, 82.9%) granules while the specific (Sp) and gelatinase (Ge) granules and the secretory vesicles/plasma membrane (Se/Pl) contain less MPO, **Figure 3B**. Neutrophils are known to release their proteinaceous granule content via degranulation upon stimulation (Bjornsdottir et al., 2016). To expand on these findings, we monitored the release and glycosylation patterns of degranulated MPO (Dg-MPO) longitudinally upon neutrophil activation mediated by low-level short-term infection by *S. aureus*, an opportunistic pathogen present in neutrophil-rich tissues undergoing inflammation including, for example, the upper respiratory tract of individuals living with cystic fibrosis, **Figure 3C**.

**Figure 3.**
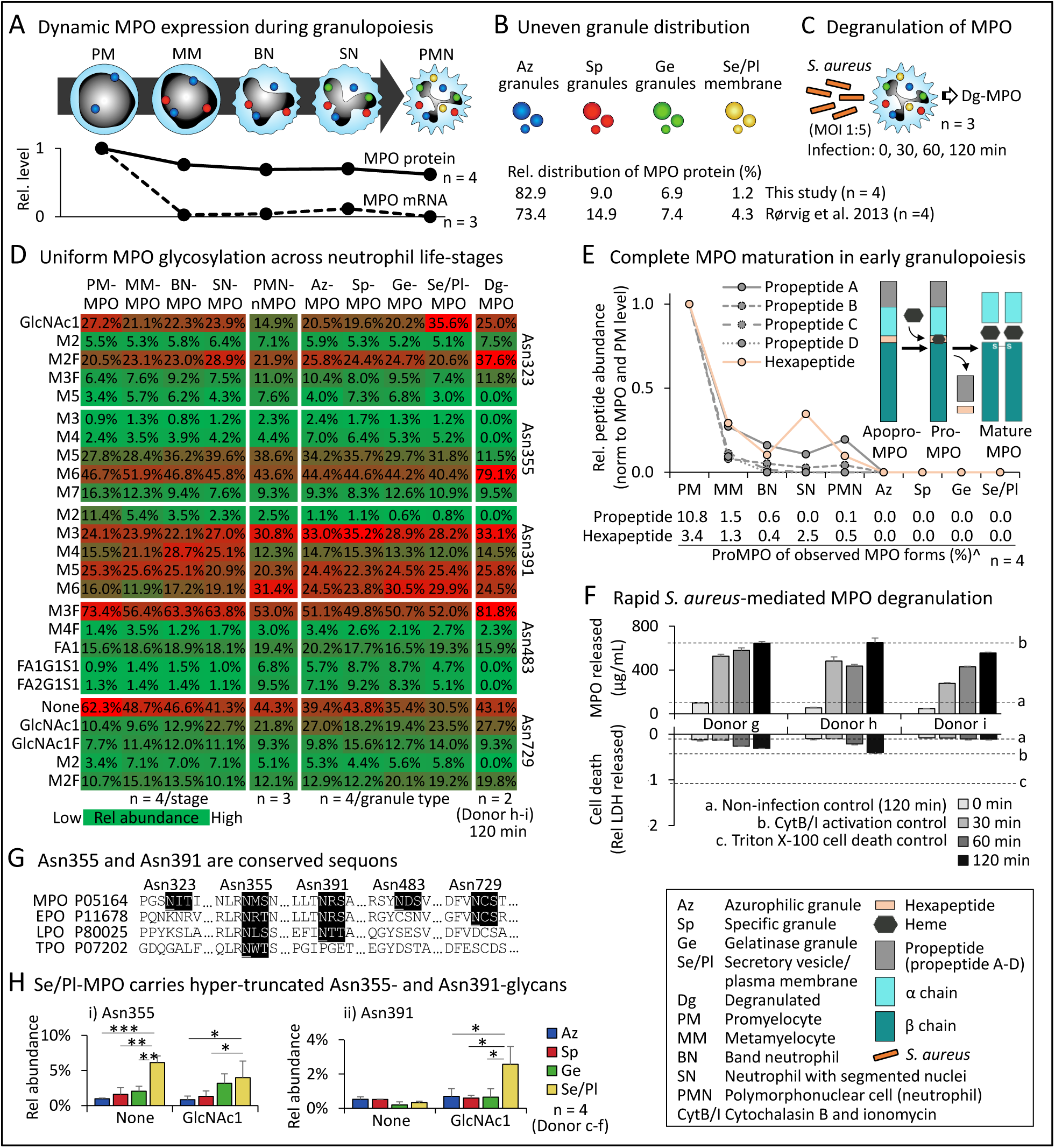
Dynamics, granule distribution, polypeptide processing, site-specific glycosylation, and degranulation of MPO across the neutrophil life stages. **A**) Dynamic expression of MPO mRNA and protein during granulopoiesis. The relative MPO expression levels, normalised to the PM-stage, were established by data re-interrogation (Hoogendijk et al., 2019). See key for nomenclature. **B**) Quantitative proteomics of granule-separated neutrophils showed an uneven MPO distribution across granules congruent with literature (Rørvig et al., 2013). **C**) *S. aureus*-mediated neutrophil activation. Longitudinal profiling of the levels and glycosylation of released Dg-MPO (see Panel F, D for data and controls). **D**) Site-specific glycoprofiling of MPO from maturing, resting (PMN-nMPO, see Figure 1D), granule-separated, and activated neutrophils. Prominent *N*-glycans at each site were plotted (**Supplementary Figure S5** and **Supplementary Table S10**). The MPO glycosylation of maturing neutrophils was determined by data re-interrogation (Hoogendijk et al., 2019). See key for intensity scale and biological replicates and **Figure 1** for short-hand nomenclature of glycans. **E**) Longitudinal tracking of the proMPO-to-MPO conversion within maturing, mature, and granule-separated neutrophils. Relative levels of peptides arising from the proMPO regions (normalised to PM-stage) were plotted. ^ProMPO levels were determined from the intensity of peptide pairs arising from proMPO and mature MPO. The data do not discriminate apoproMPO and proMPO. See insert for schematics of MPO polypeptide maturation. **F**) *S. aureus*-mediated activation of neutrophils (MOI 1:5) demonstrating a rapid time-dependent and cell death-independent degranulation of Dg-MPO as measured by ELISA and LDH release assays. Controls (a–c, mean plotted as broken lines) were included for both assays. **G**) Sequence alignment showed that Asn355 and Asn391 are conserved sequons across the human peroxidases implying functional importance. EPO, eosinophil peroxidase; LPO, lactoperoxidase; TPO, thyroid peroxidase. See **Supplementary Figure S6** for alignment to mammalian MPO. **H**) Hyper-truncated GlcNAcβAsn and absent glycosylation were elevated features at i) Asn355 and ii) Asn391 of Se/Pl-MPO relative to MPO from other granules. For all panels, data plotted as mean ± SD. ns, not significant (*p* ≥ 0.05), \**p* < 0.05, \*\**p* < 0.01, \*\*\**p* < 0.005, \*\*\*\**p* < 0.001, \*\*\*\*\**p* < 0.00005.

Site-specific glycoprofiling of MPO from maturing, mature (resting), granule-separated, and pathogen-activated neutrophils showed that MPO carries relatively similar glycosylation across the neutrophil life-stages and under the conditions that were analysed, **Figure 3D**. Similar MPO glycosylation patterns of all five sites were observed across all granule types (correlation coefficients, r > 0.95), which, as expected, matched the glycosylation of unfractionated nMPO from resting neutrophils (PMN-nMPO, r > 0.95, see **Supplementary Figure S5** and **Supplementary Table S10** for all data). Given our recent glycomics-based report of granule-specific *N*-glycosylation of resting neutrophils (Venkatakrishnan et al., 2020) and the observation of uniform *N*-glycosylation of neutrophil elastase across granules (Loke et al., 2017), the observation of relatively similar *N*-glycosylation of MPO distributed across granules indicates that both the different granule protein populations and differential glycan processing of the proteins trafficking to the individual granules contribute to the compartment-specific glycosylation of neutrophils.

MPO glycoprofiling of maturing neutrophils, facilitated by data re-interrogation (Hoogendijk et al., 2019), indicated that PM-MPO and MPO from all subsequent neutrophil development steps surprisingly carry fully processed *N*-glycosylation signatures at all sites (r > 0.93). Supporting the MPO maturation in early-stage granulopoiesis, longitudinal profiling of the proMPO-to-MPO conversion within maturing, mature, and granule-separated neutrophils showed a near-complete pro- and hexapeptide removal from metamyelocyte-derived MPO (MM-MPO, ∼1% proMPO) and from MPO of more advanced cellular maturation stages, **Figure 3E**. Our data could not discriminate between the apoproMPO and proMPO forms (without and with heme, respectively), but the unprocessed form observed in this study is likely proMPO since heme acquisition reportedly occurs rapidly in the ER (Nauseef et al., 1992). Taken together, our data indicate that the heavily processed MPO undergoes rapid glycan and polypeptide maturation upon expression at the PM-stage during granulopoiesis.

In addition to the characterisation during neutrophil maturation, we also glycoprofiled MPO during pathogenic infection. *S. aureus*-mediated activation of neutrophils performed at sub-stoichiometric levels to simulate the relatively weak chronic infection levels experienced by individuals with cystic fibrosis and to prevent cell lysis (MOI 1:5) demonstrated a rapid (30– 120 min) time-dependent and cell death-independent degranulation of Dg-MPO, **Figure 3F**. The Dg-MPO carried similar glycosylation to PMN-nMPO (r > 0.91) indicating a glycoform-independent degranulation process of the protein. It remains unknown if the weakly altered glycans observed at selected sites (e.g. Asn355, r = 0.81) functionally impact the pathogen-killing ability of MPO or result from technical variation of the analytically challenging site-specific profiling of MPO directly from biological mixtures (Thaysen-Andersen et al., 2016).

Sequence alignments showed that Asn355 and Asn391 are conserved sequons within the family of human peroxidases and within MPO expressed across mammalian species, **Figure 3G** and **Supplementary Figure S6**. The sequon conservation of Asn355 and Asn391, which recently was strengthened by the observation of similar glycosylation patterns of these two sites on human and mouse MPO (Caval et al., 2020), imply a functional relevance of the Asn355- and Asn391-glycans. Interestingly, absence of glycosylation and hyper-truncated GlcNAcβAsn were found to be elevated features of Se/Pl-MPO at Asn355 (*p* = 9.2 × 10^−5^ and 0.04, respectively, Se/Pl- vs Az-MPO) and Asn391 (*p* = 0.022 for GlcNAcβAsn) relative to MPO from other granules, **Figure 3H**. It may be speculated that these minimal Asn355- and Asn391-glycosylation features of Se/Pl-MPO arise from the action of endoglycosidase H- (Endo H-) and peptide:*N*-glycosidase F-like hydrolases e.g. di-*N*-acetylchitobiase (Q01459) and N(4)- (beta-*N*-acetylglucosaminyl)-L-asparaginase (P20933) and/or other glycoside hydrolases of the truncation pathway that may be co-expressed and co-sorted with MPO trafficking to these compartments (Damme et al., 2010; Tjondro et al., 2019). Regardless of the underlying biosynthesis, the position-specific enrichment of uncommon glycan signatures in specific compartments of the neutrophil is intriguing and without precedence in the literature.

### Hyper-Truncated Asn355- and Asn391-GlcNAc Residues Allosterically Augment MPO Activity

Next, we explored the function of MPO glycans, which are unlikely to interfere directly with the substrate-product exchange to the heme-containing catalytic site as these are located distal to the active site, **Figure 4Ai**. Hyper-oxidation of Met251, Met253 and Trp255 lining the catalytic site was observed indicating extensive auto-oxidation that may impact the MPO activity by possibly reconfiguring the active site, **Figure 4Aii**. Activity assays performed on granules of Donor a–b neutrophils fractionated on two-layered gradients demonstrated that Se/Pl-MPO exhibits a higher chlorination activity (3.7–9.0-fold) and oxidation activity (5.0– 51.5-fold, Se/Pl- vs Az-MPO) than MPO from other granules, **Figure 4Aiii** and **Supplementary Table S11**. In support, peptide analysis of granules from Donor c–f neutrophils fractionated on three-layered gradients confirmed that Se/Pl-MPO exhibits a higher chlorination activity than MPO from other granules based on a higher Tyr chlorination level of the granule proteins (*p* = 4.7 × 10^−3^, Se/Pl- vs Az-MPO), **Figure 4Aiv** and **Supplementary Table S12**.

**Figure 4.**
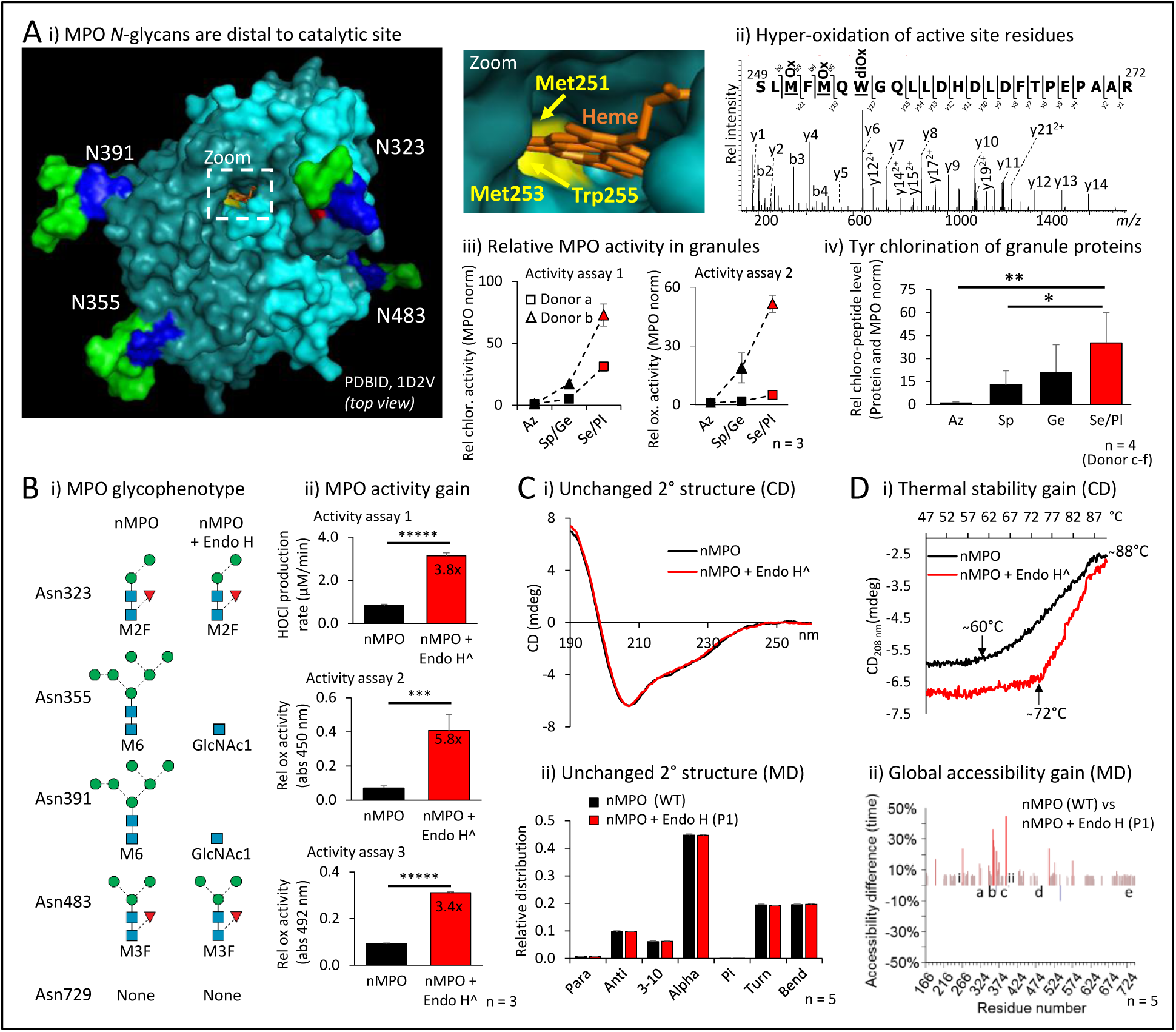
Hyper-truncated Asn355- and Asn391-GlcNAc signatures positioned distal to the catalytic site augment the MPO activity. **A**) i) The MPO *N*-glycans are positioned distal to the catalytic site. Zoom: Oxidation-prone Met251, Met253 and Trp255 (yellow) in the heme-containing (orange) catalytic site. Mapped on monoprotomeric MPO (PDBID, 1D2V), see **Figure 1** for key. ii) Hyper-oxidation of Met251, Met253 and Trp255 indicates extensive auto-oxidation. iii) Se/Pl-MPO from Donor a–b neutrophils displayed a higher chlorination and oxidation activity than MPO from other granules (**Supplementary Table S11**). Adjusted for MPO levels, n = 3, technical replicates. iv) Relative Tyr chlorination level of proteins from granules fractionated with high resolution of neutrophils from Donor c–f (**Supplementary Table S12**). Adjusted for total protein and MPO levels, n = 4, biological replicates. **B**) i) Endo H-treatment of nMPO produced an Asn355- and Asn391-GlcNAc glycophenotype (mimicking the enriched GlcNAcβAsn signatures of Se/Pl-MPO, see **Figure 3H**) as validated using LC-MS/MS (**Figure 5B**). Main glycoforms are depicted for each site. ii) The Endo H-treated nMPO exhibited a higher enzyme activity than native nMPO based on technical triplicate measurements of HOCl production (activity assay 1) and the oxidation rate (activity assays 2–3) (**Supplementary Table S5A-C**). **C**) Endo H-treated and untreated nMPO displayed indistinguishable secondary structure profiles based on i) CD profiling (**Supplementary Table S13A-B**) and ii) MD-based predictions. **D**) Relative to native nMPO (WT), Endo H-treated MPO (P1) showed i) higher thermal stability as determined by CD_208 nm_ (arrows indicate initial melting temperatures) and ii) greater global polypeptide accessibility based on MD data (**Supplementary Figure S8-S9**). Key: a–e and i–ii indicate positions of the MPO glycosylation sites and key catalytic residues, respectively. For panel Bii, Ci, and Di: ^Data from Endo H only controls were subtracted from Endo H-treated nMPO data to enable comparison to untreated nMPO. For all panels: Data plotted as mean ± SD, ns, not significant (*p* ≥ 0.05), \**p* < 0.05, \*\**p* < 0.01, \*\*\**p* < 0.005, \*\*\*\**p* < 0.001, \*\*\*\*\**p* < 0.00005.

Endo H-treatment of nMPO under native conditions produced an Asn355- and Asn391-GlcNAc-rich glycophenotype mimicking the enriched glycoforms of Se/Pl-MPO, **Figure 4Bi**. Biochemical characterisation validated that the oligomannose-rich Asn355- and Asn391-glycans were fully converted to GlcNAcβAsn while the processed *N*-glycans in other positions were Endo H-insensitive, **Supplementary Figure S7**. The Endo H-treated nMPO exhibited higher chlorination and oxidation activity than untreated nMPO as established using three different activity assays (3.4–5.8x, all *p* < 0.005), **Figure 4Bii** and **Supplementary Table S5A-C**.

The molecular basis of the intriguing glycoform-dependent MPO activity was investigated using CD and MD simulations of relevant MPO glycoforms (WT and P1–P3), **Supplementary Table S4A-B**. Endo H-treated nMPO (P1) and untreated nMPO (WT) displayed indistinguishable secondary structure profiles rich in helical content as reported (Banerjee et al., 2011; Paumann-Page et al., 2013) based on CD data and predictions based on MD data, **Figure 4C** and **Supplementary Table S13A-B**. Notably, however, the Endo H-treated nMPO showed a higher initial melting temperature (72°C) than untreated nMPO (60°C) as determined by CD_208 nm_, **Figure 4Di** and **Supplementary Table S13C**. Both glycoforms completed their transition by 88°C, though about half the helical structure remained at this temperature. Thus, our data suggest that Endo H-treated nMPO displays an enhanced thermal stability relative to native nMPO. MD indicated that the relatively higher thermal stability of Endo H-treated nMPO was accompanied by a significantly higher global polypeptide accessibility relative to nMPO, **Figure 4Dii** and **Supplementary Figure S8-S9**. Similar accessibility gains distributed throughout the polypeptide chains, but with particular “hot-spots” of dramatically enhanced accessibility of residues C-terminal to Asn355 and Asn391 (labelled b–c) and distal to the heme and active site residues (i–ii, His261/Arg405), were observed for P2 (mimicking the Se/Pl-MPO-enriched glycoforms). The P3 control glycoform carrying semi-truncated *N*-glycans at Asn355 and Asn391 (M2–M3) did not show elevated accessibility relative to nMPO suggesting that minimal glycosylation at Asn355 and Asn391 is required to allosterically impact the global MPO structure and augment activity. In support, prolonged contacts were consistently observed between the β1,4-GlcNAc and β1,4-Man of the trimannosylchitobiose core (<u>Manβ1,4GlcNAcβ1,</u>4GlcNAcβAsn) and the polypeptide region immediately C-terminal to Asn355 and Asn391 during the MD simulation of glycans elongated beyond the GlcNAcβAsn at these positions (data not shown). Such glycan-protein contacts, which may modulate protein stability and structure, are particularly interesting given that Asn355 and Asn391 are positioned on flexible loops proximal to the active site. The MD data including RMSD-based mobility measurements, however, were insufficiently sensitive to unravel the molecular basis of the observed structure-activity relationships of glycosylated MPO in greater details, which, consequently, await future exploration, **Supplementary Figure S8D**.

### Hyper-Truncated Asn355-Glycosylation Enhances Ceruloplasmin-Mediated MPO Inhibition

The impact of glycosylation on MPO inhibition by ceruloplasmin, an endogenous inhibitor of MPO (Chapman et al., 2013), was explored by comparing the relative activity of Endo H-treated and untreated nMPO in the absence and presence of serum-derived ceruloplasmin, **Figure 5A**. The Endo H-treatment completely converted the oligomannose-rich Asn355 and Asn391 to GlcNAcβAsn-containing sites while sites containing processed glycans (e.g. Asn323) were Endo H-insensitive, **Figure 5B** and **Supplementary Figure S7**. The chlorination and relative oxidation rates of Endo H-treated and untreated nMPO were determined with and without equimolar ceruloplasmin using three activity assays, **Figure 5C** and **Supplementary Table S5D-F**. Activity assays 1 and 3 showed significant ceruloplasmin-mediated inhibition of Endo H-treated nMPO (p = 3.8 × 10^−4^ and 8.5 × 10^−5^) while untreated nMPO exhibited weak to no ceruloplasmin-mediated inhibition. For activity assay 2, an activity gain was experienced by untreated nMPO upon the addition of ceruloplasmin; this unexplained activity gain was quenched for the Endo H-treated nMPO. The molecular mechanisms underpinning the glycoform-dependent MPO inhibition by ceruloplasmin were explored using MD data of relevant Asn323-, Asn355- and Asn391-glycans modelled on a structure of the ceruloplasmin-MPO complex (PDBID, 4EJX), **Figure 5D**. Modelling demonstrated that only Asn355 is positioned directly in the MPO:ceruloplasmin interface, and, importantly, that Asn355-M6 and not Asn355-GlcNAc sterically clashes with ceruloplasmin thereby providing a molecular basis for the glycoform-dependent inhibition of MPO by ceruloplasmin.

**Figure 5.**
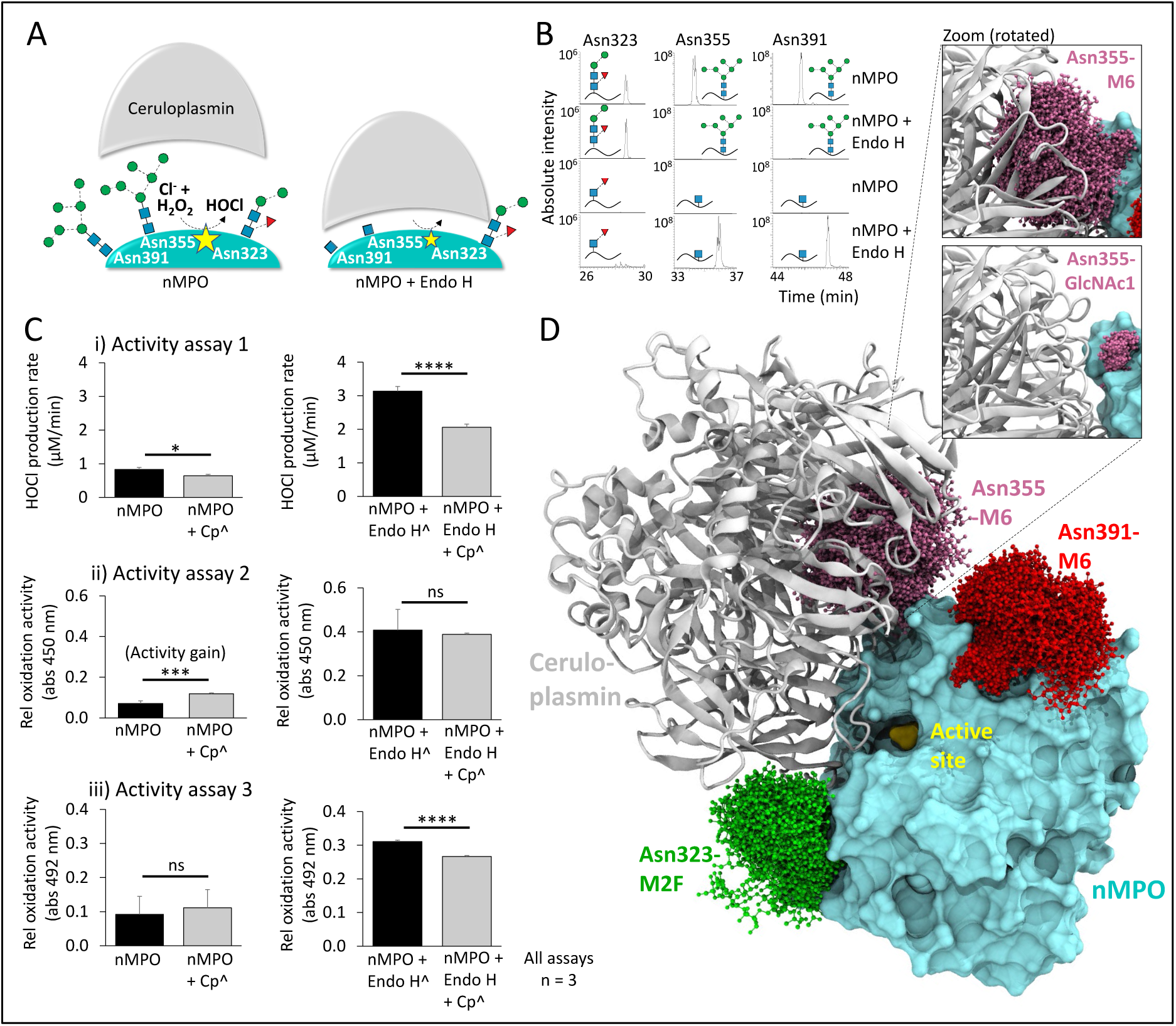
Hyper-truncated Asn355-glycans augment the ceruloplasmin-mediated MPO inhibition. **A**) Schematics of the glycoform-dependent ceruloplasmin-based inhibition of Endo H-treated and untreated nMPO. Common glycans decorating Asn323, Asn355, and Asn391 in proximity to the MPO-ceruloplasmin interface and the HOCl-producing active site (yellow star) are portrayed. **B**) Glycopeptide data (selected EICs) demonstrating complete conversion of Asn355-M6 and Asn391-M6 to GlcNAcβAsn upon Endo H-treatment (**Supplementary Figure S7**). The Endo H-insensitive Asn323-M2F was included as a control. **C**) The chlorination (i) and relative oxidation (ii-iii) levels of native nMPO (left graphs) and Endo H-treated nMPO (right) incubated with (grey bars) and without (black) serum ceruloplasmin (Cp) were determined in technical triplicates using activity assay 1–3, respectively (**Supplementary Table S5D-F**). ^Data from Endo H and Cp only controls were subtracted from Endo H-and Cp-treated nMPO data to enable comparison to untreated nMPO. **D**) MD data of Asn323-M2F (green), Asn355-M6 (magenta) and Asn391-M6 (red) modelled on a crystal structure of the ceruloplasmin-MPO complex (PDBID, 4EJX) demonstrated that Asn355-M6 and not Asn355-GlcNAc clashes with ceruloplasmin (see zoom). Data plotted as mean ± SD, ns, not significant (*p* ≥ 0.05), \**p* < 0.05, \*\**p* < 0.01, \*\*\**p* < 0.005, \*\*\*\**p* < 0.001.

### Conclusions

We have characterised the structure-biosynthesis-activity relationship of neutrophil granule MPO, and longitudinally profiled the spatiotemporal expression, protein processing, site-specific glycosylation and degranulation of MPO from maturing, mature, granule-separated and pathogen-activated neutrophils from healthy donors. Powered by multi-omics tools, our quantitative and longitudinal data add important molecular-level knowledge to our growing understanding of the complex MPO biology governing many innate immune processes central to human health and disease (Nauseef, 2014b; Winterbourn et al., 2016).

Complementary mass spectrometry and novel mass photometry approaches helped unravel the molecular complexity displayed by the heavily glycosylated and post-translationally modified nMPO. Critically, nMPO is site-specifically modified by unique glycans that are rarely reported in human glycobiology, but indeed are common to neutrophils including the hyper-truncated paucimannosidic- and chitobiose core-type *N*-glycans (Tjondro et al., 2019; Ugonotti et al., 2020), as well as elaborate oxidation, chlorination and polypeptide truncation variants. The structural glycan data presented herein align with and significantly expand on the existing knowledge base (Ravnsborg et al., 2010; Reiding et al., 2019; Van Antwerpen et al., 2010) by providing not only details of the glycan isomers but also the molecular mechanisms contributing to the distinctive site-specific MPO glycosylation. Several protein factors including the spatial environment and dimerisation status were found to affect the local *N*-glycan processing producing a position-specific glycan-code, which, intriguingly, was found to impact both the MPO structure and the activity and inhibition potential of the enzyme. The rarely reported GlcNAcβAsn-type glycans were elevated at strategic sites that are important for the activity and ceruloplasmin-mediated inhibition of MPO within mobile compartments of neutrophils. Taken together, our data suggest that neutrophils dynamically produce, process, package, store, and - upon activation, release a repertoire of related MPO glycoforms displaying a continuum of different activity and inhibition profiles. The strategically positioned Asn355 glycosylation site carrying both hyper-truncated and elongated *N*-glycans were found to be particularly important for MPO function, a finding that may guide future glycoengineering efforts aiming to generate therapeutically relevant recombinant MPO products with tuneable activity and inhibition potential tailored to specific biomedical applications involving persistent and severe pathogen infections.

In conclusion, this study has provided new molecular-level insights into the intriguingly complex sugar code of MPO of importance to fundamental neutrophil glycobiology and MPO-mediated immune processes central to human health and disease.

## Supporting information

This study contains supporting information. Three supporting files have been provided: 1) Supplementary Information containing Extended Methods and Supplementary Figure S1–S9 (PDF), 2) Supplementary Tables containing Supplementary Table S1–S13 (Microsoft Excel), and 3) Supplementary Data containing Supplementary Data S1–S3 (PDF).

The LC-MS/MS raw data files are available via ProteomeXchange with identifier PXD021131. Username, Password: nOuGDU6Q.

## Acknowledgments and funding sources

HCT was supported by an Australian Cystic Fibrosis postgraduate studentship award and an international Macquarie University Research Excellence Scholarship (iMQRES). SC was supported by an iMQRES. JU was supported by a Macquarie University Research Excellence Scholarship (MQRES). RK was supported by an Early Career Fellowship from the Cancer Institute NSW. MTA was supported by a Macquarie University Safety Net Grant.

The late Professor Niels Borregaard is thanked for valuable ideas and discussions seeding this research.

## Author contributions

Conception of research, HCT, MTA; Experimental design, HCT, RK, IL, RJW, JB, AKB, WBS, MTA; Experimental execution, HCT, JU, RK, SC, IL, SC, FS, HH, BLP, VV, RD, OCG, AR, WBS; Data analysis and interpretation: HCT, JU, RK, SC, IL, SC, FS, HH, BLP, VV, RD, OCG, AR, RJW, WBS, MTA; Manuscript writing and editing: HCT, JU, RK, SC, OCG, AR, JB, AKB, WBS, MTA.

## Author declarations

No conflict of interest has been declared.

## Abbreviations

AUC: area-under-the-curve
Az granule: azurophilic granule
Az-MPO: azurophilic granule-resident MPO
BN: band neutrophil
CD: circular dichroism
CytB/I: cytochalasin B and ionomycin
Dg-MPO: degranulated MPO
EIC: extracted ion chromatogram
Endo H: endoglycosidase H
ER: endoplasmic reticulum
Fuc (F): α-L-fucose
Ge granule: gelatinase granule
Ge-MPO: gelatinase granule-resident MPO
GlcNAc: *N*-acetyl-β-D-glucosamine
H_2_O_2_: hydrogen peroxide
HOCl: hypochlorous acid
LC-MS/MS: liquid chromatography tandem mass spectrometry
LDH: lactate dehydrogenase
Man: α/β-D-mannose
MD: molecular dynamics
MM: metamyelocyte
MOI: multiplicity-of-infection
MPO: myeloperoxidase
nMPO: neutrophil-derived myeloperoxidase (unfractionated)
PDB: Protein Data Bank
PM: promyelocyte
PMN: polymorphonuclear cell (neutrophil)
PMN-nMPO: myeloperoxidase from derived from resting (circulating) neutrophils
RMSD: root mean squared deviation
RMSF: root mean squared fluctuation
SD: standard deviation
Se/Pl: secretory vesicle and plasma membrane fraction
Se/Pl-MPO: secretory vesicle/plasma membrane-resident MPO
SN: maturing neutrophil with segmented nuclei
Sp granule: specific granule
Sp-MPO: specific granule-resident MPO
TMB: 3,3’,5,5’-tetramethylbenzidine.

## References

Agner, K. (1958). Crystalline myeloperoxidase. Acta Chemica Scandinavica 12, 89–94.

Aratani, Y., Kura, F., Watanabe, H., Akagawa, H., Takano, Y., Ishida-Okawara, A., Suzuki, K., Maeda, N., and Koyama, H. (2006). Contribution of the myeloperoxidase-dependent oxidative system to host defence against Cryptococcus neoformans. Journal of Medical Microbiology 55, 1291–1299.

Ashwood, C., Lin, C.-H., Thaysen-Andersen, M., and Packer, N.H. (2018). Discrimination of Isomers of Released N- and O-Glycans Using Diagnostic Product Ions in Negative Ion PGC-LC-ESI-MS/MS. Journal of The American Society for Mass Spectrometry 29, 1194–1209.

Banerjee, S., Stampler, J., Furtmuller, P.G., and Obinger, C. (2011). Conformational and thermal stability of mature dimeric human myeloperoxidase and a recombinant monomeric form from CHO cells. Biochim Biophys Acta 1814, 375–387.

Bjornsdottir, H., Welin, A., Dahlgren, C., Karlsson, A., and Bylund, J. (2016). Quantification of heterotypic granule fusion in human neutrophils by imaging flow cytometry. Data Brief 6, 386–393.

Blair-Johnson, M., Fiedler, T., and Fenna, R. (2001). Human Myeloperoxidase:? Structure of a Cyanide Complex and Its Interaction with Bromide and Thiocyanate Substrates at 1.9 Å Resolution. Biochemistry 40, 13990–13997.

Borregaard, N., and Cowland, J.B. (1997). Granules of the Human Neutrophilic Polymorphonuclear Leukocyte. Blood 89, 3503.

Borregaard, N., Sørensen, O.E., and Theilgaard-Mönch, K. (2007). Neutrophil granules: a library of innate immunity proteins. Trends in Immunology 28, 340–345.

Carpena, X., Vidossich, P., Schroettner, K., Calisto, B.M., Banerjee, S., Stampler, J., Soudi, M., Furtmüller, P.G., Rovira, C., Fita, I., et al. (2009). Essential role of proximal histidine-asparagine interaction in mammalian peroxidases. The Journal of biological chemistry 284, 25929–25937.

Caval, T., Heck, A.J.R., and Reiding, K.R. (2020). Meta-heterogeneity: evaluating and describing the diversity in glycosylation between sites on the same glycoprotein. Mol Cell Proteomics.

Chapman, A.L., Mocatta, T.J., Shiva, S., Seidel, A., Chen, B., Khalilova, I., Paumann-Page, M.E., Jameson, G.N., Winterbourn, C.C., and Kettle, A.J. (2013). Ceruloplasmin is an endogenous inhibitor of myeloperoxidase. J Biol Chem 288, 6465–6477.

Cheng, D., Talib, J., Stanley, C.P., Rashid, I., Michaelsson, E., Lindstedt, E.L., Croft, K.D., Kettle, A.J., Maghzal, G.J., and Stocker, R. (2019). Inhibition of MPO (Myeloperoxidase) Attenuates Endothelial Dysfunction in Mouse Models of Vascular Inflammation and Atherosclerosis. Arteriosclerosis, thrombosis, and vascular biology, Atvbaha119312725.

Clemmensen, S.N., Udby, L., and Borregaard, N. (2014). Subcellular Fractionation of Human Neutrophils and Analysis of Subcellular Markers. In Neutrophil Methods and Protocols, M.T. Quinn, and F.R. DeLeo, eds. (Totowa, NJ: Humana Press), pp. 53–76.

Cowland, J.B., and Borregaard, N. (2016). Granulopoiesis and granules of human neutrophils. Immunol Rev 273, 11–28.

Cox, J., and Mann, M. (2008). MaxQuant enables high peptide identification rates, individualized p.p.b.-range mass accuracies and proteome-wide protein quantification. Nat Biotechnol 26, 1367–1372.

Damme, M., Morelle, W., Schmidt, B., Andersson, C., Fogh, J., Michalski, J.C., and Lubke, T. (2010). Impaired lysosomal trimming of N-linked oligosaccharides leads to hyperglycosylation of native lysosomal proteins in mice with alpha-mannosidosis. Mol Cell Biol 30, 273–283.

Davies, M.J., Hawkins, C.L., Pattison, D.I., and Rees, M.D. (2008). Mammalian heme peroxidases: From molecular mechanisms to health implications. Antioxid Redox Signal 10, 1199–1234.

Delporte, C., Zouaoui Boudjeltia, K., Furtmueller, P.G., Maki, R.A., Dieu, M., Noyon, C., Soudi, M., Dufour, D., Coremans, C., Nuyens, V., et al. (2018). Myeloperoxidase-catalyzed oxidation of cyanide to cyanate: A potential carbamylation route involved in the formation of atherosclerotic plaques? Journal of Biological Chemistry.

Dypbukt, J.M., Bishop, C., Brooks, W.M., Thong, B., Eriksson, H., and Kettle, A.J. (2005). A sensitive and selective assay for chloramine production by myeloperoxidase. Free Radical Biology and Medicine 39, 1468–1477.

Feuk-Lagerstedt, E., Movitz, C., Pellme, S., Dahlgren, C., and Karlsson, A. (2007). Lipid raft proteome of the human neutrophil azurophil granule. Proteomics 7, 194–205.

Fiedler, T.J., Davey, C.A., and Fenna, R.E. (2000). X-ray Crystal Structure and Characterization of Halide-binding Sites of Human Myeloperoxidase at 1.8 Å Resolution. Journal of Biological Chemistry 275, 11964–11971.

Gault, J., Donlan, J.A.C., Liko, I., Hopper, J.T.S., Gupta, K., Housden, N.G., Struwe, W.B., Marty, M.T., Mize, T., Bechara, C., et al. (2016). High-resolution mass spectrometry of small molecules bound to membrane proteins. Nature Methods 13, 333.

Gorudko, I.V., Grigorieva, D.V., Sokolov, A.V., Shamova, E.V., Kostevich, V.A., Kudryavtsev, I.V., Syromiatnikova, E.D., Vasilyev, V.B., Cherenkevich, S.N., and Panasenko, O.M. (2018). Neutrophil activation in response to monomeric myeloperoxidase. Biochem Cell Biol 96, 592–601.

Gray, E., Thomas, T.L., Betmouni, S., Scolding, N., and Love, S. (2008). Elevated activity and microglial expression of myeloperoxidase in demyelinated cerebral cortex in multiple sclerosis. Brain pathology (Zurich, Switzerland) 18, 86–95.

Grishkovskaya, I., Paumann-Page, M., Tscheliessnig, R., Stampler, J., Hofbauer, S., Soudi, M., Sevcnikar, B., Oostenbrink, C., Furtmuller, P.G., Djinovic-Carugo, K., et al. (2017). Structure of human promyeloperoxidase (proMPO) and the role of the propeptide in processing and maturation. J Biol Chem 292, 8244–8261.

Harrison, J.E., and Schultz, J. (1976). Studies on the chlorinating activity of myeloperoxidase. Journal of Biological Chemistry 251, 1371–1374.

Haussermann, K., Young, G., Kukura, P., and Dietz, H. (2019). Dissecting FOXP2 Oligomerization and DNA Binding. Angew Chem Int Ed Engl 58, 7662–7667.

Hinneburg, H., Chatterjee, S., Schirmeister, F., Nguyen-Khuong, T., Packer, N.H., Rapp, E., and Thaysen-Andersen, M. (2019). Post-Column Make-Up Flow (PCMF) Enhances the Performance of Capillary-Flow PGC-LC-MS/MS-Based Glycomics. Analytical chemistry 91, 4559–4567.

Hoogendijk, A.J., Pourfarzad, F., Aarts, C.E.M., Tool, A.T.J., Hiemstra, I.H., Grassi, L., Frontini, M., Meijer, A.B., van den Biggelaar, M., and Kuijpers, T.W. (2019). Dynamic Transcriptome-Proteome Correlation Networks Reveal Human Myeloid Differentiation and Neutrophil-Specific Programming. Cell Rep 29, 2505–2519 e2504.

Hubber, S.J., and Thornton, J.M. (1993). NACCESS computer program. Department of Biochemistry and Molecular Biology, University College, London.

Jensen, P.H., Karlsson, N.G., Kolarich, D., and Packer, N.H. (2012). Structural analysis of N- and O-glycans released from glycoproteins. Nature Protocols 7, 1299.

Kawahara, R., Ortega, F., Rosa-Fernandes, L., Guimarães, V., Quina, D., Nahas, W., Schwämmle, V., Srougi, M., Leite, K.R.M., Thaysen-Andersen, M., et al. (2018). Distinct urinary glycoprotein signatures in prostate cancer patients. Oncotarget 9, 33077–33097.

Khalilova, I.S., Dickerhof, N., Mocatta, T.J., Bhagra, C.J., McClean, D.R., Obinger, C., and Kettle, A.J. (2018). A myeloperoxidase precursor, pro-myeloperoxidase, is present in human plasma and elevated in cardiovascular disease patients. PLoS One 13, e0192952.

Kim, H.J., Wei, Y., Wojtkiewicz, G.R., Lee, J.Y., Moskowitz, M.A., and Chen, J.W. (2018). Reducing myeloperoxidase activity decreases inflammation and increases cellular protection in ischemic stroke. Journal of Cerebral Blood Flow & Metabolism, 0271678×18771978.

Klebanoff, S.J. (1968). Myeloperoxidase-Halide-Hydrogen Peroxide Antibacterial System. Journal of Bacteriology 95, 2131.

Klebanoff, S.J. (2005). Myeloperoxidase: Friend and foe. J Leukocyte Biol 77, 598–625.

Lehrer, R.I., and Cline, M.J. (1969). Leukocyte myeloperoxidase deficiency and disseminated candidiasis: the role of myeloperoxidase in resistance to Candida infection. J Clin Invest 48, 1478–1488.

Loke, I., Ostergaard, O., Heegaard, N.H.H., Packer, N.H., and Thaysen-Andersen, M. (2017). Paucimannose-Rich N-glycosylation of Spatiotemporally Regulated Human Neutrophil Elastase Modulates Its Immune Functions. Mol Cell Proteomics 16, 1507–1527.

Loke, I., Packer, N.H., and Thaysen-Andersen, M. (2015). Complementary LC-MS/MS-Based N-Glycan, N-Glycopeptide, and Intact N-Glycoprotein Profiling Reveals Unconventional Asn71-Glycosylation of Human Neutrophil Cathepsin G. Biomolecules 5, 1832–1854.

Marty, M.T., Baldwin, A.J., Marklund, E.G., Hochberg, G.K.A., Benesch, J.L.P., and Robinson, C.V. (2015). Bayesian Deconvolution of Mass and Ion Mobility Spectra: From Binary Interactions to Polydisperse Ensembles. Analytical chemistry 87, 4370–4376.

Nauseef, W.M. (1986). Myeloperoxidase biosynthesis by a human promyelocytic leukemia cell line: insight into myeloperoxidase deficiency. Blood 67, 865.

Nauseef, W.M. (1987). Posttranslational processing of a human myeloid lysosomal protein, myeloperoxidase. Blood 70, 1143.

Nauseef, W.M. (2014a). Isolation of Human Neutrophils from Venous Blood. In Neutrophil Methods and Protocols, M.T. Quinn, and F.R. DeLeo, eds. (Totowa, NJ: Humana Press), pp. 13–18.

Nauseef, W.M. (2014b). Myeloperoxidase in human neutrophil host defence. Cell Microbiol 16, 1146–1155.

Nauseef, W.M. (2018). Biosynthesis of human myeloperoxidase. Archives of Biochemistry and Biophysics 642, 1–9.

Nauseef, W.M., Cogley, M., Bock, S., and Petrides, P.E. (1998). Pattern of inheritance in hereditary myeloperoxidase deficiency associated with the R569W missense mutation. J Leukoc Biol 63, 264–269.

Nauseef, W.M., McCormick, S., and Yi, H. (1992). Roles of heme insertion and the mannose-6-phosphate receptor in processing of the human myeloid lysosomal enzyme, myeloperoxidase. Blood 80, 2622–2633.

Paumann-Page, M., Furtmuller, P.G., Hofbauer, S., Paton, L.N., Obinger, C., and Kettle, A.J. (2013). Inactivation of human myeloperoxidase by hydrogen peroxide. Arch Biochem Biophys 539, 51–62.

Ravnsborg, T., Houen, G., and Hojrup, P. (2010). The glycosylation of myeloperoxidase. Biochim Biophys Acta 1804, 2046–2053.

Reiding, K.R., Franc, V., Huitema, M.G., Brouwer, E., Heeringa, P., and Heck, A.J.R. (2019). Neutrophil myeloperoxidase harbors distinct site-specific peculiarities in its glycosylation. Journal of Biological Chemistry.

Rørvig, S., Østergaard, O., Heegaard, N.H.H., and Borregaard, N. (2013). Proteome profiling of human neutrophil granule subsets, secretory vesicles, and cell membrane: correlation with transcriptome profiling of neutrophil precursors. J Leukocyte Biol 94, 711–721.

Samygina, V.R., Sokolov, A.V., Bourenkov, G., Petoukhov, M.V., Pulina, M.O., Zakharova, E.T., Vasilyev, V.B., Bartunik, H., and Svergun, D.I. (2013). Ceruloplasmin: macromolecular assemblies with iron-containing acute phase proteins. PloS one 8, e67145–e67145.

Silvescu, C.I., and Sackstein, R. (2014). G-CSF induces membrane expression of a myeloperoxidase glycovariant that operates as an E-selectin ligand on human myeloid cells. Proc Natl Acad Sci U S A 111, 10696–10701.

Soltermann, F., Foley, E., Pagnoni, V., Galpin, M., Benesch, J., Kukura, P., and Struwe, W. (2020). Quantifying protein-protein interactions by molecular counting with mass photometry. Angew Chem Int Ed Engl.

Stamp, L.K., Khalilova, I., Tarr, J.M., Senthilmohan, R., Turner, R., Haigh, R.C., Winyard, P.G., and Kettle, A.J. (2012). Myeloperoxidase and oxidative stress in rheumatoid arthritis. Rheumatology 51, 1796–1803.

Takeuchi, K., Umeki, Y., Matsumoto, N., Yamamoto, K., Yoshida, M., Suzuki, K., and Aratani, Y. (2012). Severe neutrophil-mediated lung inflammation in myeloperoxidase-deficient mice exposed to zymosan. Inflammation Research 61, 197–205.

Thaysen-Andersen, M., and Packer, N.H. (2012). Site-specific glycoproteomics confirms that protein structure dictates formation of N-glycan type, core fucosylation and branching. Glycobiology 22, 1440–1452.

Thaysen-Andersen, M., Packer, N.H., and Schulz, B.L. (2016). Maturing Glycoproteomics Technologies Provide Unique Structural Insights into the N-glycoproteome and Its Regulation in Health and Disease. Mol Cell Proteomics 15, 1773–1790.

Thaysen-Andersen, M., Venkatakrishnan, V., Loke, I., Laurini, C., Diestel, S., Parker, B.L., and Packer, N.H. (2015). Human Neutrophils Secrete Bioactive Paucimannosidic Proteins from Azurophilic Granules into Pathogen-Infected Sputum. The Journal of Biological Chemistry 290, 8789–8802.

Tjondro, H.C., Loke, I., Chatterjee, S., and Thaysen-Andersen, M. (2019). Human protein paucimannosylation: cues from the eukaryotic kingdoms. Biol Rev Camb Philos Soc 94, 2068–2100.

Ugonotti, J., Chatterjee, S., and Thaysen-Andersen, M. (2020). Structural and functional diversity of neutrophil glycosylation in innate immunity and related disorders. Molecular aspects of medicine, 100882.

Van Antwerpen, P., Slomianny, M.C., Boudjeltia, K.Z., Delporte, C., Faid, V., Calay, D., Rousseau, A., Moguilevsky, N., Raes, M., Vanhamme, L., et al. (2010). Glycosylation pattern of mature dimeric leukocyte and recombinant monomeric myeloperoxidase: glycosylation is required for optimal enzymatic activity. J Biol Chem 285, 16351–16359.

Vanhamme, L., Zouaoui Boudjeltia, K., Van Antwerpen, P., and Delporte, C. (2018). The other myeloperoxidase: Emerging functions. Archives of Biochemistry and Biophysics 649, 1–14.

Varki, A., Cummings, R.D., Aebi, M., Packer, N.H., Seeberger, P.H., Esko, J.D., Stanley, P., Hart, G., Darvill, A., Kinoshita, T., et al. (2015). Symbol Nomenclature for Graphical Representations of Glycans. Glycobiology 25, 1323–1324.

Venkatakrishnan, V., Dieckmann, R., Loke, I., Tjondro, H., Chatterjee, S., Bylund, J., Thaysen-Andersen, M., Karlsson, N.G., and Karlsson-Bengtsson, A. (2020). Glycan analysis of human neutrophil granules implicates a maturation-dependent glycosylation machinery. J Biol Chem.

Wang, J., Li, J.N., Cui, Z., and Zhao, M.H. (2018). Deglycosylation influences the oxidation activity and antigenicity of myeloperoxidase. Nephrology (Carlton) 23, 46–52.

Winterbourn, C.C., Kettle, A.J., and Hampton, M.B. (2016). Reactive Oxygen Species and Neutrophil Function. Annu Rev Biochem 85, 765–792.

Young, G., Hundt, N., Cole, D., Fineberg, A., Andrecka, J., Tyler, A., Olerinyova, A., Ansari, A., Marklund, E.G., Collier, M.P., et al. (2018). Quantitative mass imaging of single biological macromolecules. Science 360, 423–427.

Yu, J.T., Li, J.N., Wang, J., Jia, X.Y., Cui, Z., and Zhao, M.H. (2017). Deglycosylation of myeloperoxidase uncovers its novel antigenicity. Kidney Int 91, 1410–1419.

