## Supplementary Figures (S1-S9) and Extended Methods for "Hyper-Truncated Glycans Augment the Activity of Neutrophil Granule Myeloperoxidase"

for

[1]*Contributed equally*

**Running title**: The Sugar Code of Neutrophil Myeloperoxidase

***Corresponding author**:

Dr Morten Thaysen-Andersen, PhD

Department of Molecular Sciences

Macquarie University

NSW-2109, Sydney, Australia

Ph: +61 2 9850 7487

#### List of Content of SI


**Abbreviations used in SI** **S3**

**Extended Methods** **S4-S22**

**Supplementary Figures** **S23-S37**

Supplementary Figure S1: Structural characterisation of signature *N*-glycans released from nMPO

Supplementary Figure S2: Sequence coverage of the α- and β-chains of nMPO

Supplementary Figure S3: Structural characterisation of prominent *N*-glycopeptides from nMPO

Supplementary Figure S4: Characterisation of monoprotomeric (αβ) and diprotomeric (ααββ) nMPO

Supplementary Figure S5: Glycoprofiling of MPO from maturing, resting, and activated neutrophils

Supplementary Figure S6: MPO-centric sequence alignments

Supplementary Figure S7: Biochemical validation of native nMPO treated with Endo H

Supplementary Figure S8: Polypeptide accessibility and RMSD of WT and P1-P3 (MD data)

Supplementary Figure S9: Depiction of protein accessibility differences of WT and P1-P3 (MD data)

**Supplementary Tables *(provided as a separate excel file)***

Supplementary Table S1: Overview of donors, neutrophil/granule isolation methods and assays

Supplementary Table S2: Overview of LC-MS/MS (glyco)peptide profiling setting and strategies

Supplementary Table S3: Top down and native MS profiling of intact nMPO

Supplementary Table S4: Molecular dynamics simulations of MPO variants

Supplementary Table S5: MPO activity of Endo H-treated and untreated nMPO

Supplementary Table S6: Distribution and fine structures of the *N*-glycans decorating nMPO

Supplementary Table S7: Site-specific *N*-glycoprofiling and glycan processing features of nMPO

Supplementary Table S8: Asn and glycan accessibility of monoprotomeric and diprotomeric MPO

Supplementary Table S9: Site-specific *N*-glycoprofiling of monoprotomeric and diprotomeric nMPO

Supplementary Table S10: Glycoprofiling of MPO of maturing, mature, and activated neutrophils

Supplementary Table S11: Relative MPO chlorination and oxidation activity of two donors

Supplementary Table S12: Relative MPO activity of four donors based on the Tyr chlorination level

Supplementary Table S13: CD profiling and thermal stability of Endo H-treated and untreated nMPO

**Supplementary Data *(provided as a separate PDF)***

Supplementary Data S1: CID-MS/MS spectral evidence of nMPO *N*-glycans

Supplementary Data S2: HCD-MS/MS spectral evidence of non-glycan PTMs of nMPO peptides

Supplementary Data S3: HCD-MS/MS spectral evidence of nMPO *N*-glycopeptides

**References used in SI** **S38-S40**

#### Abbreviations used in SI

3D, three-dimensional; ACN, acetonitrile; AGC, automatic gain control; ALP, alkaline phosphatase; AUC, area-under-the-curve; Az granule, azurophilic granule; Az-MPO, azurophilic granule-resident MPO; BN, band neutrophil; CD, circular dichroism; CID, collision-induced dissociation; CytB/I, cytochalasin B and ionomycin; Dg-MPO, degranulated MPO; DDA, data-dependent acquisition; DTT, dithiothreitol; EIC, extracted ion chromatogram; Endo H, endoglycosidase H; FA, formic acid; FDR, false discovery rate; Fuc (F), α-L-fucose; FWHM, full width at half maximum; Ge granule, gelatinase granule; Ge-MPO, gelatinase granule-resident MPO; GlcNAc, *N*-acetyl-β-D-glucosamine; $H_2O_2$, hydrogen peroxide; HCD, higher-energy collisional dissociation; Hex, *N*-acetyl-β-hexosaminidase; HOCl, hypochlorous acid; IAA, iodoacetamide; kDa, kilo Dalton; KRG buffer, Krebs-Ringer buffer with glucose; LC-MS/MS, liquid chromatography tandem mass spectrometry; LDH, lactate dehydrogenase; LF, lactoferrin; LFQ, label-free quantitation; Man, α/β-D-mannose; MD, molecular dynamics; MM, metamyelocyte; MMP-9, matrix metalloproteinase-9; MOI, multiplicity-of-infection; MPO, myeloperoxidase; NCE, normalised collision energy; nMPO, neutrophil-derived (unfractionated) myeloperoxidase; NRMSD, normalised root mean squared deviation; PBS, phosphate buffered saline; PDB, Protein Data Bank; PGC, porous graphitised carbon; PM, promyelocyte; PMN, polymorphonuclear cell (neutrophil); PSM, peptide-to-spectral match; RMSD, root mean squared deviation; RMSF, root mean squared fluctuation; SD, standard deviation; Se/Pl, secretory vesicle and plasma membrane fraction; Se/Pl-MPO, secretory vesicle/plasma membrane-resident MPO; SN, neutrophil with segmented nuclei; Sp granule, specific granule; Sp-MPO, specific granule-resident MPO; SPE, solid phase extraction; TMB, 3,3',5,5'-tetramethylbenzidine.

#### Extended Methods

This section elaborates on the experimental procedures briefly described in the main text.

*Origin, purification, structural integrity, and activity of neutrophil MPO (nMPO)*

Enzymatically active human myeloperoxidase (MPO, UniProtKB, P05164) derived from unfractionated resting neutrophils (nMPO) was isolated from pooled donor blood (product number 426-10, Lee BioSolutions Inc, MO, USA). The nMPO was isolated to greater than 95% purity as established by gel electrophoresis using preparation-scale ion exchange and concanavalin A-based chromatography (data not shown). Upon receiving nMPO, the purity of the product was confirmed using reductive SDS-PAGE, which, as expected, showed two characteristic bands corresponding to the MPO light chain (α-chain, ~12.5 kDa) and heavy chain (β-chain, ~65 kDa) after Coomassie Blue staining. No other significant protein bands were observed, e.g. **Supplementary Figure S7A**. In addition, the enzymatic activity of nMPO was validated using a 3,3',5,5'-tetramethylbenzidine (TMB) assay (described below). The concentration of nMPO was determined with a Bradford assays (Sigma-Aldrich, Australia) using bovine serum albumin (Sigma-Aldrich, Australia) as a protein standard in the concentration range 0–2.5 mg/mL. nMPO was aliquoted and stored in a soluble form at a concentration of 1 µg/µL in 50 mM aqueous sodium acetate, 100 mM aqueous sodium chloride, pH 6.0 at –20°C until use.

*Donors, neutrophil isolation, granule separation and pathogen-based activation of neutrophils*

<u>Donors</u>

Six 40 mL donor pools of buffy coats each obtained from four healthy individuals (6 donor pools x 4 individuals/pool x 10 mL buffy coat/individual) were provided by The Blood Center, Sahlgrenska University Hospital, Gothenburg, Sweden. These six donated buffy coat pools are hereafter referred to as "Donor a–f", see **Supplementary Table S1** for an overview of the donor samples, handling and assays. Further, for the experiments involving pathogen-based activation of neutrophils, three 20 mL non-pooled buffy coats obtained from three healthy individual were provided by The Blood Center, Sahlgrenska University Hospital, Gothenburg, Sweden. These non-pooled buffy coat samples are hereafter referred to as "Donor g–i".

Details of the donors, isolation procedures and sample handling of maturing neutrophils used to establish the protein and transcript profile of maturing neutrophils undergoing granulopoiesis based on data reinterrogation have been published (Hoogendijk et al., 2019).

<u>Neutrophil isolation</u>

Primary neutrophils were isolated from the buffy coat from Donor a-i as previously described (Clemmensen et al., 2014; Rørvig et al., 2013). Briefly, the erythrocytes were removed by dextran (1%, w/v) sedimentation. Monocytes and lymphocytes were removed by centrifugation at 400 x g, 4°C, 30 min on a Ficoll–Paque density gradient. The cells were then washed two times for 30 s in Krebs-Ringer buffer with 10 mM aqueous glucose (KRG buffer) to remove the remaining erythrocytes by hypotonic lysis. The neutrophils were then spun at 200 x g, 4°C, 10 min and resuspended in the KRG buffer. The total neutrophil count for each of the buffy coat pools was approximately $10^9$, the purity was greater than 95%, and the viability was greater than 99% as measured by a haematology cell counter (Sysmex, Sweden), trypan blue stain microscopy, and Annexin V and propidium iodide staining using flow cytometry (Thermo Fisher Scientific, Sweden), respectively. Aqueous diisopropyl fluorophosphate was added to a final concentration of 5 mM, and kept in the cell suspension for 5 min before excess protease inhibitor was removed by centrifugation at 200 x g, 4°C, 6 min.

Granule separation

Granules were separated from resting neutrophils isolated from Donor a-f. For this purpose, the plasma membranes of the isolated neutrophils were gently disrupted without compromising the integrity of the granule membranes using nitrogen cavitation employing scientific grade nitrogen (≥ 99.99% purity, v/v) in a Parr bomb. The released granules from Donor a-b neutrophils were crudely separated using a two-layered Percoll separation method while the released neutrophil granules from Donor c-f were subjected to a three-layered Percoll separation method facilitating high-resolution granule separation as described previously (Clemmensen et al., 2014; Nauseef, 2014).

The three-layered Percoll separation method, which enabled the isolation of the azurophilic (Az) granules, specific (Sp) granules, and gelatinase (Ge) granules and a combined fraction containing the secretory vesicles and the plasma membrane (Se/Pl), employed a density gradient generated by 1.12 g/mL, 1.09 g/mL, and 1.05 g/mL aqueous Percoll (catalogue number GE17-0891-01, GE Healthcare, Sweden). Separation was achieved by centrifugation of the released granules loaded on the Percoll density gradient at 37,000 x g, 4°C, 30 min. The relative distribution or activity of four proteins recognised as being robust compartment markers were determined for the resulting density fractions to identify the location of the four neutrophil compartments within the gradient. The enzyme activity of MPO was determined in order to identify the Az granules; the relative abundance of lactoferrin (LF) was determined to identify the Sp granules; the relative abundance of matrix metalloproteinase-9 (MMP-9) was determined to identify the Ge granules; and the enzyme activity of alkaline phosphatase (ALP) was determined to identify the Se/Pl fraction. The relative enzyme activities of MPO and ALP were determined as previously described (Feuk-Lagerstedt et al., 2007). The relative abundances of

LF and MMP-9 were assessed by immunoblotting experiments using rabbit polyclonal anti-LF antibodies (1:2500 dilution, Dako, Sweden) and rabbit polyclonal anti-MMP9 antibodies (1:2000 dilution, product number 444236, Calbiochem, Sweden), respectively. Density fractions containing the same marker signatures were combined.

The two-layered Percoll separation method, which facilitated the separation of the Az granules from a fraction containing both the Sp and Ge granules (called Sp/Ge) and a fraction containing the Se/Pl membrane, employed a density gradient generated by 1.05 g/mL and 1.12 g/mL aqueous Percoll (catalogue number GE17-0891-01, GE Healthcare, Sweden). Separation was achieved by centrifugation of the released granules loaded on the density gradient at 37,000 x g, 4°C, 30 min. The relative distribution and/or activity of complementary compartment markers were determined for the resulting density fractions to identify the location of the Az, Sp/Ge and Se/Pl fractions within the gradient. Specifically, the enzyme activity of MPO was determined to identify the Az granule; the relative abundance of LF was determined to identify the Sp/Ge granule fractions and the enzyme activity of ALP was determined to identify the Se/Pl fraction as described above. Density fractions containing the same granule marker signatures were combined.

Debris, Percoll and other insoluble components were removed from the granule fractions isolated using both the two- and three-layered Percoll separation methods by employing ultra-centrifugation at 100,000 x g, 4°C, 1 h, which physically separated the pelleted granules from the unwanted debris and Percoll in the bottom of the tubes (Clemmensen et al., 2014; Nauseef, 2014). The pelleted granules were resuspended in a homogenisation buffer consisting of 100 mM aqueous potassium chloride, 3 mM aqueous sodium chloride, 3.5 mM aqueous magnesium chloride, 10 mM aqueous 1,4-piperazinediethanesulfonic acid, pH 7.4. The granules were then lysed using a tip sonicator at 1/3 power (8 MHz) three times for 30 s on ice. The luminal (soluble) proteins were separated from the membrane proteins of each granule fraction by ultracentrifugation at 100,000 x g, 4°C, 90 min. Only the MPO-containing soluble protein (supernatant) fractions were investigated in this study. The supernatants were carefully collected, the protein concentrations determined using Bradford assays. The protein samples were stored at a concentration of 4–15 µg/µL at –20°C until being used for biochemical characterisation and various functional assays.

Neutrophil activation, MPO degranulation and determination of cell death

*Staphylococcus aureus* from the pathogenic LS1 lab strain kindly donated by Dr Tao Jin, Department of Rheumatology and Inflammation Research, Institute of Medicine, University of Gothenburg, Sweden was used for the pathogen-based activation of resting neutrophils. Stimulation was performed at an multiplicity-of-infection (MOI) of 1:5 (bacteria:neutrophil) to simulate low-level infection sufficient to trigger neutrophil activation (and MPO degranulation) while avoiding neutrophil cell death. Approximately $10^7$ neutrophils isolated from Donor g-i were inoculated with approximately 2 x $10^6$ *S. aureus* bacteria. The inoculate was separated into four tubes and subjected to different incubation periods at 37°C: 0 min, 30 min, 60 min and 120 min. For the 0 min incubation, the samples were briefly inoculated with *S. aureus*, but were not incubated for any significant length of time (approximately 2-3 min). As negative infection controls, the same number of isolated neutrophils ($10^7$) from the same donors (Donor g-i) were incubated under the same conditions, but without *S. aureus* inoculation for 0 min and 120 min. As a positive activation control, the same number of isolated neutrophils from the same donors were incubated with 5 µg/mL aqueous cytochalasin B for 5 min at 37°C followed by 0.5 µM aqueous ionomycin (hereafter collectively referred to as CytB/I) for 10 min at 37°C. As a positive control for neutrophil cell death, the same number of isolated neutrophils from the same donors were incubated with 1% (w/v) Triton-X 100, 37°C, 5 min.

To investigate the degranulated MPO (Dg-MPO), an MPO ELISA kit (catalogue number E-80PX, ICL LAB, OR, USA) was used as per the manufacturer's instructions to measure the level of Dg-MPO secreted into the extracellular space across various study conditions. For this purpose, the supernatant fractions of all neutrophil samples were collected after spinning the samples at 2,000 x g, 4°C, 15 min. An MPO standard from the ELISA kit was used to determine the concentration of Dg-MPO in the supernatant fractions. The level of neutrophil cell death was assessed by measuring the level of cytosolic lactate dehydrogenase (LDH) released into the same supernatant fractions using the Cytotoxicity Detection KitPLUS (catalogue number 4744934001, Sigma, Sweden) as per the manufacturer's instructions.

*Glycomics*

<u>Glycan release and clean-up</u>

For glycan profiling, nMPO was reduced using 10 mM aqueous dithiothreitol (DTT), 56°C, 45 min, and alkylated using a freshly prepared solution of 25 mM aqueous iodoacetamide (IAA), 25°C, 30 min in the dark (both final concentrations) and otherwise handled as previously described (Jensen et al., 2012). In short, isolated nMPO (20 µg/replicate) was blotted in triplicates on a primed PVDF membrane (Merck-Millipore, Australia) and stained with Direct Blue (Sigma-Aldrich, Australia). Protein spots were cut out and transferred to a flat bottom polypropylene 96-well plate (Corning Life Science, Australia), blocked with 1% (w/v) polyvinylpyrrolidone in 50% (v/v) aqueous methanol and washed with water. The *N*-glycans were exhaustively released from nMPO using 2 U recombinant *Elizabethkingia miricola* peptide-*N*-glycosidase F produced in *Escherichia coli* (Promega, Australia) per

20 µg nMPO in 10 µL water/well, 37°C, 16 h. Subsequently, the *N*-glycans were hydroxylated in 100 mM aqueous ammonium acetate, pH 5, 25°C, 1 h, and reduced to alditols in 1 M aqueous sodium borohydride in 50 mM aqueous potassium hydroxide, 50°C, 3 h. The reaction was quenched using glacial acetic acid. Desalting was performed using solid phase extraction (SPE) micro-columns packed with strong cation exchange resin (AG 50W-X8, Bio-Rad, Australia) on top of C18 discs serving to remove cationic salts and particulates, followed by porous graphitised carbon (PGC) SPE micro-columns to perform a final desalting of the *N*-glycans. For the final step, the retained *N*-glycans were eluted using 40% (v/v) aqueous acetonitrile (ACN) containing 0.1% (v/v) aqueous trifluoroacetic acid, dried and redissolved in 10 µL MilliQ water (Merck-Millipore, Australia) for LC-MS/MS. To assess for *O*-glycosylation, the de-*N*-glycosylated nMPO was subsequently treated with 1 M aqueous sodium borohydride in 50 mM aqueous potassium hydroxide, 50°C, 16 h and handled as described (Jensen et al., 2012). For all glycomics experiments, bovine fetuin (Sigma, Australia) was used as a standard glycoprotein to ensure efficient *N*- and *O*-glycan release and clean-up, and LC-MS/MS performance.

Profiling and data analysis of nMPO glycans

Glycans were separated and detected by PGC-LC-MS/MS performed on an LTQ Velos Pro ion trap mass spectrometer (Thermo Scientific, Australia) coupled to a Dionex Ultimate 3000 HPLC (Thermo Scientific, Australia). The *N*-glycan samples were loaded on a PGC-LC capillary column (Hypercarb KAPPA, 5 µm particle size, 250 Å pore size, 0.18 mm inner diameter x 100 mm length, Thermo Scientific, Australia) and separated using a linear multi-step gradient consisting of 2.6–64% (v/v) ACN in10 mM aqueous ammonium bicarbonate over 86 min at a constant flow rate of 3 µL/min and with a post-column make-up flow supplying pure methanol (Sigma-Aldrich, Australia) after the analytical column at a constant flow rate of 4 µL/min (Hinneburg et al., 2019). The MS1 acquisition range was *m/z* 500–2,000, the resolution was *m/z* 0.25 full width at half maximum (FWHM, measured at *m/z* 200), and the source voltage was +2.7 kV. Detection was performed in negative ionisation polarity mode. The automatic gain control (AGC) for the MS1 scans was 5 x $10^4$ with a maximum accumulation time of 50 ms. For MS/MS, the resolution was *m/z* 0.35 FWHM, the AGC was 2 x $10^4$ and the maximum accumulation time was 300 ms. Using a data-dependent acquisition (DDA) strategy, the six most abundant precursors in each MS1 full scan were selected for resonance activation (ion trap) collision-induced dissociation (CID)-MS/MS employing a 33% normalised collision energy (NCE). The *O*-glycomics method utilised similar LC-MS/MS settings, but the experimental details and resulting data have been omitted here since no *O*-glycans were detected for nMPO.

The *N*-glycan fine structures were manually characterised from the PGC-LC-MS/MS data using Xcalibur v2.2 (Thermo Scientific, Australia) based the monoisotopic precursor mass, the glycan fragmentation pattern arising from the negative-ion CID-MS/MS, and the relative and absolute PGC-LC retention time of each candidate glycan (Abrahams et al., 2018; Everest-Dass et al., 2013; Sethi et al., 2015; Stadlmann et al., 2008). RawMeat v2.1 (Vast Scientific, www.vastscientific.com) and GlycoMod (https://web.expasy.org/glycomod/) aided the glycan identification process. The biosynthetic relationship between the observed *N*-glycans served to further enhance the confidence of the reported structures. The relative abundances of the identified *N*-glycans were established based on the area-under-the-curve (AUC) measurements of extracted ion chromatograms (EICs) performed for all relevant charge states of the monoisotopic glycan precursors using Skyline 64-bit v20.1.0.76 (Ashwood et al., 2018; Loziuk et al., 2017).

*Glycoproteomics and glycopeptide profiling*

<u>Generation of glycopeptide mixtures</u>

Mixtures of peptides and glycopeptides were prepared from i) purified nMPO, ii) the separated monoprotomer (αβ) and diprotomer (ααββ) of nMPO, iii) endoglycosidase H- (Endo H-) treated and untreated nMPO, iv) separated neutrophil granules and v) supernatant fractions of pathogen-activated neutrophils.

i) Isolated nMPO (20 µg) was reduced and alkylated in technical triplicates as described above. The alkylation reactions were quenched using 30 mM aqueous DTT in 100 mM aqueous ammonium bicarbonate, 25$^{o}$C, 30 min. The proteins were then exhaustively digested using sequencing grade modified porcine trypsin (Promega, Australia) at a 1:30 enzyme:substrate ratio (w/w) in 100 mM aqueous ammonium bicarbonate, pH 8.4, 37$^{o}$C, 16 h. Digestions were stopped by acidification with 1% (v/v) aqueous formic acid (FA) (final concentration). The resulting peptide mixtures were desalted using C18-SPE micro-columns (Merck-Millipore, Australia). The desalted peptide mixtures were dried and redissolved in 10 µL 0.1% (v/v) aqueous FA for LC-MS/MS.

ii) The monoprotomeric (αβ) and diprotomeric (ααββ) forms of nMPO (20 µg protein/replicate) were separated in technical triplicates using non-reductive SDS-PAGE on a pre-cast 4–12% gradient gel (Invitrogen, Australia). Separations were performed using a constant potential of 120 V, 45 min, and was carried out on ice to prevent spontaneous dissociation of diprotomeric nMPO. The gels were stained using Coomassie Blue and the protein bands corresponding to the monoprotomeric (αβ) and diprotomeric (ααββ) nMPO were excised, in-gel digested at 37°C, 16 h using 33.3 ng/µL sequencing grade modified porcine trypsin (Promega, Australia). The resulting peptide mixtures were extracted using repeated washing cycles of ACN and water, and then desalted using C18-SPE micro-columns

(Merck-Millipore). The desalted peptide mixtures were dried and redissolved in 10 µL 0.1% (v/v) aqueous FA for LC-MS/MS.

iii) Reducing LDS-PAGE sample loading buffer (4x, pH 8.4, Invitrogen, Australia) and 50 mM aqueous DTT (final concentration) was added to 2 µg Endo H-treated and 2 µg untreated nMPO (see details of Endo H treatment below) and the mixtures were heated at 70°C, 10 min. The samples were then applied to SDS-PAGE on a pre-cast 4–12% gradient gel (Invitrogen, Australia). Following Coomassie Blue staining, the β-chains of nMPO (53-58 kDa) were excised, and in-gel digested at 37°C, 16 h using 33.3 ng/µL sequencing grade modified porcine trypsin (Promega, Australia). The resulting peptide mixtures were extracted using repeated washing cycles of ACN and water, and then desalted using C18-SPE micro-columns (Merck-Millipore, Australia). The desalted peptide mixtures were dried and redissolved in 10 µL 0.1% (v/v) aqueous FA for LC-MS/MS.

iv) Reducing LDS-PAGE sample loading buffer (4x, pH 8.4, Invitrogen, Australia) and 50 mM aqueous DTT (final concentration) was added to each of the isolated granule fractions (~300 µg total protein/replicate). The samples were heated to 95°C for 10 min before the granule protein extracts were introduced briefly into a pre-cast 4–12% gradient gel (Invitrogen, Australia) to form a single protein band on the top of the gel. This approach was chosen since the soluble protein fractions from the isolated granules contained residual Percoll and/or other agents incompatible with the downstream (glyco)proteomics workflow despite the multiple ultracentrifugation steps. The single protein band from each granule sample was excised and in-gel digested at 37°C, 16 h using 100 ng/µL sequencing grade modified porcine trypsin (Promega, Australia). The resulting peptide mixtures were extracted using repeated washing cycles of ACN and water, and then desalted using C18-SPE micro-columns (Merck-Millipore, Australia). The desalted peptide mixtures were dried and redissolved in 10 µL 0.1% (v/v) aqueous FA for LC-MS/MS.

v) Proteins in the supernatant fractions of activated neutrophils (and control samples) were precipitated using acetone at -20°C with overnight incubation. The pelleted proteins were then resuspended in 100 mM aqueous ammonium bicarbonate to a final protein concentration of 0.1 µg/mL. Proteins were then reduced using 10 mM aqueous DTT, 56°C, 45 min, and alkylated using 25 mM aqueous IAA, 25°C, 30 min in the dark (both final concentrations). The alkylation reactions were quenched using 30 mM aqueous DTT in 100 mM aqueous ammonium bicarbonate, 25°C, 30 min and the proteins were then exhaustively digested using sequencing-grade modified porcine trypsin (Promega, Australia) at a 1:30 enzyme:substrate ratio (w/w) in 100 mM aqueous ammonium bicarbonate, pH 8.4, 37°C, 16 h. Digestions were stopped by acidification with 1% (v/v) aqueous FA (final concentration). The resulting peptide mixtures were desalted using C18-SPE micro-columns

(Merck-Millipore, Australia). The desalted peptide mixtures were dried and redissolved in 10 µL 0.1% (v/v) aqueous FA for LC-MS/MS.

<u>LC-MS/MS (glyco)peptide profiling of neutrophil MPO from various origins and conditions</u>

The generated peptide mixtures were analysed using multiple high-resolution LC-MS/MS strategies as outlined below, see also **Supplementary Table S2** for overview and details of all LC-MS/MS-based glycopeptide profiling experiments. For all experiments, the peptides were detected using a Q-Exactive HF-X Hybrid Quadrupole-Orbitrap mass spectrometer (Thermo Scientific, Australia) coupled to either an UltiMate™ 3000 RSLCnano System (Thermo Scientific, Australia) or an Easy nLC1200 system (Thermo Scientific, Australia). The mass spectrometer was operated in positive ion polarity mode. The peptide mixtures were separated using reversed-phase C18 chromatography using custom-made nano-LC columns packed with either C18 HALO resin (2.7 µm particle size, 160 Å pore size, 0.1 mm inner diameter x 300 mm length, Halo, Advanced Materials Technology, Australia) or C18AQ resin (1.9 µm particle size, 120 Å pore size, 0.075 mm inner diameter x 500 mm length, Dr Maisch HPLC GmbH, Germany) or alternatively using a commercial PepMap100 C18 LC column (3 µm particle size, 100 Å pore size, 0.075 mm inner diameter x 250 mm length, Thermo Scientific, Australia). The peptide mixtures were loaded onto trap columns packed with either C18 HALO resin (2.7 µm particle size, 160 Å pore size, 0.1 mm inner diameter x 35 mm length, Halo, Advanced Materials Technology, Australia) or C18 PepMap (5 µm particle size, 100 Å pore size, 0.3 mm inner diameter x 5 mm length, Thermo Scientific, Australia) and then separated on the analytical columns (above) using different LC gradients at flow rates of 300–600 nL/min.

The MS1 acquisition range was *m/z* 350–1,650 or *m/z* 350–1,800 using a resolution of 60,000 measured at *m/z* 200. The MS1 AGC target was 3 x $10^6$ with a maximum injection time of 50 ms. The MS/MS AGC target was 1 x $10^5$ or 2 x $10^5$, the maximum injection time was 25–28 ms and the resolution was 15,000 measured at *m/z* 200. Employing DDA strategies, the 15–20 most abundant precursors in each MS1 scan were selected for MS/MS using higher-energy collisional dissociation (HCD) at 27%-30% NCE. Precursor isolation windows of *m/z* 1.2–1.4 were used. Already selected precursors were dynamically excluded for 20–30 s. Charge state unassigned precursors, singly charged precursors and precursors with charge states Z > 7–8 were also excluded for MS/MS.

The resulting HCD-MS/MS data were searched using multiple search strategies, see **Supplementary Table S2** for an overview of the search strategies and variables. The data were searched against either the canonical form of human MPO (UniProtKB, P05164) and/or the entire human proteome (UniProtKB, 20,397 reviewed entries, downloaded October 2018 or 20,364 reviewed entries, downloaded December 2019) using Byonic v3.6.0 (Protein Metrics Inc, CA, USA) and MaxQuant v1.6.0.1 or v1.6.10.43 (Cox and Mann, 2008). Raw data were used as input for both search engines. The searches employed tryptic, semi-tryptic or non-specific cleavage specificity with up to two missed cleavages allowed, fixed Cys carbamidomethylation and several glycan and non-glycan variable modifications including Met and Trp mono- and di-oxidation and Tyr mono- and di-chlorination. The glycopeptide data were searched against a comprehensive library of 309 mammalian *N*-glycans to which several non-conventional monosaccharide compositions including phosphorylated paucimannosidic and phosphorylated oligomannosidic-type *N*-glycans were manually added in Byonic. The precursor and product ion mass tolerance was 4.5–10 ppm and 20 ppm, respectively, and decoy and contaminant databases were included in the searches.

For the Byonic output, only (glyco)peptides identified with high scores (low false discovery rates, FDRs) were considered (PEP-2D scores < 0.001) (Kawahara et al., 2018). The Byonic-identified glycopeptides were further validated by manually interrogating the precursor envelope to confirm correct monoisotopic peak picking and charge state assignment, and by manually checking the HCD-MS/MS data and C18-LC elution time to ensure the presence of the expected product ions and similar LC retention time of related glycopeptides. The confidently identified glycopeptides were quantified based on AUC measurements from narrow EICs performed for all relevant charge states of the monoisotopic glycopeptide precursors (MS1 data) using Skyline 64-bit v20.1.0.76 (Ashwood et al., 2018; Loziuk et al., 2017) or manually using the Qual Browser in Xcalibur v2.2 (Thermo Scientific, Australia). For both the Skyline-based and manual EIC-based label-free quantitation (LFQ), a narrow 5 ppm precursor mass tolerance was applied, and data were manually inspected to ensure correct monoisotopic peak integration and consistent LC retention time across sample runs and an agreement with the glycopeptide identifications provided by Byonic.

For the MaxQuant output, the peptide-to-spectral matches (PSMs) and the protein identifications were filtered to FDRs below 0.01-0.03. The MaxQuant-based LFQ of the non-modified peptides and peptides modified with non-glycan modifications was based on the relative MS1 signal strength of confidently identified precursors (Cox and Mann, 2008). The proportion of proMPO of all MPO forms was determined for both the propeptide region and the hexapeptide based on the intensity of related peptide pairs arising from proMPO and maturely processed MPO.

The granulopoiesis data published by (Hoogendijk et al., 2019) were extracted from the ProteomeXchange consortium via the PRIDE partner (data accession number, PXD013785). The dataset consisted of LC-MS/MS (glyco)peptide data of maturing neutrophils isolated from four different maturation stages obtained by fluorescence-activated cell sorted myeloid progenitor cells derived from the bone marrow of four healthy donors including promyelocytes and myelocytes (collectively referred to as PMs), metamyelocytes (MMs), immature neutrophils with band-formed nuclei (BNs), and mature neutrophils with segmented nuclei (SNs). Circulating (mature) neutrophils (polymorphonuclear cells, PMNs) derived from blood from the same four donors were also investigated. Multiple LC-MS/MS datasets (technical replicates) were provided for each of these conditions. For our study, only a single technical replicate from each one of the four biological replicates from each maturation stage was downloaded via PRIDE and re-interrogated using Proteome Discoverer v2.4 (Thermo Scientific, Australia). Peptides and glycopeptides were identified with Byonic v3.6.0 using a human proteome database (UniProtKB, 20,416 entries, downloaded August 2019) and a custom-made neutrophil-focused *N*-glycan database (206 *N*-glycans) as the protein and glycan search space, respectively. For the granulopoiesis data sets, the quantification of (glyco)peptides was performed by Minora Feature Detector with the minimum trace length set to 5 and maximum delta retention time of isotope pattern multiplets of 0.2 min within the Proteome Discoverer v2.4 platform. For this purpose, chromatographic alignment of signals of interest was performed and the quantification based on a retention time match within a 1 min tolerance and an *m/z* match within a 10 ppm tolerance. Minimum S/N thresholds were enabled. Protein expression data and relative mRNA expression levels of MPO were manually retrieved (Hoogendijk et al., 2019).

*Characterisation of intact nMPO using native MS and mass photometry*

<u>Top down and native MS of intact nMPO</u>

Intact mass analyses were performed of i) intact α-chain of nMPO and ii) intact nMPO.

i) The nMPO was reduced using 10 mM aqueous DTT, 56°C, 45 min. The dissociated α- and β-chains of nMPO were then desalted using C4 SPE ZipTips (Millipore, Australia), dried and resuspended in 0.1% (v/v) aqueous FA. Approximately 1 µg (6–7 pmol) protein material was injected on a reversed-phase C4 LC column (Proteocol C4Q, 3 µm particle size, 300 Å pore size, 0.3 mm inner diameter × 100 mm length, SGE Analytical Science, Australia) kept at a constant temperature of 25°C and operated at a constant flow rate of 5 µL/min using an Agilent 1100 HPLC system connected directly to a high-resolution Agilent 6538 quadrupole-time-of-flight mass spectrometer (Agilent Technologies, Australia). A multi-step LC gradient was applied comprising the following linear steps: a) 100% solvent A containing 0.1% (v/v) aqueous FA for 5 min, b) 0–60% solvent B containing 99% ACN in 0.1% (v/v) aqueous FA ramp over 30 min, c) immediate increase to 100% solvent B, which was kept for 5 min and d) re-equilibration in 100% solvent A for 5 min. The mass spectrometer was operated in high-resolution (4 GHz) positive ion polarity mode using the following settings: The fragmentor potential was +200 V, the MS1 scan range was *m/z* 400–2,500, the nitrogen drying gas flow rate was 8 L/min at

300°C, the nitrogen nebuliser pressure was 10 psi, the capillary potential was +4.3 kV, and the skimmer potential was +65 V. Mass spectra were viewed, deconvoluted and analysed with MassHunter work station vB.06 (Agilent Technologies, Australia). The intact α-chain of nMPO was profiled by deconvoluting the multi-charged signals into the expected mass range for the MPO α-chain (10–20 kDa) using mass stepping of 0.5. Signals corresponding to the α-chain of nMPO were assigned based on accurate matches better than 2 Da (see **Supplementary Table S3A**) to the expected α-chain proteoforms predicted based on the obtained LC-MS/MS peptide data of nMPO, see **Supplementary Figure S2** and **Supplementary Data S2**. Average masses (neutral M, Da) were used to compare the experimental mass of the observed precursors (apex of signals) to the theoretical mass of the expected precursors.

ii) nMPO was desalted and buffer exchanged into 1 M aqueous ammonium acetate, pH 7.0 using Bio-Spin 6 Columns (BioRad, UK). Further, three washes in 1 M aqueous ammonium acetate were performed using 3,000 molecular weight cut-off centrifugal filters (Amicon Ultra-0.5, Merck, UK). The concentration of nMPO was measured using a UV-VIS spectrophotometer (DeNovix, DE, USA) and adjusted to a final concentration of 10 µM with 1 M aqueous ammonium acetate immediately prior to analysis. A modified Q-Exactive (Thermo Scientific, UK) was used for the native MS profiling of intact nMPO (Gault et al., 2016). Ions were generated by static nano-electrospray using gold-coated capillaries prepared in-house as described previously (Hernández and Robinson, 2007). The capillary voltage was +1.3 kV, the source was 200°C, the S-lens RF was 200, the maximum injection time was 50 ms, the HCD potential was off with in-source trapping enabled, and the desolvation potential was –10 V (positive ion polarity mode). MS1 data were acquired in the range *m/z* 1,000–15,000 with 10 microscans. Data were processed with Xcalibur v2.2 (Thermo Scientific, UK). Resulting spectra were deconvoluted with UniDec (Marty et al., 2015), and the signals were annotated using software produced in-house.

Mass photometry of intact nMPO

Intact nMPO was also analysed using mass photometry as described (Soltermann et al., 2020). The concentration of nMPO was measured using a UV-VIS spectrophotometer (DeNovix, DE, USA) and adjusted to a final concentration of 10 nM with phosphate buffered saline (PBS). After dilution, nMPO was equilibrated for 15 min prior to analysis. A clean coverslip was assembled for sample delivery using silicone gaskets (CultureWell™ gasket, 3 mm diameter x 1 mm depth, Grace Bio-Labs, OR, USA). For data acquisition, 36 µL sample was injected and data were collected for 60 s after a 10 s initial delay. Data processing and analysis were performed using custom software written in Python (Young et al., 2018).

*Sequence alignment*

The canonical FASTA sequences of human MPO (UniProtKB, P05164), eosinophil peroxidase (EPO, P11678), lactoperoxidase (LPO, P22079) and thyroid peroxidase (TPO, P07202, all downloaded July 2020) were imported into T-Coffee (http://tcoffee.crg.cat/apps/tcoffee/do:regular), a program capable of performing multisequence alignment. The fasta_aln output option was selected. All other variables were used as the default setting. The fasta.aln output file was inserted into the Boxshade program (http://www.ch.embnet.org/software/BOX_form.html) to generate a greyscale representation of the sequence alignment. This same method was then performed with human MPO (P05164), mouse MPO (P11247), macaque MPO (F7BAA9), porcine MPO (K7GRV6) and bovine MPO (A6QPT4) as sequence input. Conserved regions and the five sequons of the human MPO polypeptide chains were highlighted.

*Visualisation, modelling, and molecular dynamics*

Amongst the 18 three-dimensional (3D) protein structures currently available of human MPO from the Protein Data Bank (PDB, http://www.rcsb.org/pdb), the structure of the intact diprotomer obtained with the highest resolution by X-ray crystallography containing both the heme, bromide substrates and crystallisable fragments of Asn-conjugated glycans (PDBID, 1D2V, 1.7 Å) was used for visualisation, modelling and structural assessments. Importantly, the MPO used to generate this crystal structure was purified from human neutrophils and covered nearly the entire α- and β-chains of the mature diprotomer (Fiedler et al., 2000). Visualisation and the initial structural assessments were performed using PyMOL Molecular Graphic System v2.3 (Schrödinger, LLC, MA, USA). Signature *N*-glycans (typically the most abundant *N*-glycans identified based on glycan and glycopeptide data, see **Figure 1B**, **Figure 1D**, **Figure 3D**, **Supplementary Table S7** and **Supplementary Table S10**) were added *in-silico* to the four highly occupied sequons of each β chain of diprotomeric MPO (Asn323, Asn355, Asn391, Asn483) to mimic key MPO glycoforms of interest to this study including i) nMPO (referred to as WT in the MD simulations), ii) Endo H-treated nMPO (P1), iii) the highly truncated Asn355-/Asn391-glycophenotype elevated in Se/Pl-MPO (P2) and iv) a semi-truncated Asn355-/Asn391-glycophenotype (P3) included to also explore the structure of MPO when intermediate length glycans were positioned at Asn355 and Asn391, see **Supplementary Table S4** for details. The Asn729 was consistently left unoccupied based on the glycoprofiling data. The 3D structures of the glycans were generated using the Carbohydrate Builder functionality available at http://glycam.org. The glycans were conjugated to the Asn side chains using in-house code. The side chain angles were initially set to common values ($\chi1 = 192°$, $\chi2 = 178°$, $\psi = 177°$, $\phi = 261°$) (Petrescu et al., 2004). The code relieved any atomic overlaps between the conjugated glycan and the underlying protein by adjusting these initial values within the observed ranges (Nivedha et al., 2014; Petrescu et al., 2004). Energy minimisation of the resulting glycoproteins was performed by placing structures a periodic box of approximately 45,000 TIP5P waters with a 10 Å buffer between the glycoprotein and the edges of the box. Energy minimisation of all atoms was performed for 20,000 steps (10,000 steepest decent, followed by 10,000 conjugant gradient).

All molecular dynamics (MD) simulations were performed with the CUDA implementation of the PMEMD (Gotz et al., 2012; Salomon-Ferrer et al., 2013) simulation code, as present in the Amber14 software suite (Case et al., 2014). The GLYCAM06j force field (Kirschner et al., 2008) and Amber14SB force field (Case et al., 2014) were employed for the carbohydrate and protein moieties, respectively. The GAFF forcefield was employed for the heme molecule and the side chains of the residues directly conjugated to the heme (Wang et al., 2004). A Berendsen barostat with a time constant of 1 ps was employed for pressure regulation, while a Langevin thermostat with two collisions per ps was employed for temperature regulation. A nonbonded interaction cut-off of 8 Å was employed. Long-range electrostatics were treated with the particle-mesh Ewald method (Darden et al., 1993). Covalent bonds involving hydrogen were constrained with the SHAKE algorithm, allowing an integration time step of 2 fs to be employed (Ryckaert et al., 1977). All simulations were performed under nPT conditions and the restraints employed were 5 kcal/mol-$Å^2$ Cartesian. The energy-minimised coordinates were equilibrated at 300 K over 400 ps with restraints on the solute heavy atoms. The system was then equilibrated with restraints on the Cα atoms of the protein for 1 ns, prior to performing five unique 500 ns production MD simulations of WT and P1-3 MPO during which no restraints were employed.

The per-residue solvent accessibilities of each of the MD simulated structures were calculated using the Cpptraj program available in Ambertools 18 and averaged across the five production MD simulations of WT and P1-3 MPO. The values from each protomer in the diprotomeric MPO were combined. Values from WT MPO were statistically compared to those from the P1-P3 simulations using $p < 0.05$ as a significance threshold. The results were plotted with Gnuplot 5.2 and visualised on the 3D structure of MPO using VMD 1.9.3. Further, the secondary structure, root mean squared deviation (RMSD) and root mean squared fluctuation (RMSF) were calculated over the trajectory using Cpptraj in AmberTools 18.

The cpptraj program was used to extract 1,000 snapshots from the 500 ns MD simulation of the WT and P1 MPO. Each snapshot was aligned with human ceruloplasmin by employing a 3D structure of the ceruloplasmin-MPO complex (PDBID, 4EJX, 4.7 Å) (Samygina et al., 2013) using VMD 1.9.3 (Humphrey et al., 1996). The degree to which each MPO glycan would overlap with the complexed ceruloplasmin was visualised by displaying each shape of the MPO glycan from the 1,000 snapshots in VMD 1.9.3.

*Biosynthesis-related solvent accessibility of MPO*

To assess the relative solvent accessibility to each of the glycosylation sites of MPO and their conjugated *N*-glycans in the context of MPO biosynthesis and processing (see **Figure 2**), five high-resolution 3D structures of diprotomeric human MPO were selected from the Protein Data Bank including 1D2V (1.75 Å), 1CXP (1.8 Å), 1DNU (1.85 Å), 1DNW (1.9 Å), and 5FIW (1.7 Å). Two separate assessments were then performed using these five structures of human MPO.

Firstly, to determine the relative solvent accessibilities to Asn323, Asn355, Asn391, Asn483, and Asn729 of monoprotomeric MPO (αβ) as the trafficking protein would appear in the early-stage biosynthesis within the *cis*-Golgi, the five PDB files were manually edited to create an artificial monoprotomeric MPO comprising a single α- and β-chain. The five monoprotomeric MPO structures were deposited and processed using the Glycoprotein Builder functionality of GLYCAM web (http://glycam.org), which stripped the structures of all hetero-atoms, water, ions, heme and glycans, and energy-minimised the resulting glycoprotein structures. The relative solvent accessibilities of the side chains of non-glycosylated Asn323, Asn355, Asn391, Asn483, and Asn729 were then established using NACCESS, which rolls a spherical probe (5 Å radii) on the surface of MPO to determine the atomic accessible areas (Van der Waal's interaction) by providing unit-less output values that are comparable across sites and protein structures (Hubber and Thornton, 1993).

Secondly, to determine the relative solvent accessibilities of *N*-glycans proximal to the dimerisation-interface of diprotomeric (ααββ) MPO in the context of late-stage MPO biosynthesis/processing in the late Golgi, or shortly before reaching or upon arrival in the granules, the five selected PDB structures were deposited and processed in the Glycoprotein Builder in their mono- and diprotomeric forms as detailed above. However, for these experiments FA1-glycans (see **Figure 1** for structure) were attached to Asn323, Asn483 and Asn729 carrying a significant level of processed *N*-glycans as determined using glycopeptide analysis. Of these three sites, Asn323 and Asn483 were proximal while Asn729 was distal to the dimerisation interface. The under-processed Asn355 and Asn391 largely carrying oligomannosidic *N*-glycans were not considered in this assessment. The PDB output files were applied to NACCESS using a 5 Å radii probe as detailed above, but the accessibility measurements differed slightly by this time considering the hetero-atoms enabling the determination of the relative solvent accessibilities to the conjugated *N*-glycans rather than the underlying Asn residues. The relative solvent accessibilities to the entire monosaccharide residue of the terminal α1,3-armpositioned β1,2-*N*-acetyl-β-D-glucosamine (GlcNAc) moiety of the FA1-glycans attached to each site of the mono- and diprotomeric MPO as it would be experienced by the processing glycoside hydrolases i.e. the *N*-acetyl-β-hexosaminidase isoenzymes were used for ratiometric analyses.

*Endoglycosidase H-treatment of nMPO*

Intact nMPO (0.5–15 µg) was incubated with or without 1 µL recombinant *Streptomyces plicatus* endoglycosidase H (Endo H) expressed in *E. coli*, 500 U/µL (Promega, Australia) in a total volume of 60-150 µL buffer containing PBS or 5 mM aqueous sodium acetate and 10 mM aqueous sodium chloride, 37°C, 16 h, under native conditions. Blanks (water) and samples containing only Endo H and ceruloplasmin (see below) were included as controls. The untreated nMPO, Endo H-treated nMPO and controls were used immediately for functional assays and characterised using circular dichroism (CD), SDS-PAGE and LC-MS/MS.

*Chlorination and oxidation activity of MPO (activity assay 1–3)*

Three independent enzyme activity assays designated activity assay 1–3 were performed to establish the chlorination and oxidation activity of the granule-separated MPO in crude protein mixtures, and the isolated Endo H-treated and untreated nMPO.

<u>Activity assay 1</u>

The chlorination activity of MPO was determined by the formation of hypochlorous acid (HOCl) captured via taurine per time as described (Dypbukt et al., 2005). Firstly, a HOCl standard curve was generated by adding known concentrations of HOCl freshly made from an aqueous sodium hypochlorite solution (catalogue number 425044, Sigma, Australia) to a final concentration of 5 mM aqueous taurine (Sigma, Australia) in PBS, which was kept on ice for 30 min prior to measurements. For the standard curve, the concentrations of hypochlorite ($OCl^-$) were determined by measuring its absorbance at 292 nm at pH 12 ($\varepsilon_{292}$ = 350 $M^{-1}cm^{-1}$), and solutions were subsequently neutralised before use. The relative and absolute chlorination activities of granule-fractionated MPO were established using 2 µg total protein extract from each of the Az, Sp/Ge and Se/Pl fractions from Donor a-b in technical triplicates and triplicate experiments using 0.5 µg Endo H-treated and untreated nMPO per replicate, respectively. The protein samples were mixed with 5 mM aqueous taurine (final concentration) and firstly incubated for 30 min at room temperature. Reactions were then initiated by the addition of fresh hydrogen peroxide ($H_2O_2$) (catalogue number 516813, Sigma, Australia) to a final concentration of 20 µM. The reactions ran for 20 min at room temperature, before being stopped by the addition of 20 µg/mL aqueous bovine catalase (final concentration, Sigma, Australia). In total,

200 µL of each reaction and the HOCl standards were mixed with 50 µL developing reagent containing 2 mM aqueous TMB in 400 mM aqueous acetate buffer, 10% (v/v) aqueous dimethylformamide and 100 µM aqueous sodium iodide, pH 5.4 in a 96-well plate. After lightly shaking the plate for 5 min, the absorbance of each sample was measured using a plate reader at 650 nm (SPECTROstar Nano, BMG Labtech, Germany). The absolute amount of HOCl produced was determined for Endo H-treated and untreated nMPO based on a pre-established standard curve after baseline subtraction using water blanks and after subtracting readings of the Endo H only samples from the Endo H-treated nMPO (called "nMPO + Endo H^"). The relative chlorination activities of the granule-fractionated MPO were determined after blank correction of absorbance measurements and normalisation to the relative MPO protein levels in each granule fraction determined using LC-MS/MS peptide-based quantitation, see **Figure 3B** for relative MPO protein levels across granules.

Activity assay 2

The 3,3',5,5'-tetramethylbenzidine (TMB) reagent is a chromogenic substrate that can report on the MPO oxidation activity. This assay was used to measure the relative oxidation activities of granule-fractionated MPO in 2 µg total protein extract from each of the Az, Sp/Ge and Se/Pl fractions from Donor a-b (each in technical triplicates), and of 0.05 µg Endo H-treated and untreated nMPO per replicate in technical triplicates. The protein samples were diluted in PBS to a total sample volume of 50 µL and subsequently mixed with 150 µL TMB solution (catalogue number T4444, Sigma-Aldrich, Australia) in a 96-well plate and incubated for 10 min at room temperature. The reaction was quenched by the addition of 100 µL 1 M aqueous sulphuric acid (Sigma, Australia) and the colour intensity was measured on a plate reader at 450 nm absorbance (SPECTROstar Nano, BMG Labtech, Germany). The relatively oxidation activities of Endo H-treated and untreated nMPO were established after baseline subtraction using water blanks and after subtracting absorbance values of an Endo H only sample from the absorbance values measured of Endo H-treated nMPO (called "nMPO + Endo H^"). The relatively oxidation activities of the granule-fractionated MPO were determined after blank correction of absorbance measurements and normalisation to the relative MPO protein levels in each granule fraction determined using LC-MS/MS peptide quantitation, see **Figure 3B**.

Activity assay 3

The 1,2-phenylenediamine reagent is another chromogenic peroxidase substrate that can be utilised to measure the oxidation activity of MPO. This assay was used to measure the relative oxidation activities of 0.05 µg Endo H-treated and untreated nMPO per replicate in technical triplicates. Insufficient material from the granule-separated MPO in crude protein mixtures from Donor a-b was available to probe these samples with this activity assay. The protein samples were diluted in PBS to a total volume of 50 µL which was mixed with 150 µL aqueous 1,2-phenylenediamine (0.4 mg/mL final concentration, Sigma, Australia) and 0.015% (v/v) aqueous $H_2O_2$ (Sigma, Australia) solution and incubated for 10 min at room temperature in the dark. The reaction was quenched by the addition of 100 µL 1 M aqueous sulphuric acid (Sigma, Australia) and the colour intensity was measured on a plate reader at 492 nm absorbance (SPECTROstar Nano, BMG Labtech, Germany). The relatively oxidation activities of Endo H-treated and untreated nMPO were established after baseline subtraction using water blanks and after subtracting absorbance values of an Endo H only sample from the absorbance values measured of the Endo H-treated nMPO (called "nMPO + Endo H^").

*Ceruloplasmin inhibition of MPO*

Glycoform-dependent inhibition of MPO by ceruloplasmin (UniProtKB, P00450) was investigated using Endo H-treated and untreated nMPO. Three enzyme activity assays (activity assay 1-3) carried out as described above were employed, but the Endo H-treated and untreated nMPO were incubated in technical triplicates in the presence or absence of 100 nM ceruloplasmin purified from human serum (final concentration, catalogue number 187-51, Lee BioSolutions Inc, MO, USA) for 30 min at room temperature, prior to the commencement of the MPO enzyme activity measurements. Samples containing only ceruloplasmin were added to the set of other controls included for the three enzyme activity assays (see above). The absolute chlorination activities were determined for Endo H-treated and untreated nMPO with and without ceruloplasmin inhibition by activity assay 1 and displayed as HOCl produced per min as determined by the pre-established standard curve. The relative oxidation activities were determined by activity assays 2 and 3 as established from relative absorbance values. For all assays, baseline subtraction using water blanks and corrections for the absorbance of ceruloplasmin and Endo H were applied called "nMPO + Cp^" and "nMPO + Endo H + Cp^" to enable direct comparisons to nMPO and nMPO + Endo H^, respectively.

*Circular dichroism profiling, temperature stability and secondary structure prediction*

Circular dichroism (CD) data were collected for two separate preparations of i) 200 µL nMPO (0.1 mg/mL (as determined by the Bradford assay, see above) in 5 mM aqueous sodium acetate and 10 mM aqueous sodium chloride, both final concentrations), ii) 200 µL Endo H-treated nMPO (0.1 mg/mL (as determined by the Bradford assay) in 5 mM aqueous sodium acetate and 10 mM aqueous sodium chloride, both final concentrations and with 500 U Endo H, see above for details of digestion conditions), and iii) 200 µL 500 U Endo H in MilliQ water (Merck-Millipore, Australia) as an enzyme control. The samples were measured separately in 1 mm pathlength cuvettes (Starna Scientific Ltd, UK) in a Jasco J-1500 spectropolarimeter (Jasco, Japan) with 2 nm bandwidth and 2 s response time.

A thermostatted 6-cell changer was used to collect wavelength scans of all protein samples and water every 5°C. The temperature ramp (0.3°C/min + the time for wavelength scans, response time of 16 s) was monitored at 208 nm for all samples. The wavelength scans were performed in the range 260–190 nm and data were plotted as the average of the spectra from 20–50°C before any of the protein samples melted. All wavelength scan spectra were baseline-corrected by subtracting a water spectrum collected in the same cuvettes used for the protein samples. The spectra obtained from the two independent sample preparations were of the same shape and very similar magnitudes. Thus, only one set of data have been presented. To enable direct comparisons between the nMPO glycoforms for the CD spectral profile and the temperature stability experiments, the readings from the Endo H only sample were subtracted from the measurements performed for the Endo H-treated nMPO (denoted “nMPO + Endo H^”), plotted and compared to the CD data of untreated nMPO.

For structure fitting, all spectra data were converted to $\Delta\varepsilon$ ($mol^{-1}cm^{-1}dm^{3}$) per residue based on the Bradford assay concentrations. CD spectra were analysed using a graphical user interface version [GUI to Implement SOMSpec, a CD Secondary Structure Fitting Approach 2018, gitlab.com] of our validated self-organising map approach to secondary structure fitting (Hall et al., 2014; Sklepari et al., 2016). In this approach a reference set, in our case SP175 (Lees et al., 2006) (truncated to 190 nm) augmented with a constructed 100% helix and 100% random coil peptide data (San Miguel et al., 2003) was organised into a map with spectra of similar shape in the same neighbourhood. Another map, which overlays the first, contains the secondary structure contents of the reference proteins. The secondary structures of the intervening nodes were determined by interpolation from five neighbouring references spectra. An unknown spectrum is placed on the map where it minimised the distance (difference) between it and the spectral shape and magnitude of a nodal spectrum. The unknown protein’s secondary structure was then assigned to be that of the node. We ran SOMSpec with a 50 × 50 map, five best matching units, and a wavelength range from 240–190 nm. The goodness of fit of the secondary structure estimates were assessed by considering the overlay of the experimental and predicted spectrum visually and by a normalised RMSD (NRMSD). Visual inspection weights maxima, minima and where a spectrum crosses the zero-line more than other points in the spectrum, whereas the NRMSD approach equally weights all wavelengths. In general, for spectra from 240–190 nm an NRMSD < 0.02 was used as a threshold to indicate a very good fit whereas higher NRMSD indicate more uncertainty in the prediction. Higher values usually require user-interference to choose the best fit.

It was visually apparent from the room temperature spectra that the ratios of the 190 nm:208 nm intensities were smaller than expected for highly helical or sheet proteins. In addition, our measured $\Delta\varepsilon$ values were small, suggesting systematic concentration errors due to the use a non-glycosylated reference protein. In accord with this, attempts to use SOMSpec to predict structures from the spectral data were not satisfactory (NRMSD > 0.08). We therefore removed fractions of a random coil spectrum (in 0.025 steps) for the SufI-KK peptide MSLSKKQFIQASGIALCAGAVPLKASA (San Miguel et al., 2003), and also scaled the spectra to adjust for overestimates of the protein concentrations (in 10% steps) (Bansal et al., 2020). We then reran SOMSpec to get the best possible fit. The structure-fitting results for best fit derandomised and scaled spectra are provided in **Supplementary Table S13C** as are the secondary structure percentages when the removed random coil was added back in. There were two options for best fit depending whether one considers the whole wavelength region or emphasises from 190–210 nm as the 222 nm region is unusually small. Consideration of the crystal structures led us to select the rows where the Bradford-determined concentration was twice the protein residue concentration and 12.5–15% random coil removed for the fitting process. Overall the CD data suggest that both Endo H-treated and untreated nMPO in solution are ~34% helix, ~11% sheet, ~16% bonded turns, ~8% bends, and ~31% other structures (including random coil) with each number being indicative not absolute.

*Data representation and statistics*

Statistical significance was tested using one- or two-tailed paired or unpaired Student's t-tests as indicated for each experiment. $p \geq 0.05$ was chosen as the universal threshold to reject various null hypotheses (e.g. *glycosylation sites are equally solvent accessible* or *MPO glycoforms display the same enzyme activity*). For each test, the statistical confidence has consistently been indicated by *$p < 0.05$, **$p < 0.01$, ***$p < 0.005$, ****$p < 0.001$ and *****$p < 0.0005$. ns signifies a non-significant test ($p \geq 0.05$). The n values representing either biological replicates or technical (sample handling or analytical) replicates have been indicated for each experiment. Data points have been plotted as the mean, and the error bars consistently represent their standard deviation (SD). The *N*-glycans were depicted according to the established symbol nomenclature (Varki et al., 2015), see also **Figure 1** for key.

#### Supplementary Figures

##### Supplementary Figure S1

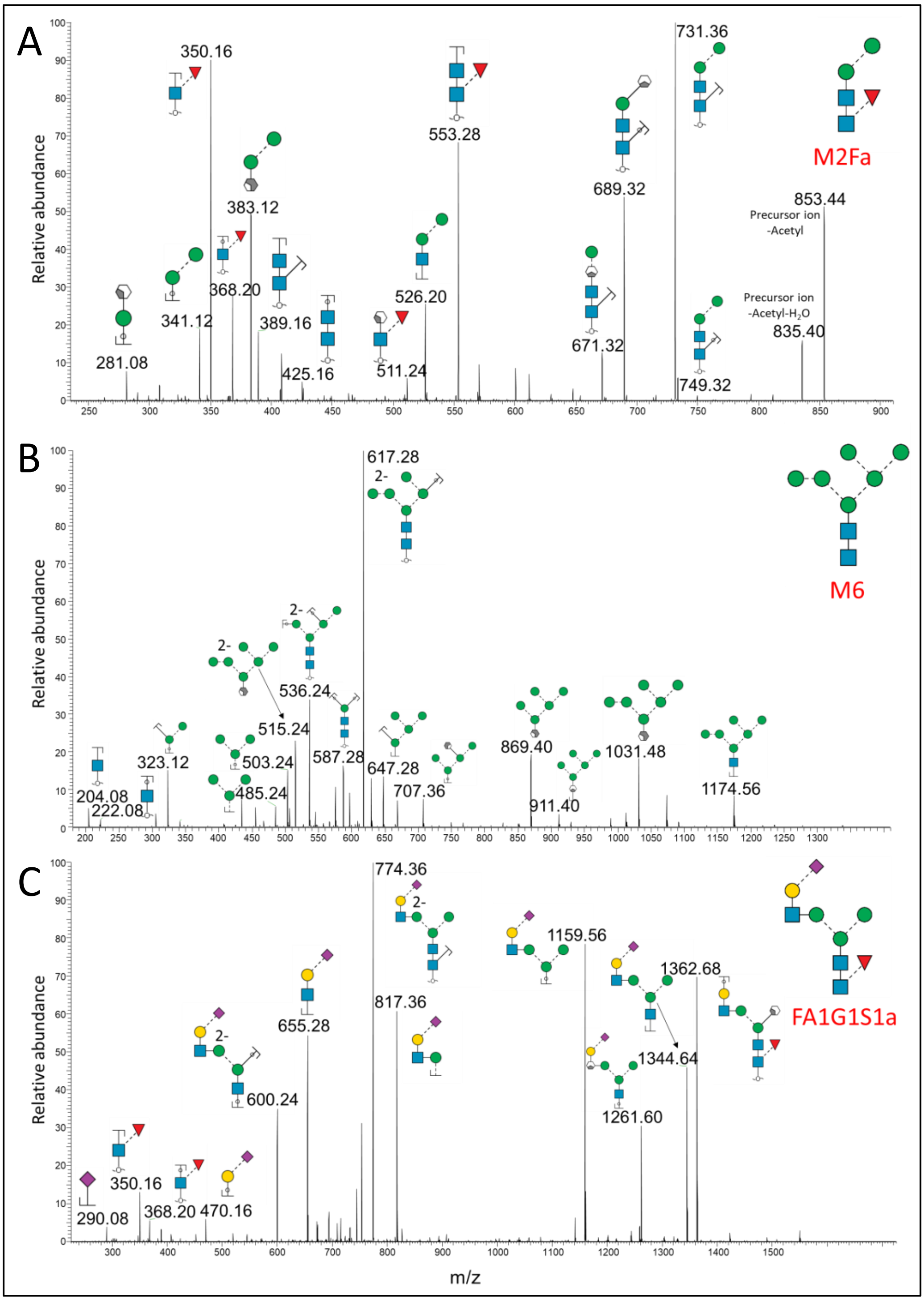


**Supplementary Figure S1.** Examples of annotated CID-MS/MS (-) spectra of signature *N*-glycans decorating nMPO obtained from the PGC-LC-MS/MS glycomics data including **A**) M2Fa, **B**) M6 and **C**) FA1G1S1a. See **Figure 1B** for overview of the *N*-glycan fine structures decorating nMPO and the key to symbols and nomenclature. See **Supplementary Data S1** for all annotated spectral evidence.

**Supplementary Figure S2**

Coverage of mature MPO α-chain: 100.0% and β-chain: 99.8%

$^{1}$MGVPFFSSLRCMVDLGPCWAGGLTAEMKLLLALAGLLAILAT
PQPSEGAAPAVLGEVDTSLVLSSMEEAKQLVDKAYKERRESIKQ
RLRSGSASPMELLSYFKQPVAATRTAVRAADYLHVALDLLERKL
RSLWRRPFNVTDVLTPAQLNVLSKSSGCAYQDV(G)(V)TCPEQ
DKYRTITGMCNNRRSPTLGASNRAFVRWLPAEYEDGFSLPYG
WTPGVKRNGFPVALARAVSNEIVRFPTDQLTPDQERSLMFM
QWGQLLDHDLDFTPEPAAR(A)SFV(T)(G)VNCETSCVQQPPC
FPLKIPPNDPRIKNQADCIPFFRSCPACPGSNITIRNQINALTSF
VDASMVYGSEEPLARNLRNMSNQLGLLAVNQRFQDNGRAL
LPFDNLHDDPCLLTNRSARIPCFLAGDTRSSEMPELTSMHTLLL
REHNRLATELKSLNPRWDGERLYQEARKIVGAMVQIITYRDYL
PLVLGPTAMRKYLPTYRSYNDSVDPRIANVFTNAFRYGHTLIQ
PFMFRLDNRYQPMEPNPRVPLSRVFFASWRVVLEGGIDPILR
GLMATPAKLNRQNQIAVDEIRERLFEQVMRIGLDLPALNMQ
RSRDHGLPGYNAWRRFCGLPQPETVGQLGTVLRNLKLARKL
MEQYGTPNNIDIWMGGVSEPLKRKGRVGPLLACIIGTQFRKL
RDGDRFWWENEGVFSMQQRQALAQISLPRIICDNTGITTVS
KNNIFMSNSYPRDFVNCSTLPALNLASWREAS$^{745}$

Signal peptide | α chain | Hexapeptide
Propeptide | β chain | **Observed residues**
( ) Truncation variant observed | X PTM observed

**Supplementary Figure S2.** Sequence coverage of nMPO as determined based on LC-MS/MS peptide profiling data. Observed regions have been mapped in bold red, see also **Figure 1C** for polypeptide termini and truncation variants of the MPO α- and β-chains and non-glycan modifications of nMPO. See **Supplementary Data S2A-B** for spectral evidence. See key for colour coding.

**Supplementary Figure S3**

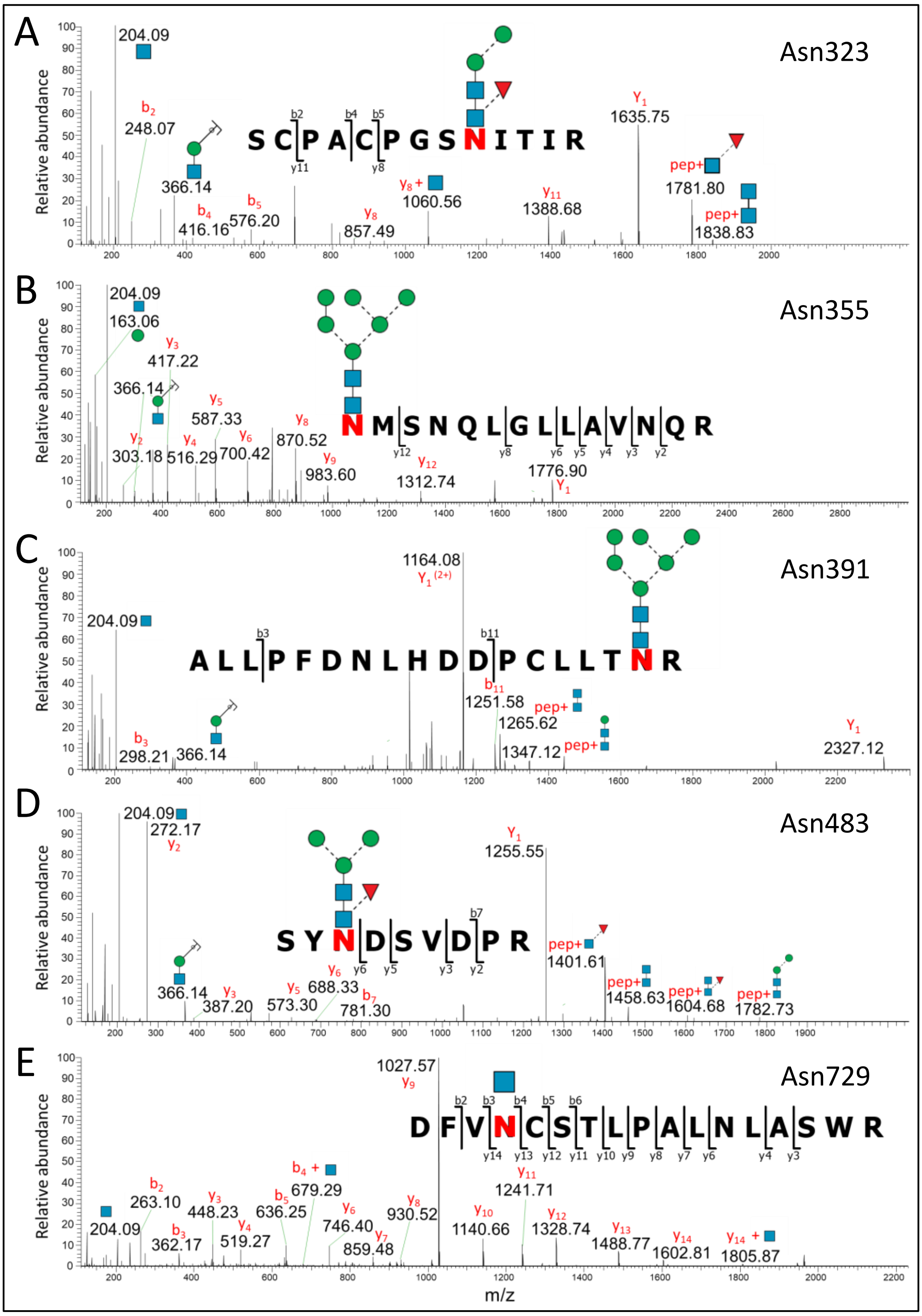


**Supplementary Figure S3.** Examples of annotated HCD-MS/MS (+) spectra of prominent *N*-glycans carried by peptides containing the five MPO sites obtained from the LC-MS/MS peptide data including key glycopeptides spanning **A**) Asn323, **B**) Asn355, **C**) Asn391, **D**) Asn483, and **E**) Asn729, see **Figure 1D** for overview of the site-specific *N*-glycoprofile of nMPO and key to symbols and nomenclature. See **Supplementary Data S3** for all annotated spectral evidence.

#### Supplementary Figure S4

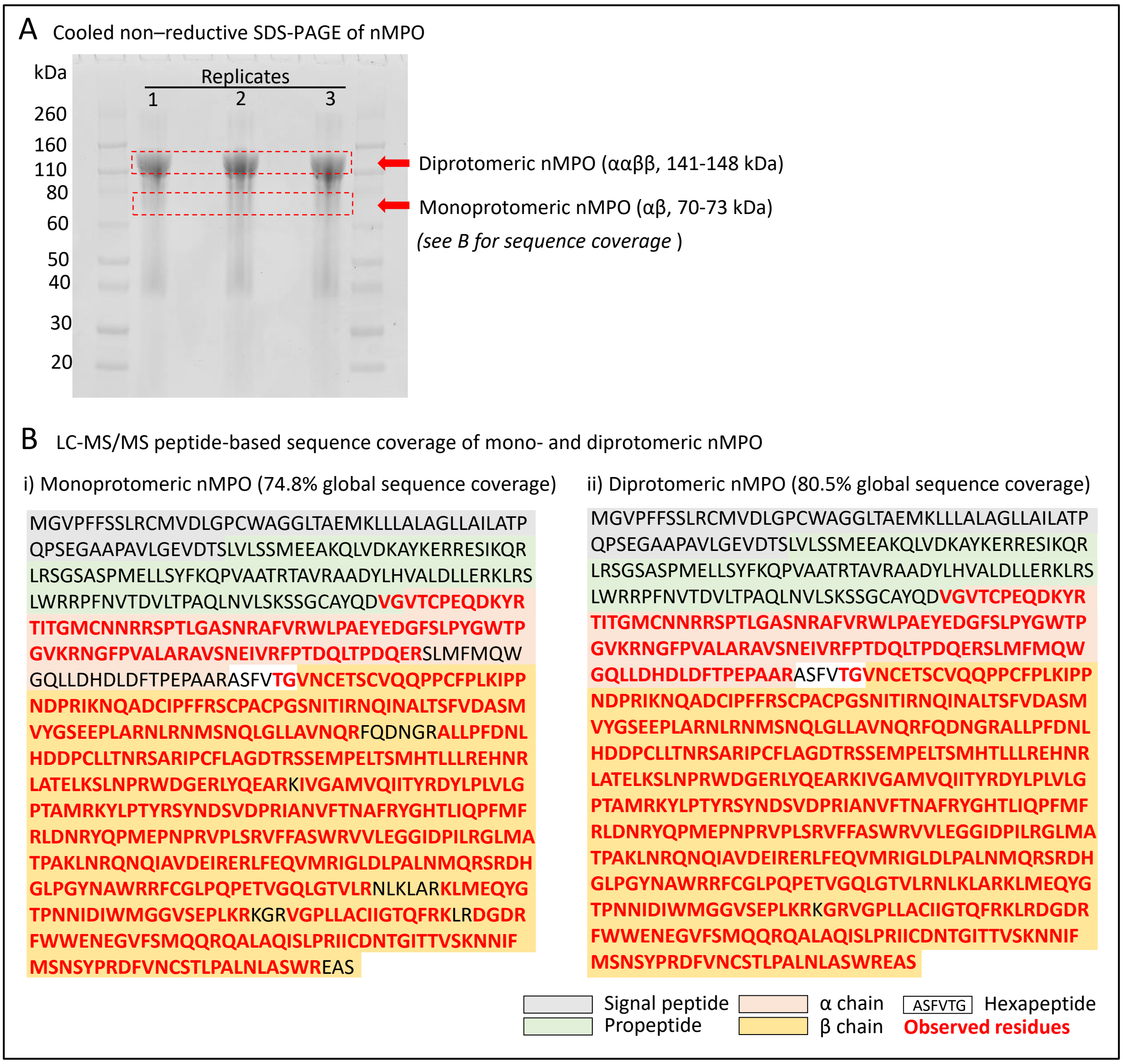


**Supplementary Figure S4.** Structural characterisation of mono- (αβ) and diprotomeric (ααββ) nMPO. **A**) Separation of a low abundance form of monoprotomeric nMPO around 70-73 kDa (7%, as determined using mass photometry, see **Figure 1G**) and the predominant form of diprotomeric nMPO at 141-148 kDa (93%) using non-reductive SDS-PAGE performed on ice to avoid complex dissociation. Gel bands from technical triplicate experiments (broken red boxes) were excised and the extracted tryptic peptide mixtures subjected to LC-MS/MS analysis. **B**) Global sequence coverage (including the proMPO polypeptide region) of the i) mono- and ii) diprotomeric nMPO as determined based on LC-MS/MS peptide data (observed regions in bold red). See key for colour coding.

#### Supplementary Figure S5

### A Asn323

##### i) Glycoprofile

Relative abundance (%)

| | PM-MPO | MM-MPO | BN-MPO | SN-MPO | PMN-nMPO | Az-MPO | Sp-MPO | Ge-MPO | Se-/Pl-MPO | Dg-MPO |
|---|---|---|---|---|---|---|---|---|---|---|
| None | 5.0% | 4.1% | 2.3% | 3.2% | 0.7% | 1.3% | 0.8% | 0.7% | 0.0% | 0.0% |
| HexNAc(1) | 27.2% | 21.1% | 22.3% | 23.9% | 14.9% | 20.5% | 19.6% | 20.2% | 35.6% | 25.0% |
| HexNAc(1)Fuc(1) | 11.3% | 7.3% | 8.0% | 9.3% | 6.3% | 9.1% | 7.2% | 7.5% | 7.2% | 10.7% |
| HexNAc(2) | 0.7% | 0.5% | 0.3% | 0.5% | 1.0% | 1.4% | 1.7% | 1.9% | 3.5% | 0.0% |
| HexNAc(2)Fuc(1) | 1.9% | 1.3% | 1.7% | 1.0% | 1.7% | 1.7% | 1.2% | 1.2% | 1.4% | 0.0% |
| HexNAc(2)Hex(1) | 0.1% | 0.1% | 0.2% | 0.2% | 0.5% | 0.6% | 1.0% | 0.5% | 0.4% | 0.0% |
| HexNAc(2)Hex(1)Fuc(1) | 0.0% | 0.0% | 0.0% | 0.0% | 0.4% | 0.4% | 0.7% | 0.4% | 0.2% | 0.0% |
| HexNAc(2)Hex(2) | 5.5% | 5.3% | 5.8% | 6.4% | 7.1% | 5.9% | 5.3% | 5.2% | 5.1% | 7.5% |
| HexNAc(2)Hex(2)Fuc(1) | 20.5% | 23.1% | 23.0% | 28.9% | 21.9% | 25.8% | 24.4% | 24.7% | 20.6% | 37.6% |
| HexNAc(2)Hex(3) | 3.5% | 4.7% | 4.1% | 2.9% | 4.4% | 3.4% | 3.3% | 3.5% | 2.9% | 0.0% |
| HexNAc(2)Hex(3)Fuc(1) | 6.4% | 7.6% | 9.2% | 7.5% | 11.0% | 10.4% | 8.0% | 9.5% | 7.4% | 11.8% |
| HexNAc(2)Hex(4) | 3.0% | 3.4% | 5.0% | 2.7% | 4.9% | 2.1% | 3.6% | 2.8% | 2.1% | 0.0% |
| HexNAc(2)Hex(4)Fuc(1) | 0.0% | 0.3% | 0.1% | 0.1% | 0.0% | 0.0% | 0.0% | 0.0% | 0.0% | 0.0% |
| HexNAc(2)Hex(5) | 3.4% | 5.7% | 6.2% | 4.3% | 7.6% | 4.0% | 7.3% | 6.8% | 3.0% | 0.0% |
| HexNAc(2)Hex(5)Fuc(1) | 0.2% | 0.7% | 0.4% | 0.4% | 0.0% | 0.0% | 0.0% | 0.0% | 0.0% | 0.0% |
| HexNAc(2)Hex(6) | 5.9% | 8.0% | 7.4% | 5.0% | 6.6% | 7.6% | 8.7% | 7.7% | 5.9% | 3.7% |
| HexNAc(2)Hex(7) | 2.3% | 4.7% | 3.0% | 3.3% | 4.2% | 4.5% | 6.1% | 6.1% | 3.9% | 1.9% |
| HexNAc(2)Hex(7)Phos(1) | 0.8% | 0.9% | 0.1% | 0.1% | 2.9% | 0.0% | 0.0% | 0.0% | 0.0% | 0.0% |
| HexNAc(2)Hex(7)Phos(2) | 0.0% | 0.0% | 0.0% | 0.0% | 0.9% | 0.0% | 0.0% | 0.0% | 0.0% | 1.8% |
| HexNAc(2)Hex(8) | 0.0% | 0.0% | 0.0% | 0.0% | 0.1% | 0.0% | 0.0% | 0.0% | 0.0% | 0.0% |
| HexNAc(2)Hex(9) | 1.1% | 0.2% | 0.0% | 0.0% | 0.0% | 0.0% | 0.0% | 0.0% | 0.0% | 0.0% |
| HexNAc(2)Hex(10) | 0.2% | 0.0% | 0.0% | 0.0% | 0.0% | 0.0% | 0.0% | 0.0% | 0.0% | 0.0% |
| HexNAc(3)Hex(3)Fuc(1) | 0.8% | 0.7% | 0.7% | 0.3% | 0.0% | 0.0% | 0.0% | 0.0% | 0.0% | 0.0% |
| HexNAc(3)Hex(3)Fuc(1)NeuAc(1) | 0.0% | 0.0% | 0.0% | 0.0% | 0.0% | 0.0% | 0.0% | 0.0% | 0.0% | 0.0% |
| HexNAc(3)Hex(4)Fuc(1) | 0.0% | 0.0% | 0.0% | 0.0% | 0.0% | 0.0% | 0.0% | 0.0% | 0.0% | 0.0% |
| HexNAc(3)Hex(4)Fuc(1)NeuAc(1) | 0.1% | 0.4% | 0.2% | 0.2% | 2.6% | 1.2% | 1.2% | 1.1% | 0.8% | 0.0% |
| HexNAc(4)Hex(4)Fuc(1)NeuAc(1) | 0.0% | 0.0% | 0.0% | 0.0% | 0.2% | 0.0% | 0.0% | 0.0% | 0.0% | 0.0% |
| HexNAc(3)Hex(4)NeuAc(1) | 0.0% | 0.0% | 0.0% | 0.0% | 0.1% | 0.0% | 0.0% | 0.0% | 0.0% | 0.0% |
| HexNAc(3)Hex(5)Fuc(1) | 0.0% | 0.0% | 0.0% | 0.0% | 0.1% | 0.0% | 0.0% | 0.0% | 0.0% | 0.0% |
| HexNAc(4)Hex(5)Fuc(1)NeuAc(2) | 0.0% | 0.0% | 0.0% | 0.0% | 0.1% | 0.0% | 0.0% | 0.0% | 0.0% | 0.0% |
| SUM | 100.0% | 100.0% | 100.0% | 100.0% | 100.0% | 100.0% | 100.0% | 100.0% | 100.0% | 100.0% |

##### ii) Correlation analysis

| | PM-MPO | MM-MPO | BN-MPO | SN-MPO | PMN-nMPO |
|---|---|---|---|---|---|
| PM-MPO | | 0.9640 | 0.9670 | 0.9639 | 0.8793 |
| MM-MPO | | | 0.9926 | 0.9851 | 0.9559 |
| BN-MPO | | | | 0.9830 | 0.9627 |
| SN-MPO | | | | | 0.9433 |
| PMN-nMPO | | | | | |

| | Az-MPO | Sp-MPO | Ge-MPO | Se/Pl-MPO | PMN-nMPO | Dg-MPO |
|---|---|---|---|---|---|---|
| Az-MPO | | 0.9870 | 0.9925 | 0.9096 | 0.9675 | 0.9702 |
| Sp-MPO | | | 0.9977 | 0.9038 | 0.9715 | 0.9374 |
| Ge-MPO | | | | 0.9100 | 0.9743 | 0.9478 |
| Se/Pl-MPO | | | | | 0.8393 | 0.8714 |
| PMN-nMPO | | | | | | 0.9212 |
| Dg-MPO | | | | | | |

### B Asn355

#### i) Glycoprofile

Relative abundance (%)

| | PM-MPO | MM-MPO | BN-MPO | SN-MPO | PMN-nMPO | Az-MPO | Sp-MPO | Ge-MPO | Se-/Pl-MPO | Dg-MPO |
|---|---|---|---|---|---|---|---|---|---|---|
| None | 1.2% | 0.5% | 0.9% | 0.5% | 0.8% | 1.0% | 1.6% | 2.0% | 6.1% | 0.0% |
| HexNAc(1) | 0.0% | 0.0% | 0.0% | 0.0% | 0.2% | 0.8% | 1.3% | 3.2% | 4.0% | 0.0% |
| HexNAc(2) | 0.0% | 0.0% | 0.0% | 0.0% | 0.0% | 0.4% | 0.1% | 1.5% | 0.4% | 0.0% |
| HexNAc(2)Hex(1) | 0.0% | 0.0% | 0.0% | 0.0% | 0.1% | 0.0% | 0.0% | 0.0% | 0.0% | 0.0% |
| HexNAc(2)Hex(2) | 0.0% | 0.0% | 0.0% | 0.0% | 0.1% | 0.0% | 0.1% | 0.1% | 0.0% | 0.0% |
| HexNAc(2)Hex(3) | 0.9% | 1.3% | 0.8% | 1.2% | 2.3% | 2.4% | 1.7% | 1.3% | 1.2% | 0.0% |
| HexNAc(2)Hex(4) | 2.4% | 3.5% | 3.9% | 4.2% | 4.4% | 7.0% | 6.4% | 5.3% | 5.2% | 0.0% |
| HexNAc(2)Hex(5) | 27.8% | 28.4% | 36.2% | 39.6% | 38.6% | 34.2% | 35.7% | 29.7% | 31.8% | 11.5% |
| HexNAc(2)Hex(6) | 46.7% | 51.9% | 46.8% | 45.8% | 43.6% | 44.4% | 44.6% | 44.2% | 40.4% | 79.1% |
| HexNAc(2)Hex(7) | 16.3% | 12.3% | 9.4% | 7.6% | 9.3% | 9.3% | 8.3% | 12.6% | 10.9% | 9.5% |
| HexNAc(2)Hex(8) | 0.7% | 0.1% | 0.0% | 0.0% | 0.4% | 0.3% | 0.0% | 0.0% | 0.0% | 0.0% |
| HexNAc(2)Hex(9) | 1.7% | 0.0% | 0.0% | 0.0% | 0.0% | 0.0% | 0.0% | 0.0% | 0.0% | 0.0% |
| HexNAc(3)Hex(4) | 0.1% | 0.2% | 0.2% | 0.1% | 0.0% | 0.0% | 0.0% | 0.0% | 0.0% | 0.0% |
| HexNAc(3)Hex(5) | 0.3% | 0.7% | 0.5% | 0.0% | 0.0% | 0.0% | 0.0% | 0.0% | 0.0% | 0.0% |
| HexNAc(3)Hex(6) | 2.0% | 1.0% | 1.1% | 1.0% | 0.0% | 0.0% | 0.0% | 0.0% | 0.0% | 0.0% |
| SUM | 100.0% | 100.0% | 100.0% | 100.0% | 100.0% | 100.0% | 100.0% | 100.0% | 100.0% | 100.0% |

#### ii) Correlation analysis

| | PM-MPO | MM-MPO | BN-MPO | SN-MPO | PMN-nMPO |
|---|---|---|---|---|---|
| PM-MPO | | 0.9940 | 0.9795 | 0.9642 | 0.9661 |
| MM-MPO | | | 0.9840 | 0.9698 | 0.9686 |
| BN-MPO | | | | 0.9976 | 0.9966 |
| SN-MPO | | | | | 0.9989 |
| PMN-nMPO | | | | | |

| | Az-MPO | Sp-MPO | Ge-MPO | Se/Pl-MPO | PMN-nMPO | Dg-MPO |
|---|---|---|---|---|---|---|
| Az-MPO | | 0.9992 | 0.9926 | 0.9904 | 0.9958 | 0.8525 |
| Sp-MPO | | | 0.9903 | 0.9919 | 0.9972 | 0.8431 |
| Ge-MPO | | | | 0.9922 | 0.9840 | 0.8851 |
| Se/Pl-MPO | | | | | 0.9879 | 0.8424 |
| PMN-nMPO | | | | | | 0.8159 |
| Dg-MPO | | | | | | |

### C Asn391

#### i) Glycoprofile

Relative abundance (%)

| | PM-MPO | MM-MPO | BN-MPO | SN-MPO | PMN-nMPO | Az-MPO | Sp-MPO | Ge-MPO | Se-/Pl-MPO | Dg-MPO |
|---|---|---|---|---|---|---|---|---|---|---|
| None | 5.1% | 11.7% | 3.1% | 5.2% | 0.7% | 0.5% | 0.5% | 0.2% | 0.3% | 2.0% |
| HexNAc(1) | 0.0% | 0.0% | 0.0% | 0.0% | 0.3% | 0.7% | 0.6% | 0.7% | 2.6% | 0.0% |
| HexNAc(2) | 0.0% | 0.0% | 0.0% | 0.0% | 0.1% | 0.3% | 0.1% | 0.2% | 0.5% | 0.0% |
| HexNAc(2)Hex(1) | 0.0% | 0.0% | 0.0% | 0.0% | 0.0% | 0.0% | 0.0% | 0.0% | 0.0% | 0.0% |
| HexNAc(2)Hex(2) | 11.4% | 5.4% | 3.5% | 2.3% | 2.5% | 1.1% | 1.1% | 0.6% | 0.8% | 0.0% |
| HexNAc(2)Hex(3) | 24.1% | 23.9% | 22.1% | 27.0% | 30.8% | 33.0% | 35.2% | 28.9% | 28.2% | 33.1% |
| HexNAc(2)Hex(3)Fuc(1) | 0.0% | 0.0% | 0.0% | 0.1% | 0.1% | 0.1% | 0.1% | 0.1% | 0.0% | 0.0% |
| HexNAc(2)Hex(4) | 15.5% | 21.1% | 28.7% | 25.1% | 12.3% | 14.7% | 15.3% | 13.3% | 12.0% | 14.5% |
| HexNAc(2)Hex(5) | 25.3% | 25.6% | 25.1% | 20.9% | 20.3% | 24.4% | 22.3% | 24.5% | 25.4% | 25.8% |
| HexNAc(2)Hex(6) | 16.0% | 11.9% | 17.2% | 19.1% | 31.4% | 24.5% | 23.8% | 30.5% | 29.9% | 24.5% |
| HexNAc(2)Hex(7) | 0.6% | 0.2% | 0.2% | 0.3% | 0.3% | 0.2% | 0.2% | 0.2% | 0.0% | 0.0% |
| HexNAc(2)Hex(8) | 0.1% | 0.0% | 0.0% | 0.0% | 0.0% | 0.0% | 0.0% | 0.0% | 0.0% | 0.0% |
| HexNAc(2)Hex(9) | 1.8% | 0.0% | 0.0% | 0.0% | 0.0% | 0.0% | 0.0% | 0.0% | 0.0% | 0.0% |
| HexNAc(3)Hex(4)NeuAc(1) | 0.1% | 0.1% | 0.2% | 0.2% | 0.7% | 0.3% | 0.6% | 0.4% | 0.2% | 0.0% |
| HexNAc(3)Hex(5)NeuAc(1) | 0.0% | 0.0% | 0.0% | 0.0% | 0.3% | 0.1% | 0.2% | 0.4% | 0.1% | 0.0% |
| HexNAc(3)Hex(6)NeuAc(1) | 0.0% | 0.0% | 0.0% | 0.0% | 0.2% | 0.0% | 0.0% | 0.0% | 0.0% | 0.0% |
| SUM | 100.0% | 100.0% | 100.0% | 100.0% | 100.0% | 100.0% | 100.0% | 100.0% | 100.0% | 100.0% |

#### ii) Correlation analysis

| | PM-MPO | MM-MPO | BN-MPO | SN-MPO | PMN-nMPO |
|---|---|---|---|---|---|
| PM-MPO | | 0.9504 | 0.9215 | 0.9295 | 0.8840 |
| MM-MPO | | | 0.9481 | 0.9509 | 0.8055 |
| BN-MPO | | | | 0.9803 | 0.8375 |
| SN-MPO | | | | | 0.9019 |
| PMN-nMPO | | | | | |

| | Az-MPO | Sp-MPO | Ge-MPO | Se/Pl-MPO | PMN-nMPO | Dg-MPO |
|---|---|---|---|---|---|---|
| Az-MPO | | 0.9974 | 0.9855 | 0.9827 | 0.9797 | 0.9986 |
| Sp-MPO | | | 0.9754 | 0.9705 | 0.9759 | 0.9944 |
| Ge-MPO | | | | 0.9982 | 0.9928 | 0.9841 |
| Se/Pl-MPO | | | | | 0.9883 | 0.9817 |
| PMN-nMPO | | | | | | 0.9749 |
| Dg-MPO | | | | | | |

### D Asn483

#### i) Glycoprofile

| | Relative abundance (%) | | | | | | | | | |
|---|---|---|---|---|---|---|---|---|---|---|
| | PM-MPO | MM-MPO | BN-MPO | SN-MPO | PMN-nMPO | Az-MPO | Sp-MPO | Ge-MPO | Se-/Pl-MPO | Dg-MPO |
| None | 0.0% | 0.0% | 0.0% | 0.0% | 0.0% | 0.2% | 0.2% | 0.3% | 0.5% | 0.0% |
| HexNAc(1)Fuc(1) | 0.4% | 5.0% | 5.6% | 5.5% | 0.0% | 0.7% | 0.9% | 1.9% | 5.4% | 0.0% |
| HexNAc(2)Hex(2)Fuc(1) | 1.0% | 2.1% | 1.3% | 2.2% | 2.4% | 2.0% | 2.1% | 2.1% | 2.8% | 0.0% |
| HexNAc(2)Hex(3) | 0.0% | 0.0% | 0.0% | 0.0% | 0.1% | 0.1% | 0.2% | 0.2% | 0.4% | 0.0% |
| HexNAc(2)Hex(3)Fuc(1) | 73.4% | 56.4% | 63.3% | 63.8% | 53.0% | 51.1% | 49.8% | 50.7% | 52.0% | 81.8% |
| HexNAc(2)Hex(4)Fuc(1) | 1.4% | 3.5% | 1.2% | 1.7% | 3.0% | 3.4% | 2.6% | 2.1% | 2.7% | 2.3% |
| HexNAc(2)Hex(5) | 0.2% | 0.4% | 0.1% | 0.0% | 0.1% | 0.2% | 0.4% | 0.3% | 0.3% | 0.0% |
| HexNAc(2)Hex(5)Fuc(1) | 1.5% | 3.6% | 1.9% | 1.8% | 2.6% | 2.5% | 2.0% | 2.0% | 2.0% | 0.0% |
| HexNAc(2)Hex(6) | 0.1% | 0.2% | 0.1% | 0.0% | 0.0% | 0.0% | 0.0% | 0.0% | 0.0% | 0.0% |
| HexNAc(2)Hex(7) | 0.0% | 0.0% | 0.0% | 0.0% | 0.0% | 0.0% | 0.0% | 0.0% | 0.0% | 0.0% |
| HexNAc(2)Hex(9) | 0.8% | 0.0% | 0.0% | 0.0% | 0.0% | 0.0% | 0.0% | 0.0% | 0.0% | 0.0% |
| HexNAc(2)Hex(10) | 0.1% | 0.0% | 0.0% | 0.0% | 0.0% | 0.0% | 0.0% | 0.0% | 0.0% | 0.0% |
| HexNAc(3)Hex(3)Fuc(1) | 15.6% | 18.6% | 18.9% | 18.1% | 19.4% | 20.2% | 17.7% | 16.5% | 19.3% | 15.9% |
| HexNAc(3)Hex(4)Fuc(1) | 1.2% | 1.9% | 0.9% | 1.3% | 0.0% | 2.9% | 3.3% | 3.2% | 2.5% | 0.0% |
| HexNAc(3)Hex(4)Fuc(1)NeuAc(1) | 0.9% | 1.4% | 1.5% | 1.0% | 6.8% | 5.7% | 8.7% | 8.7% | 4.7% | 0.0% |
| HexNAc(3)Hex(4)Fuc(2) | 0.8% | 1.6% | 0.9% | 0.6% | 0.0% | 0.0% | 0.0% | 0.0% | 0.0% | 0.0% |
| HexNAc(3)Hex(6)Fuc(1)NeuAc(1) | 0.1% | 0.1% | 0.0% | 0.0% | 0.6% | 0.4% | 0.2% | 0.4% | 0.2% | 0.0% |
| HexNAc(4)Hex(3)Fuc(1) | 0.6% | 2.1% | 1.6% | 2.1% | 1.8% | 2.0% | 1.5% | 1.8% | 1.4% | 0.0% |
| HexNAc(4)Hex(4)Fuc(1) | 0.2% | 0.7% | 0.3% | 0.3% | 0.0% | 0.9% | 1.0% | 1.0% | 0.6% | 0.0% |
| HexNAc(4)Hex(4)Fuc(1)NeuAc(1) | 1.3% | 1.4% | 1.4% | 1.1% | 9.5% | 7.1% | 9.2% | 8.3% | 5.1% | 0.0% |
| HexNAc(4)Hex(4)Fuc(2) | 0.4% | 0.9% | 1.0% | 0.6% | 0.0% | 0.0% | 0.0% | 0.0% | 0.0% | 0.0% |
| HexNAc(4)Hex(5)Fuc(1)NeuAc(1) | 0.0% | 0.0% | 0.0% | 0.0% | 0.7% | 0.4% | 0.4% | 0.5% | 0.3% | 0.0% |
| SUM | 100.0% | 100.0% | 100.0% | 100.0% | 100.0% | 100.0% | 100.0% | 100.0% | 100.0% | 100.0% |

#### ii) Correlation analysis

| | PM-MPO | MM-MPO | BN-MPO | SN-MPO | PMN-nMPO |
|---|---|---|---|---|---|
| PM-MPO | | 0.9902 | 0.9935 | 0.9944 | 0.9730 |
| MM-MPO | | | 0.9983 | 0.9981 | 0.9769 |
| BN-MPO | | | | 0.9996 | 0.9765 |
| SN-MPO | | | | | 0.9744 |
| PMN-nMPO | | | | | |

| | Az-MPO | Sp-MPO | Ge-MPO | Se/Pl-MPO | PMN-nMPO | Dg-MPO |
|---|---|---|---|---|---|---|
| Az-MPO | | 0.9959 | 0.9948 | 0.9943 | 0.9974 | 0.9710 |
| Sp-MPO | | | 0.9991 | 0.9886 | 0.9978 | 0.9637 |
| Ge-MPO | | | | 0.9909 | 0.9963 | 0.9702 |
| Se/Pl-MPO | | | | | 0.9899 | 0.9769 |
| PMN-nMPO | | | | | | 0.9679 |
| Dg-MPO | | | | | | |

### E Asn729

#### i) Glycoprofile

| | Relative abundance (%) | | | | | | | | | |
|---|---|---|---|---|---|---|---|---|---|---|
| | PM-MPO | MM-MPO | BN-MPO | SN-MPO | PMN-nMPO | Az-MPO | Sp-MPO | Ge-MPO | Se-/Pl-MPO | Dg-MPO |
| None | 62.3% | 48.7% | 46.6% | 41.3% | 44.3% | 39.4% | 43.8% | 35.4% | 30.5% | 43.1% |
| HexNAc(1) | 10.4% | 9.6% | 12.9% | 22.7% | 21.8% | 27.0% | 18.2% | 19.4% | 23.5% | 27.7% |
| HexNAc(1)Fuc(1) | 7.7% | 11.4% | 12.0% | 11.1% | 9.3% | 9.8% | 15.6% | 12.7% | 14.0% | 9.3% |
| HexNAc(2) | 0.2% | 0.5% | 0.6% | 0.7% | 0.9% | 0.1% | 0.2% | 0.1% | 1.0% | 0.0% |
| HexNAc(2)Fuc(1) | 1.1% | 1.8% | 1.9% | 2.0% | 0.7% | 0.8% | 0.6% | 0.8% | 0.8% | 0.0% |
| HexNAc(2)Hex(1) | 0.5% | 1.2% | 1.3% | 1.5% | 1.2% | 1.1% | 2.2% | 1.0% | 0.9% | 0.0% |
| HexNAc(2)Hex(1)Fuc(1) | 0.7% | 1.8% | 1.9% | 1.8% | 1.7% | 2.7% | 1.9% | 2.6% | 2.7% | 0.0% |
| HexNAc(2)Hex(2) | 3.4% | 7.1% | 7.0% | 7.1% | 5.1% | 5.3% | 4.4% | 5.6% | 5.8% | 0.0% |
| HexNAc(2)Hex(2)Fuc(1) | 10.7% | 15.1% | 13.5% | 10.1% | 12.1% | 12.9% | 12.2% | 20.1% | 19.2% | 19.8% |
| HexNAc(2)Hex(3) | 0.4% | 0.7% | 0.5% | 0.4% | 0.1% | 0.1% | 0.1% | 0.5% | 0.2% | 0.0% |
| HexNAc(2)Hex(3)Fuc(1) | 1.5% | 0.7% | 0.8% | 0.3% | 0.8% | 0.1% | 0.3% | 1.0% | 0.6% | 0.0% |
| HexNAc(2)Hex(4) | 0.2% | 0.5% | 0.6% | 0.6% | 0.6% | 0.1% | 0.3% | 0.6% | 0.8% | 0.0% |
| HexNAc(2)Hex(4)Fuc(1) | 0.2% | 0.2% | 0.1% | 0.0% | 0.1% | 0.0% | 0.0% | 0.0% | 0.0% | 0.0% |
| HexNAc(2)Hex(5) | 0.1% | 0.3% | 0.2% | 0.1% | 0.5% | 0.0% | 0.0% | 0.0% | 0.0% | 0.0% |
| HexNAc(2)Hex(5)Fuc(1) | 0.2% | 0.0% | 0.0% | 0.0% | 0.1% | 0.0% | 0.0% | 0.0% | 0.0% | 0.0% |
| HexNAc(2)Hex(6) | 0.2% | 0.1% | 0.1% | 0.0% | 0.2% | 0.0% | 0.0% | 0.0% | 0.0% | 0.0% |
| HexNAc(2)Hex(8) | 0.2% | 0.0% | 0.0% | 0.0% | 0.0% | 0.0% | 0.0% | 0.0% | 0.0% | 0.0% |
| HexNAc(3)Hex(3)Fuc(1) | 0.1% | 0.0% | 0.0% | 0.0% | 0.0% | 0.0% | 0.0% | 0.0% | 0.0% | 0.0% |
| HexNAc(3)Hex(4)Fuc(1)NeuAc(1) | 0.1% | 0.2% | 0.1% | 0.1% | 0.5% | 0.5% | 0.2% | 0.2% | 0.1% | 0.0% |
| HexNAc(3)Hex(6)Fuc(1)NeuAc(1) | 0.0% | 0.0% | 0.0% | 0.0% | 0.0% | 0.0% | 0.0% | 0.0% | 0.0% | 0.0% |
| HexNAc(4)Hex(5)Fuc(1)NeuAc(2) | 0.0% | 0.0% | 0.0% | 0.0% | 0.0% | 0.0% | 0.0% | 0.0% | 0.0% | 0.0% |
| HexNAc(6)Hex(6) | 0.0% | 0.0% | 0.0% | 0.1% | 0.0% | 0.0% | 0.0% | 0.0% | 0.0% | 0.0% |
| SUM | 100.0% | 100.0% | 100.0% | 100.0% | 100.0% | 100.0% | 100.0% | 100.0% | 100.0% | 100.0% |

#### ii) Correlation analysis

| | PM-MPO | MM-MPO | BN-MPO | SN-MPO | PMN-nMPO |
|---|---|---|---|---|---|
| PM-MPO | | 0.9838 | 0.9798 | 0.9317 | 0.9516 |
| MM-MPO | | | 0.9966 | 0.9470 | 0.9605 |
| BN-MPO | | | | 0.9697 | 0.9782 |
| SN-MPO | | | | | 0.9955 |
| PMN-nMPO | | | | | |

| | Az-MPO | Sp-MPO | Ge-MPO | Se/Pl-MPO | PMN-nMPO | Dg-MPO |
|---|---|---|---|---|---|---|
| Az-MPO | | 0.9697 | 0.9693 | 0.9684 | 0.9879 | 0.9852 |
| Sp-MPO | | | 0.9689 | 0.9427 | 0.9875 | 0.9594 |
| Ge-MPO | | | | 0.9878 | 0.9660 | 0.9779 |
| Se/Pl-MPO | | | | | 0.9427 | 0.9710 |
| PMN-nMPO | | | | | | 0.9767 |
| Dg-MPO | | | | | | |

#### F All five MPO *N*-glycosylation sites

##### Correlation analysis

| | PM-MPO | MM-MPO | BN-MPO | SN-MPO | PMN-nMPO |
|---|---|---|---|---|---|
| PM-MPO | | 0.9737 | 0.9688 | 0.9531 | 0.9322 |
| MM-MPO | | | 0.9854 | 0.9704 | 0.9419 |
| BN-MPO | | | | 0.9885 | 0.9563 |
| SN-MPO | | | | | 0.9652 |
| PMN-nMPO | | | | | |

| | Az-MPO | Sp-MPO | Ge-MPO | Se/Pl-MPO | PMN-nMPO | Dg-MPO |
|---|---|---|---|---|---|---|
| Az-MPO | | 0.9905 | 0.9870 | 0.9715 | 0.9876 | 0.9321 |
| Sp-MPO | | | 0.9852 | 0.9606 | 0.9882 | 0.9163 |
| Ge-MPO | | | | 0.9785 | 0.9821 | 0.9348 |
| Se/Pl-MPO | | | | | 0.9538 | 0.9141 |
| PMN-nMPO | | | | | | 0.9106 |
| Dg-MPO | | | | | | |

##### Nomenclature

| | | | |
|---|---|---|---|
| PM | Promyelocyte | Az | Azurophilic granule |
| MM | Metamyelocyte | Sp | Specific granule |
| BN | Band neutrophil | Ge | Gelatinase granule |
| SN | Neutrophil with segmented nuclei | Se/Pl | Secretory vesicle/plasma membrane |
| PMN | Polymorphonuclear cell (neutrophil) | Dg | Degranulated |
| nMPO | Neutrophil MPO (unfractionated) | | |

##### Key to colour coding

Glycoprofiling heat scale

Low Rel abundance High

Correlation analysis heat scale

Low Correlation coefficient (r) High

**Supplementary Figure S5. A-E**) Glycoprofiling (i) and correlation analysis (ii) of the five *N*-glycosylation sites of MPO of maturing (PM-, MM-, BN-, and SN-MPO), mature/resting (PMN-nMPO), granule-separated (Az-, Sp-, Ge- and Se/Pl-MPO), and activated (Dg-MPO) neutrophils. See **Figure 3D** for comparison of top 5 glycoforms of each site and **Supplementary Table S10** for all quantitative data. **F**) Correlation analysis of all five MPO *N*-glycosylation sites. The correlation coefficients (r) have consistently been provided. See key for colour coding of the quantitative glycoprofiling data, the correlation analyses, and the used nomenclature.

#### Supplementary Figure S6

A Sequence alignment of human peroxidases

```
MPO    1 MGVPFFSSLRCMVDLGPCWAGGLTAEMKLLLALAGLLAILAT----PQPSEGAAPAVLGE
EPO    1 --------------------------MHLLPALAGVLATLVL----AQPCEGTDPASPGA
LPO    1 --------------------------MRVLLHLPALLASLIL----LQAAASTTR--AQT
TPO    1 --------------------------MRALAVLSVTLVMACTEAFFPFISRGK-ELLWGK

MPO   57 VDTSLVLSSMEEAKQLVDKAYK-ERRESIKQRLRSGSASPMELLSYFKQPVAATRTAVRA
EPO   31 VETSVLRDCIAEAKLLVDAAYN-WTQKSIKQRLRSGSASPMDLLSYFKQPVAATRTVVRA
LPO   29 TRTSAISDTVSQAKVQVNKAFL-DSRTRLKTAMSSETPTSRQLSEYLKHAKGRTRTAIRN
TPO   34 PEESRVSSVLEESKRLVDTAMYATMQRNLKKR---GILSPAQLLSFSKLPEPTSGVIARA

MPO  116 ADYLHVALDLLERKLRSLWRRPFNVTDVLTPA-QLNVLSKSSGCAYQDVGVTCPE---QD
EPO   90 ADYMHVALGLLEEKLQPQRSGPFNVTDVLTEP-QLRLLSQASGCALRDQAERCS-----D
LPO   88 GQVWEESLKRLRQKASL--------TNVTDPSLDLTSLSLEVGCGAPAPVVRCDP---CS
TPO   91 AEIMETSIQAMKRKVNLKTQQSQHPTDALSED-LLSIIANMSGCLPYMLPPKCPNTCLAN

MPO  172 KYRTITGMCNNRRSPTLGASNRAFVRWLPAEYEDGFSLPYGWTPGVKRNGFPVALARAVS
EPO  144 KYRTITGRCNNKRRPLLGASNQALARWLPAEYEDGLSLPFGWTPSRRRNGFLLPLVRAVS
LPO  137 PYRTITGDCNNRRKPALGAANRALARWLPAEYEDGLSLPFGWTPGKTRNGFPLPLAREVS
TPO  150 KYRPITGACNNRDHPRWGASNTALARWLPPVYEDGFSQPRGWNPGFLYNGFPLPPVREVT

MPO  232 NEIVRFPTDQLTPDQERSLMFMQWGQLLDHDLDFTPEPAARASFVTGVNCETSCVQQPPC
EPO  204 NQIVRFPNERLTSDRGRALMFMQWGQFIDHDLDFSPESPARVAFTAGVDCERTCAQLPPC
LPO  197 NKIVGYLNEEGVLDQNRSLLFMQWGQIVDHDLDFAPDTELGSSEYSKAQCDEYCIQGDNC
TPO  210 RHVIQVSNEVVTDDDRYSDLLMAWGQYIDHDIAFTPQSTSKAAFGGGADCQMTCENQNPC
                                        *
MPO  292 FPLKIPPNDPRIKNQADCIPFFRSCPACPGSN----------ITIRNQINALTSFVDASM
EPO  264 FPIKIPPNDPRIKNQRDCIPFFRSAPSCPQNK----------NRVRNQINALTSFVDASM
LPO  257 FPIMFPPNDPKAGTQGKCMPFFRAGFVCPTPPY--------KSLAREQINALTSFLDASF
TPO  270 FPIQLPE-EARPAAGTACLPFYRSSAACGTGDQGALFGNLSTANPRQQMNGLTSFLDAST
                      *                                       *
MPO  342 VYGSEEPLARNLRNMSNQLGLLAVNQRFQDNGRALLPFDNLH-DDPCL---LTNRSARIP
EPO  314 VYGSEVSLSLRLRNRTNYLGLLAINQRFQDNGRALLPFDNLH-DDPCL---LTNRSARIP
LPO  309 VYSSEPSLASRLRNLSSPLGLMAVNQEVSDHGLPYLPYDSKK-PSPCE---FINTTARVP
TPO  329 VYGSSPALERQLRNWTSAEGLLRVHARLRDSGRAYLPFVPPRAPAACAPEPGIPGETRGP

MPO  398 CFLAGDTRSSEMPELTSMHTLLLREHNRLATELKSLNPRWDGERLYQEARKIVGAMVQII
EPO  370 CFLAGDTRSTETPKLAAMHTLFMREHNRLATELRRLNPRWNGDKLYNEARKIMGAMVQII
LPO  365 CFLAGDSRASEHILLATSHTLFLREHNRLARELKRLNPQWDGEKLYQEARKILGAFVQII
TPO  389 CFLAGDGRASEVPSLTALHTLWLREHNRLAAALKALNAHWSADAVYQEARKVVGALHQII
                                  *
MPO  458 TYRDYLPLVLGPTAMRKYLPTYRSYNDSVDPRIANVFT-NAFRYGHTLIQPFMFRLDNRY
EPO  430 TYRDFLPLVLGKARARRTLGHYRGYCSNVDPRVANVFT-LAFRFGHTMLQPFMFRLDSQY
LPO  425 TFRDYLPIILGD-HMQKWIPPYQGYSESVDPRISNVFT-FAFRFGHLEVPSSMFRLDENY
TPO  449 TLRDYIPRILGPEAFQQYVGPYEGYDSTANPTVSNVFSTAAFRFGHATIHPLVRRLDASF

MPO  517 QPMEPNPRVPLSRVFFASWRVVLEGGIDPILRGLMATPAKLNRQNQIAVDEIRERLFEQV
EPO  489 RASAPNSHVPLSSAFFASWRIVYEGGIDPILRGLMATPAKLNRQDAMLVDELRDRLFRQV
LPO  483 QPWGPEPELPLHTLFFNTWRMVKDGGIDPLVRGLLAKKSKLMKQNKMMTGELRNKLFQPT
TPO  509 QEHPDLPGLWLHQAFFSPWTLLRGGGLDPLIRGLLARPAKLQVQDQLMNEELTERLFVLS

MPO  577 MRI-GLDLPALNMQRSRDHGLPGYNAWRRFCGLPQPETVGQLGTVLRNLKLARKLMEQYG
EPO  549 RRI-GLDLAALNMQRSRDHGLPGYNAWRRFCGLSQPRNLAQLSRVLKNQDLARKFLNLYG
LPO  543 HRIHGFDLAAINTQRCRDHGQPGYNSWRAFCDLSQPQTLEELNTVLKSKMLAKKLLGLYG
TPO  569 NSS-TLDLASINLQRGRDHGLPGYNEWREFCGLPRLETPADLSTAIASRSVADKILDLYK

MPO  636 TPNNIDIWMGGVSEPLKRKGRVGPLLACIIGTQFRKLRDGDRFWWENEGVFSMQQRQALA
EPO  608 TPDNIDIWIGAIAEPLLPGARVGPLLACLFENQFRRARDGDRFWWQKRGVFTKRQRKALS
LPO  603 TPDNIDIWIGAIAEPLVERGRVGPLLACLLGKQFQQIRDGDRFWWENPGVFTNEQKDSLQ
TPO  628 HPDNIDVWLGGLAENFLPRARTGPLFACLIGKQMKALRDGDWFWWENSHVFTDAQRRELE
                                          *
MPO  696 QISLPRIICDNTGITTVSKNNIFMSNSYPRDFVNCSTLPALNLASWRE------------
EPO  668 RISLSRIICDNTGITTVSR-DIFRANIYPRGFVNCSRIPRLNLSAWRG------------
LPO  663 KMSFSRLVCDNTRITKVPR-DPFWANSYPYDFVDCSAIDKLDLSPWAS------------
TPO  688 KHSLSRVICDNTGLTRVPM-DAFQVGKFPEDFESCDSITGMNLEAWRETFPQDDKCGFPE

MPO  744 -A-S--------------------------------------------------------
EPO  715 ---T--------------------------------------------------------
LPO  710 -VKN--------------------------------------------------------
TPO  747 SVENGDFVHCEESGRRVLVYSCRHGYELQGREQLTCTQEGWDFQPPLCKDVNECADGAHP

MPO      ------------------------------------------------------------
EPO      ------------------------------------------------------------
LPO      ------------------------------------------------------------
TPO  807 PCHASARCRNTKGGFQCLCADPYELGDDGRTCVDSGRLPRVTWISMSLAALLIGGFAGLT

MPO      ------------------------------------------------------------
EPO      ------------------------------------------------------------
LPO      ------------------------------------------------------------
TPO  867 STVICRWTRTGTKSTLPISETGGGTPELRCGKHQAVGTSPQRAAAQDSEQESAGMEGRDT

MPO      -------
EPO      -------
LPO      -------
TPO  927 HRLPRAL
```

Black Conserved residue
White Conserved property (basic, acidic…)
* Sequon (glycosylated Asn highlighted)

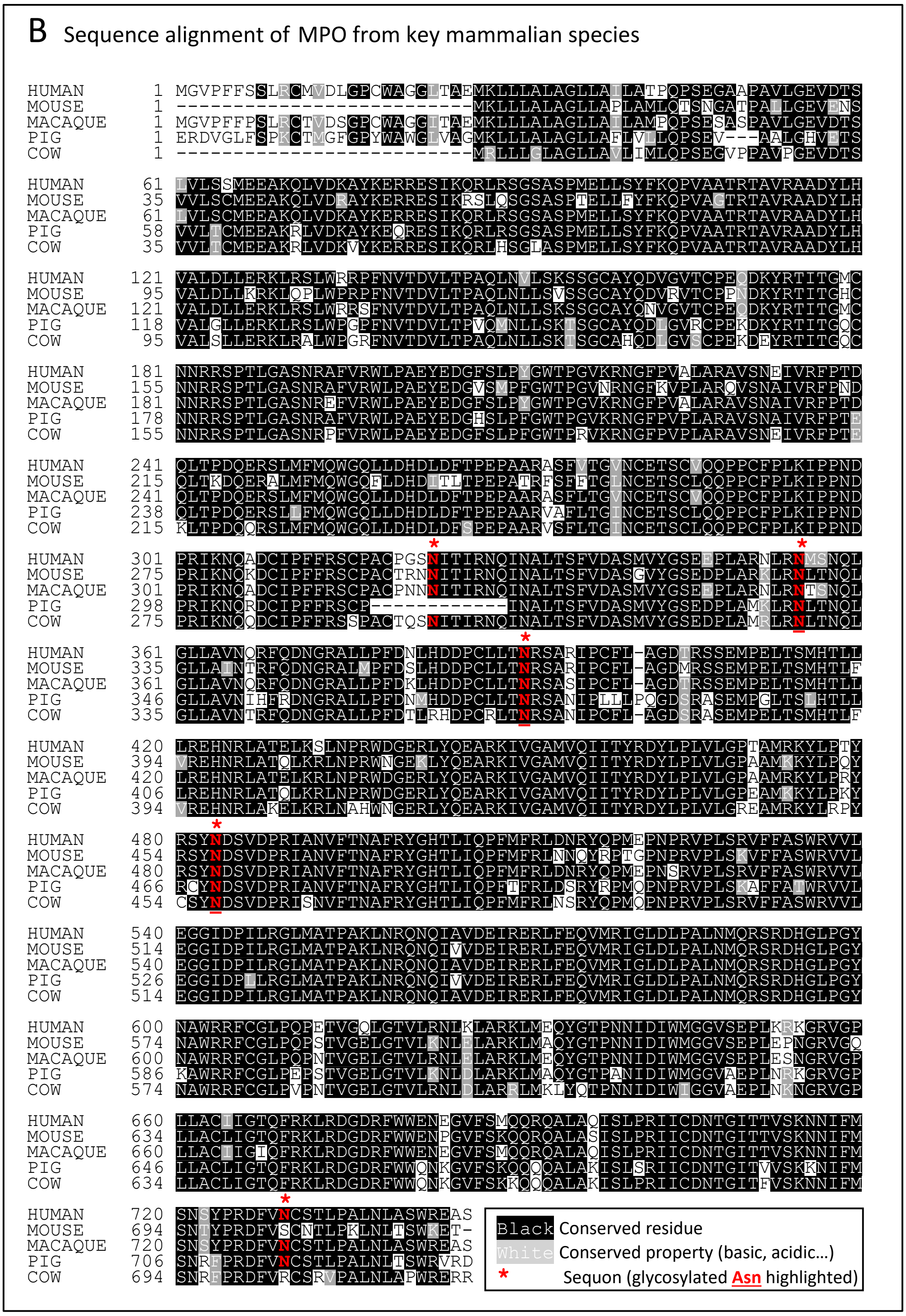


**Supplementary Figure S6**. Sequence alignment of **A**) the human peroxidases including human MPO (UniProtKB, P05164), human eosinophil peroxidase (EPO, P11678), human lactoperoxidase (LPO, P80025) and human thyroid peroxidase (TPO, P07202) and **B**) MPO from key mammalian species including mouse MPO (P11247), macaque MPO (F7BAA9), porcine MPO (K7GRV6) and bovine MPO (A6QPT4). Asn355 and Asn391 of human MPO are conserved sequons in both analyses. See key for details.

#### Supplementary Figure S7

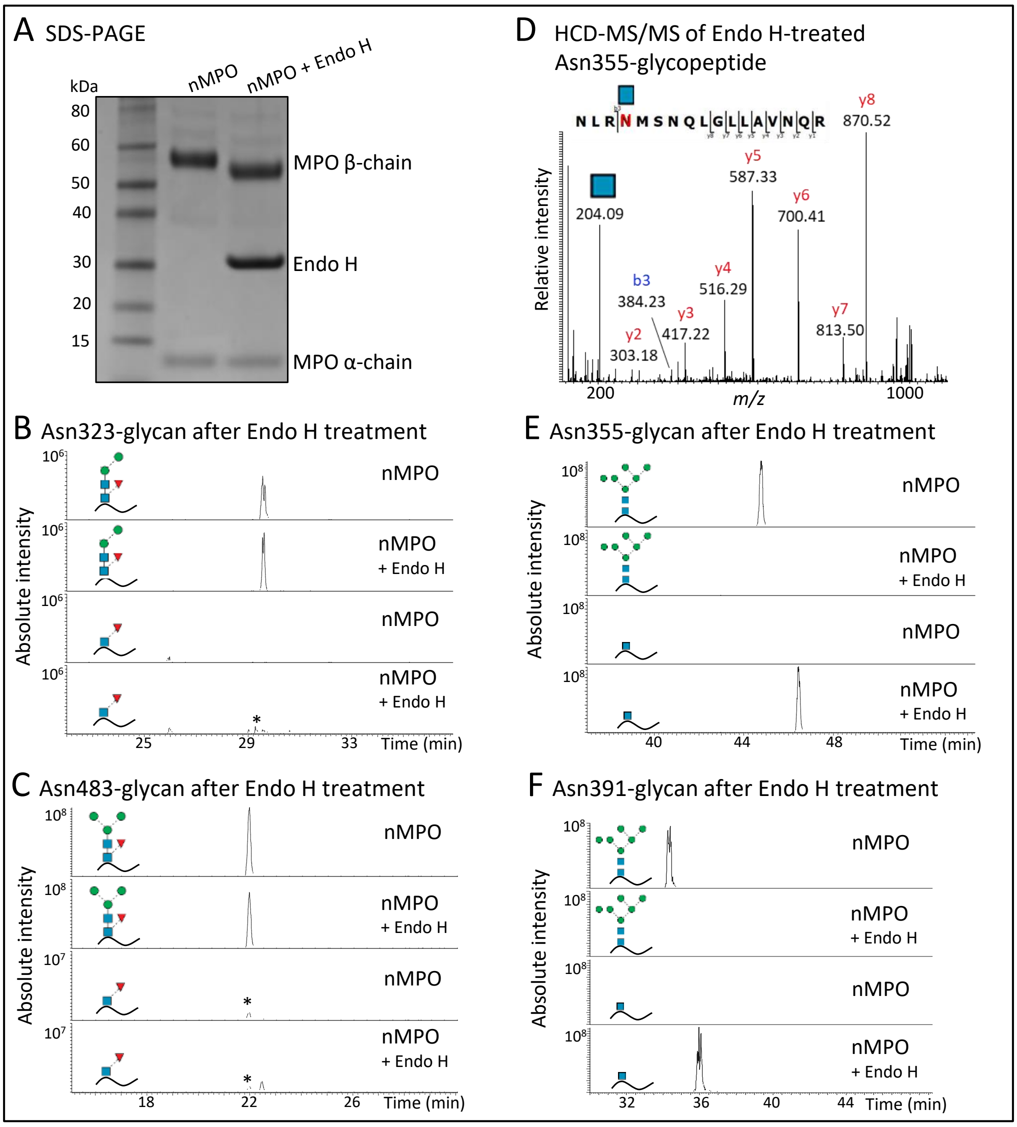


**Supplementary Figure S7**. Site-specific manipulation of the Asn355- and Asn391-glycosylation by Endo H treatment of native nMPO as demonstrated using **A**) SDS-PAGE and **B-F**) LC-MS/MS glycopeptide data. The extracted ion chromatograms (EICs) confirmed that the processed *N*-glycans decorating Asn323 (B) and Asn483 (C) were unaffected by the Endo H treatment, whilst the oligomannosidic *N*-glycans of Asn355 (E) and Asn391 (F) were fully converted to GlcNAcβAsn by Endo H. *Artificial *in-source* fragment ion. The largely unoccupied Asn729 was not assessed in this analysis. The specific glycopeptides monitored using EICs have been indicated with cartoons, see **Supplementary Table S7** for details of the monitored glycopeptides. **D**) Example of an annotated HCD-MS/MS spectrum of the Endo H-treated Asn355-glycopeptide (*m/z* 721.04, 3+) demonstrating that Endo H generates GlcNAcβAsn residues at the two oligomannose-rich sites of MPO.

**Supplementary Figure S8**

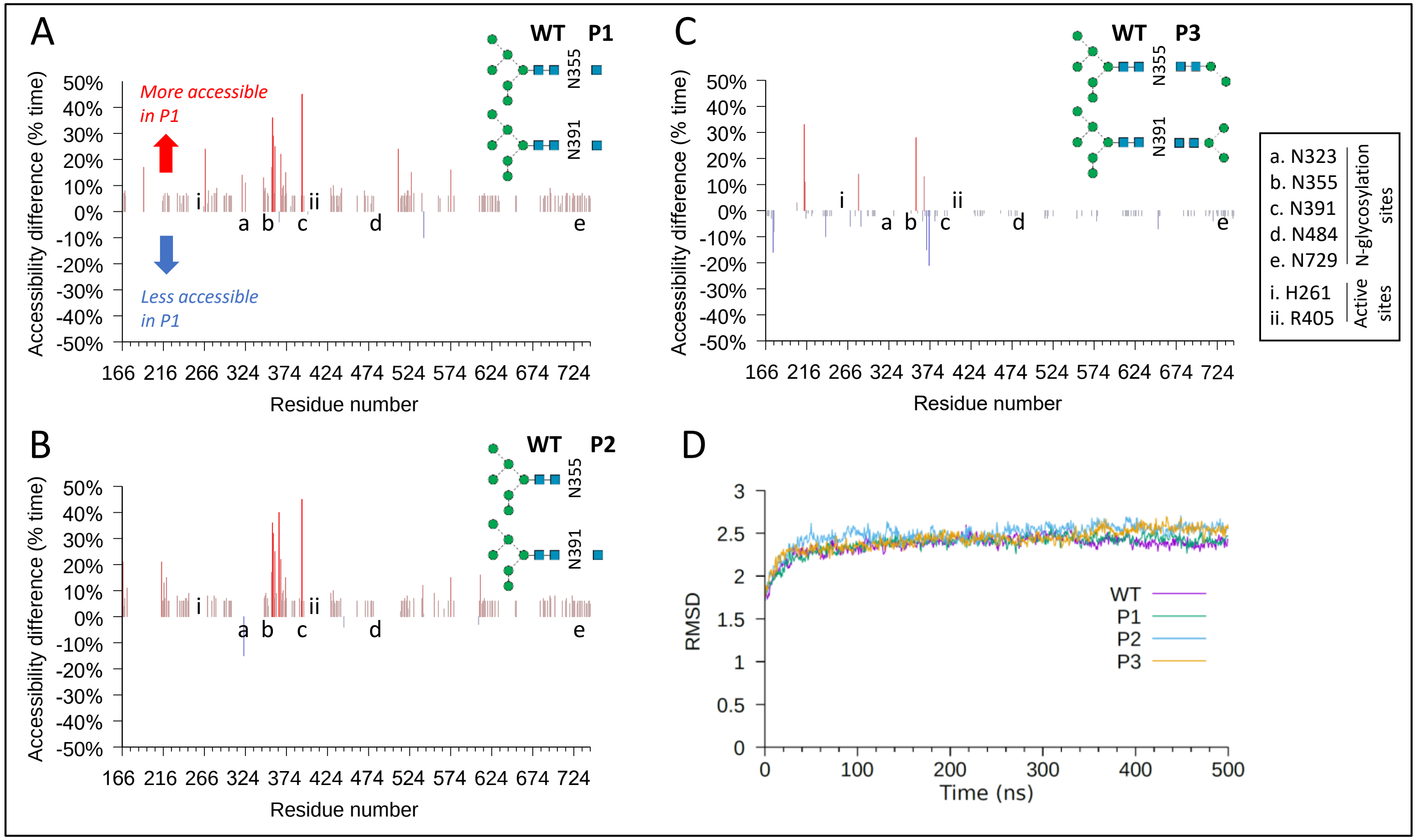


**Supplementary Figure S8.** Impact of Asn355-/Asn391-glycans on the MPO structure as determined using MD simulations. Differential per-residue accessibility of the WT MPO versus **A**) P1, **B**) P2 and **C**) P3 plotted as the proportional duration (% time) difference ($p < 0.05$) was observed during the 500 ns MD simulation. P1 and P2 (but not P3) show significant enhanced accessibility along most of the polypeptide chain relative to WT. See insert for key to important MPO residues and **Supplementary Table S4** for structures and nomenclature of simulated MPO glycoforms. **D**) Assessment of the global movement of the MPO polypeptide chain using RMSD monitored during the 500 ns MD simulations.

#### Supplementary Figure S9

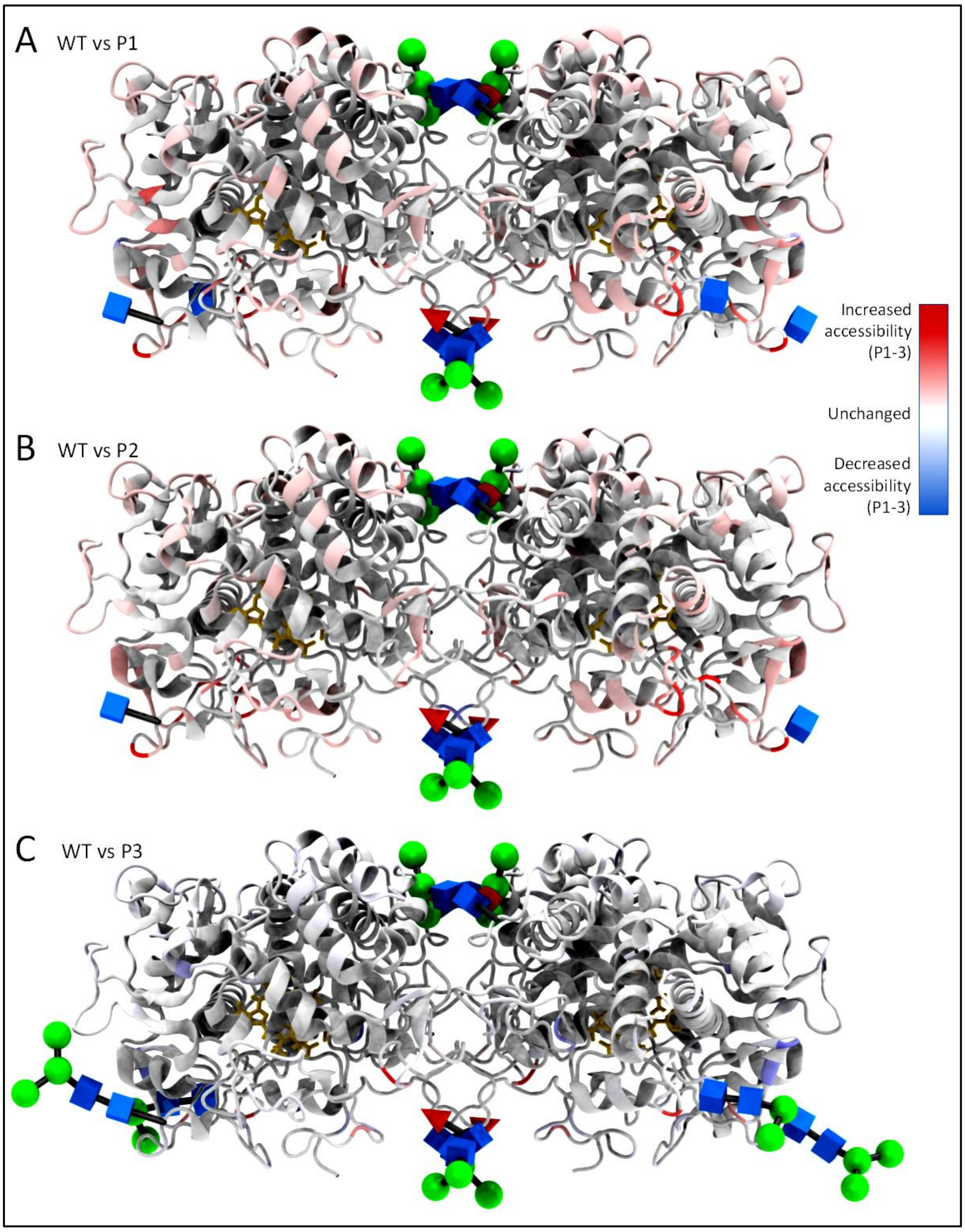


**Supplementary Figure S9**. Differential accessibility of the WT (resembling nMPO) and **A**) P1 (Endo H-treated nMPO i.e. Asn355-/Asn391-GlcNAc glycophenotype), **B**) P2 (minimal Asn355-/Asn391-glycophenotype elevated in Se/Pl-MPO), and **C**) P3 (semi-truncated Asn355/Asn391-glycophenotype of MPO) depicted on diprotomeric MPO (PDBID, 1D2V) as monitored over the 500 ns MD simulation. Unaffected residues are in grey, see also **Supplementary Figure S8** for other representation of data. Residues with reduced (blue) or increased (red) accessibilities ($p < 0.05$) are mapped with an intensity shading reflecting the proportional duration of the differential accessibility, see key. Darker residue intensity signifies a longer duration of differential accessibility during the simulations. The glycans are shown as 3D-SNFG symbols. The heme moiety is represented in yellow sticks.

#### References used in SI


Abrahams, J.L., Campbell, M.P., and Packer, N.H. (2018). Building a PGC-LC-MS N-glycan retention library and elution mapping resource. Glycoconjugate Journal *35*, 15-29.

Ashwood, C., Lin, C.-H., Thaysen-Andersen, M., and Packer, N.H. (2018). Discrimination of Isomers of Released N- and O-Glycans Using Diagnostic Product Ions in Negative Ion PGC-LC-ESI-MS/MS. Journal of The American Society for Mass Spectrometry *29*, 1194-1209.

Bansal, R., Elgundi, Z., Goodchild, S.C., Care, A., Lord, M.S., Rodger, A., and Sunna, A. (2020). The Effect of Oligomerization on A Solid-Binding Peptide Binding to Silica-Based Materials. Nanomaterials (Basel) *10*.

Case, D.A., Babin, V., Berryman, J.T., Betz, R.M., Cai, Q., Cerutti, D.S., T.E. Cheatham, I., Darden, T.A., Duke, R.E., Gohlke, H.*, et al.* (2014). AMBER 14 (University of California, San Francisco).

Clemmensen, S.N., Udby, L., and Borregaard, N. (2014). Subcellular Fractionation of Human Neutrophils and Analysis of Subcellular Markers. In Neutrophil Methods and Protocols, M.T. Quinn, and F.R. DeLeo, eds. (Totowa, NJ: Humana Press), pp. 53-76.

Cox, J., and Mann, M. (2008). MaxQuant enables high peptide identification rates, individualized ppb-range mass accuracies and proteome-wide protein quantification. Nature biotechnology *26*, 1367-1372.

Darden, T., York, D., and Pedersen, L. (1993). Particle mesh Ewald: An N· log (N) method for Ewald sums in large systems. The Journal of chemical physics *98*, 10089.

Dypbukt, J.M., Bishop, C., Brooks, W.M., Thong, B., Eriksson, H., and Kettle, A.J. (2005). A sensitive and selective assay for chloramine production by myeloperoxidase. Free Radical Biology and Medicine *39*, 1468-1477.

Everest-Dass, A.V., Abrahams, J.L., Kolarich, D., Packer, N.H., and Campbell, M.P. (2013). Structural Feature Ions for Distinguishing N- and O-Linked Glycan Isomers by LC-ESI-IT MS/MS. Journal of The American Society for Mass Spectrometry *24*, 895-906.

Feuk-Lagerstedt, E., Movitz, C., Pellme, S., Dahlgren, C., and Karlsson, A. (2007). Lipid raft proteome of the human neutrophil azurophil granule. Proteomics *7*, 194-205.

Fiedler, T.J., Davey, C.A., and Fenna, R.E. (2000). X-ray Crystal Structure and Characterization of Halide-binding Sites of Human Myeloperoxidase at 1.8 Å Resolution. Journal of Biological Chemistry *275*, 11964-11971.

Gault, J., Donlan, J.A.C., Liko, I., Hopper, J.T.S., Gupta, K., Housden, N.G., Struwe, W.B., Marty, M.T., Mize, T., Bechara, C.*, et al.* (2016). High-resolution mass spectrometry of small molecules bound to membrane proteins. Nature Methods *13*, 333.

Gotz, A.W., Williamson, M.J., Xu, D., Poole, D., Le Grand, S., and Walker, R.C. (2012). Routine Microsecond Molecular Dynamics Simulations with AMBER on GPUs. 1. Generalized Born. J Chem Theory Comput *8*, 1542-1555.

Hall, V., Sklepari, M., and Rodger, A. (2014). Protein secondary structure prediction from circular dichroism spectra using a self-organizing map with concentration correction. Chirality *26*, 471-482.

Hernández, H., and Robinson, C.V. (2007). Determining the stoichiometry and interactions of macromolecular assemblies from mass spectrometry. Nature Protocols *2*, 715-726.

Hinneburg, H., Chatterjee, S., Schirmeister, F., Nguyen-Khuong, T., Packer, N.H., Rapp, E., and Thaysen-Andersen, M. (2019). Post-Column Make-Up Flow (PCMF) Enhances the Performance of Capillary-Flow PGC-LC-MS/MS-Based Glycomics. Analytical chemistry *91*, 4559-4567.

Hoogendijk, A.J., Pourfarzad, F., Aarts, C.E.M., Tool, A.T.J., Hiemstra, I.H., Grassi, L., Frontini, M., Meijer, A.B., van den Biggelaar, M., and Kuijpers, T.W. (2019). Dynamic Transcriptome-Proteome Correlation Networks Reveal Human Myeloid Differentiation and Neutrophil-Specific Programming. Cell Rep *29*, 2505-2519 e2504.

Hubber, S.J., and Thornton, J.M. (1993). NACCESS computer program. Department of Biochemistry and Molecular Biology, University College, London.

Humphrey, W., Dalke, A., and Schulten, K. (1996). VMD: visual molecular dynamics. J Mol Graph *14*, 33-38, 27-38.

Jensen, P.H., Karlsson, N.G., Kolarich, D., and Packer, N.H. (2012). Structural analysis of N- and O-glycans released from glycoproteins. Nature Protocols *7*, 1299.

Kawahara, R., Ortega, F., Rosa-Fernandes, L., Guimarães, V., Quina, D., Nahas, W., Schwämmle, V., Srougi, M., Leite, K.R.M., Thaysen-Andersen, M.*, et al.* (2018). Distinct urinary glycoprotein signatures in prostate cancer patients. Oncotarget *9*, 33077-33097.

Kirschner, K.N., Yongye, A.B., Tschampel, S.M., Gonzalez-Outeirino, J., Daniels, C.R., Foley, B.L., and Woods, R.J. (2008). GLYCAM06: a generalizable biomolecular force field. Carbohydrates. J Comput Chem *29*, 622-655.

Lees, J.G., Miles, A.J., Wien, F., and Wallace, B.A. (2006). A reference database for circular dichroism spectroscopy covering fold and secondary structure space. Bioinformatics *22*, 1955-1962.

Loziuk, P.L., Hecht, E.S., and Muddiman, D.C. (2017). N-linked glycosite profiling and use of Skyline as a platform for characterization and relative quantification of glycans in differentiating xylem of Populus trichocarpa. Analytical and bioanalytical chemistry *409*, 487-497.

Marty, M.T., Baldwin, A.J., Marklund, E.G., Hochberg, G.K.A., Benesch, J.L.P., and Robinson, C.V. (2015). Bayesian Deconvolution of Mass and Ion Mobility Spectra: From Binary Interactions to Polydisperse Ensembles. Analytical chemistry *87*, 4370-4376.

Nauseef, W.M. (2014). Isolation of Human Neutrophils from Venous Blood. In Neutrophil Methods and Protocols, M.T. Quinn, and F.R. DeLeo, eds. (Totowa, NJ: Humana Press), pp. 13-18.

Nivedha, A.K., Makeneni, S., Foley, B.L., Tessier, M.B., and Woods, R.J. (2014). Importance of ligand conformational energies in carbohydrate docking: Sorting the wheat from the chaff. J Comput Chem *35*, 526-539.

Petrescu, A.J., Milac, A.L., Petrescu, S.M., Dwek, R.A., and Wormald, M.R. (2004). Statistical analysis of the protein environment of N-glycosylation sites: implications for occupancy, structure, and folding. Glycobiology *14*, 103-114.

Rørvig, S., Østergaard, O., Heegaard, N.H.H., and Borregaard, N. (2013). Proteome profiling of human neutrophil granule subsets, secretory vesicles, and cell membrane: correlation with transcriptome profiling of neutrophil precursors. J Leukocyte Biol *94*, 711-721.

Ryckaert, J.-P., Ciccotti, G., and Berendsen, H.J.C. (1977). Numerical integration of the Cartesian Equations of Motion of a System with Constraints: Molecular Dynamics of n-Alkanes. Journal of Computational Physics *23*, 327-341.

Salomon-Ferrer, R., Götz, A.W., Poole, D., Le Grand, S., and Walker, R.C. (2013). Routine Microsecond Molecular Dynamics Simulations with AMBER on GPUs. 2. Explicit Solvent Particle Mesh Ewald. Journal of Chemical Theory and Computation *9*, 3878-3888.

Samygina, V.R., Sokolov, A.V., Bourenkov, G., Petoukhov, M.V., Pulina, M.O., Zakharova, E.T., Vasilyev, V.B., Bartunik, H., and Svergun, D.I. (2013). Ceruloplasmin: macromolecular assemblies with iron-containing acute phase proteins. PloS one *8*, e67145-e67145.

San Miguel, M., Marrington, R., Rodger, P.M., Rodger, A., and Robinson, C. (2003). An Escherichia coli twin-arginine signal peptide switches between helical and unstructured conformations depending on the hydrophobicity of the environment. Eur J Biochem *270*, 3345-3352.

Sethi, M.K., Kim, H., Park, C.K., Baker, M.S., Paik, Y.-K., Packer, N.H., Hancock, W.S., Fanayan, S., and Thaysen-Andersen, M. (2015). In-depth N-glycome profiling of paired colorectal cancer and non-tumorigenic tissues reveals cancer-, stage- and EGFR-specific protein N-glycosylation. Glycobiology *25*, 1064-1078.

Sklepari, M., Rodger, A., Reason, A., Jamshidi, S., Prokes, I., and Blindaer, C.A. (2016). Biophysical characterization of a protein for structure comparison: methods for identifying insulin structural changes. Analytical Methods *8*, 7460-7471.

Soltermann, F., Foley, E., Pagnoni, V., Galpin, M., Benesch, J., Kukura, P., and Struwe, W. (2020). Quantifying protein-protein interactions by molecular counting with mass photometry. Angew Chem Int Ed Engl.

Stadlmann, J., Pabst, M., Kolarich, D., Kunert, R., and Altmann, F. (2008). Analysis of immunoglobulin glycosylation by LC-ESI-MS of glycopeptides and oligosaccharides. Proteomics *8*, 2858-2871.

Varki, A., Cummings, R.D., Aebi, M., Packer, N.H., Seeberger, P.H., Esko, J.D., Stanley, P., Hart, G., Darvill, A., Kinoshita, T.*, et al.* (2015). Symbol Nomenclature for Graphical Representations of Glycans. Glycobiology *25*, 1323-1324.

Wang, J., Wolf, R.M., Caldwell, J.W., Kollman, P.A., and Case, D.A. (2004). Development and testing of a general amber force field. J Comput Chem *25*, 1157-1174.

Young, G., Hundt, N., Cole, D., Fineberg, A., Andrecka, J., Tyler, A., Olerinyova, A., Ansari, A., Marklund, E.G., Collier, M.P.*, et al.* (2018). Quantitative mass imaging of single biological macromolecules. Science *360*, 423-427.
