## Supplementary Data (S1-S3) for "Hyper-Truncated Glycans Augment the Activity of Neutrophil Granule Myeloperoxidase"

for

[1]*Contributed equally*

### Annotation and Fragmentation Key

- Mannose (162.0528 Da)
- Galactose (162.0528 Da)
- *N*-Acetylglucosamine (203.0794 Da)
- *N*-Acetylgalactosamine (203.0794 Da)
- *N*-Acetylneuraminic acid (291.0954 Da)
- Fucose (146.0579 Da)
- Cross-ring fragment (unspecified)
- Ac Acetyl group (42.0106 Da)
- Indicates mostly Y ions (includes oxygen of glyosidic linkage)
- Indicates mostly Z ions (excludes oxygen of glyosidic linkage)
- Reduced reducing end
- P Phosphate (79.9700 Da)

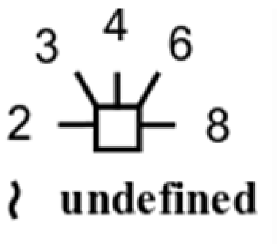


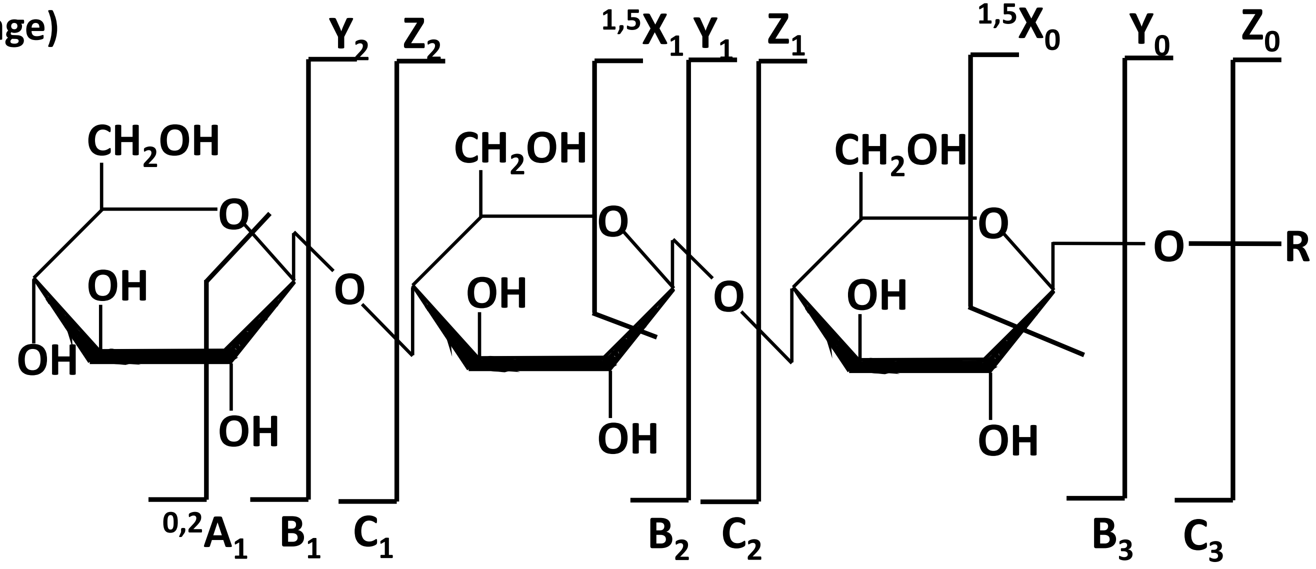


Han, L. & Costello, C.E. Mass spectrometry of glycans. *Biochemistry (Moscow)* 78, 710-720 (2013).

**Supplementary Data S1**

Manually annotated PGC-LC-ESI-CID-MS/MS (-) spectra of reduced *N*-glycans (alditols) released from nMPO

**Glycan #1**

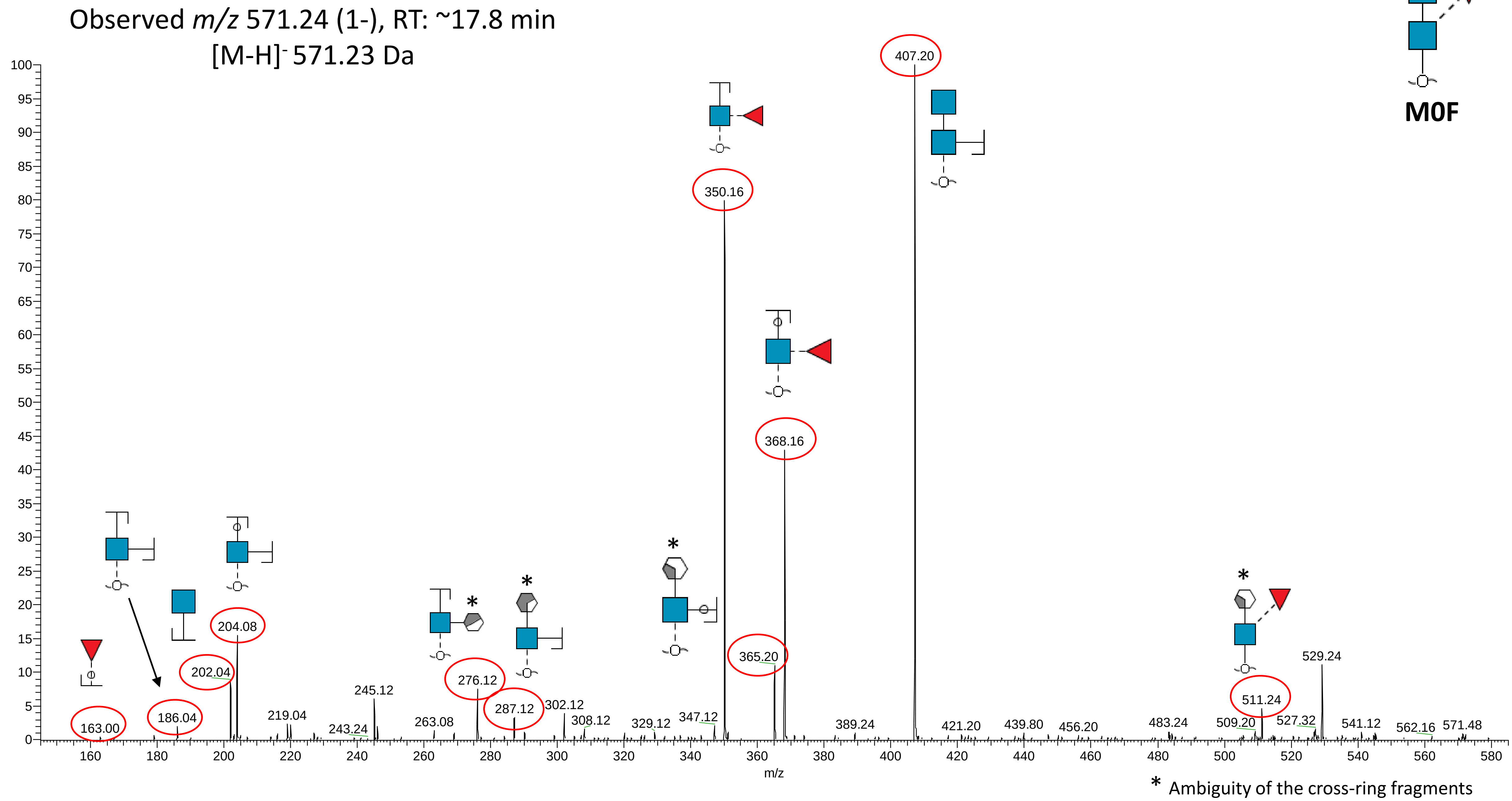
Observed *m/z* 571.24 (1-), RT: ~17.8 min
[M-H]- 571.23 Da
M0F
407.20
350.16
368.16
365.20
204.08
202.04
186.04
163.00
276.12
287.12
245.12
219.04
243.24
263.08
302.12
308.12
329.12
347.12
389.24
421.20
439.80
456.20
483.24
509.20
511.24
527.32
529.24
541.12
562.16
571.48
m/z
*
* Ambiguity of the cross-ring fragments

**Glycan #2**

Observed *m/z* 587.30 (1-), RT: ~16.9 min
$[M-H]^-$ 587.23 Da

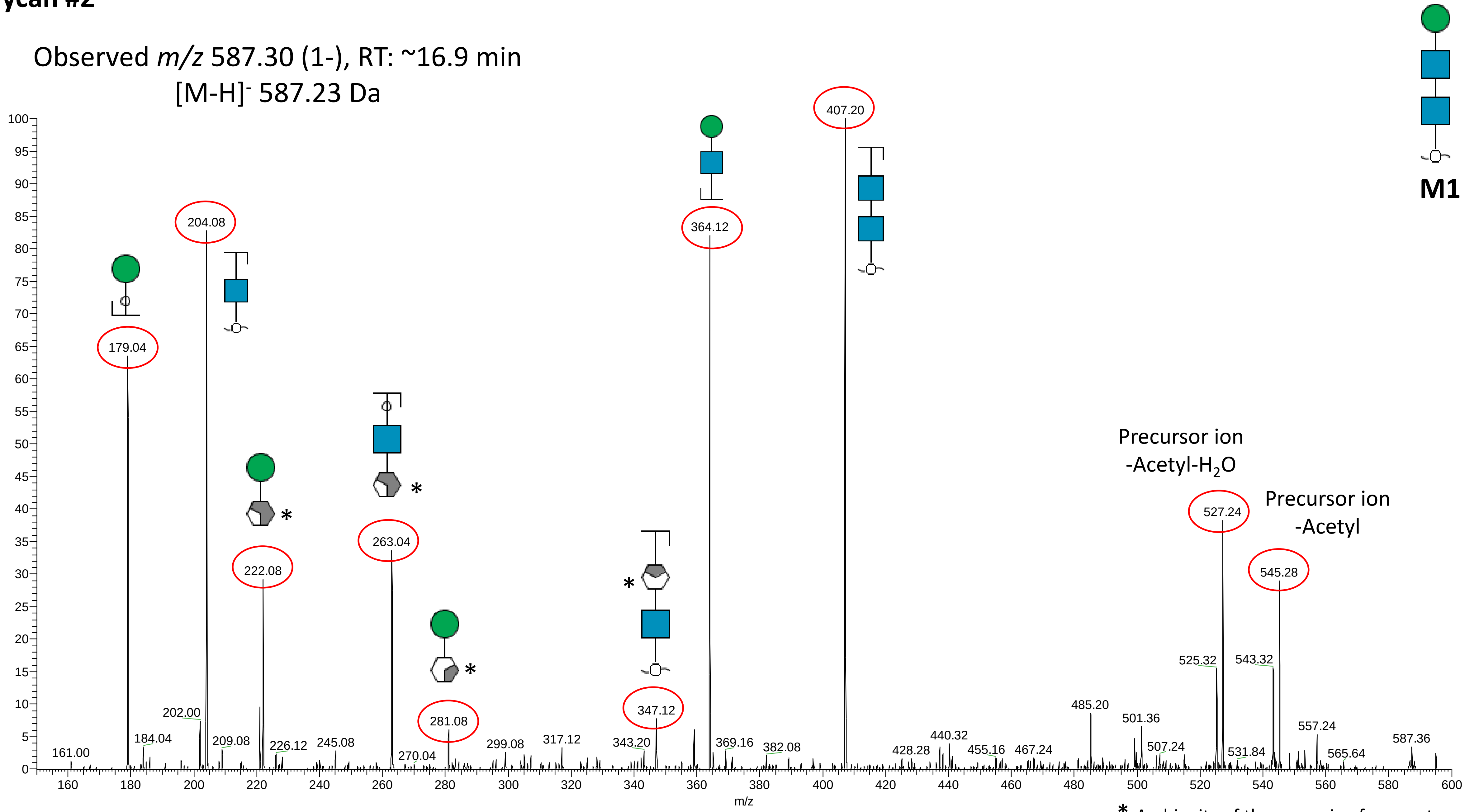


* Ambiguity of the cross-ring fragments

#### Glycan #3

Observed *m/z* 733.36 (1-), RT: ~21.1 min
$[M-H]^-$ 733.28 Da

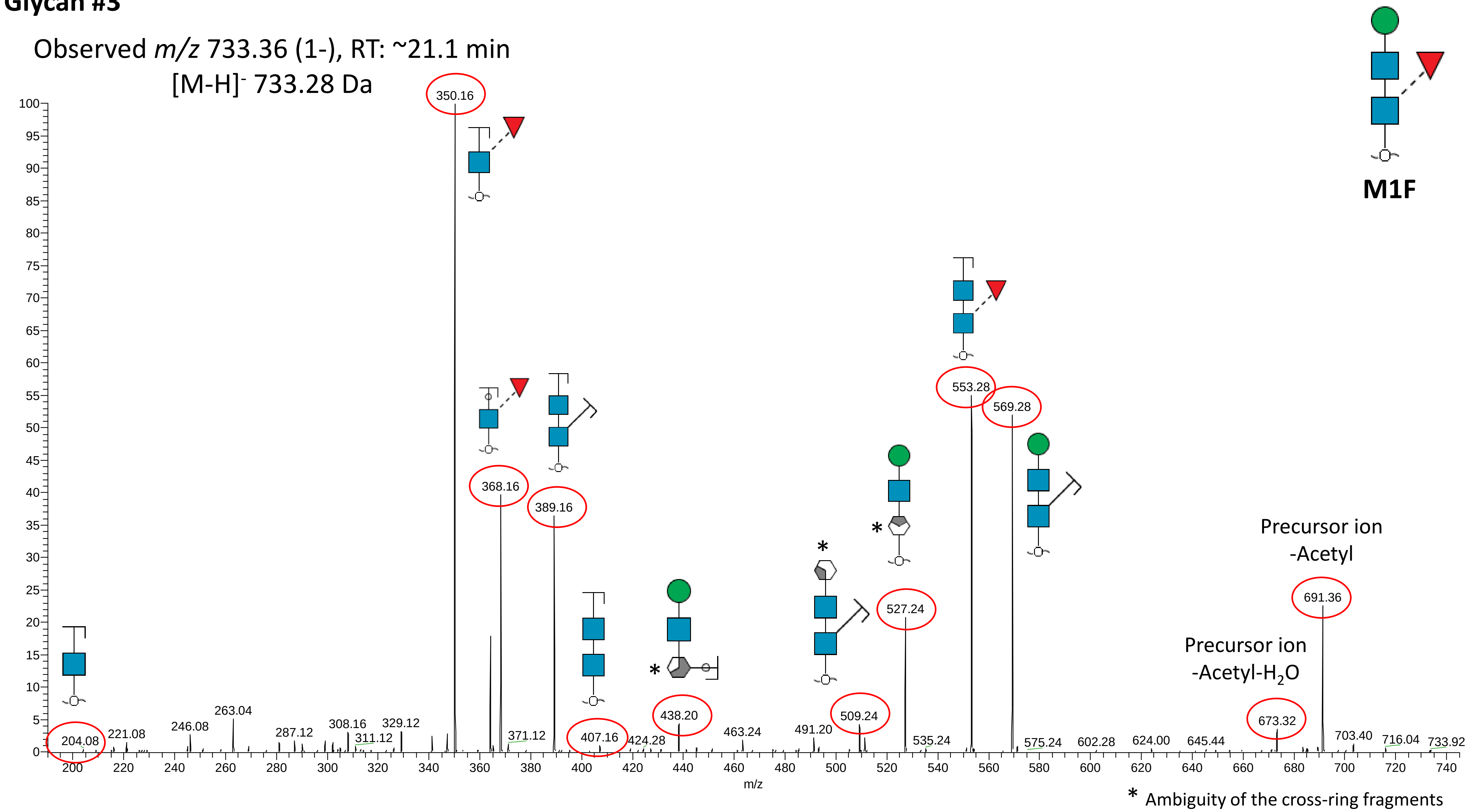

#### Glycan #4

Observed *m/z* 749.36 (1-), RT: ~18.4 min
[M-H]⁻ 749.28 Da

Note: Only one isomer was observed for M2, which is biosynthetically predicted to be α1,6-linkage isomer based on experience with neutrophil glycosylation. The α1,3-isomer, typically of lower abundance in human neutrophils, was not detected.

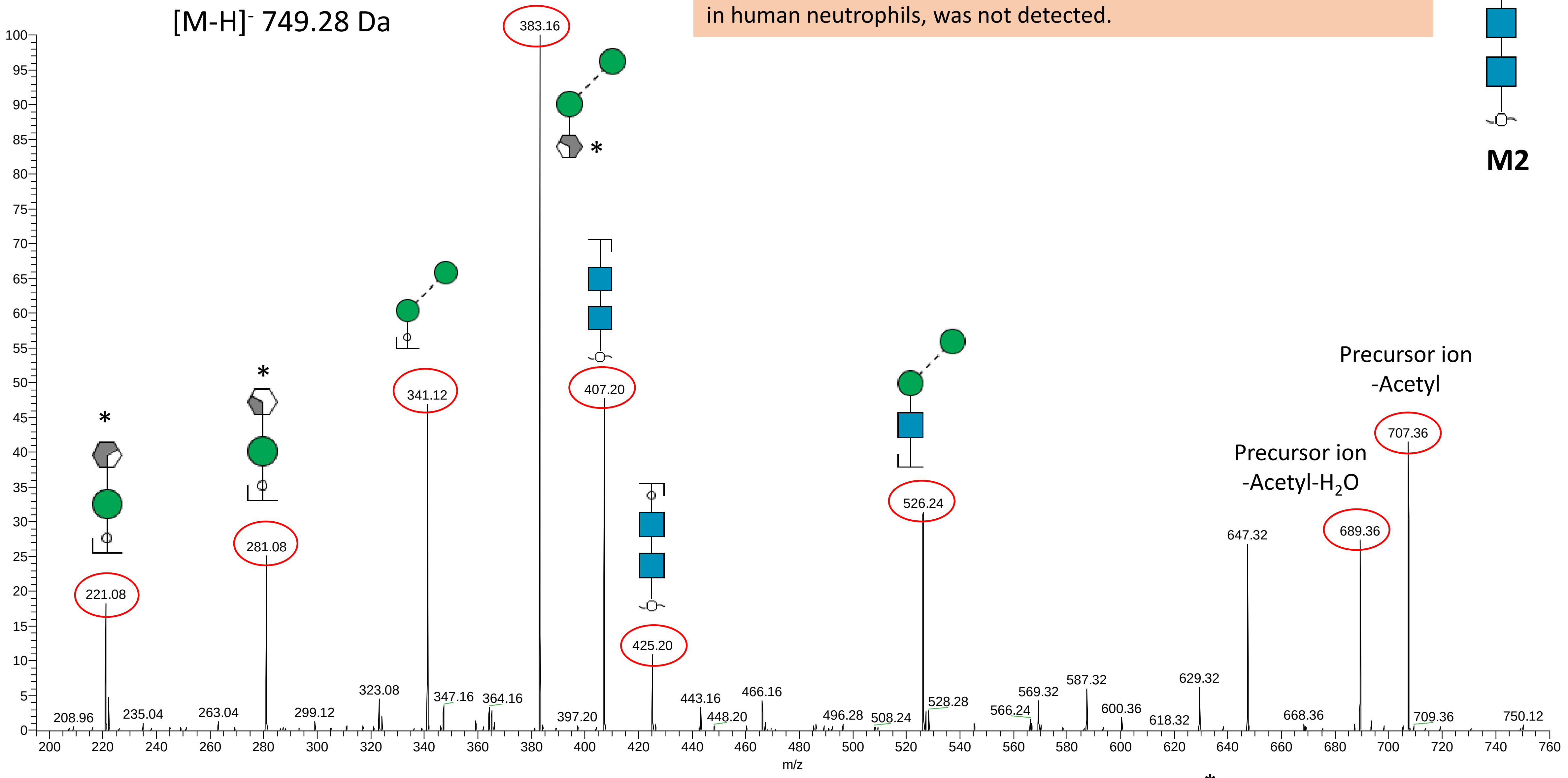


* Ambiguity of the cross-ring fragments

#### Glycan #5a

Note: Identified as the α1,6-Man isomer based on PGC LC elution time, its high abundance and experience with neutrophil *N*-glycosylation.

Observed *m/z* 895.46 (1-), RT: ~24.3 min
$[M-H]^-$ 895.34 Da

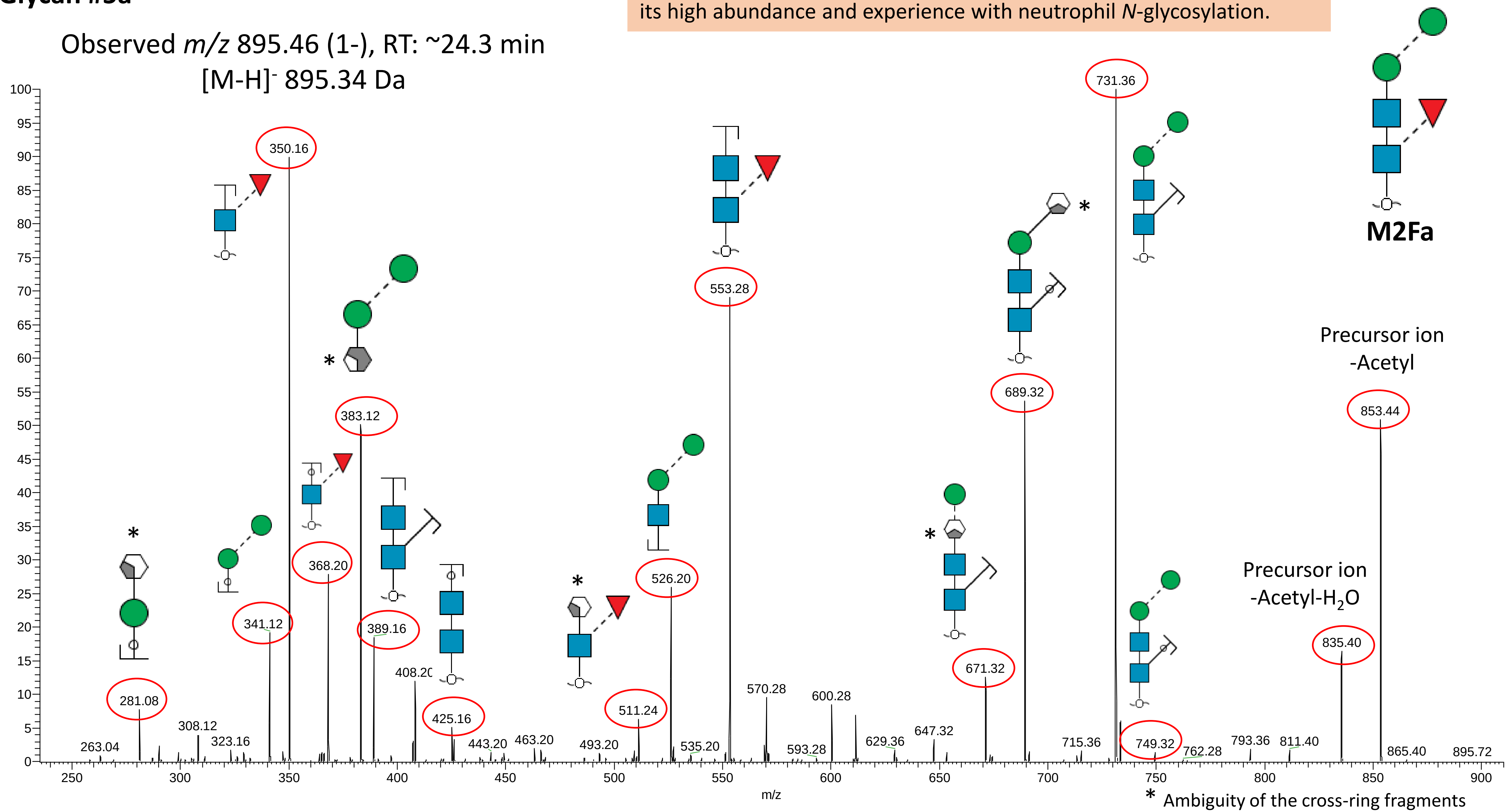

**Glycan #5b**

Note: This low abundance glycan was annotated as the α1,3-Man isomer based on PGC LC elution time and experience with neutrophil *N*-glycosylation.

Observed *m/z* 895.46 (1-), RT: ~26.4 min
[M-H]⁻ 895.34 Da

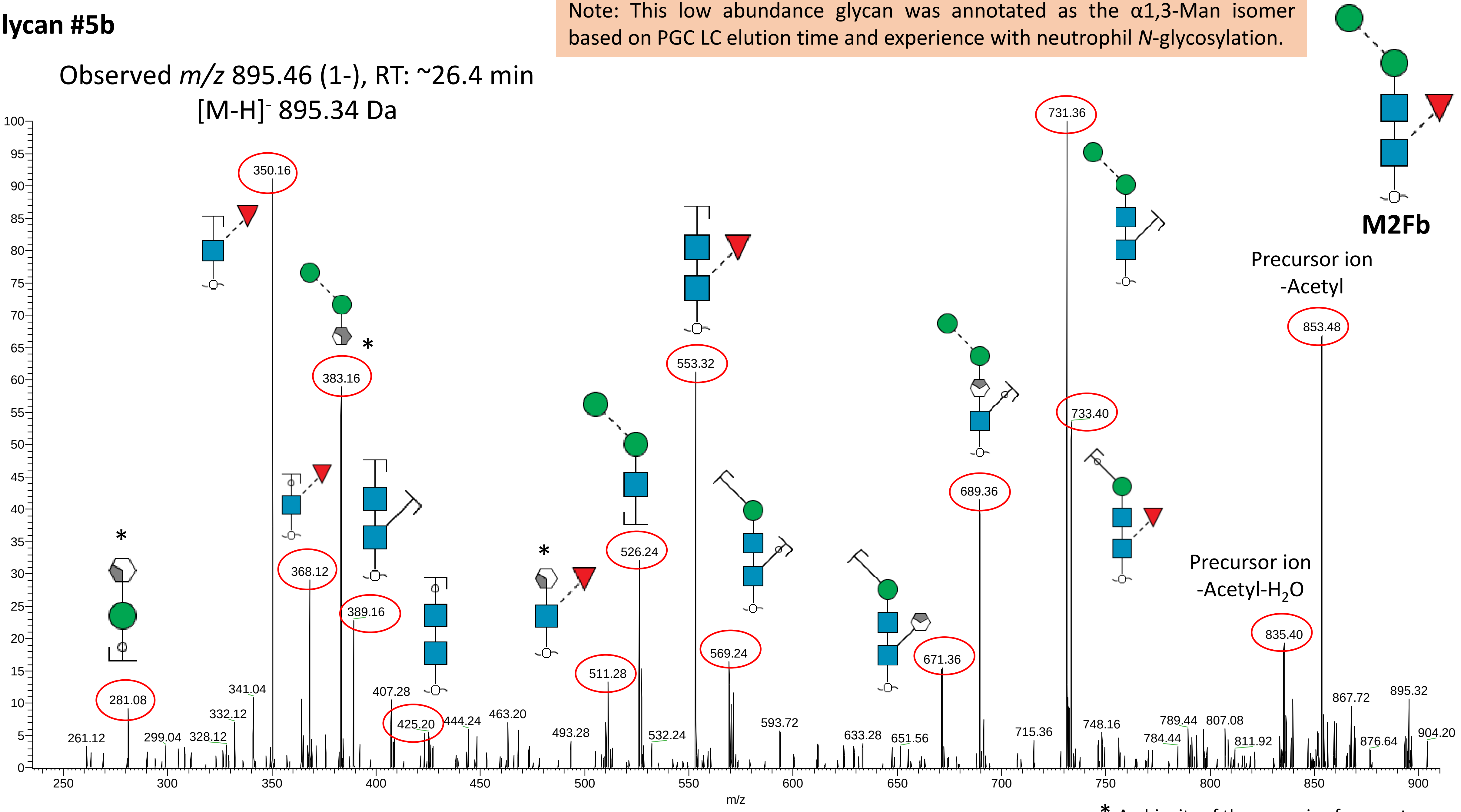


* Ambiguity of the cross-ring fragments

### Glycan #6

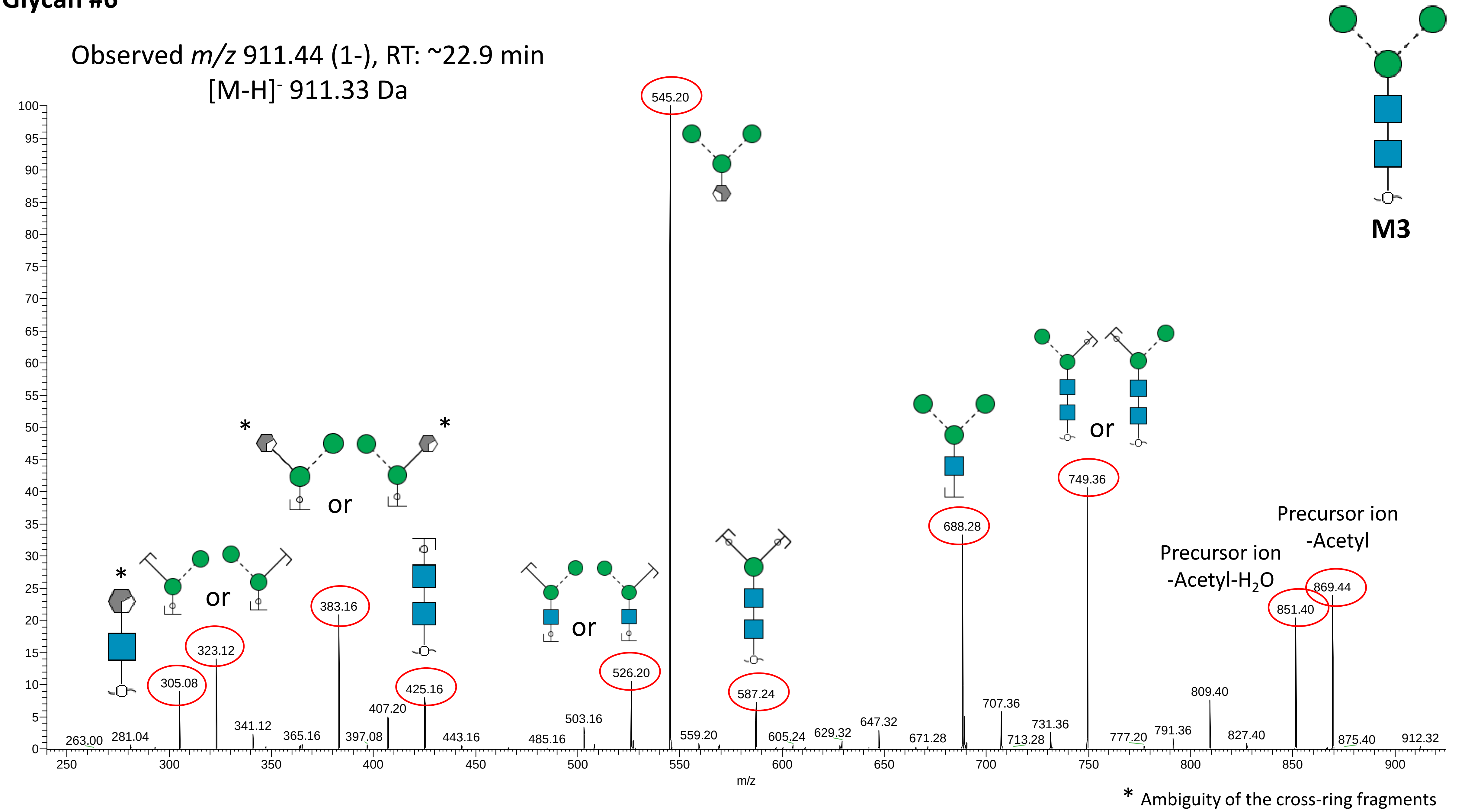

Observed m/z 911.44 (1-), RT: ~22.9 min
[M-H]- 911.33 Da
M3
545.20
305.08
323.12
383.16
425.16
526.20
587.24
688.28
749.36
851.40
869.44
Precursor ion -Acetyl-H2O
Precursor ion -Acetyl
or
m/z
* Ambiguity of the cross-ring fragments

#### Glycan #7

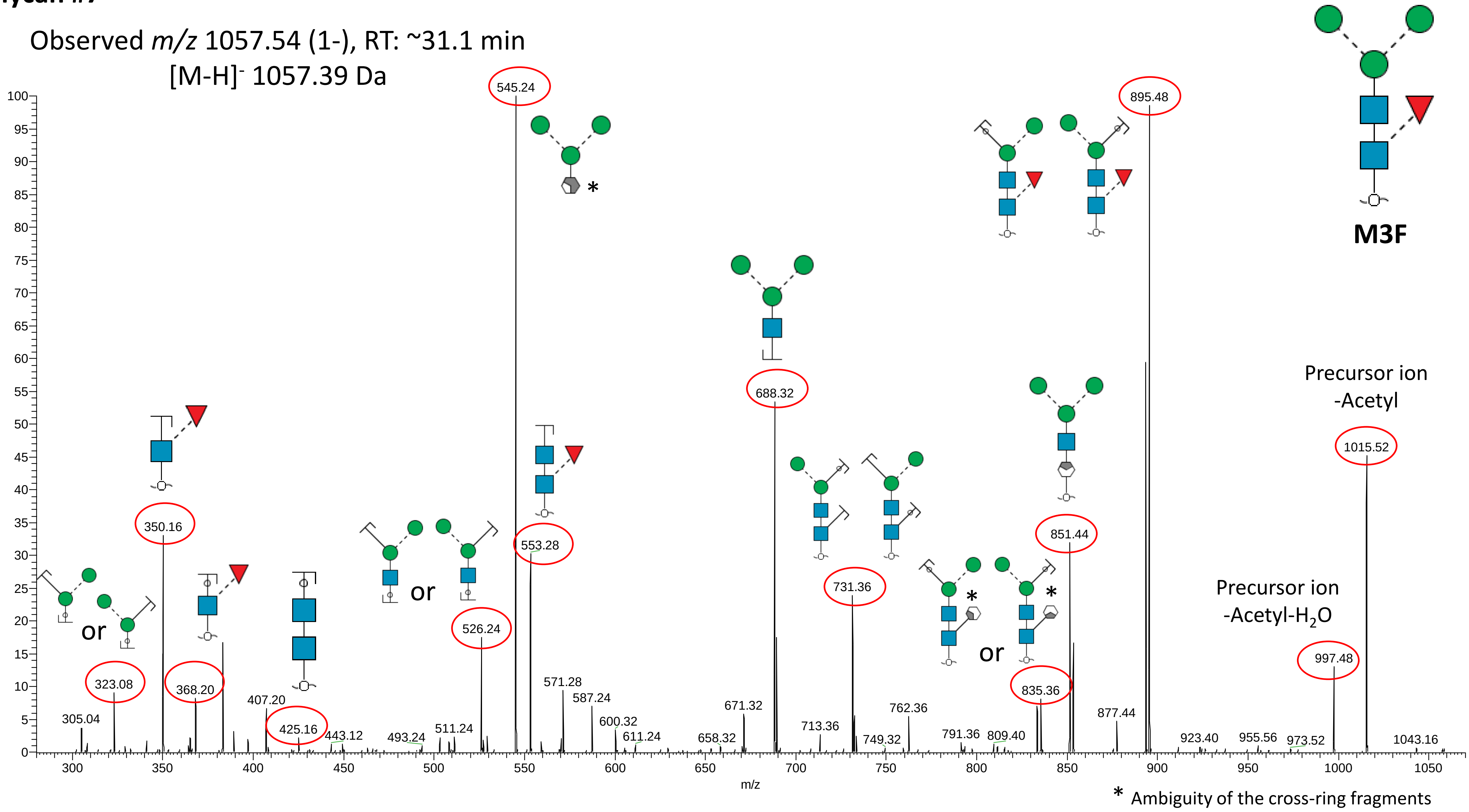
Observed m/z 1057.54 (1-), RT: ~31.1 min
[M-H]- 1057.39 Da
M3F
545.24
895.48
688.32
350.16
553.28
526.24
731.36
851.44
835.36
323.08
368.20
425.16
1015.52
997.48
Precursor ion -Acetyl
Precursor ion -Acetyl-H2O
305.04
407.20
443.12
493.24
511.24
571.28
587.24
600.32
611.24
658.32
671.32
713.36
749.32
762.36
791.36
809.40
877.44
923.40
955.56
973.52
1043.16
or
m/z
* Ambiguity of the cross-ring fragments

#### Glycan #8a

Note: The outer mannose residue has been placed on the α1,6-Man arm based on biosynthetic rules and may be predicted to be α1,6-linked based on PGC LC elution pattern (relative to the M4b isomer). However the mannosyl linkage of this residue was left unassigned since no MS/MS evidence is present to support this.

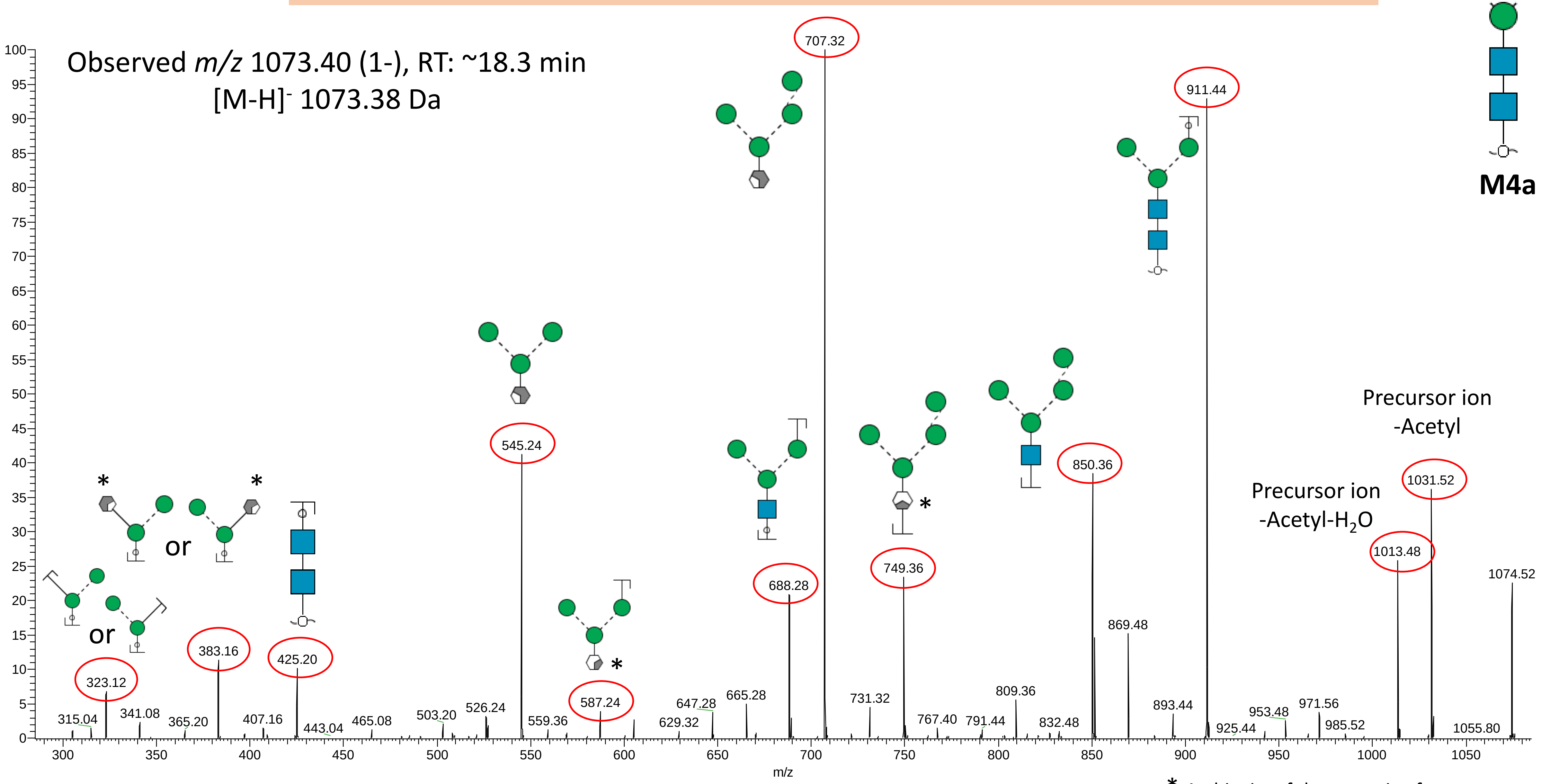


* Ambiguity of the cross-ring fragments

#### Glycan #8b

Note: The outer mannose residue has been placed on the α1,6-Man arm based on biosynthetic rules and may be predicted to be α1,3-linked based on PGC LC elution pattern (relative to the M4a isomer). However the mannosyl linkage of this residue was left unassigned since no MS/MS evidence is present to support this.

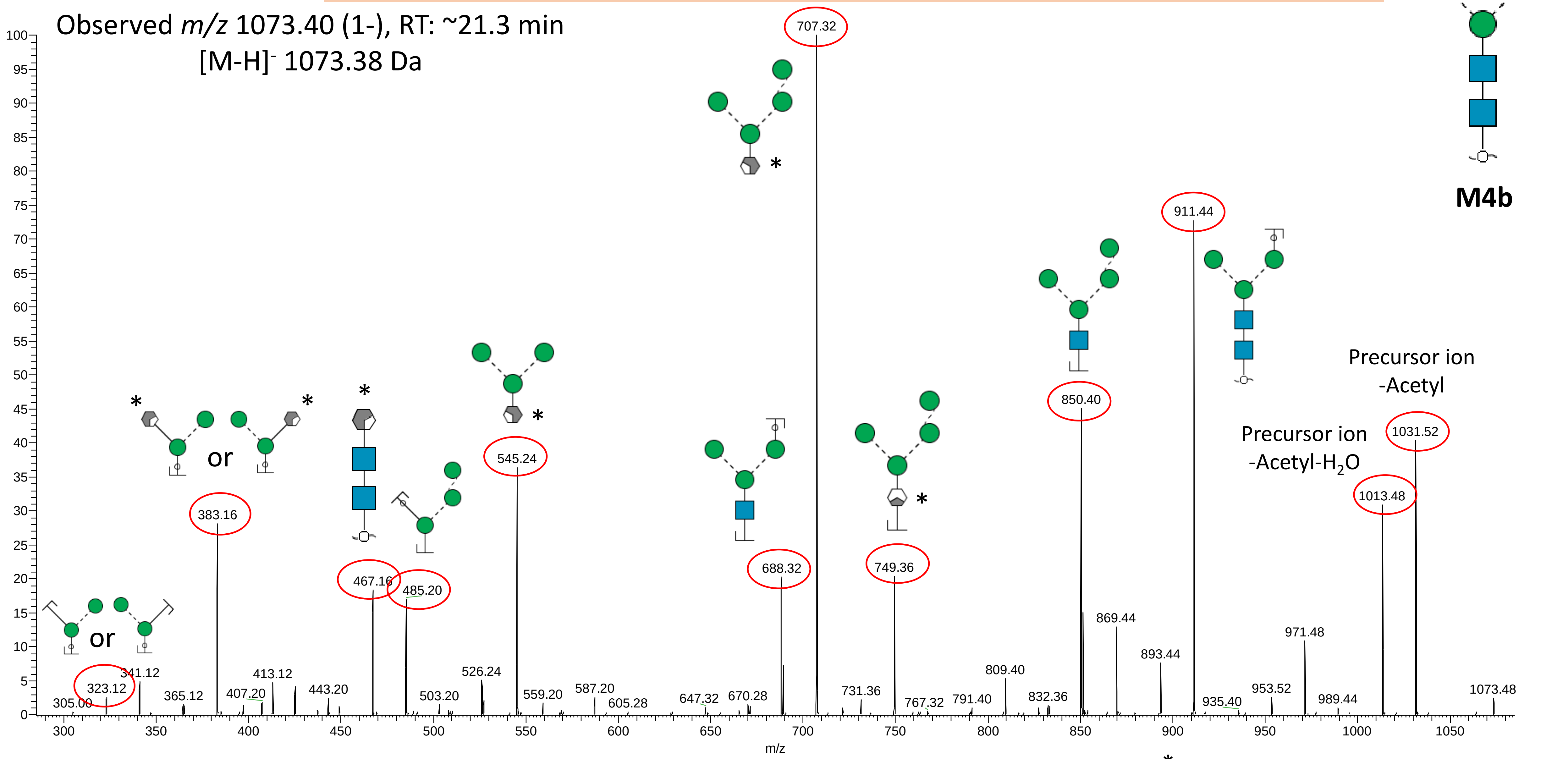


* Ambiguity of the cross-ring fragments

**Glycan #9**

Note: The outer mannose residue has been placed on the α1,6-Man arm based on biosynthetic rules. Unknown if this residue is found in a α1,3- or α1,6-linkage since no MS/MS evidence to support either so left unassigned.

Observed *m/z* 1219.44 (1-), RT: ~28.5 min

[M-H]$^-$ 1219.44 Da

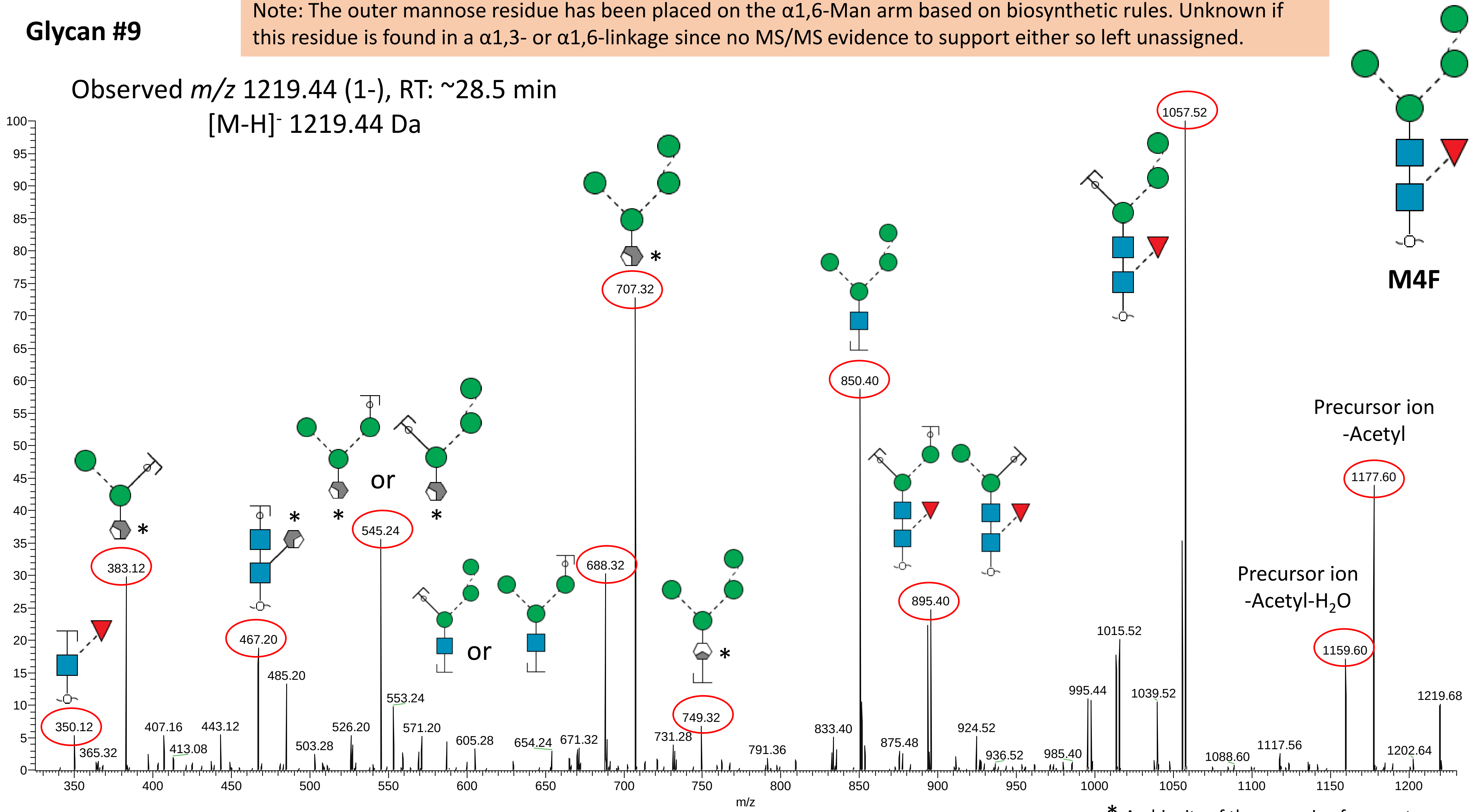


* Ambiguity of the cross-ring fragments

#### Glycan #10

Observed *m/z* 617.22 (2-), RT: ~18.1 min
[M-H]⁻ 1235.44 Da

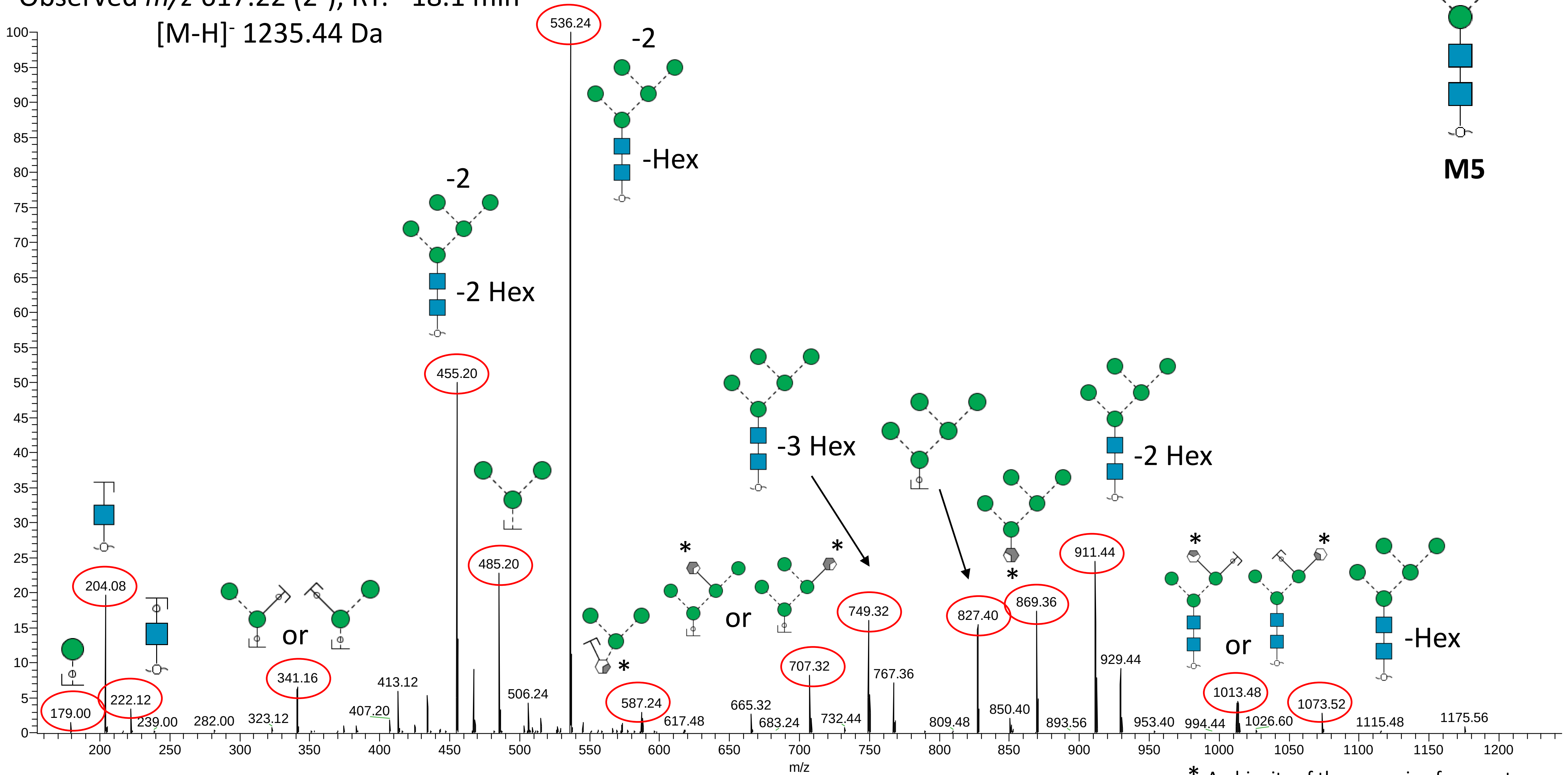


* Ambiguity of the cross-ring fragments

**Glycan #11a**

Note: Structure predicted from known biosynthetic rules, but no MS/MS support for exact topology of outer mannose residues.

Observed *m/z* 1381.62 (1-), RT: ~30.4 min

[M-H]- 1381.49 Da

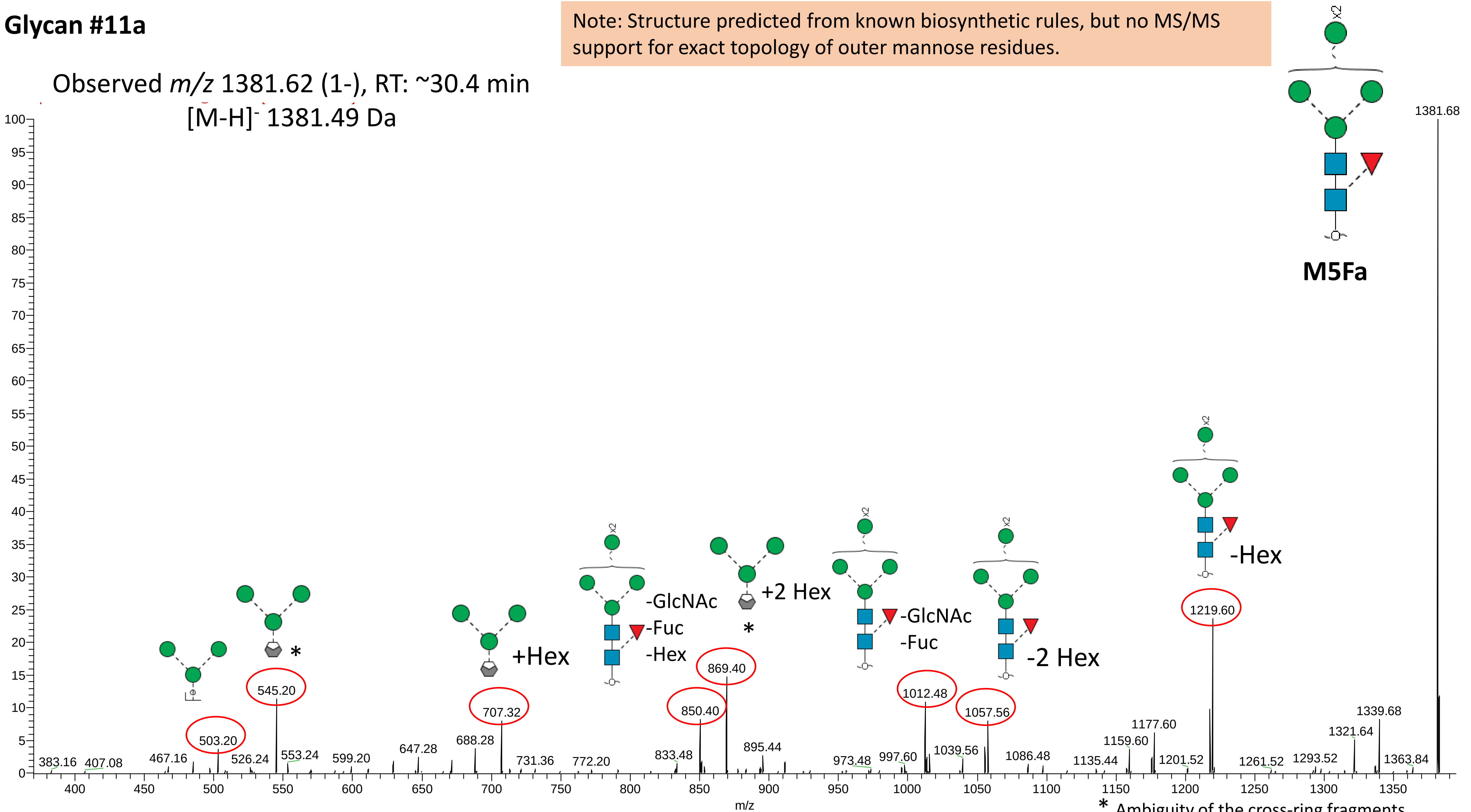


* Ambiguity of the cross-ring fragments

#### Glycan #11b

Note: Structure predicted from known biosynthetic rules, but no MS/MS support for exact topology of outer mannose residues.

Observed *m/z* 1381.62 (1-), RT: ~33.2 min
$[M-H]^-$ 1381.49 Da

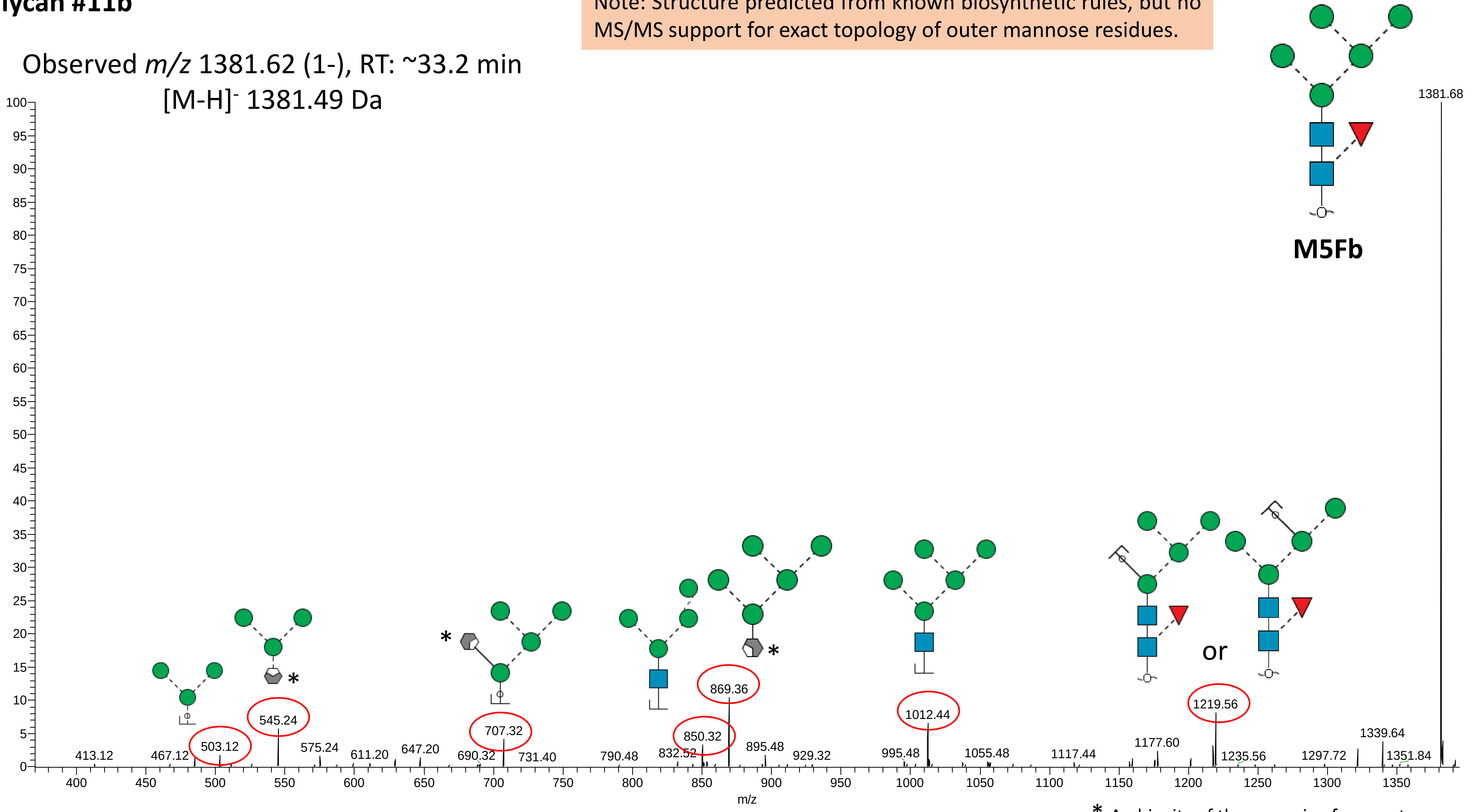


* Ambiguity of the cross-ring fragments

### Glycan #12

Observed *m/z* 698.36 (2-), RT: ~20.7 min

[M-H]⁻ 1397.49 Da

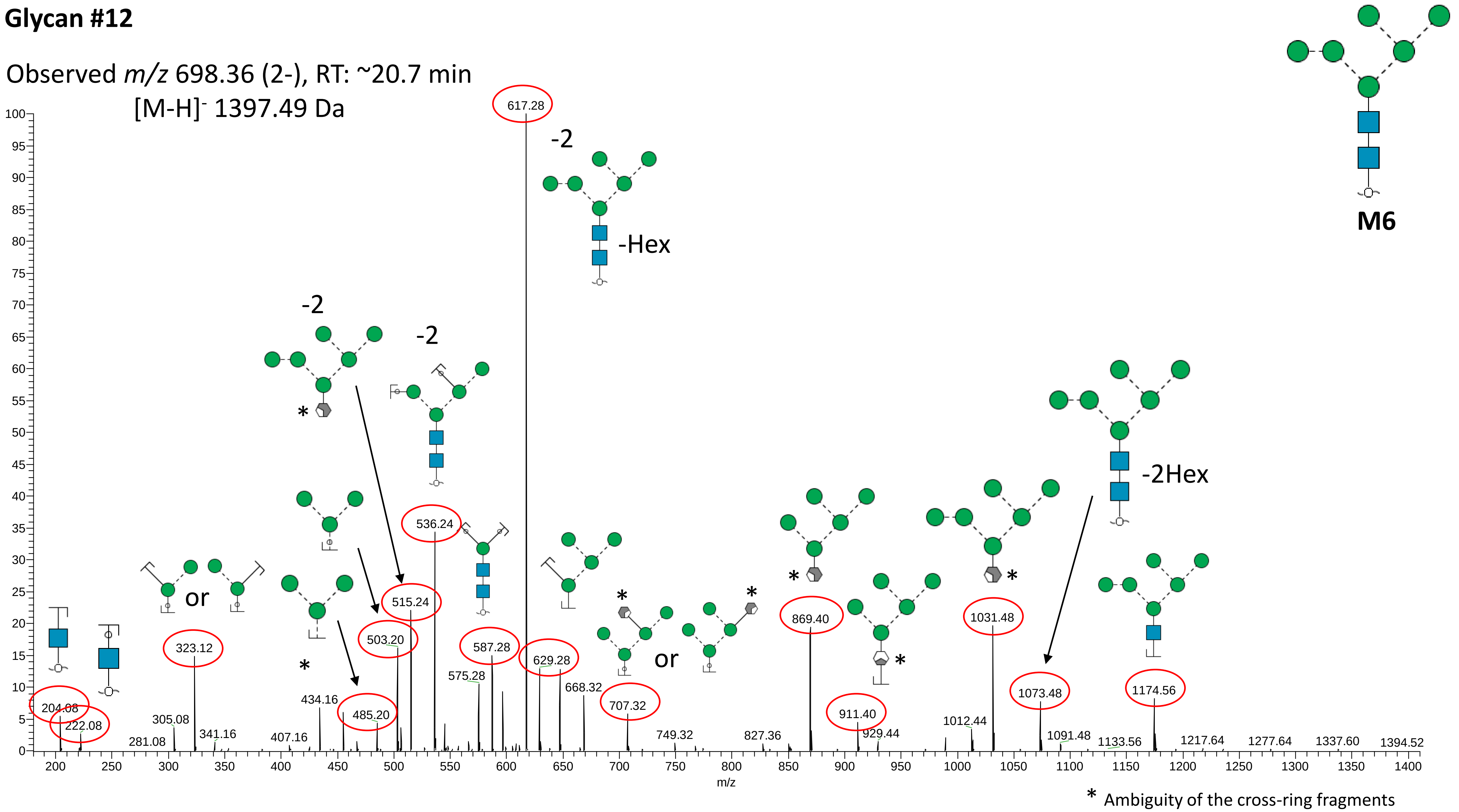

**Glycan #13**

Observed *m/z* 779.38 (2-), RT: ~20.9 min

[M-H]⁻ 1559.54 Da

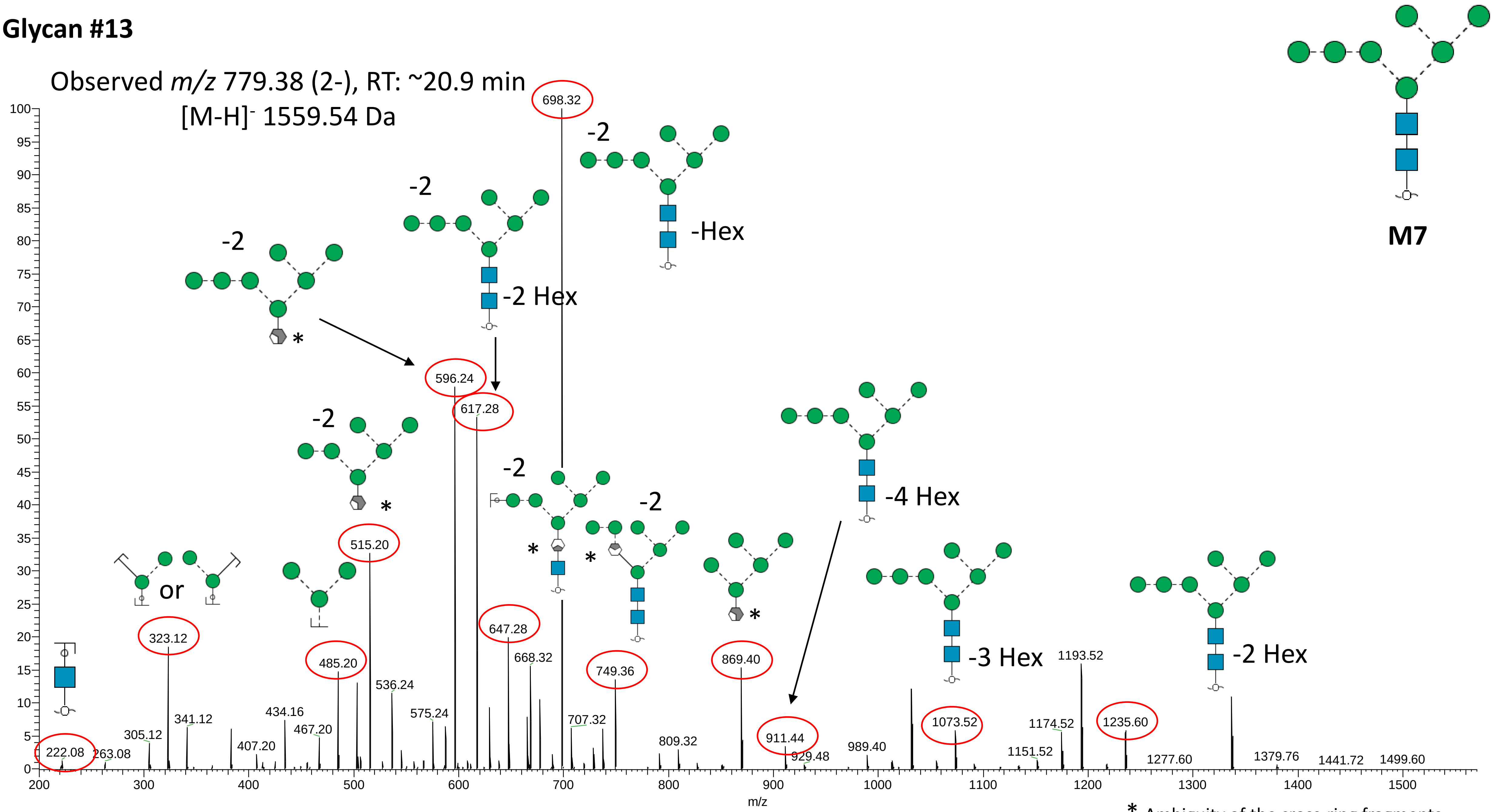


* Ambiguity of the cross-ring fragments

**Glycan #14a**

Observed *m/z* 738.23 (2-), RT: ~17.6 min

[M-H]⁻ 1477.46 Da

Note: Based on knowledge of the M6P pathway, the phosphate moiety appears on outer mannose residue, but no direct MS/MS evidence to support the exact position of the phosphate.

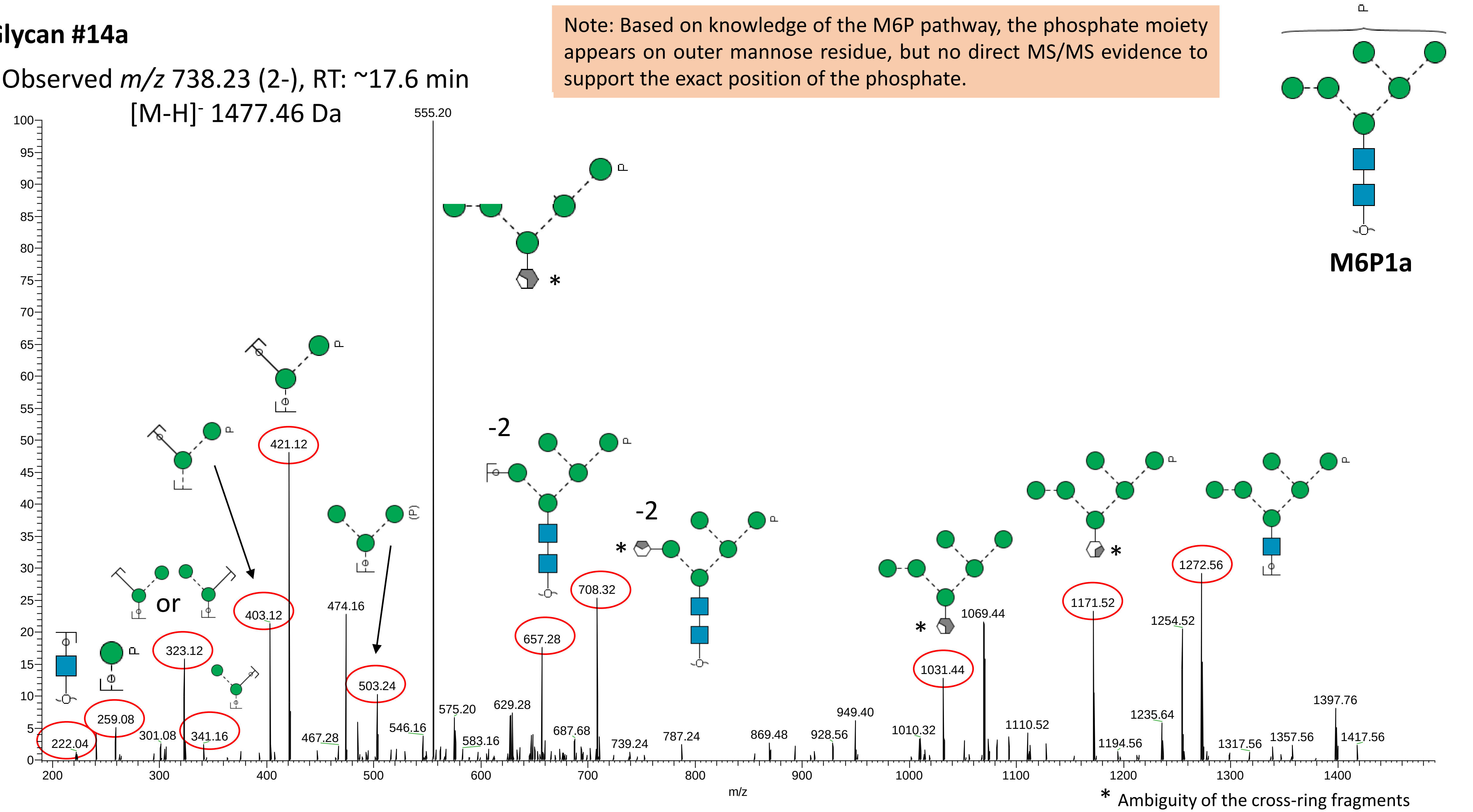

**Glycan #14b**

Observed *m/z* 738.23 (2-), RT: ~18.5 min
[M-H]⁻ 1477.46 Da

Note: Based on knowledge of the M6P pathway, the phosphate moiety appears on outer mannose residue, but no MS/MS evidence to support the exact position.

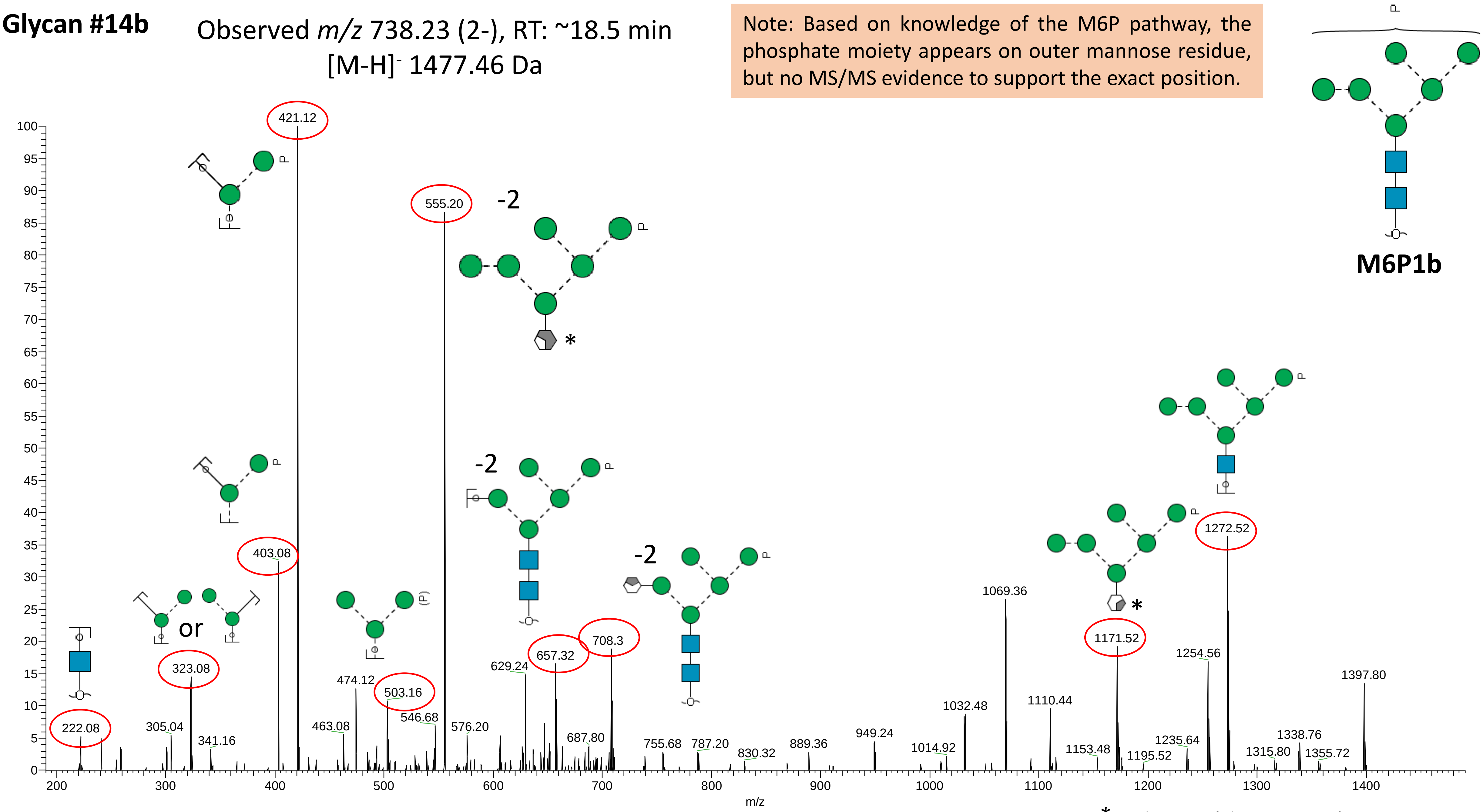


* Ambiguity of the cross-ring fragments

**Glycan #15**

Observed *m/z* 819.32 (2-), RT: ~17.8 min
$[M-H]^{-}$ 1639.51 Da

Note: Based on knowledge of the M6P pathway, the phosphate moiety appears on outer mannose residue, but no MS/MS evidence to support the exact position of the phosphate.

**M7P1**

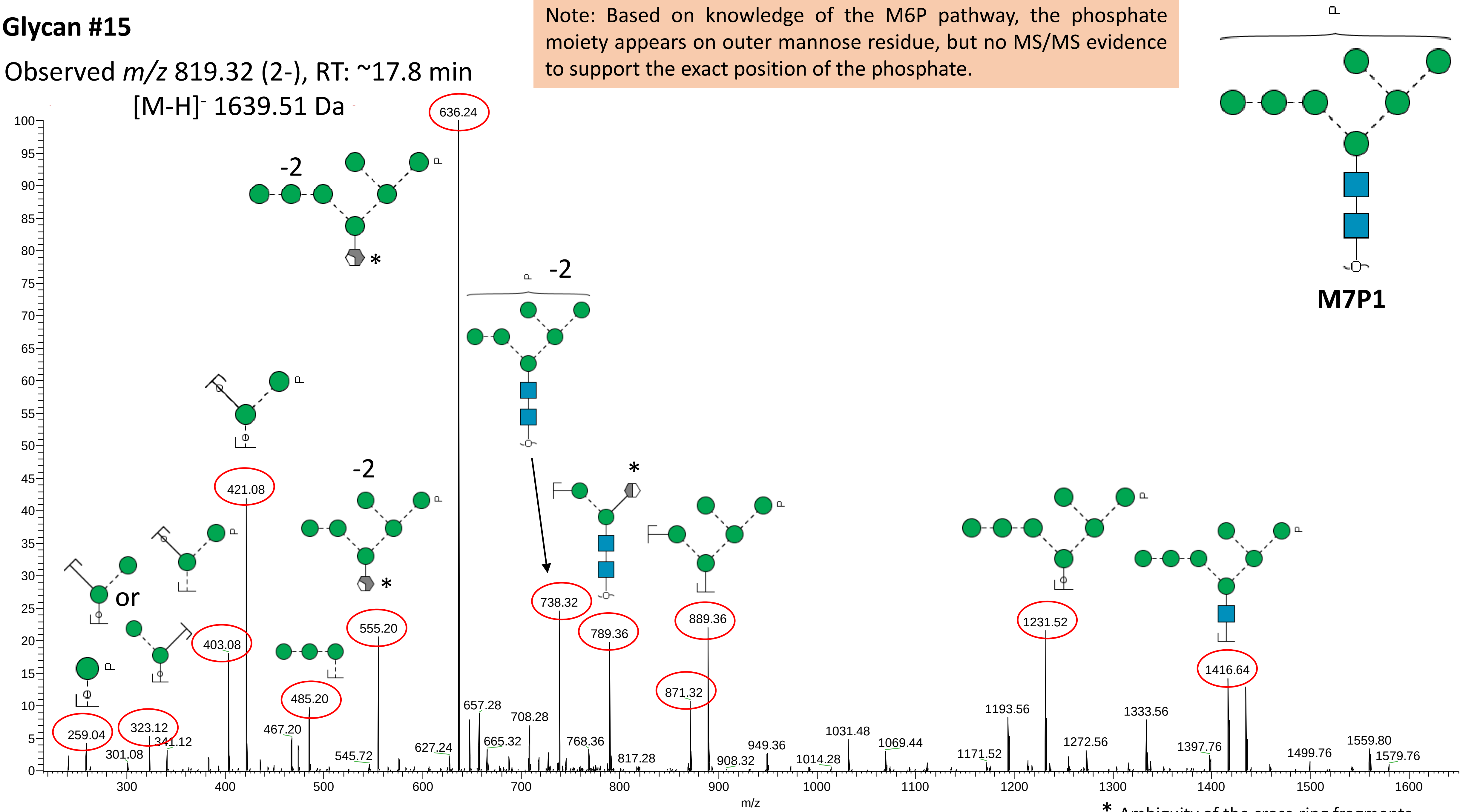


* Ambiguity of the cross-ring fragments

#### Glycan #16

Observed *m/z* 1260.60 (1-), RT: ~33.7 min
$[M-H]^-$ 1260.47 Da

Note: From biosynthetic pathway, the single β1,2-GlcNAc residue is predicted to occupy the α-1,3 arm but no MS/MS evidence to support this, so here left unassigned.

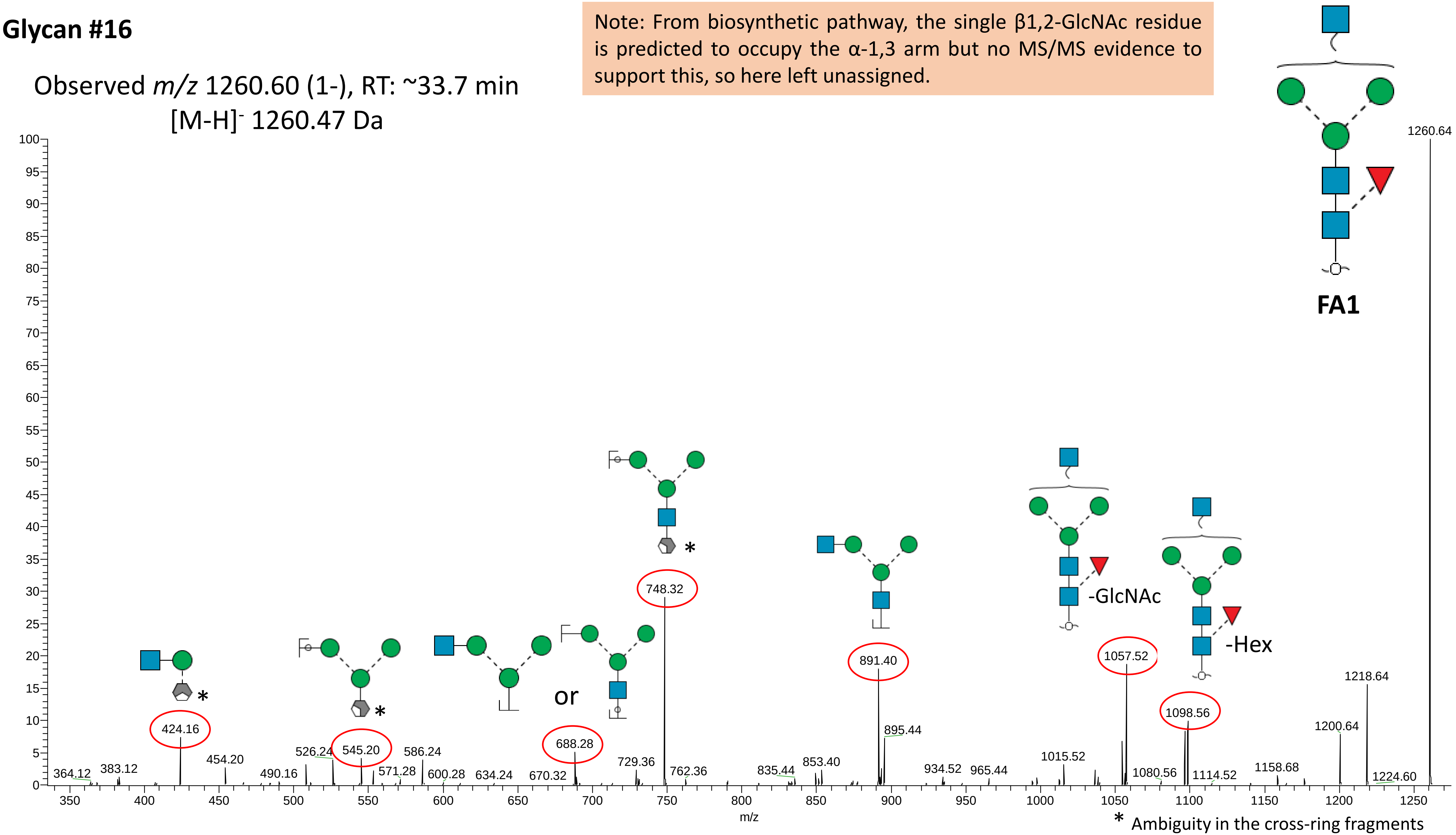

**Glycan #17**

Observed *m/z* 731.28 (2-), RT: ~26.1 min

[M-H]⁻ 1463.55 Da

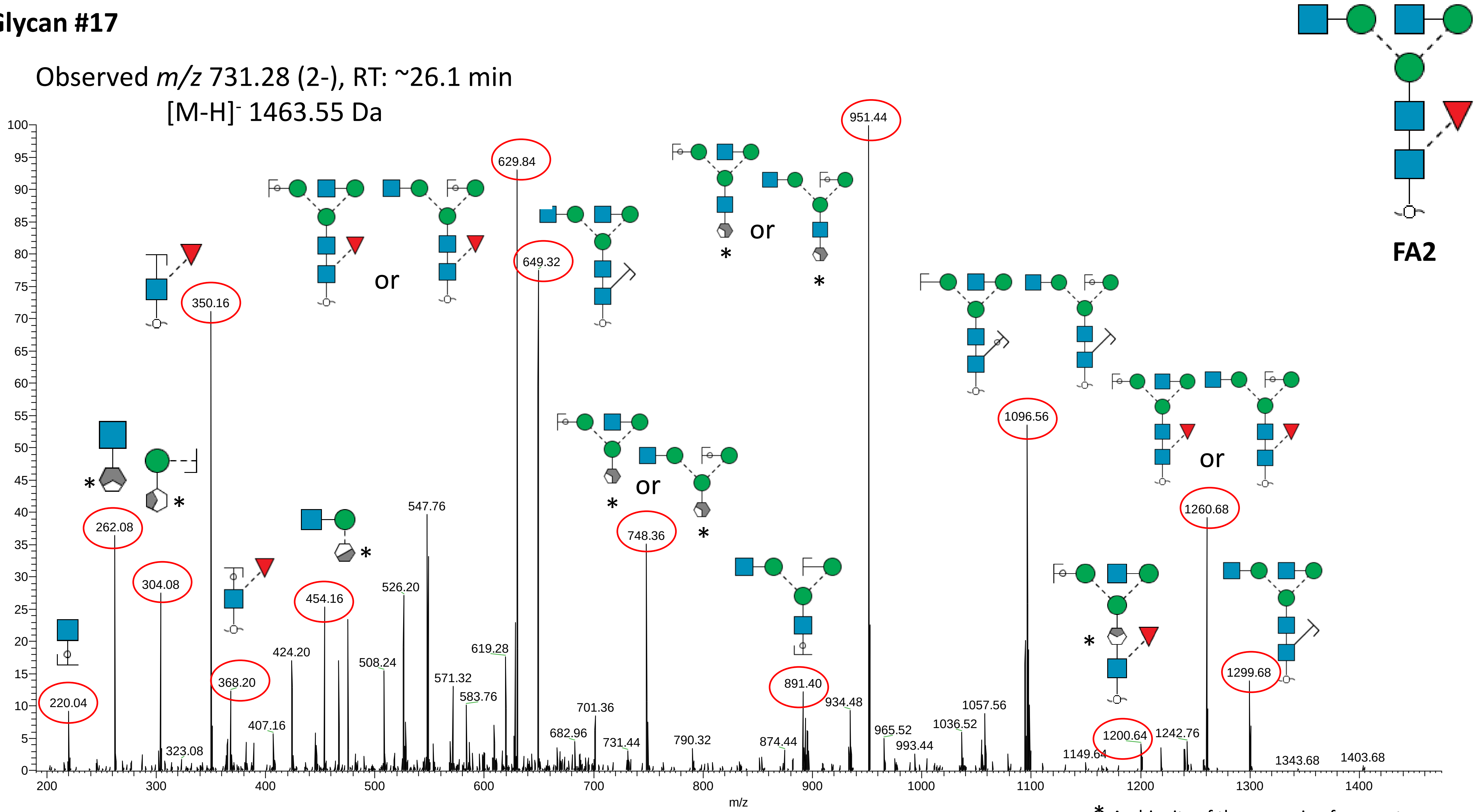


* Ambiguity of the cross-ring fragments

#### Glycan #18

Observed $m/z$ 783.28 (2-), RT: ~22.3 min
$[M-H]^-$ 1567.56 Da

Note: This glycan is annotated as the α2,6-sialyl linkage isomer based on the early PGC-LC elution and experience with neutrophil N-glycan profiling. The elongated antenna is known to frequently occupy the α1,3-arm rather than the α1,6-arm of neutrophil *N*-glycans and have thus been assigned as such.

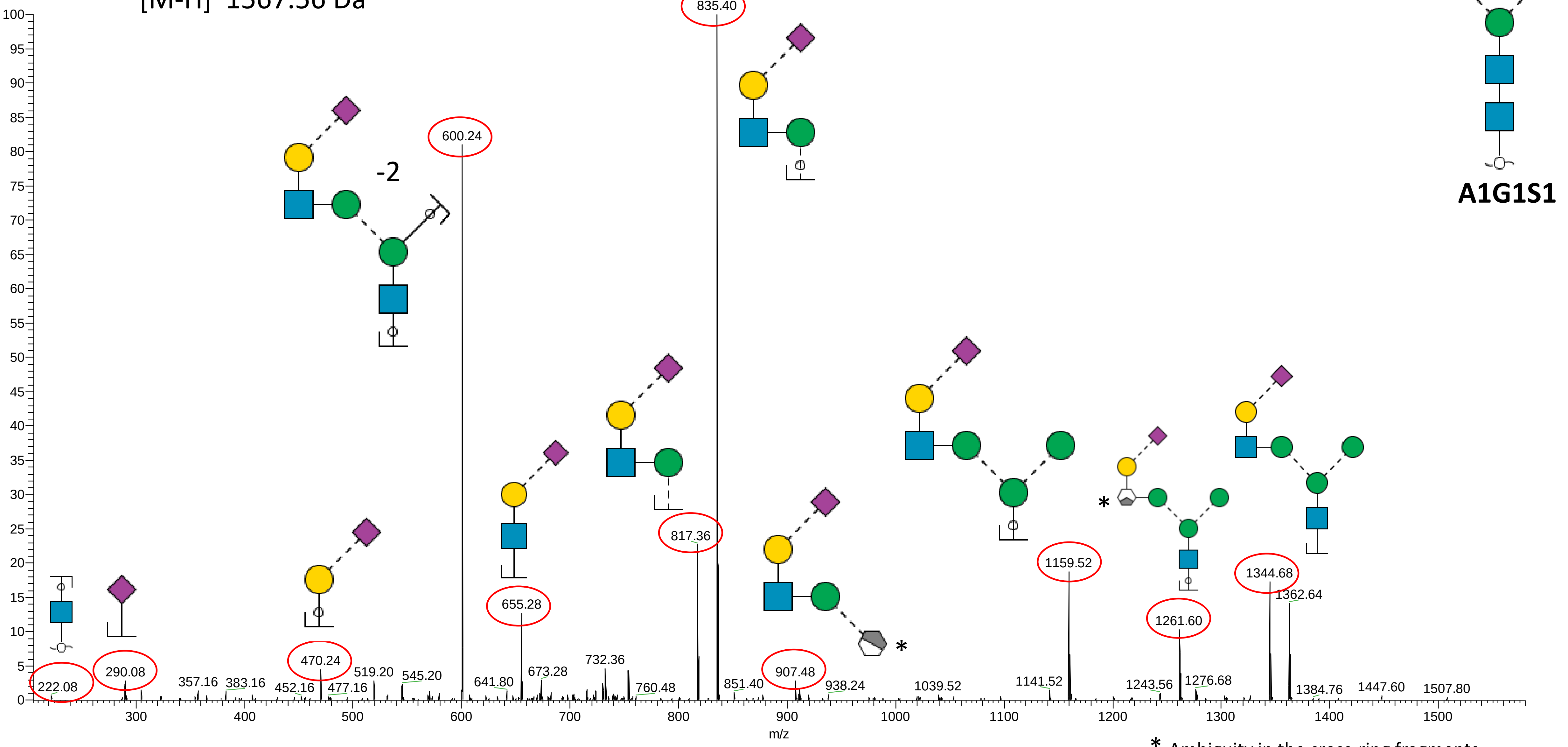


* Ambiguity in the cross-ring fragments

**Glycan #19a**

Note: This glycan is annotated as the α2,6-sialyl linkage isomer based on the early PGC-LC elution and experience with neutrophil *N*-glycan profiling. The elongated antenna is known to frequently occupy the α1,3-arm rather than the α1,6-arm of neutrophil *N*-glycans and have thus been assigned as such.

Observed *m/z* 856.31 (2-), RT: ~27.9 min
$[M-H]^-$ 1713.62 Da

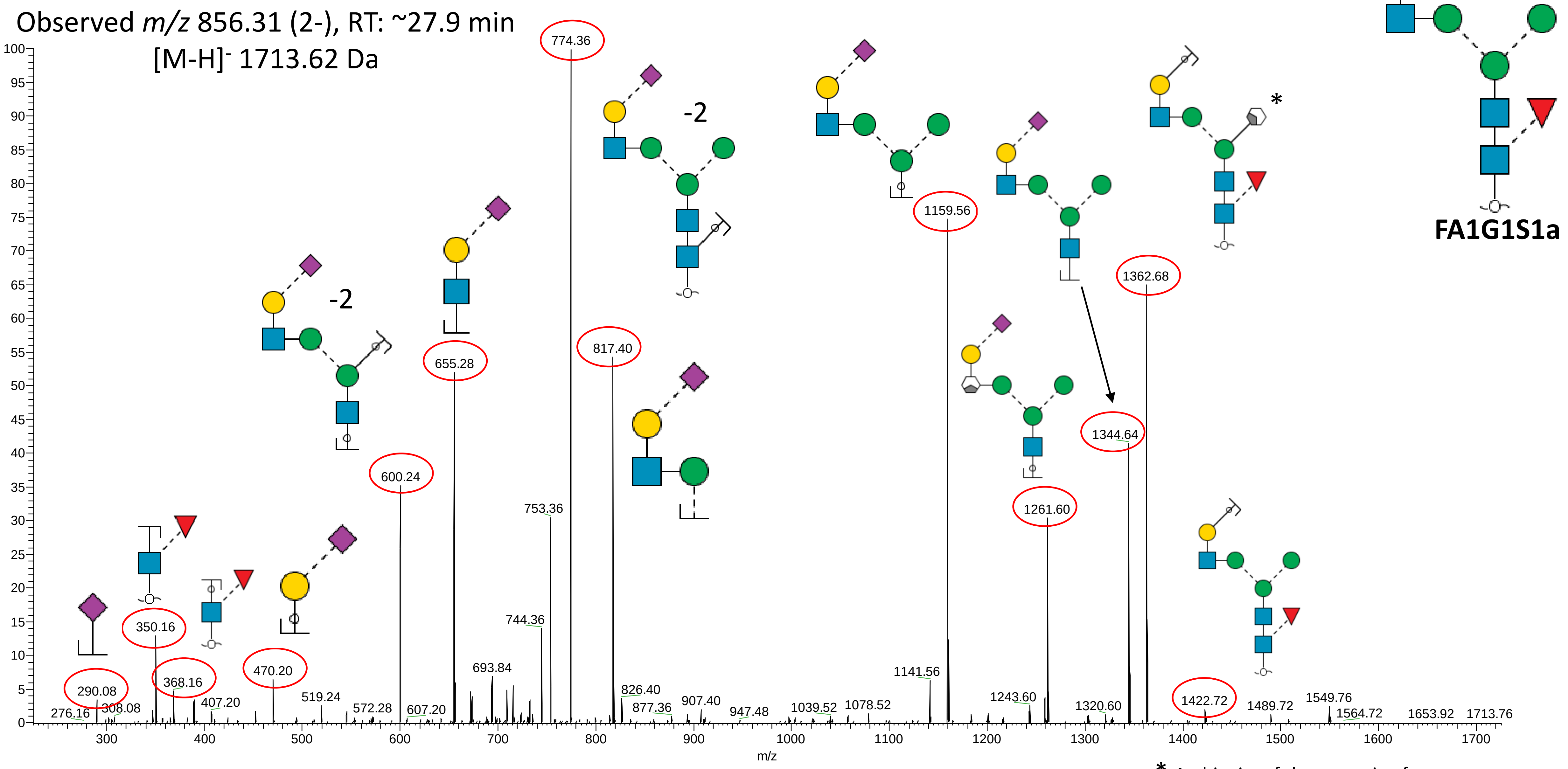


* Ambiguity of the cross-ring fragments

#### Glycan #19b

Note: This glycan is annotated as the α2,3-sialyl linkage isomer based on the late PGC-LC elution and experience with neutrophil *N*-glycan profiling. The elongated antenna is known to frequently occupy the α1,3-arm rather than the α1,6-arm of neutrophil *N*-glycans and is thus assigned as such.

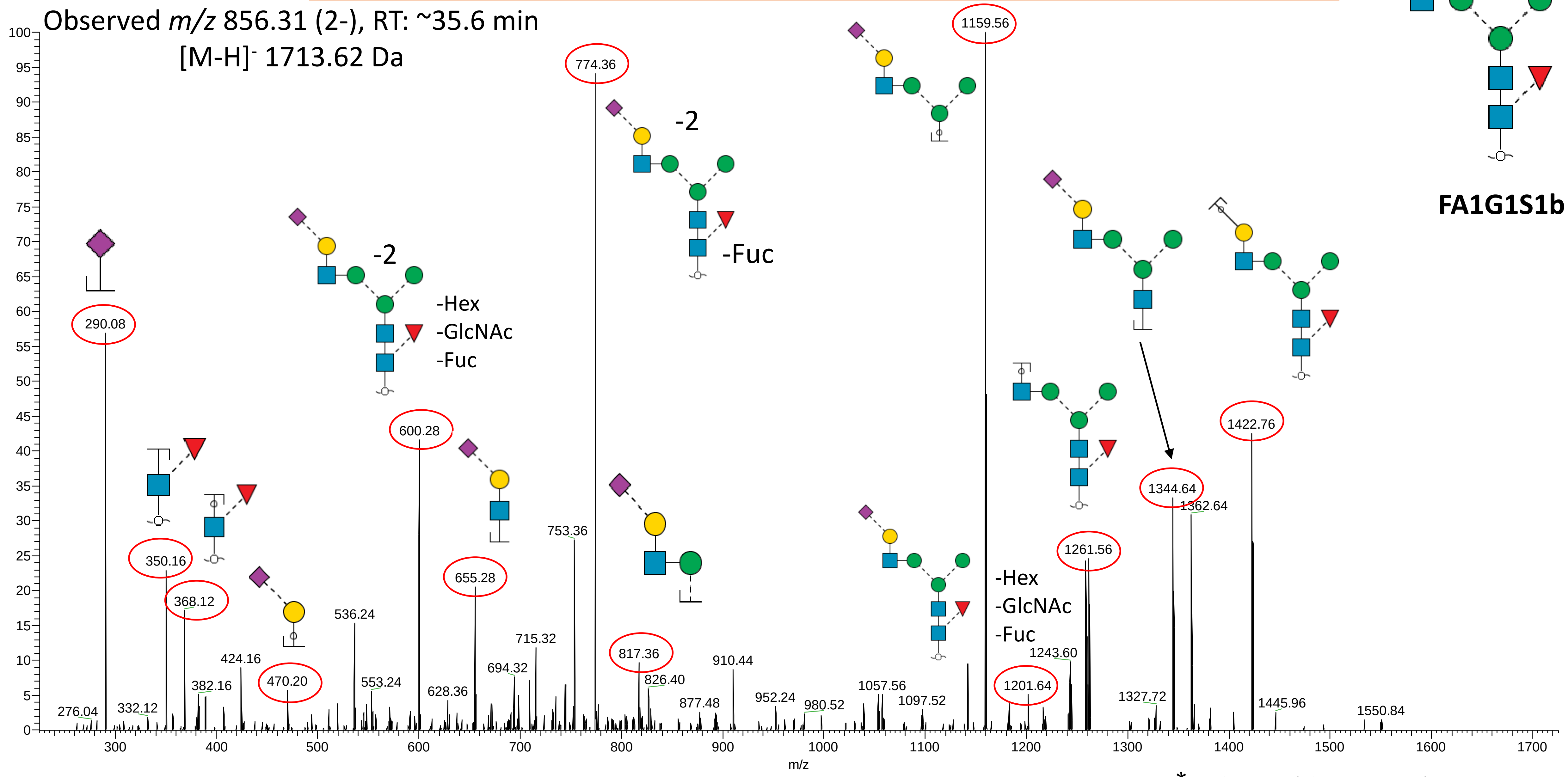


* Ambiguity of the cross-ring fragments

**Glycan #19c**

Note: This low abundance glycan is possibly the isomer where the Sia-Gal-GlcNAc antenna occupy the α1,6-arm rather than the α1,3-arm common for neutrophil *N*-glycans but no direct MS/MS evidence to support this

Observed *m/z* 856.31 (2-), RT: ~43.8 min
[M-H]- 1713.62 Da

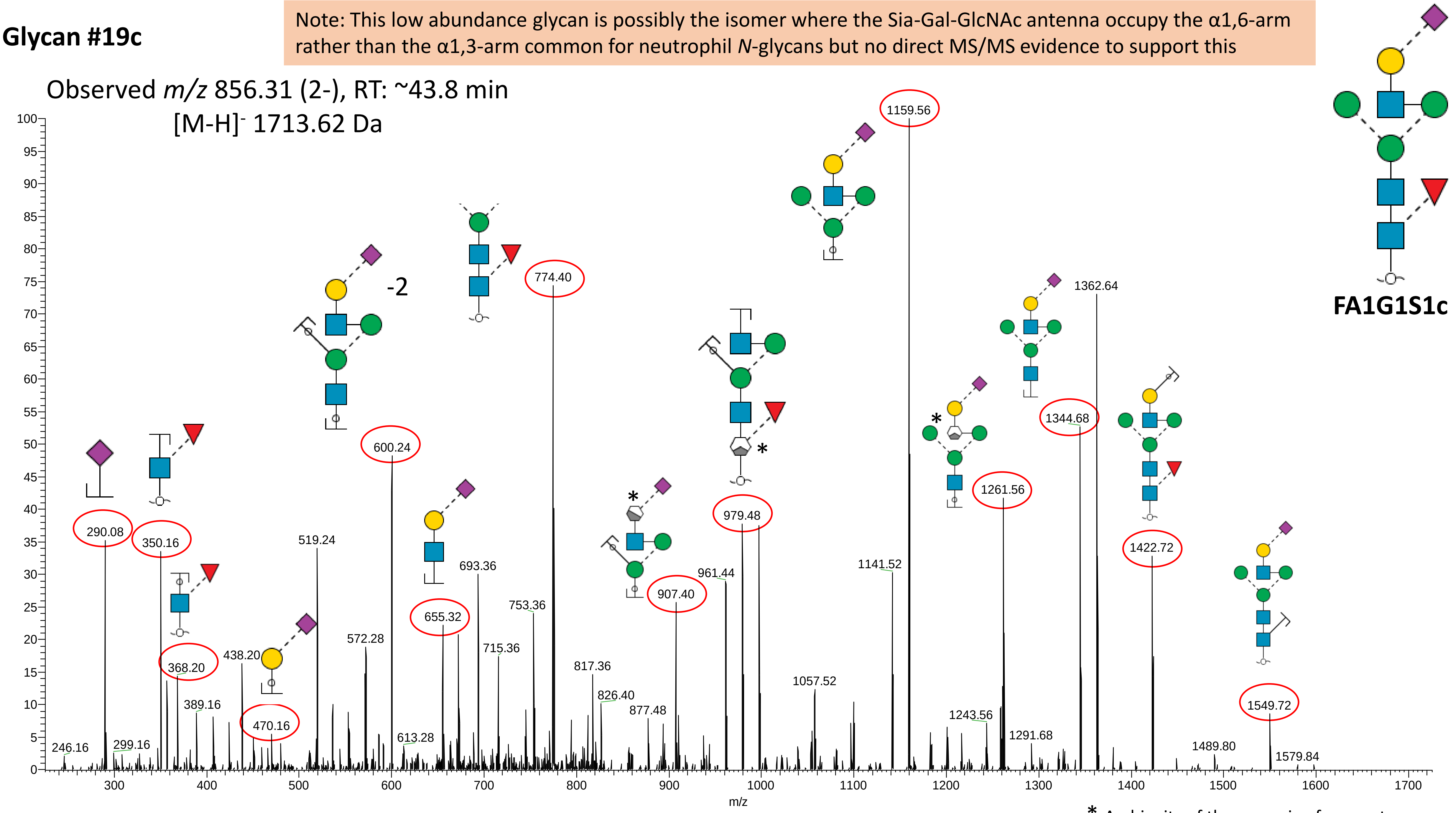


* Ambiguity of the cross-ring fragments

**Glycan #20a**

Note: This glycan is annotated as the α2,6-sialyl linkage isomer based on the early PGC-LC elution and higher abundance (relative to Glycan #20b) and experience with neutrophil *N*-glycan profiling. The elongated antenna is known to frequently occupy the α1,3-arm rather than the α1,6-arm of neutrophil *N*-glycans and is thus represented as such.

Observed *m/z* 957.85 (2-), RT: ~29.3 min

[M-H]$^{-}$ 1916.70 Da

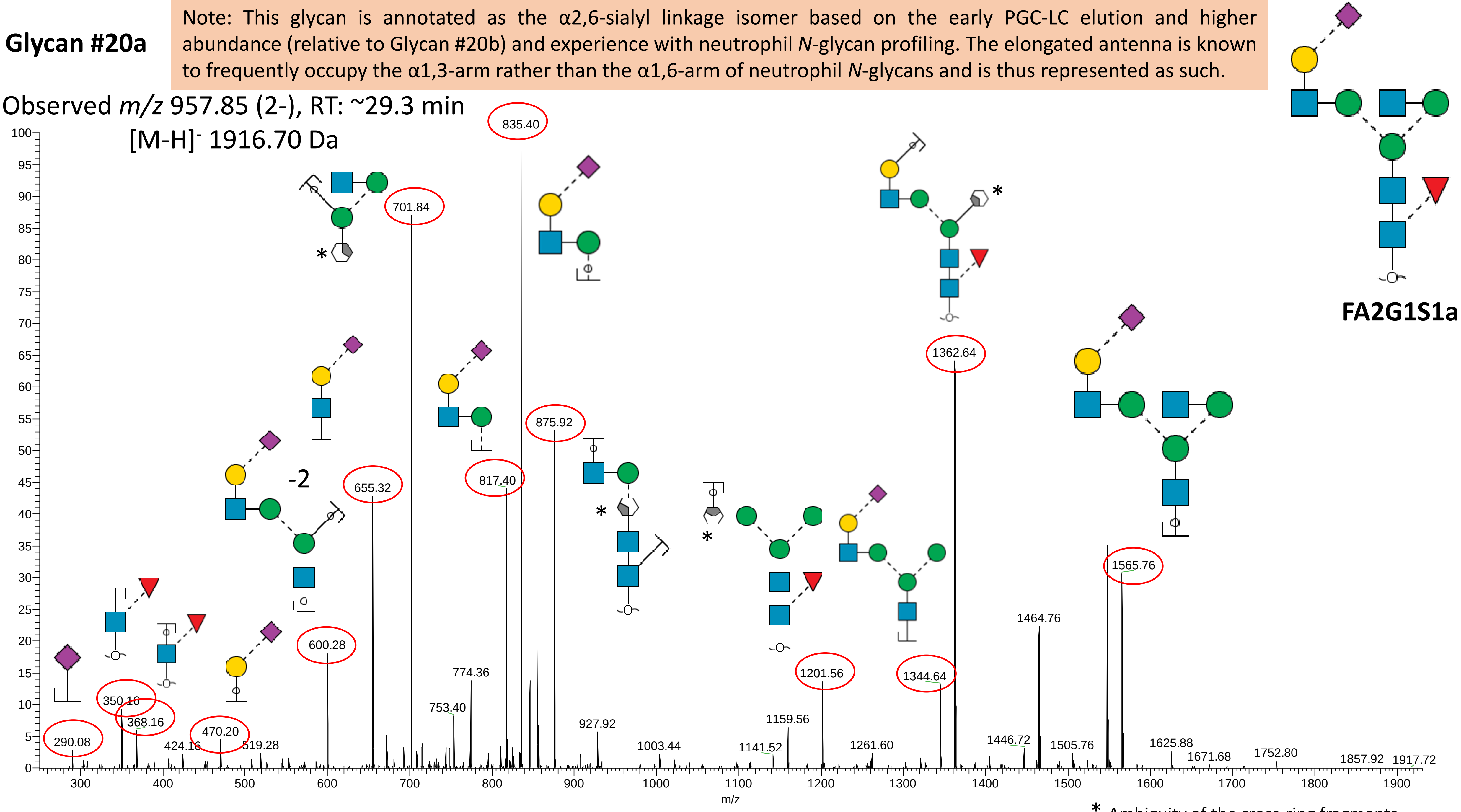


* Ambiguity of the cross-ring fragments

**Glycan #20b**

Note: This glycan is annotated as the α2,3-sialyl linkage isomer based on the later PGC-LC elution and lower abundance (relative to Glycan #20a) and experience with neutrophil *N*-glycan profiling. The elongated antenna is known to frequently occupy the α1,3-arm rather than the α1,6-arm of neutrophil *N*-glycans and have thus been assigned as such.

Observed *m/z* 957.85 (2-), RT: ~37.3 min

$[M-H]^-$ 1916.70 Da

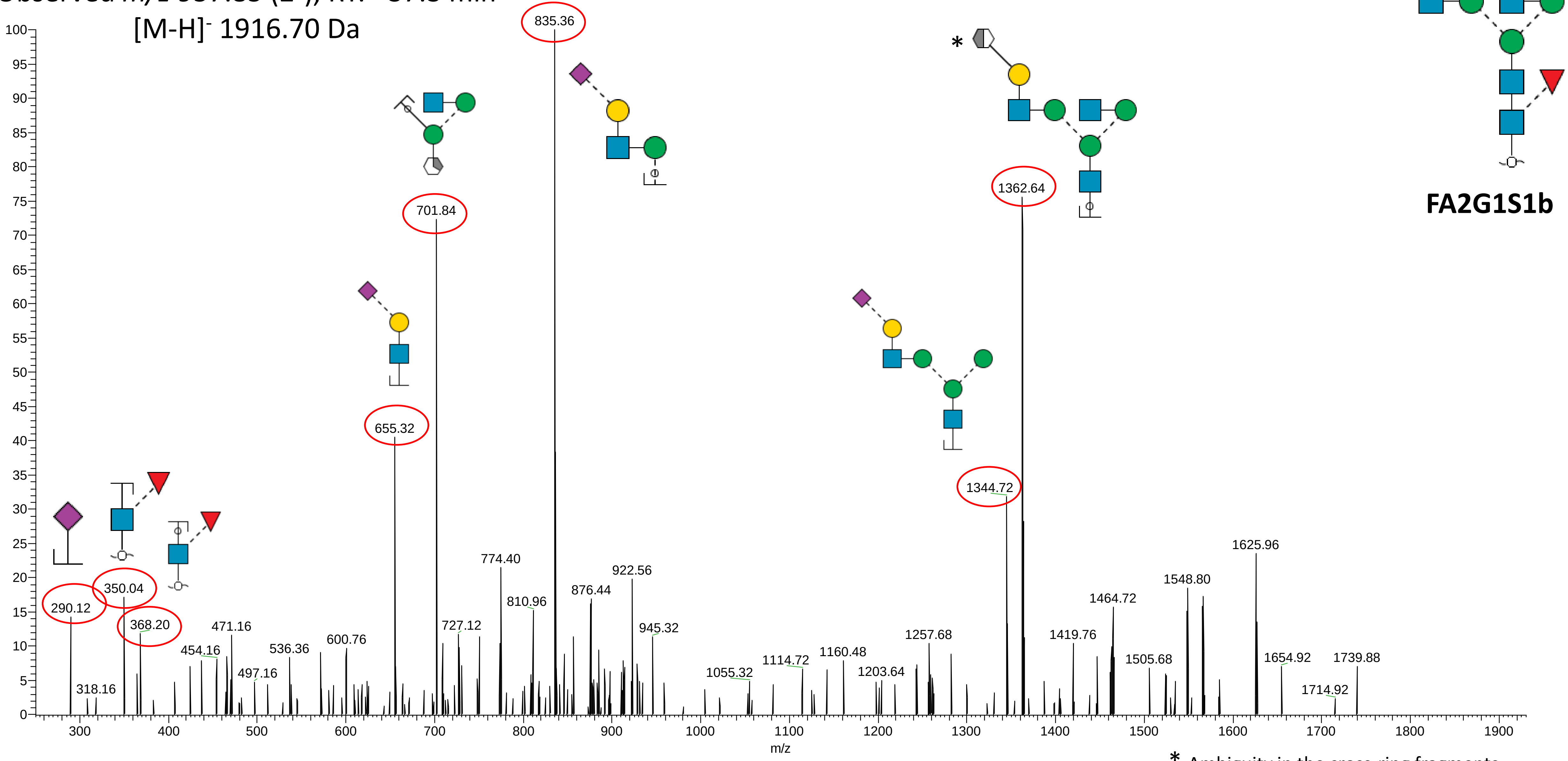


* Ambiguity in the cross-ring fragments

**Glycan #21**

Note: This glycan is annotated as the α2,6-sialyl linkage isomer based on the early PGC-LC elution and experience with neutrophil *N*-glycan profiling. The arm position of the sialic acid could not be deduced.

Observed *m/z* 1038.88 (2-), RT: ~31.3 min

[M-H]$^{-}$ 2078.75 Da

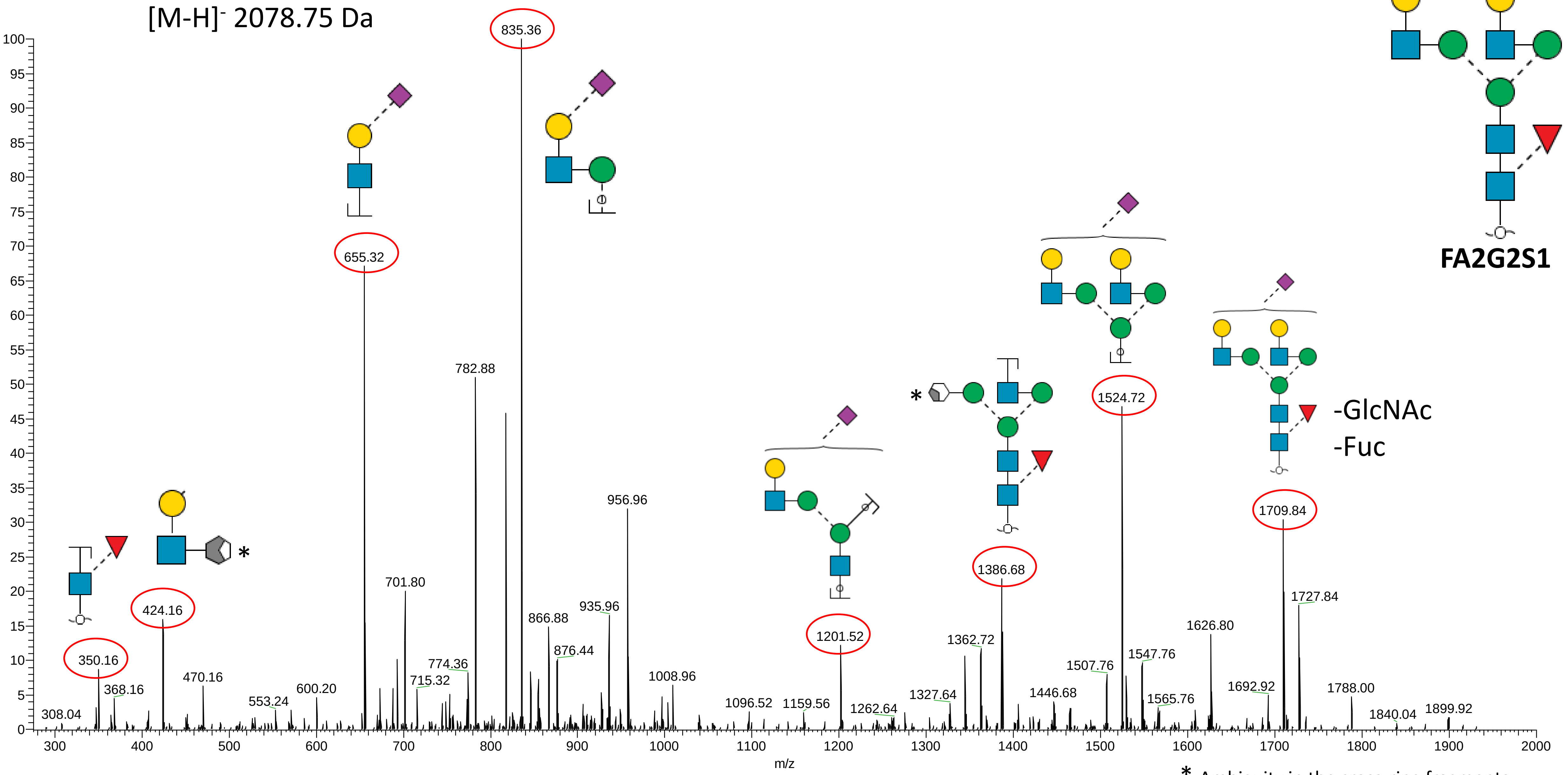


* Ambiguity in the cross-ring fragments

**Glycan #22a**

Observed *m/z* 1184.52 (2-), RT: ~31.1 min

[M-H]⁻ 2369.84 Da

Note: This glycan has been annotated as the α2,6-/α2,6-sialyl isomer based on the early PGC-LC elution time.

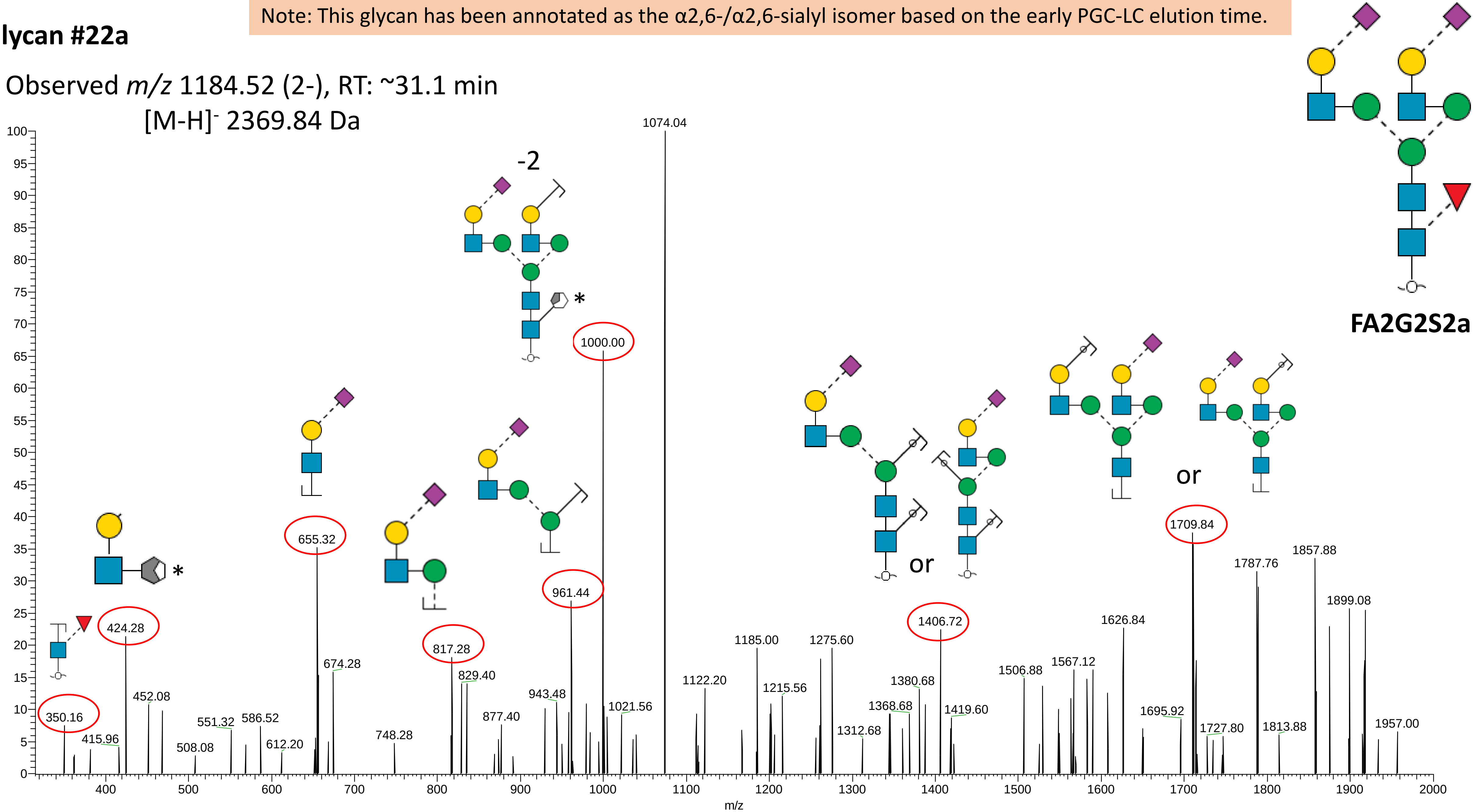


* Ambiguity of the cross-ring fragments

#### Glycan #22b

Observed *m/z* 1184.52 (2-), RT: ~39.4 min
[M-H]- 2369.84 Da

Note: This glycan has been annotated as the α2,6-/α2,3-sialyl isomer based on the late PGC-LC elution time (relative to Glycan #22a) and experience with neutrophil *N*-glycosylation. Not possible to position the α2,6- and α2,3-sialic acid to specific arms.

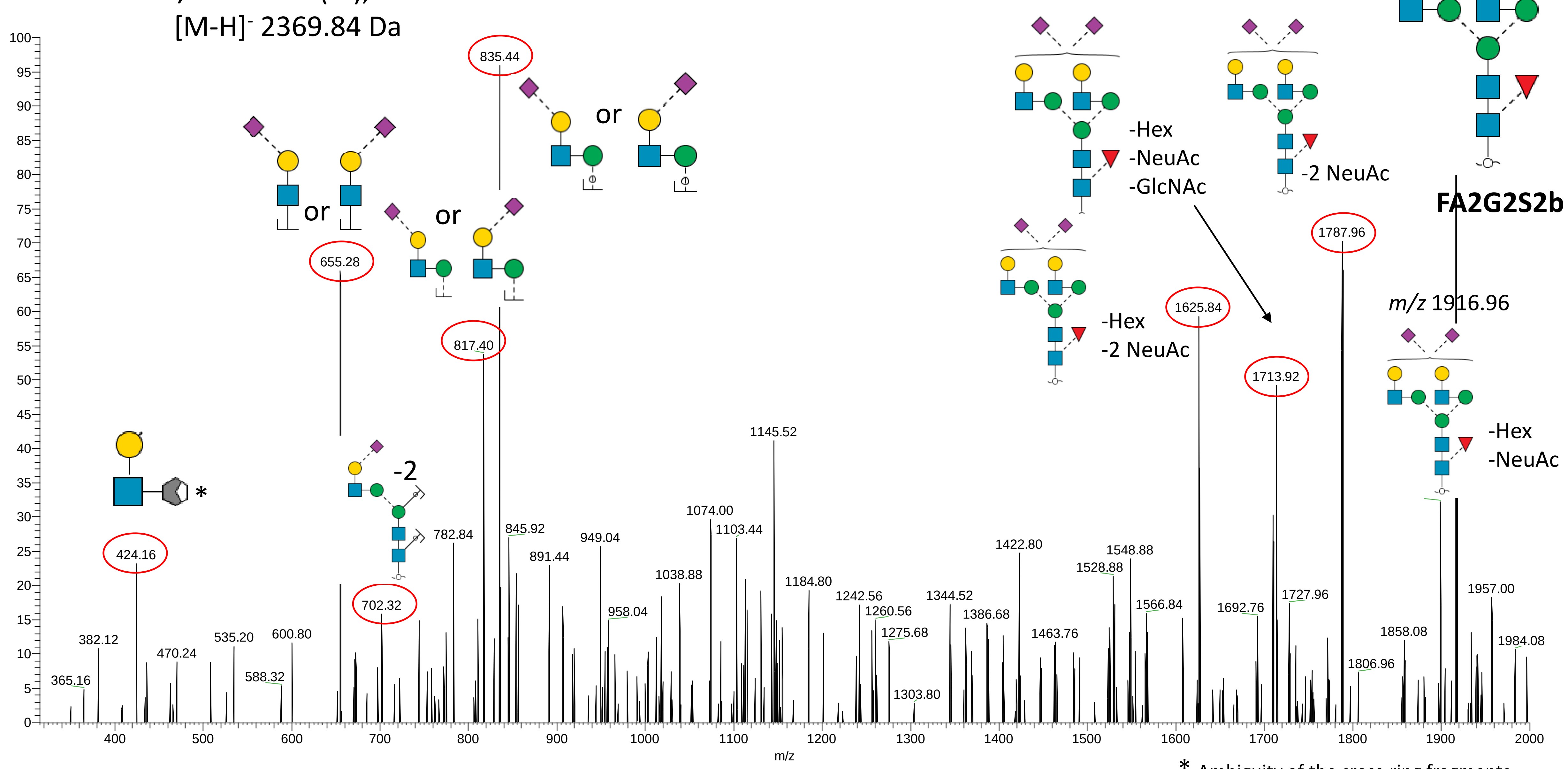


* Ambiguity of the cross-ring fragments

**Glycan #23**

Observed *m/z* 1111.91 (2-), RT: ~30.1 min

[M-H]- 2224.82 Da

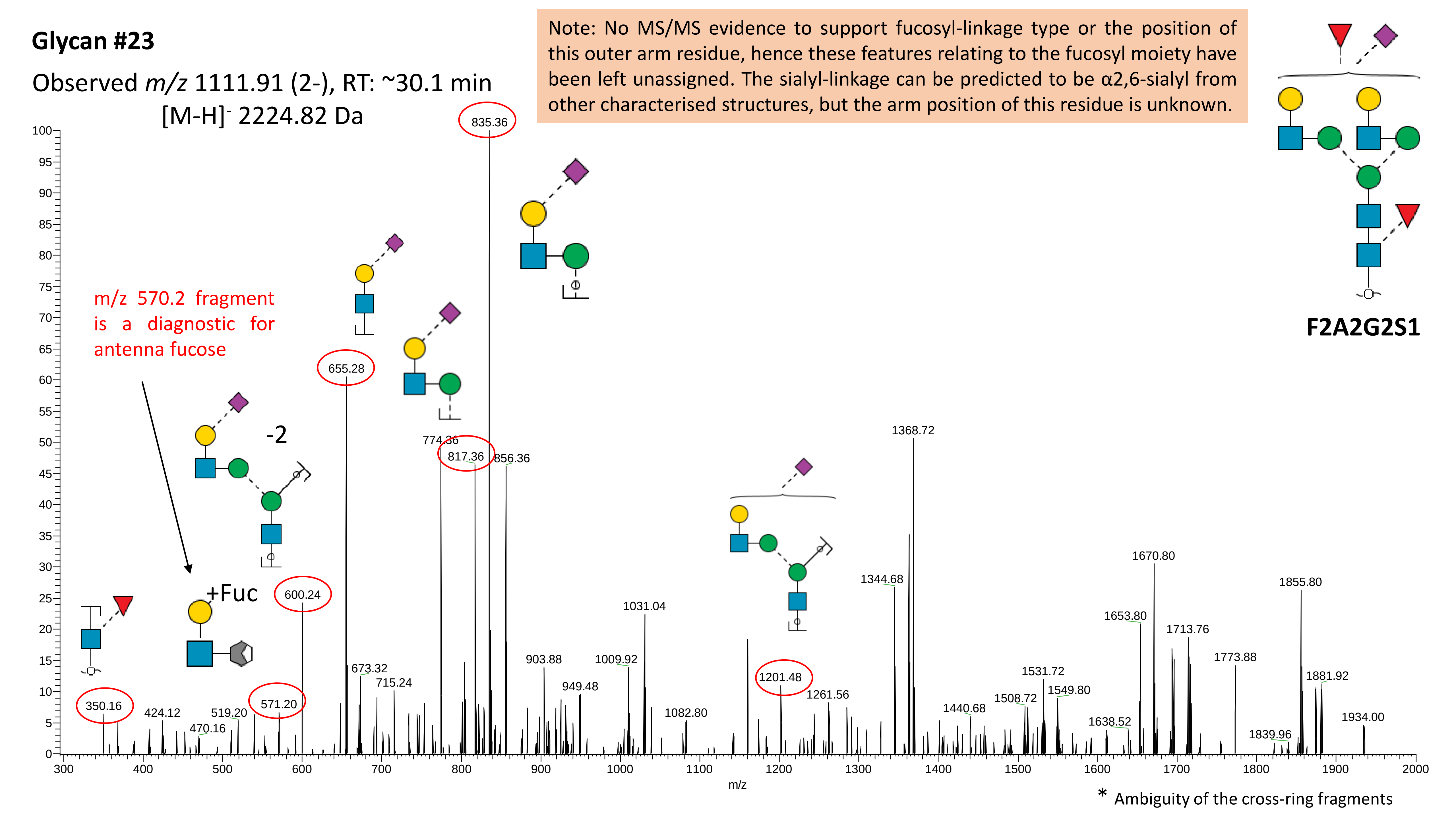

**Supplementary Data S2A**

Examples of Byonic-annotated reversed phase-LC-ESI-HCD-MS/MS (+) spectra of peptides identified with non-glyco PTMs (i.e. Met and Trp mono- and di-oxidation and Tyr mono-chlorination) from the analysis of unenriched peptide mixtures of nMPO

Human myeloperoxidase (P05164)

**Mono-oxidised methionine-containing peptide**

629KLMEQYGTPNNIDIWMGGVSEPLKR641 (identification confidence level, PEP 2D: 2.4x10$^{-20}$)

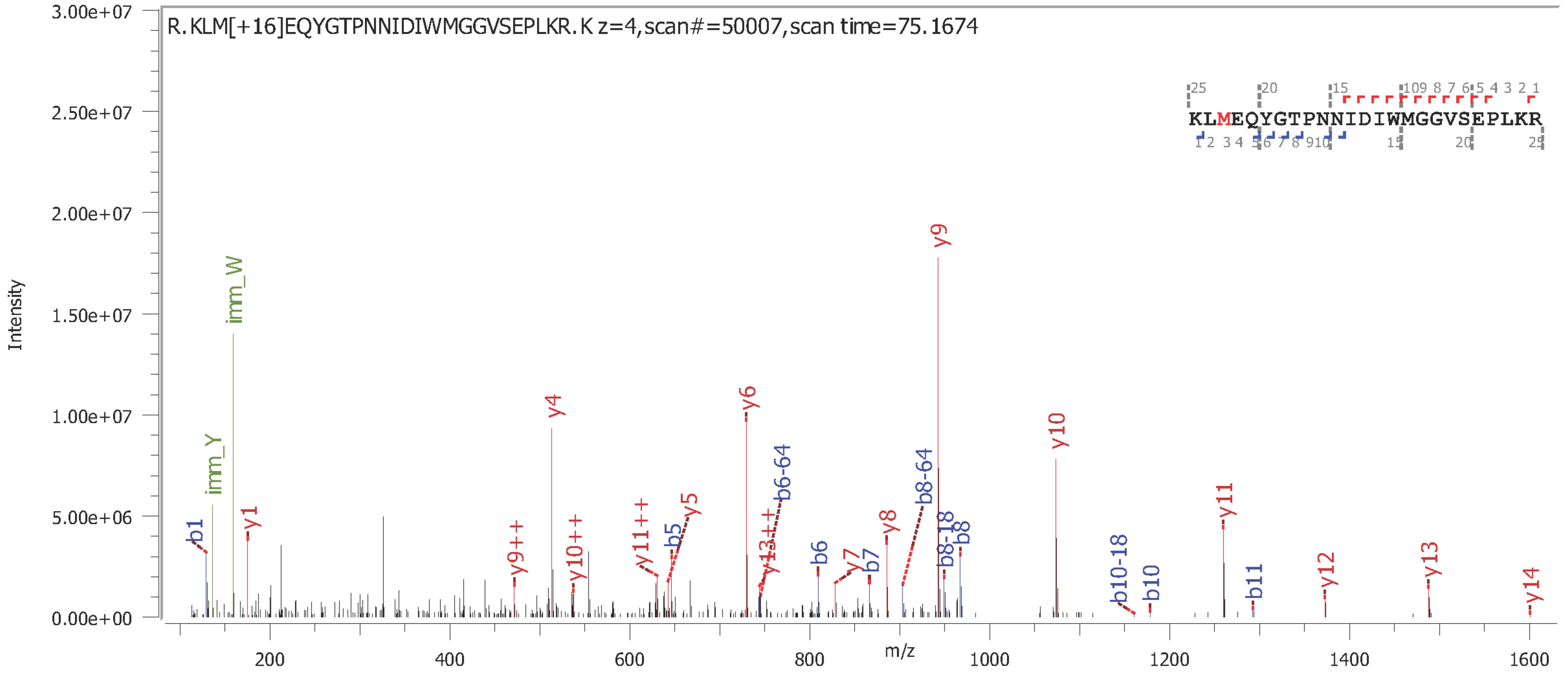

Human myeloperoxidase (P05164)

**Mono-oxidised tryptophan-containing peptide**

[198]WLPAEYEDGFSLPYG**W**TPGVKR[219] (identification confidence level, PEP 2D: $1.9 \times 10^{-15}$)

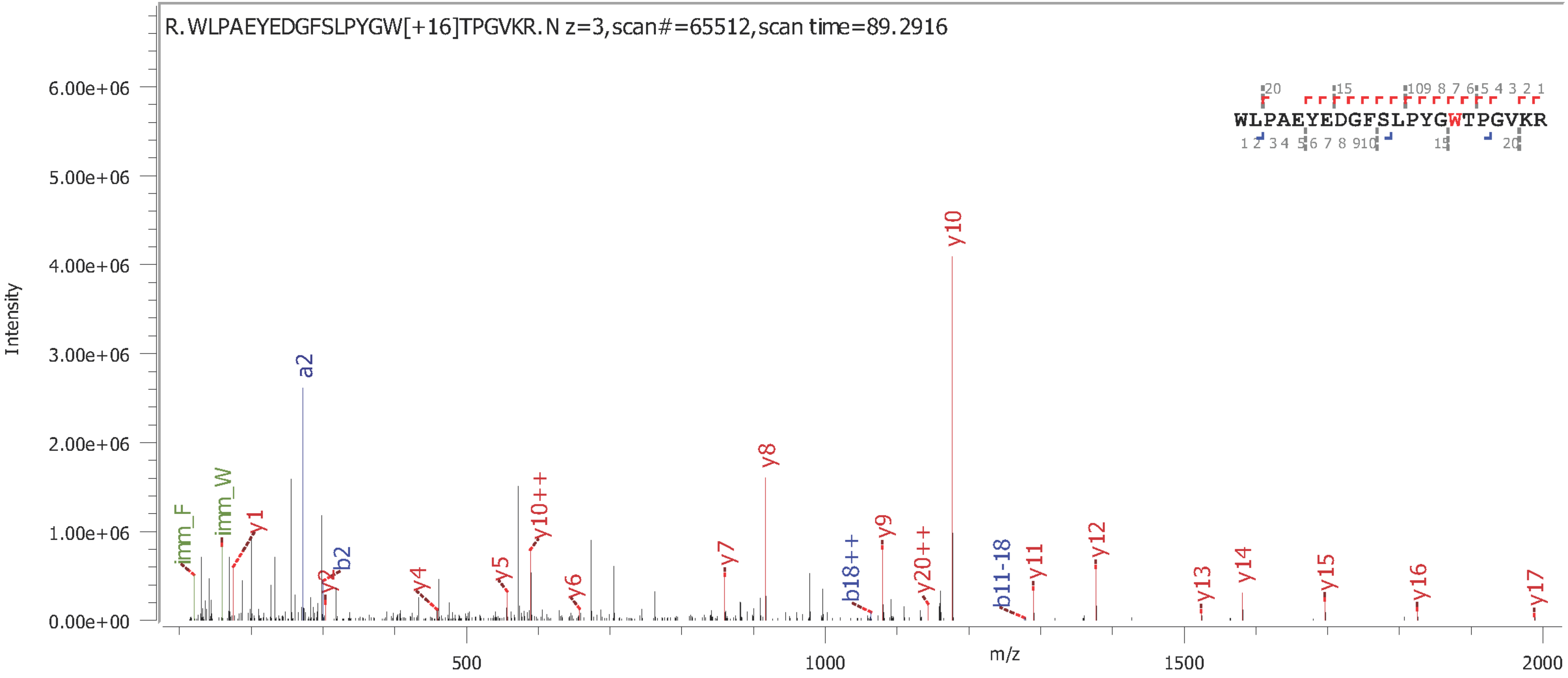

Human myeloperoxidase (P05164)

**Di-oxidised tryptophan-containing peptide**

[629]KLMEQYGTPNNIDIWMGGVSEPLKR[641] (identification confidence level, PEP 2D: $1.9 \times 10^{-18}$)

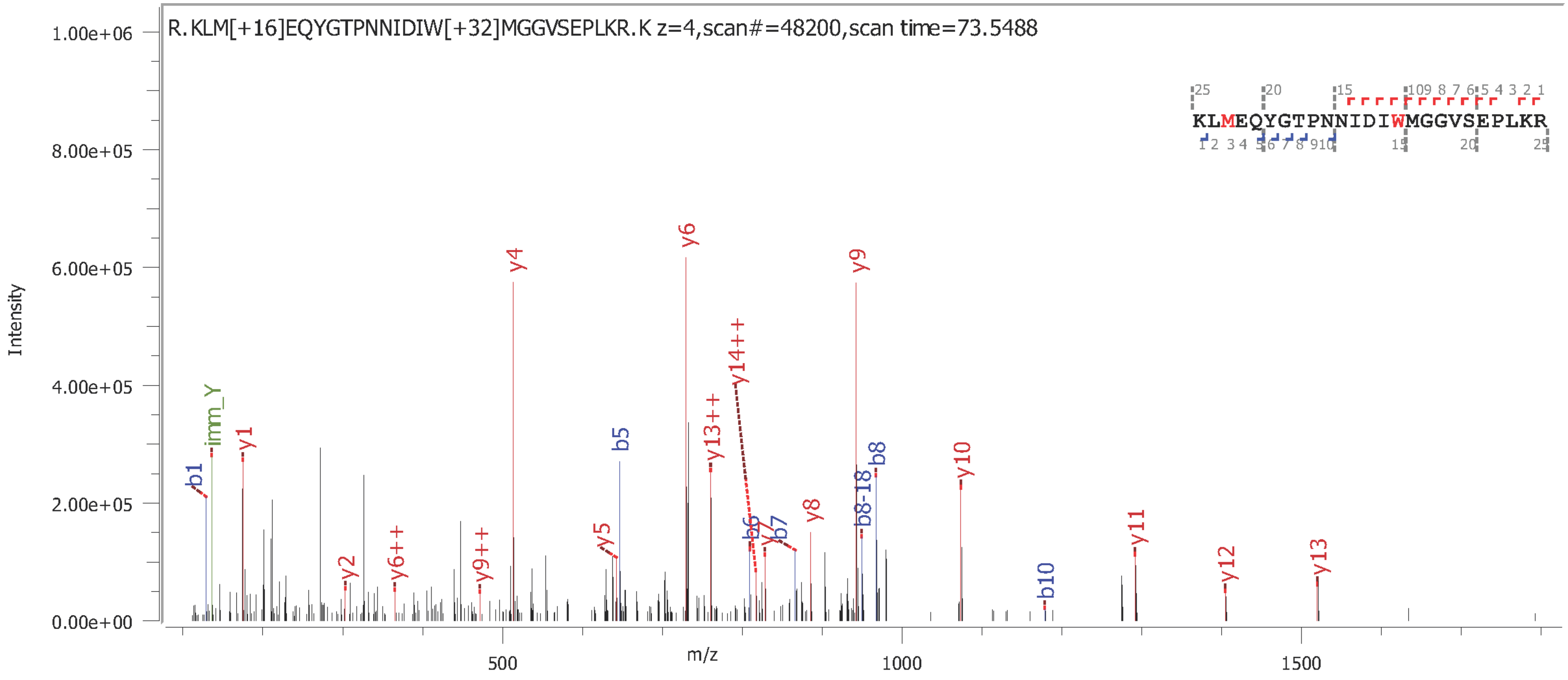

Human myeloperoxidase (P05164)

**Mono-chlorinated tyrosine-containing peptide**

[715]NNIFMSNS**Y**PRDFVNCSTLPALNLASWR[742] (identification confidence level, PEP 2D: $3.3x10^{-8}$)

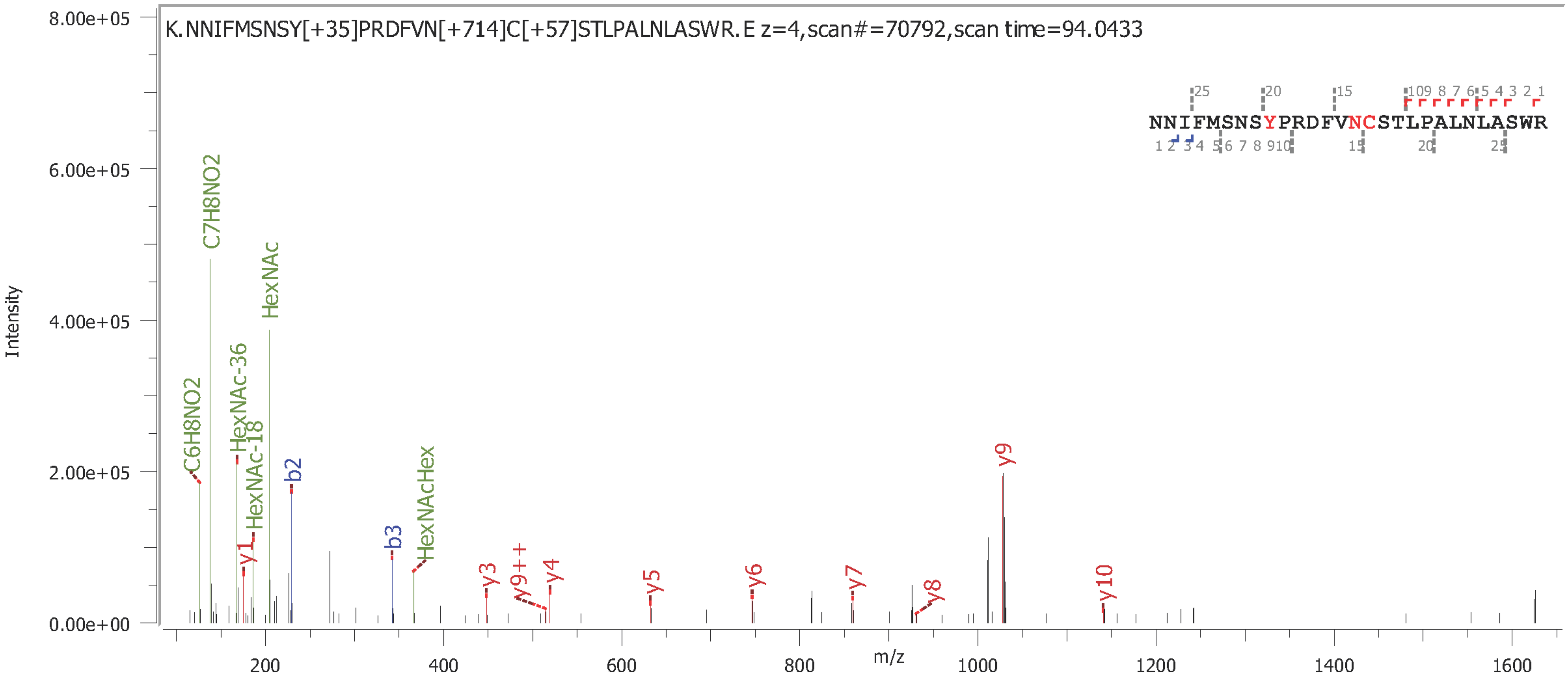

**Supplementary Data S2B**

Byonic-annotated reversed phase-LC-ESI-HCD-MS/MS (+) spectra of N- and C-terminal truncation variants of the MPO α- and β-chain identified from the analysis of unenriched peptide mixtures of nMPO

Human myeloperoxidase (P05164)

**α-chain N-terminal** (truncation variant)

164**G**VTCPEQDKYR (identification confidence level, PEP 2D: $4.0 \times 10^{-11}$)

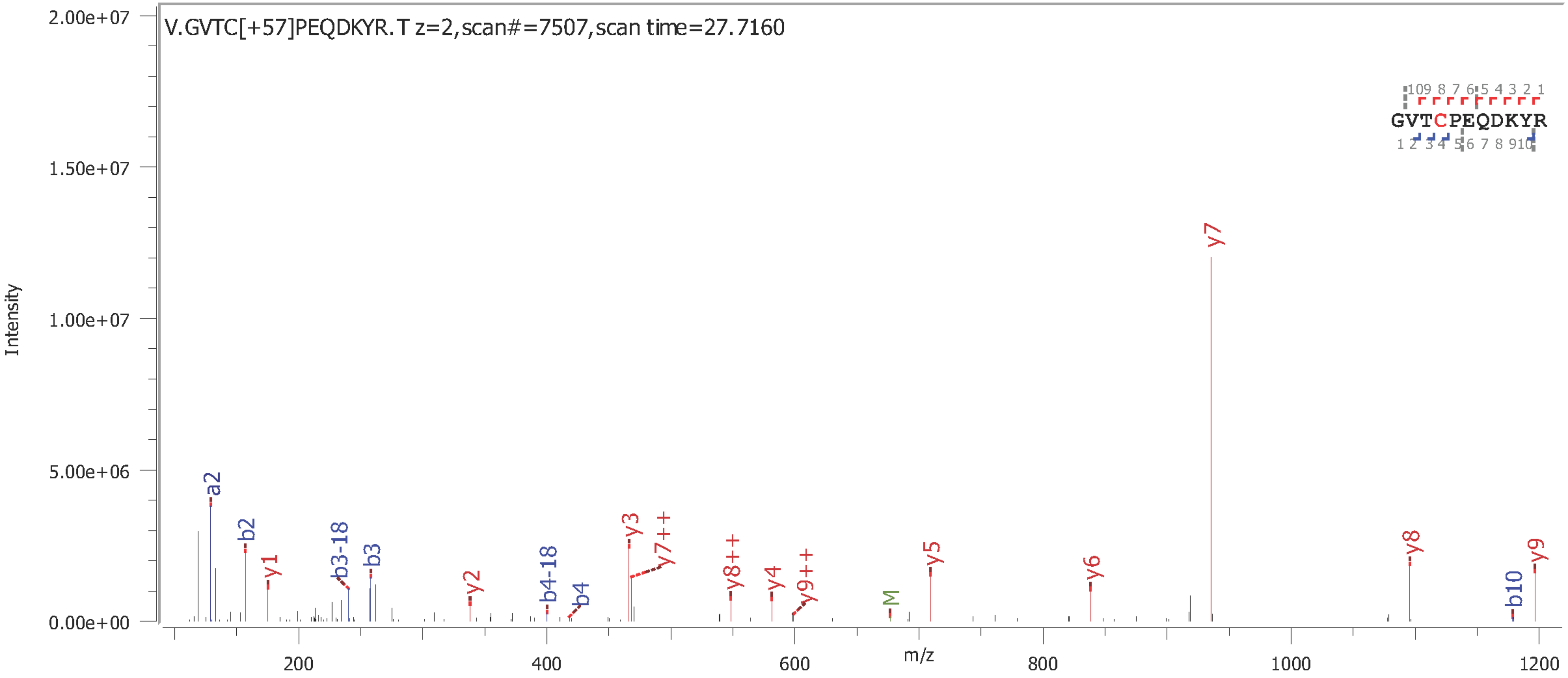

Human myeloperoxidase (P05164)

**α-chain N-terminal** (truncation variant)

165VTCPEQDKYR (identification confidence level, PEP 2D: $5.7 \times 10^{-7}$)

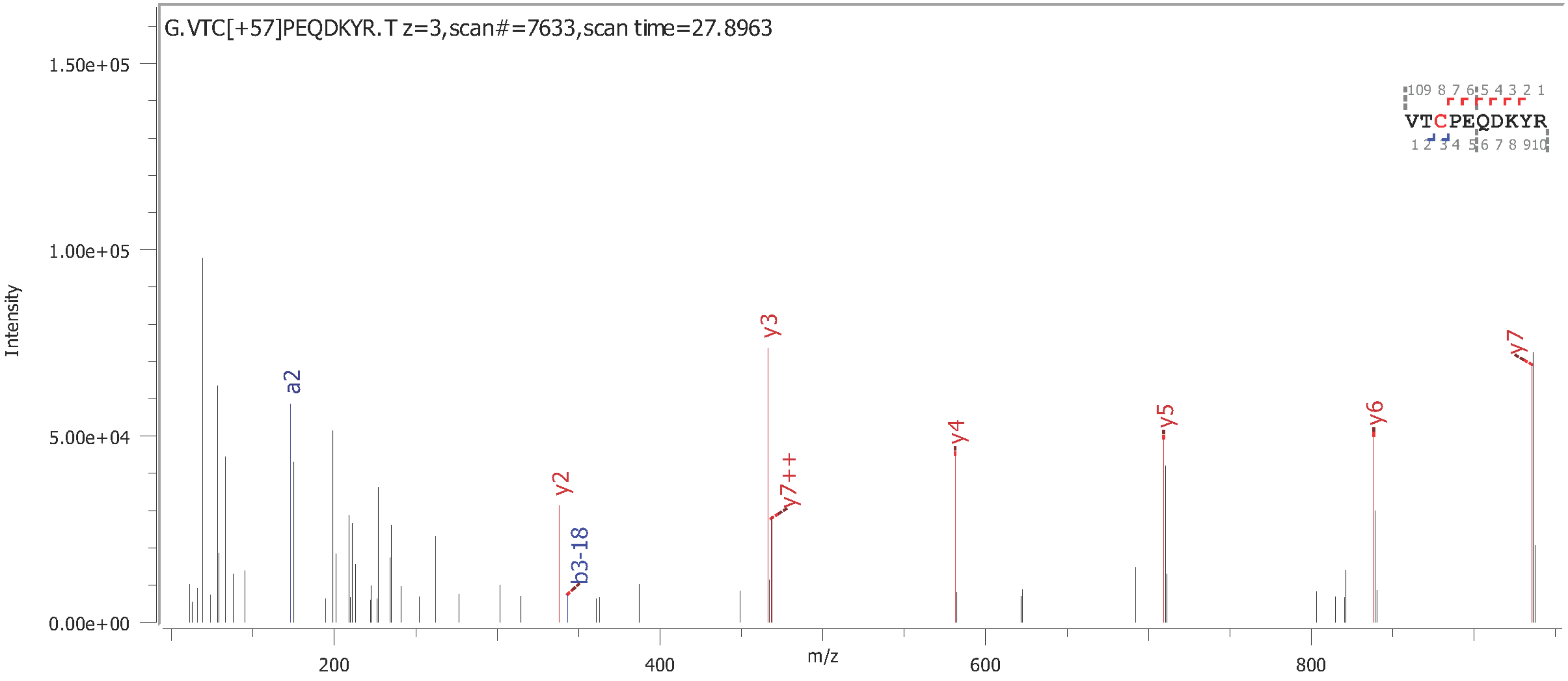

Human myeloperoxidase (P05164)

**α-chain N-terminal** (truncation variant)

$^{166}$TCPEQDKYR (identification confidence level, PEP 2D: $3.3x10^{-8}$)

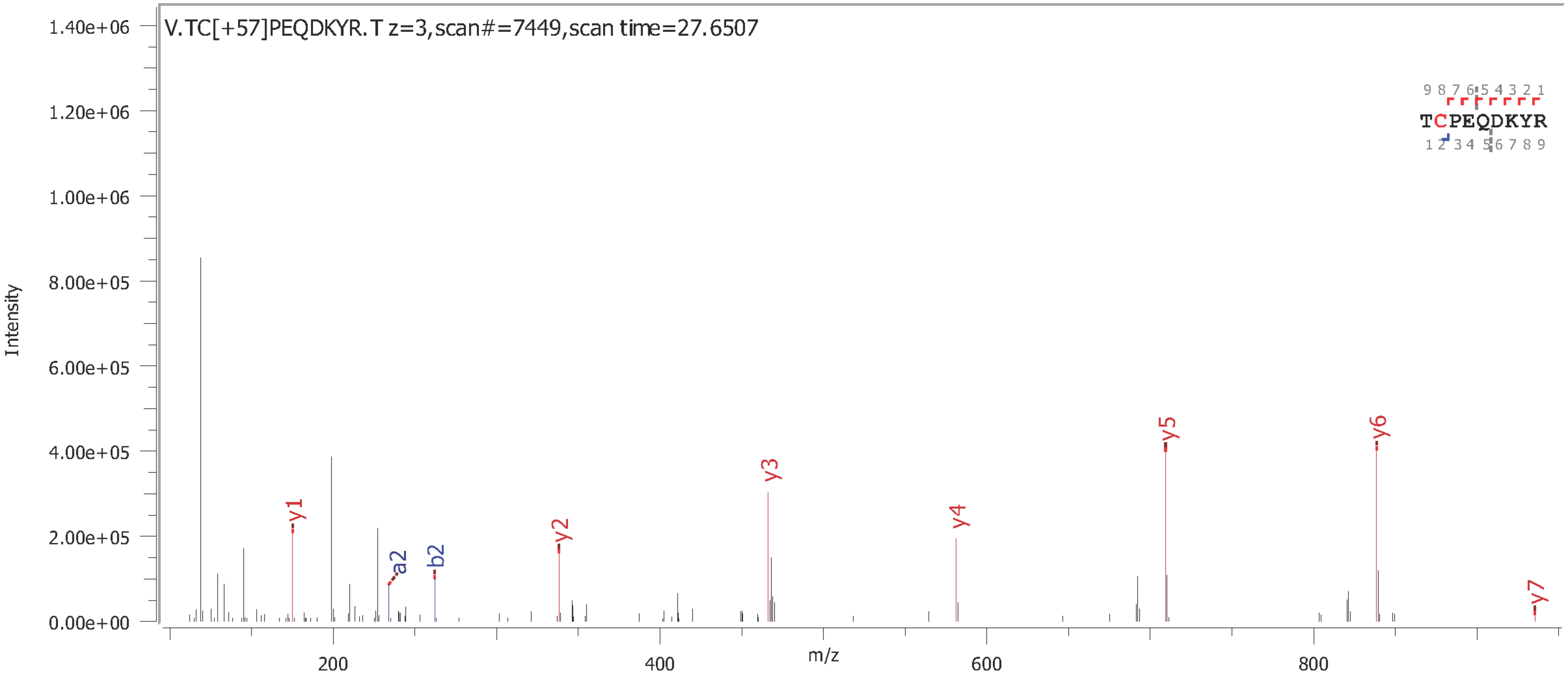

Human myeloperoxidase (P05164)

**α-chain N-terminal** (truncation variant)

SLMFMQWGQLLDHDLDFTPEPAA**R**[272] (identification confidence level, PEP 2D: $1.9 \times 10^{-23}$)

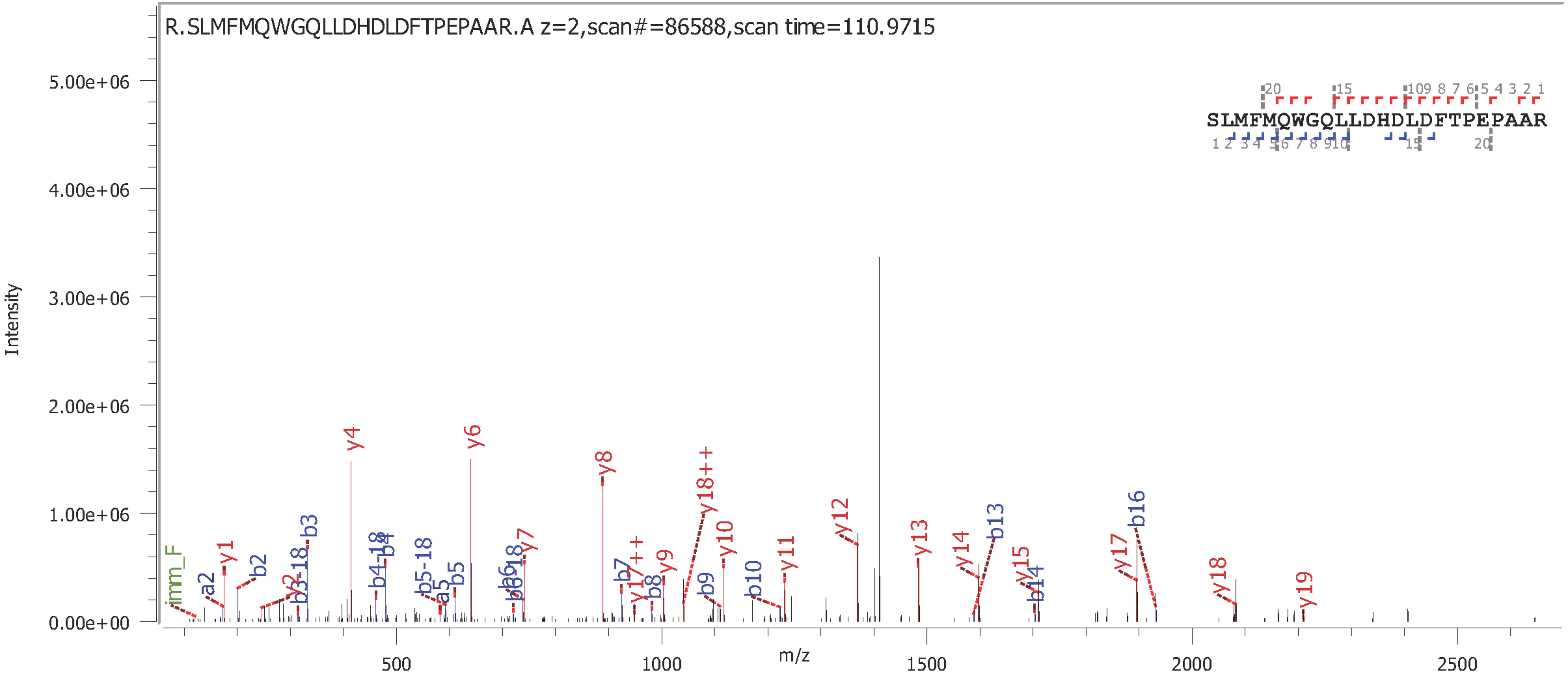

Human myeloperoxidase (P05164)

**β-chain N-terminal** (truncation variant)

277TGVNCETSCVQQPPCFPLKIPPNDPR (identification confidence level, PEP 2D: $8.7x10^{-14}$)

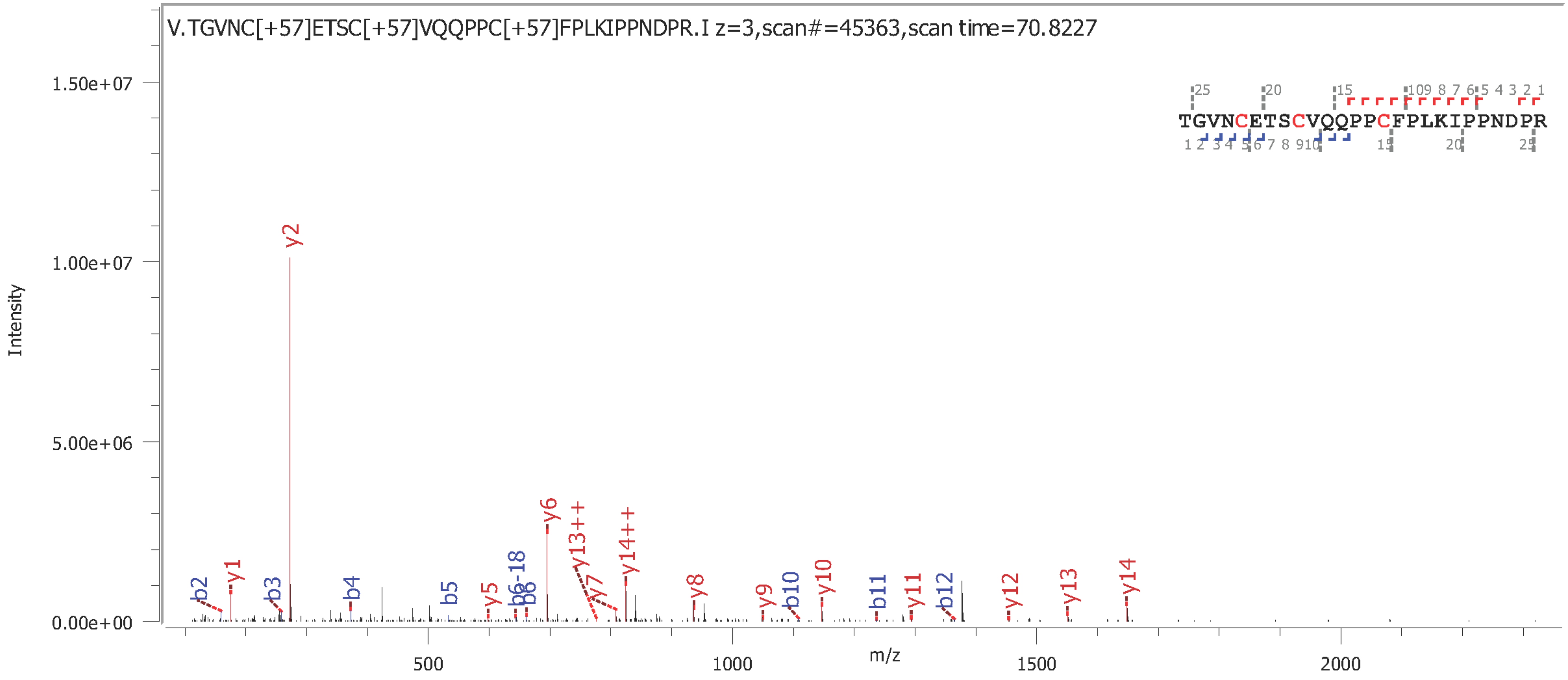

Human myeloperoxidase (P05164)

**β-chain N-termini** (truncation variant)

[278]GVNCETSCVQQPPCFPLKIPPNDPR (identification confidence level, PEP 2D: $1.1x10^{-15}$)

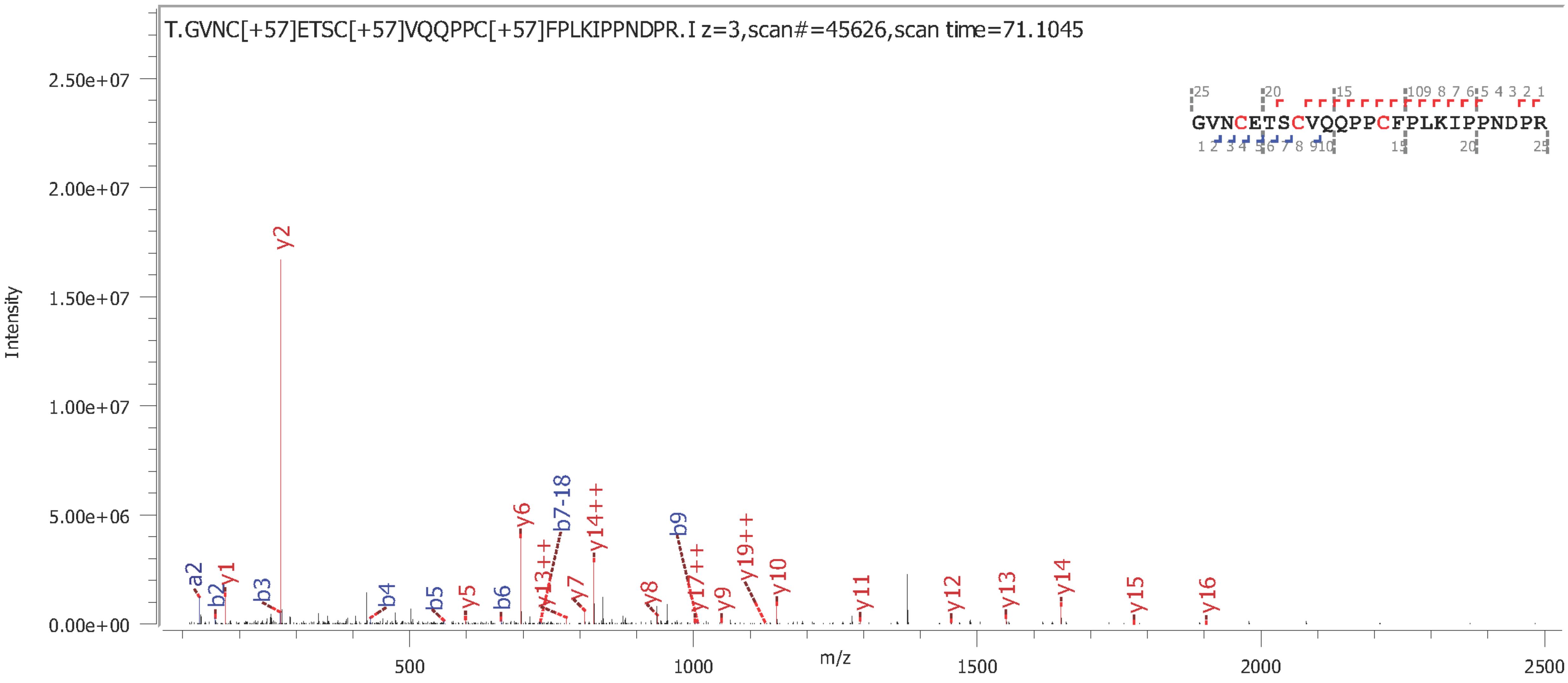

Human myeloperoxidase (P05164)

**β-chain N-terminal** (truncation variant)

[279]VNCETSCVQQPPCFPLKIPPNDPR (identification confidence level, PEP 2D: $1.5 \times 10^{-17}$)

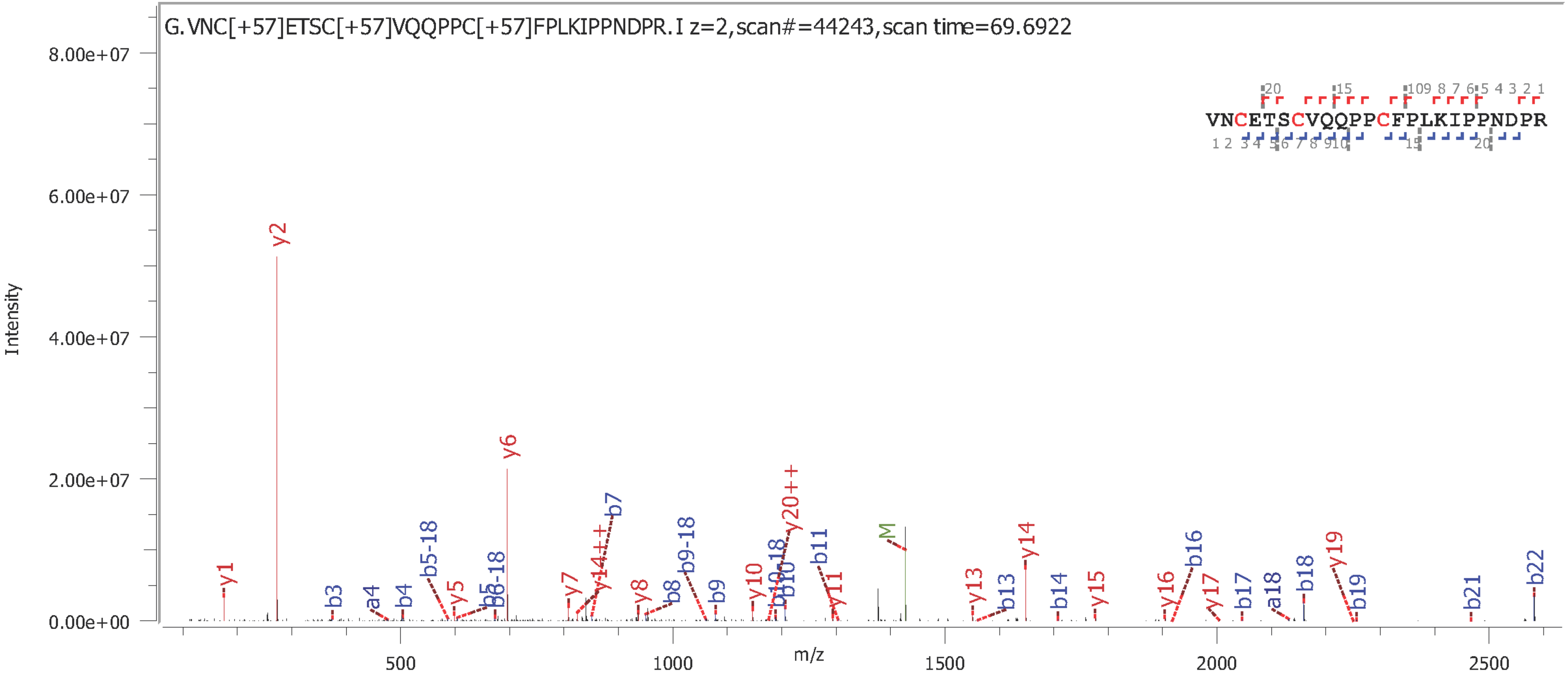

Human myeloperoxidase (P05164)

**β-chain C-terminal**

DFVNCSTLPALNLASWREA**S**[745] (identification confidence level, PEP 2D: $1.4 \times 10^{-8}$)

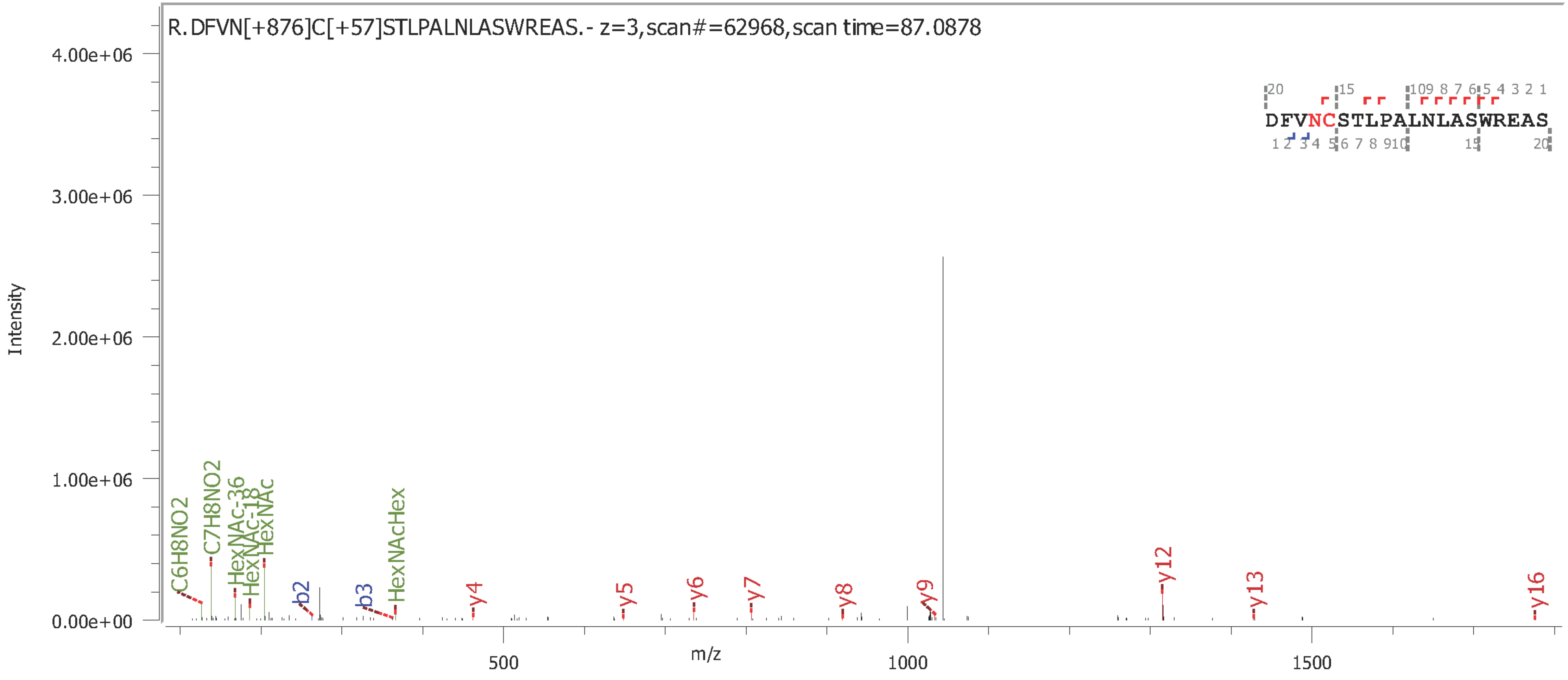

**Supplementary Data S3A**

Manually annotated reversed phase-LC-ESI-HCD-MS/MS (+) spectra of all glycosylated and non-glycosylated peptides containing Asn323 identified from the analysis of the unenriched peptide mixtures of nMPO.

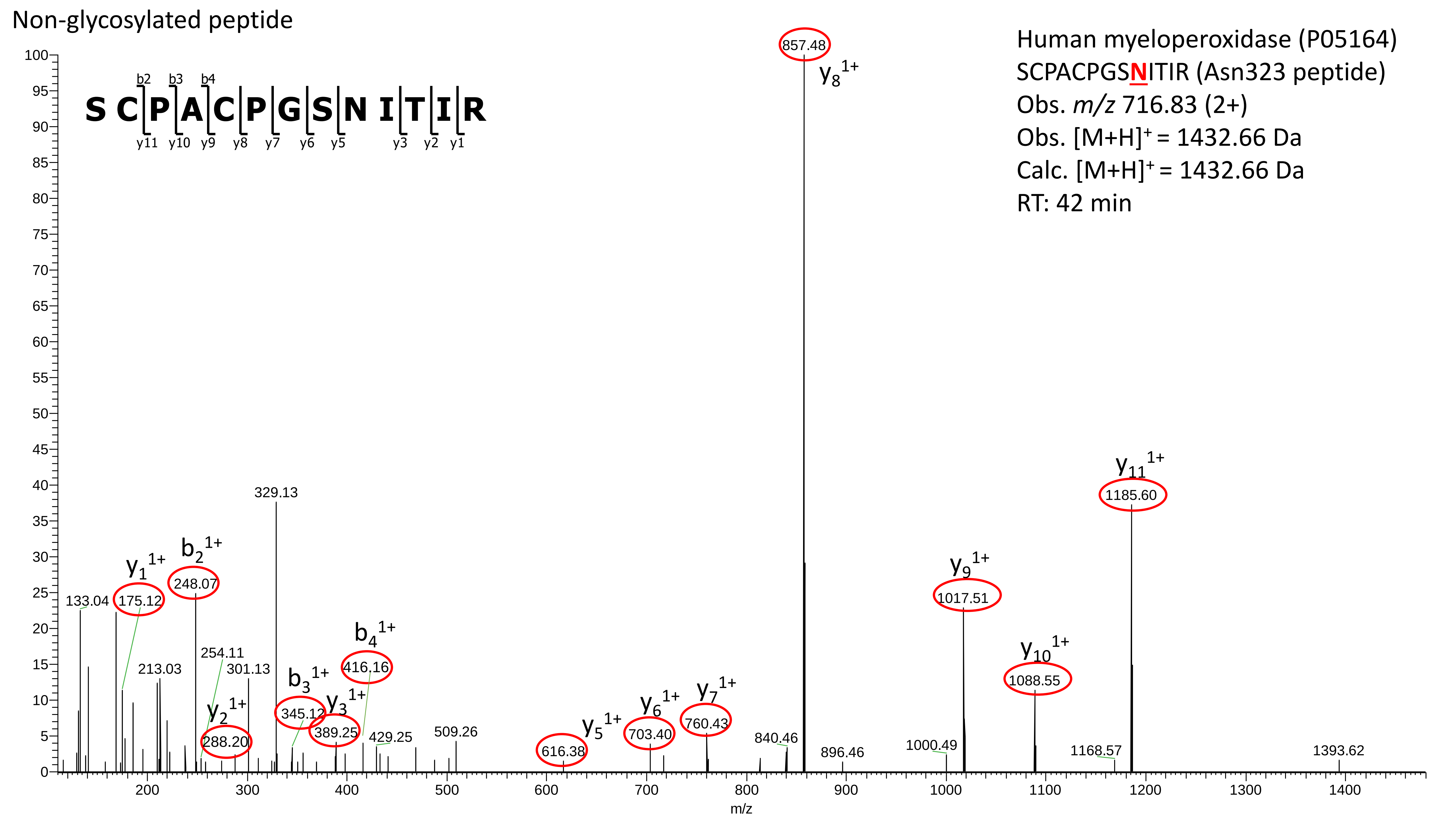
Non-glycosylated peptide
Human myeloperoxidase (P05164)
SCPACPGSNITIR (Asn323 peptide)
Obs. m/z 716.83 (2+)
Obs. [M+H]+ = 1432.66 Da
Calc. [M+H]+ = 1432.66 Da
RT: 42 min
S C P A C P G S N I T I R
b2 b3 b4
y11 y10 y9 y8 y7 y6 y5 y3 y2 y1
857.48 y8 1+
y11 1+ 1185.60
y9 1+ 1017.51
y10 1+ 1088.55
329.13
y1 1+ 175.12
b2 1+ 248.07
133.04
254.11
213.03
301.13
b4 1+ 416.16
b3 1+ 345.12
y3 1+ 389.25
y2 1+ 288.20
429.25
509.26
y5 1+ 616.38
y6 1+ 703.40
y7 1+ 760.43
840.46
896.46
1000.49
1168.57
1393.62
m/z

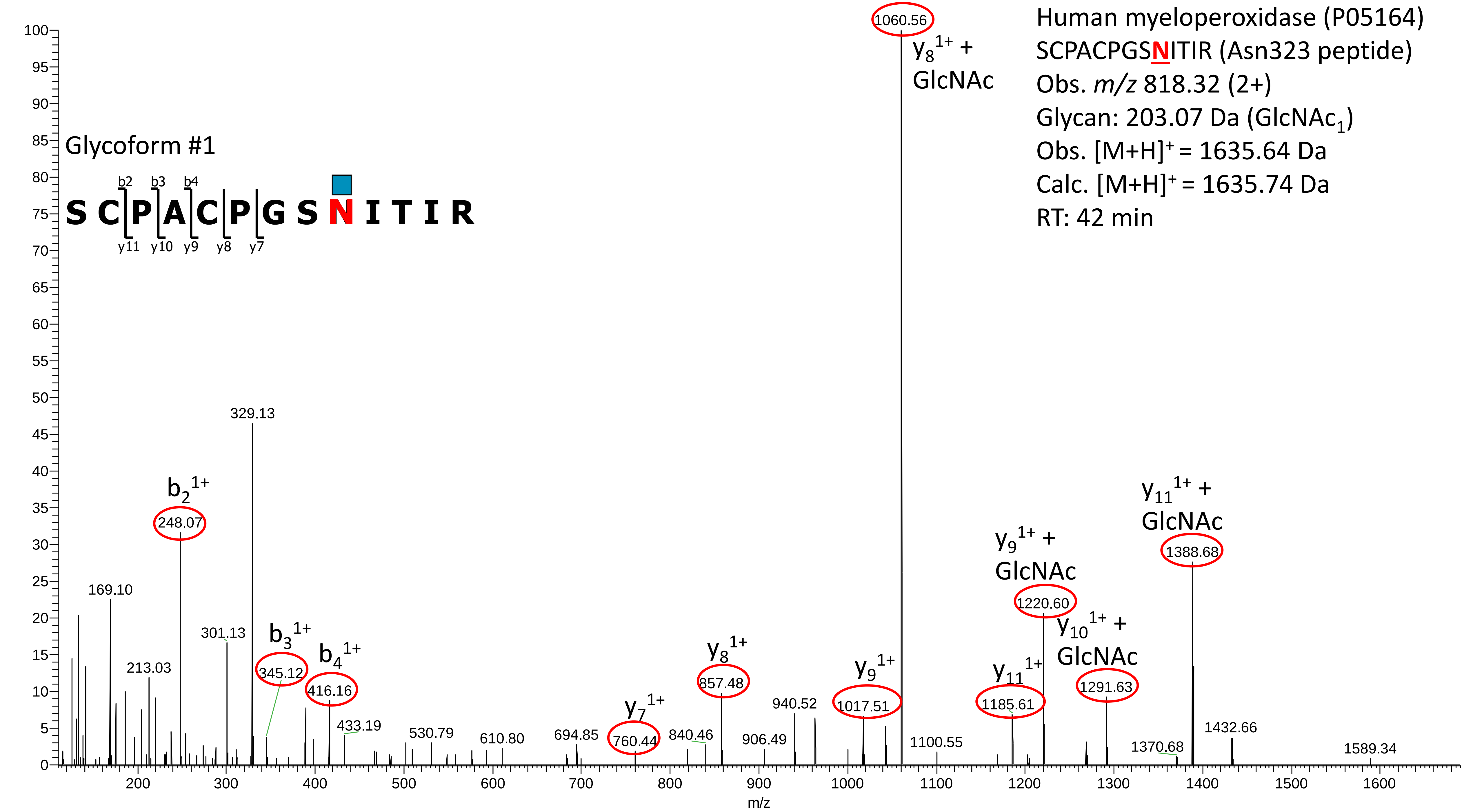

Human myeloperoxidase (P05164)
SCPACPGSNITIR (Asn323 peptide)
Obs. m/z 818.32 (2+)
Glycan: 203.07 Da (GlcNAc1)
Obs. [M+H]+ = 1635.64 Da
Calc. [M+H]+ = 1635.74 Da
RT: 42 min
Glycoform #1
S C P A C P G S N I T I R
b2 b3 b4
y11 y10 y9 y8 y7
y8 1+ + GlcNAc
1060.56
b2 1+
248.07
329.13
169.10
301.13
213.03
b3 1+
345.12
b4 1+
416.16
433.19
530.79
610.80
694.85
y7 1+
760.44
840.46
y8 1+
857.48
906.49
940.52
y9 1+
1017.51
1100.55
y11 1+
1185.61
y9 1+ + GlcNAc
1220.60
y10 1+ + GlcNAc
1291.63
1370.68
y11 1+ + GlcNAc
1388.68
1432.66
1589.34
m/z

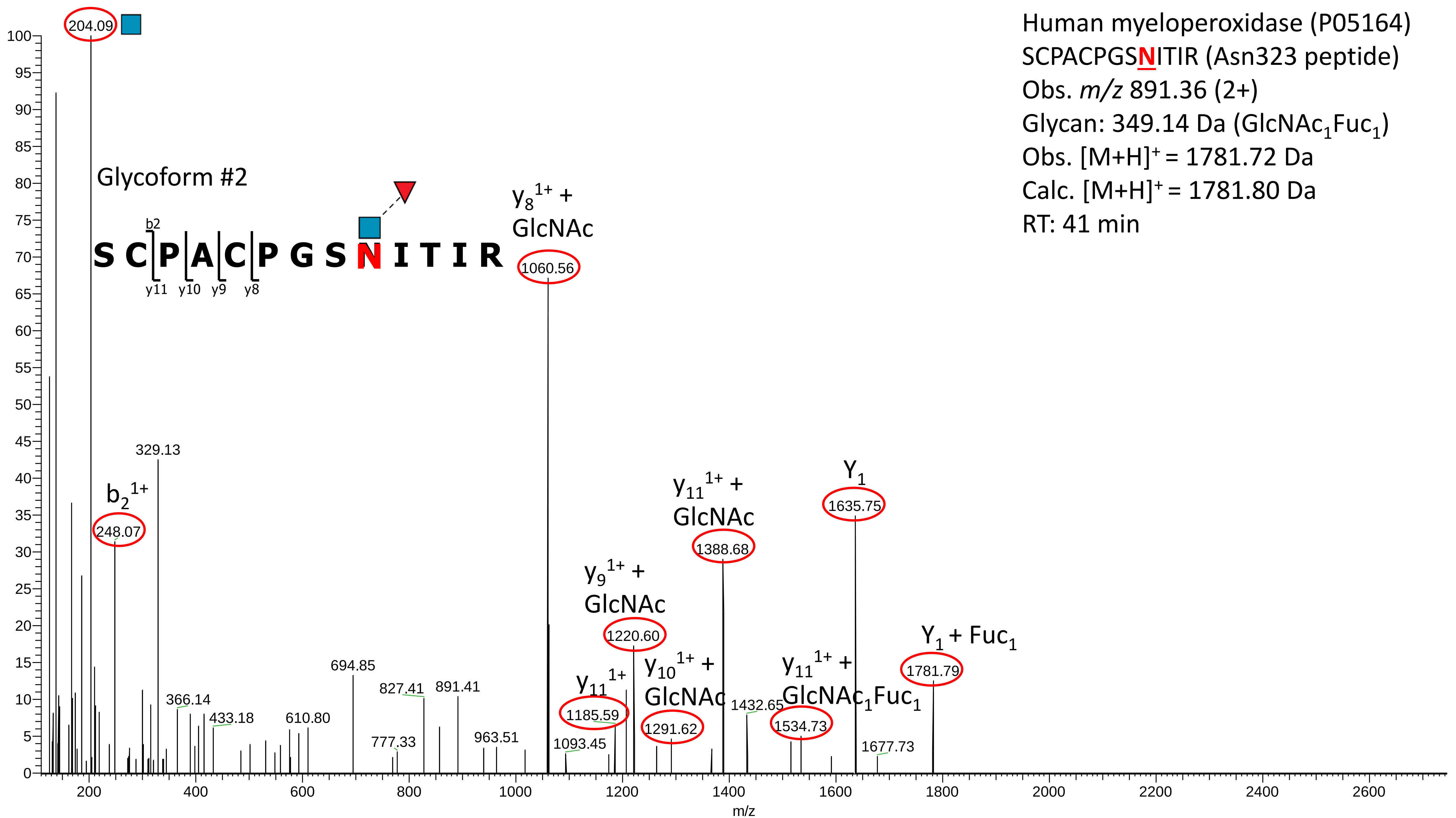
Human myeloperoxidase (P05164)
SCPACPGSNITIR (Asn323 peptide)
Obs. m/z 891.36 (2+)
Glycan: 349.14 Da (GlcNAc1Fuc1)
Obs. [M+H]+ = 1781.72 Da
Calc. [M+H]+ = 1781.80 Da
RT: 41 min
Glycoform #2
S C P A C P G S N I T I R
b2
y11 y10 y9 y8
204.09
329.13
b2 1+
248.07
y8 1+ + GlcNAc
1060.56
y9 1+ + GlcNAc
1220.60
y11 1+
1185.59
y10 1+ + GlcNAc
1291.62
y11 1+ + GlcNAc
1388.68
1432.65
y11 1+ + GlcNAc1Fuc1
1534.73
Y1
1635.75
1677.73
Y1 + Fuc1
1781.79
366.14
433.18
610.80
694.85
777.33
827.41
891.41
963.51
1093.45
m/z

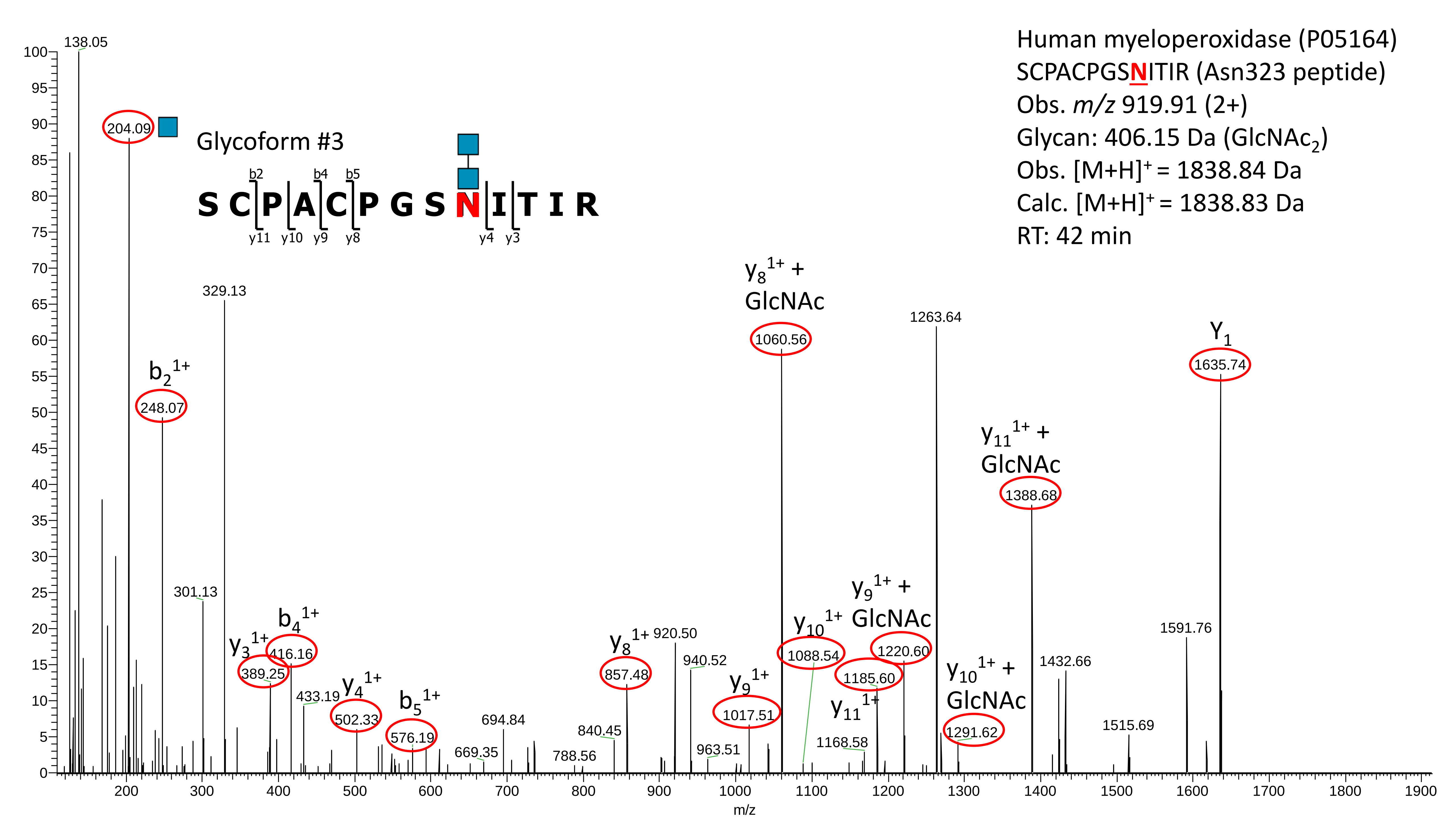

Human myeloperoxidase (P05164)
SCPACPGSNITIR (Asn323 peptide)
Obs. m/z 919.91 (2+)
Glycan: 406.15 Da (GlcNAc2)
Obs. [M+H]+ = 1838.84 Da
Calc. [M+H]+ = 1838.83 Da
RT: 42 min
Glycoform #3
b2 b4 b5
S C P A C P G S N I T I R
y11 y10 y9 y8 y4 y3
138.05
204.09
b2 1+
248.07
329.13
301.13
y3 1+
389.25
b4 1+
416.16
433.19
y4 1+
502.33
b5 1+
576.19
669.35
694.84
788.56
840.45
y8 1+
857.48
920.50
940.52
963.51
y9 1+
1017.51
y8 1+ + GlcNAc
1060.56
y10 1+
1088.54
1168.58
y11 1+
1185.60
y9 1+ + GlcNAc
1220.60
1263.64
y10 1+ + GlcNAc
1291.62
y11 1+ + GlcNAc
1388.68
1432.66
1515.69
1591.76
Y1
1635.74
m/z

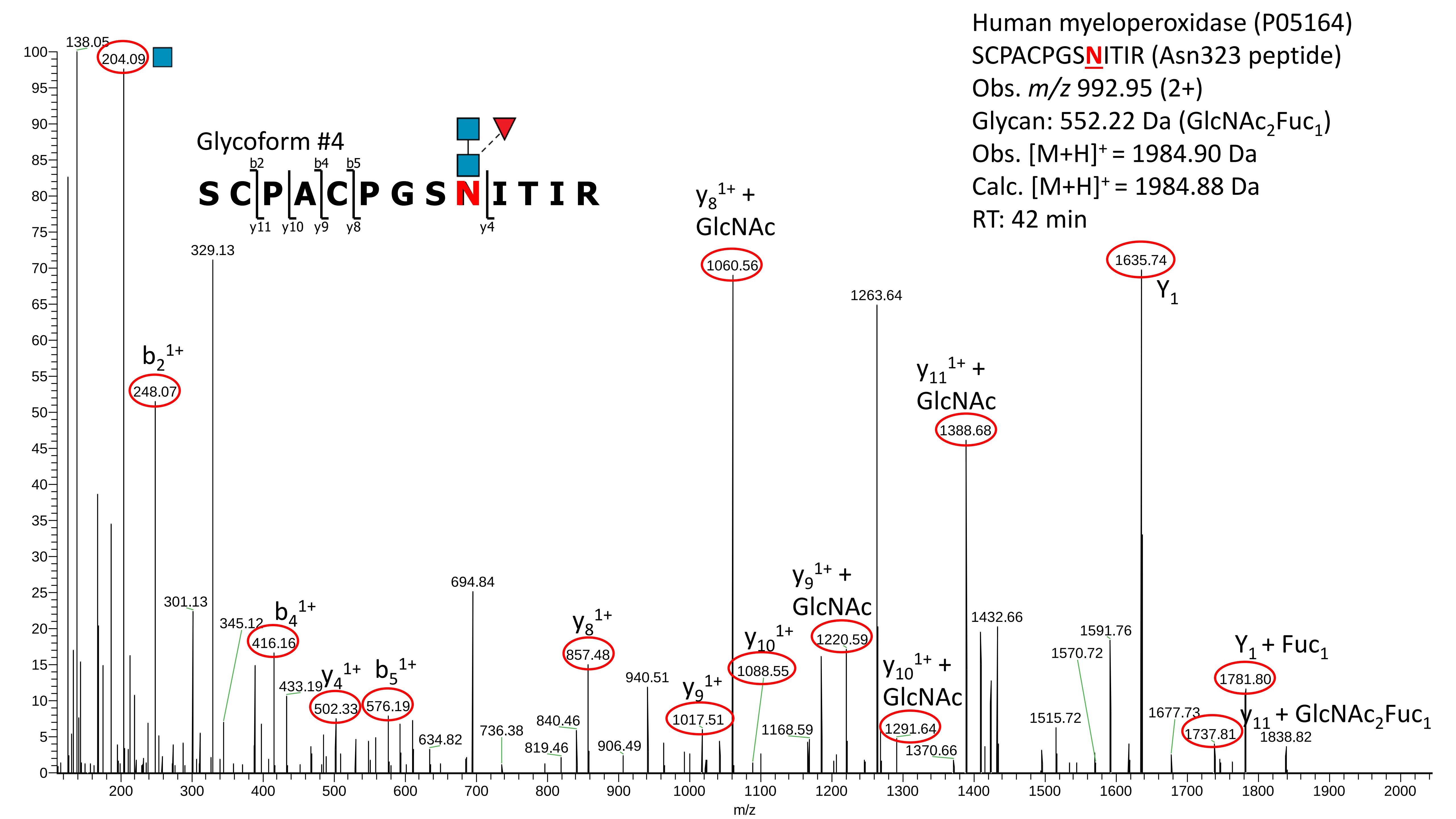
Human myeloperoxidase (P05164)
SCPACPGSNITIR (Asn323 peptide)
Obs. m/z 992.95 (2+)
Glycan: 552.22 Da (GlcNAc2Fuc1)
Obs. [M+H]+ = 1984.90 Da
Calc. [M+H]+ = 1984.88 Da
RT: 42 min
Glycoform #4
S C P A C P G S N I T I R
b2 b4 b5
y11 y10 y9 y8 y4
138.05
204.09
b2 1+
248.07
329.13
301.13
345.12
b4 1+
416.16
433.19
y4 1+
502.33
b5 1+
576.19
634.82
694.84
736.38
819.46
840.46
y8 1+
857.48
906.49
940.51
y9 1+
1017.51
y8 1+ + GlcNAc
1060.56
y10 1+
1088.55
1168.59
y9 1+ + GlcNAc
1220.59
1263.64
y10 1+ + GlcNAc
1291.64
1370.66
y11 1+ + GlcNAc
1388.68
1432.66
1515.72
1570.72
1591.76
1635.74
Y1
1677.73
1737.81
y11 + GlcNAc2Fuc1
Y1 + Fuc1
1781.80
1838.82
m/z

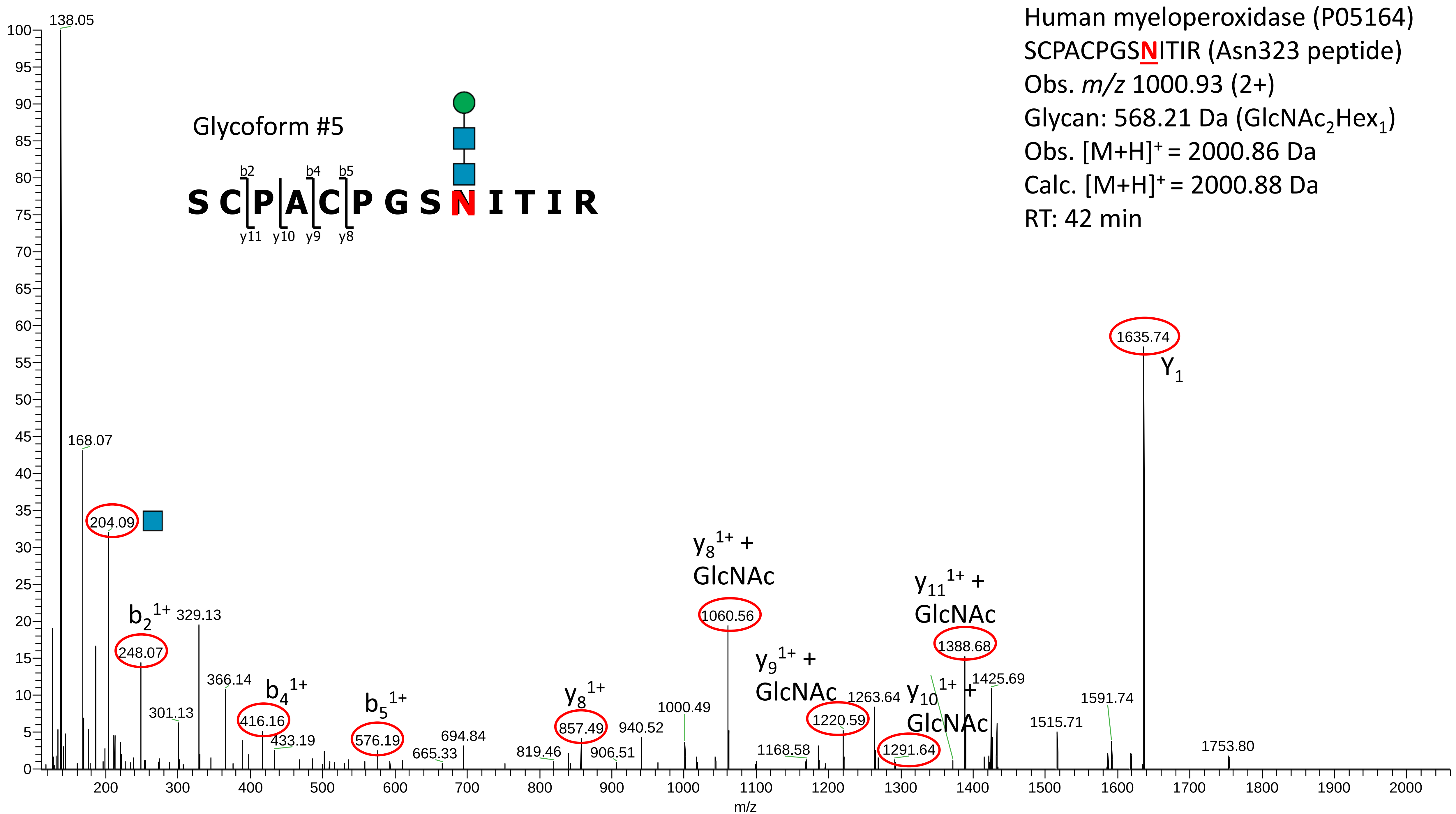

Human myeloperoxidase (P05164)
SCPACPGSNITIR (Asn323 peptide)
Obs. m/z 1000.93 (2+)
Glycan: 568.21 Da (GlcNAc2Hex1)
Obs. [M+H]+ = 2000.86 Da
Calc. [M+H]+ = 2000.88 Da
RT: 42 min
Glycoform #5
S C P A C P G S N I T I R
b2 b4 b5
y11 y10 y9 y8
138.05
168.07
204.09
b2 1+ 248.07
329.13
301.13
366.14
b4 1+ 416.16
433.19
b5 1+ 576.19
665.33
694.84
819.46
y8 1+ 857.49
906.51
940.52
1000.49
y8 1+ + GlcNAc 1060.56
1168.58
y9 1+ + GlcNAc 1220.59
1263.64
y10 1+ + GlcNAc 1291.64
y11 1+ + GlcNAc 1388.68
1425.69
1515.71
1591.74
1635.74 Y1
1753.80
m/z

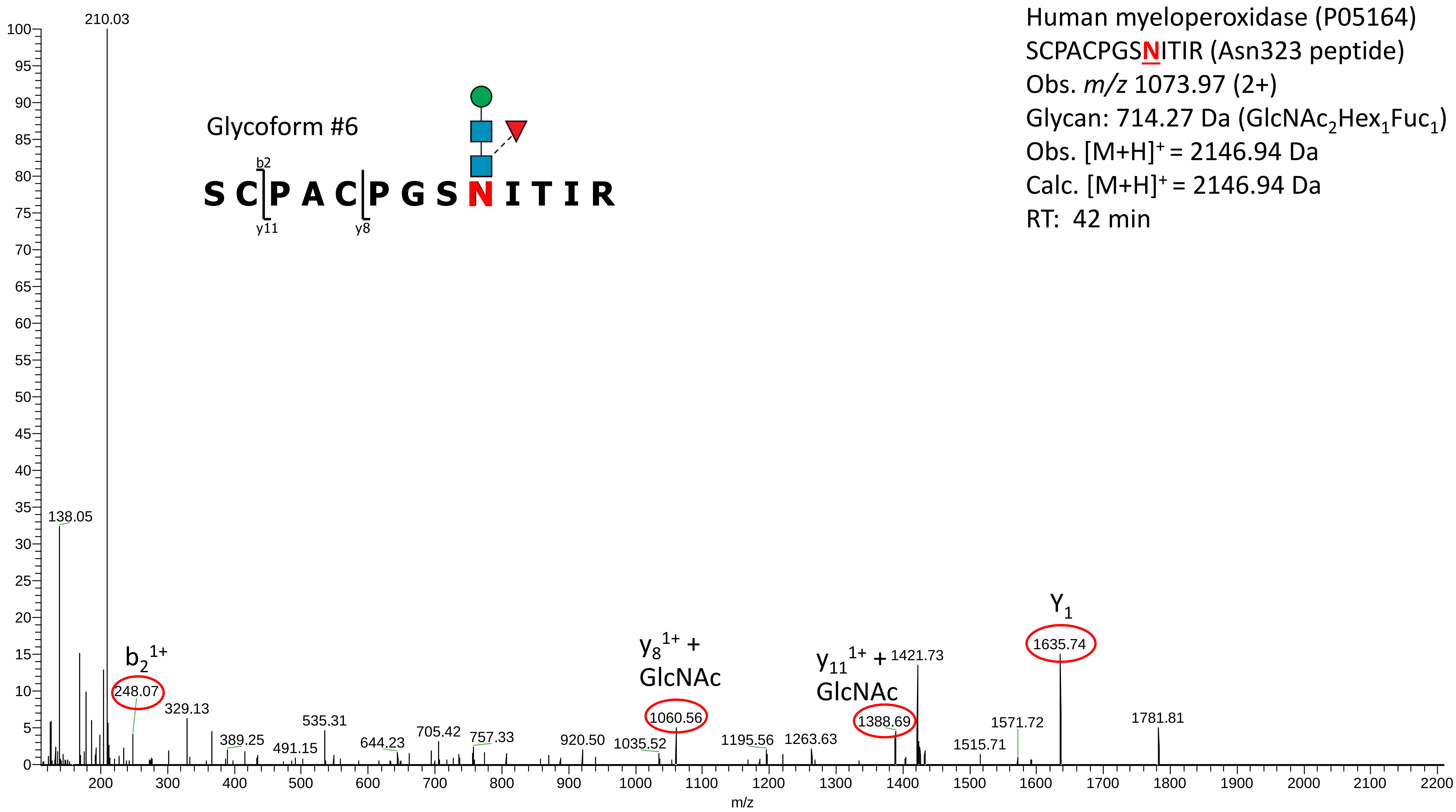

Human myeloperoxidase (P05164)
SCPACPGSNITIR (Asn323 peptide)
Obs. m/z 1073.97 (2+)
Glycan: 714.27 Da (GlcNAc2Hex1Fuc1)
Obs. [M+H]+ = 2146.94 Da
Calc. [M+H]+ = 2146.94 Da
RT: 42 min
Glycoform #6
S C P A C P G S N I T I R
b2
y11
y8
210.03
138.05
b2 1+
248.07
329.13
389.25
491.15
535.31
644.23
705.42
757.33
920.50
1035.52
y8 1+ + GlcNAc
1060.56
1195.56
1263.63
y11 1+ + GlcNAc
1388.69
1421.73
1515.71
1571.72
Y1
1635.74
1781.81
m/z

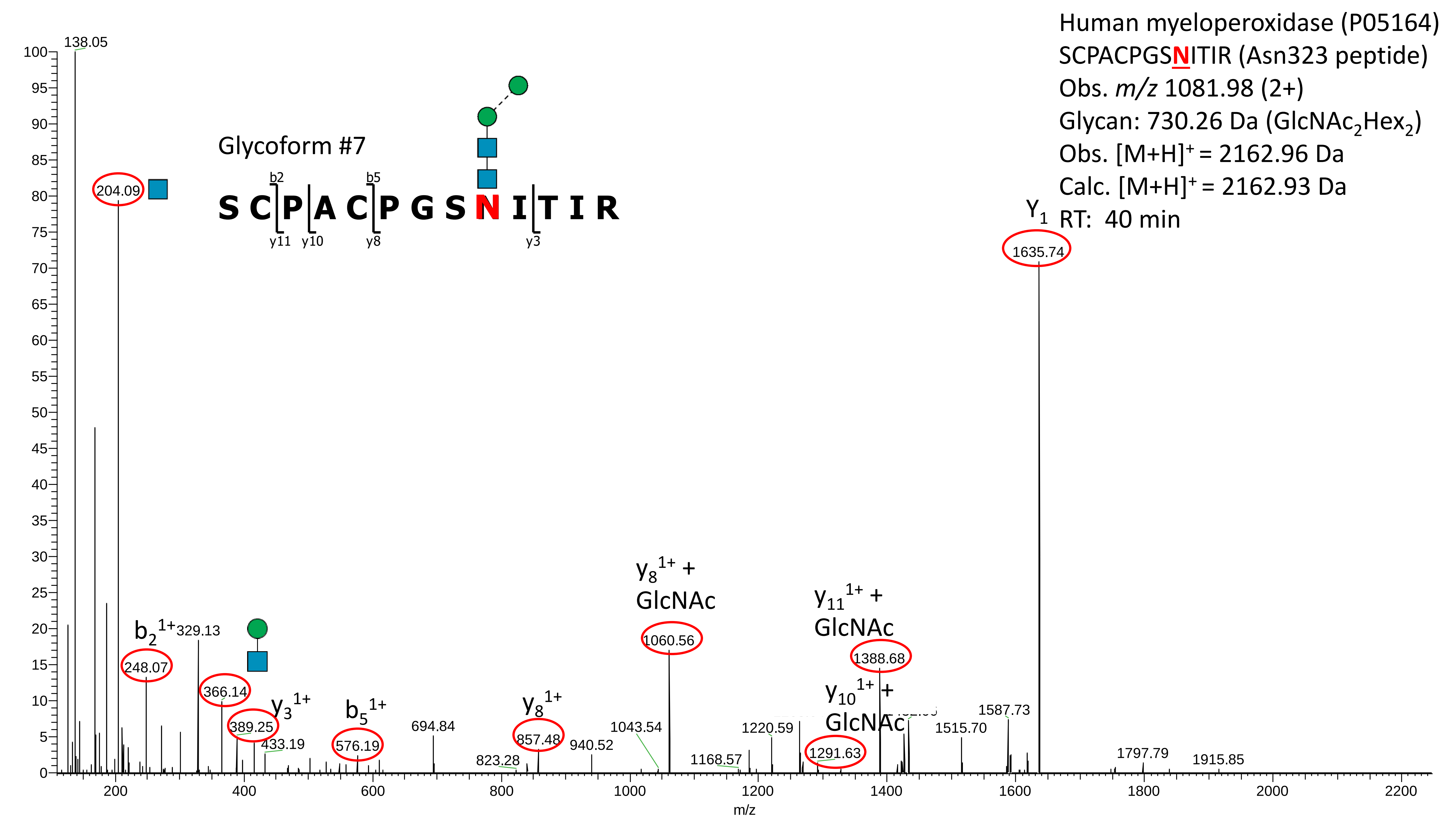
Human myeloperoxidase (P05164)
SCPACPGSNITIR (Asn323 peptide)
Obs. m/z 1081.98 (2+)
Glycan: 730.26 Da (GlcNAc2Hex2)
Obs. [M+H]+ = 2162.96 Da
Calc. [M+H]+ = 2162.93 Da
RT: 40 min
Glycoform #7
S C P A C P G S N I T I R
b2
b5
y11
y10
y8
y3
Y1
1635.74
138.05
204.09
b2 1+
248.07
329.13
366.14
389.25
y3 1+
433.19
b5 1+
576.19
694.84
823.28
y8 1+
857.48
940.52
1043.54
y8 1+ + GlcNAc
1060.56
1168.57
1220.59
y11 1+ + GlcNAc
1388.68
y10 1+ + GlcNAc
1291.63
1515.70
1587.73
1797.79
1915.85
m/z

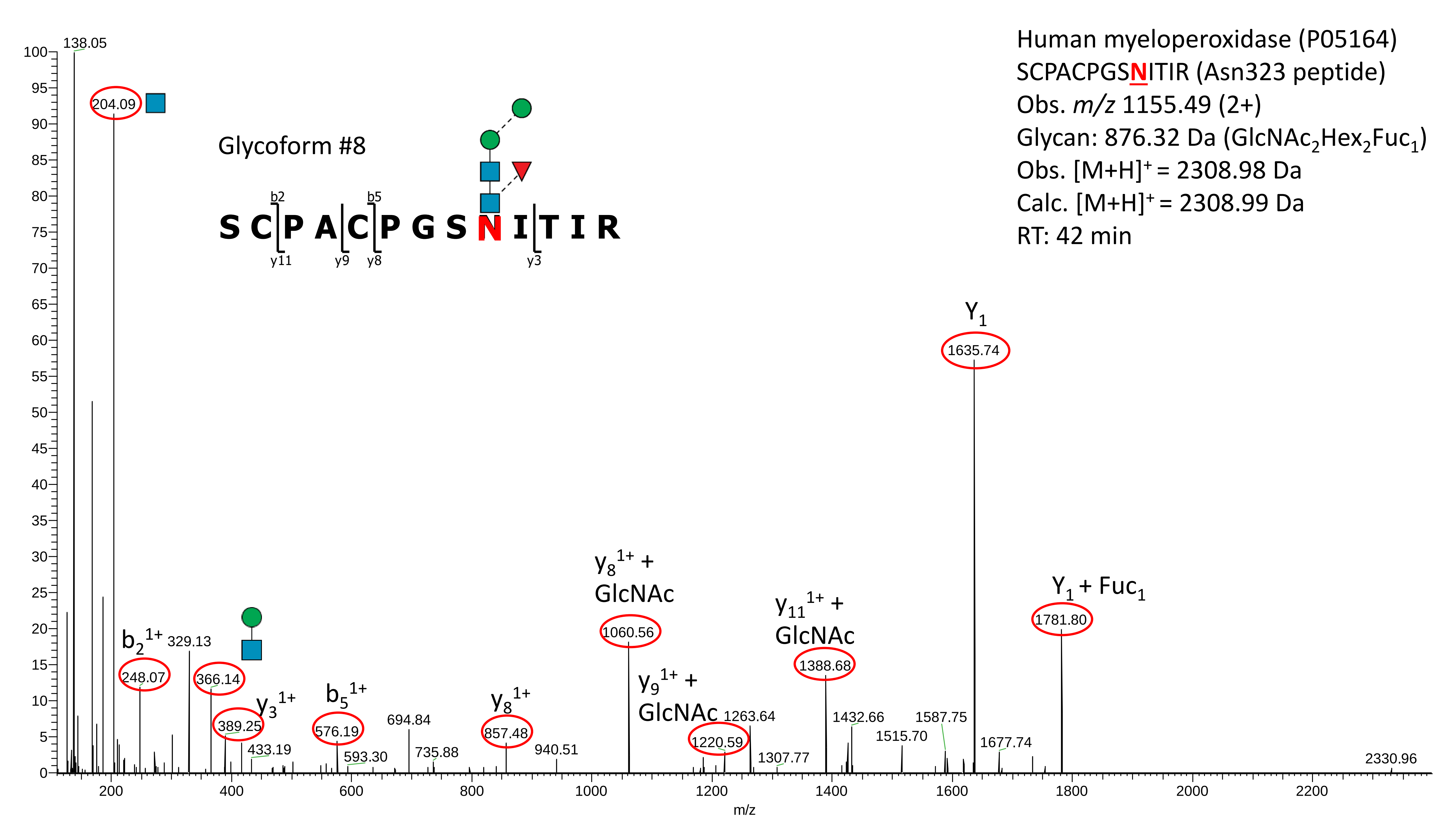
Human myeloperoxidase (P05164)
SCPACPGSNITIR (Asn323 peptide)
Obs. m/z 1155.49 (2+)
Glycan: 876.32 Da (GlcNAc2Hex2Fuc1)
Obs. [M+H]+ = 2308.98 Da
Calc. [M+H]+ = 2308.99 Da
RT: 42 min
Glycoform #8
S C P A C P G S N I T I R
b2
b5
y11
y9
y8
y3
138.05
204.09
b2 1+
248.07
329.13
366.14
389.25
y3 1+
433.19
b5 1+
576.19
593.30
694.84
735.88
y8 1+
857.48
940.51
y8 1+ + GlcNAc
1060.56
y9 1+ + GlcNAc
1220.59
1263.64
1307.77
y11 1+ + GlcNAc
1388.68
1432.66
1515.70
1587.75
Y1
1635.74
1677.74
Y1 + Fuc1
1781.80
2330.96
m/z

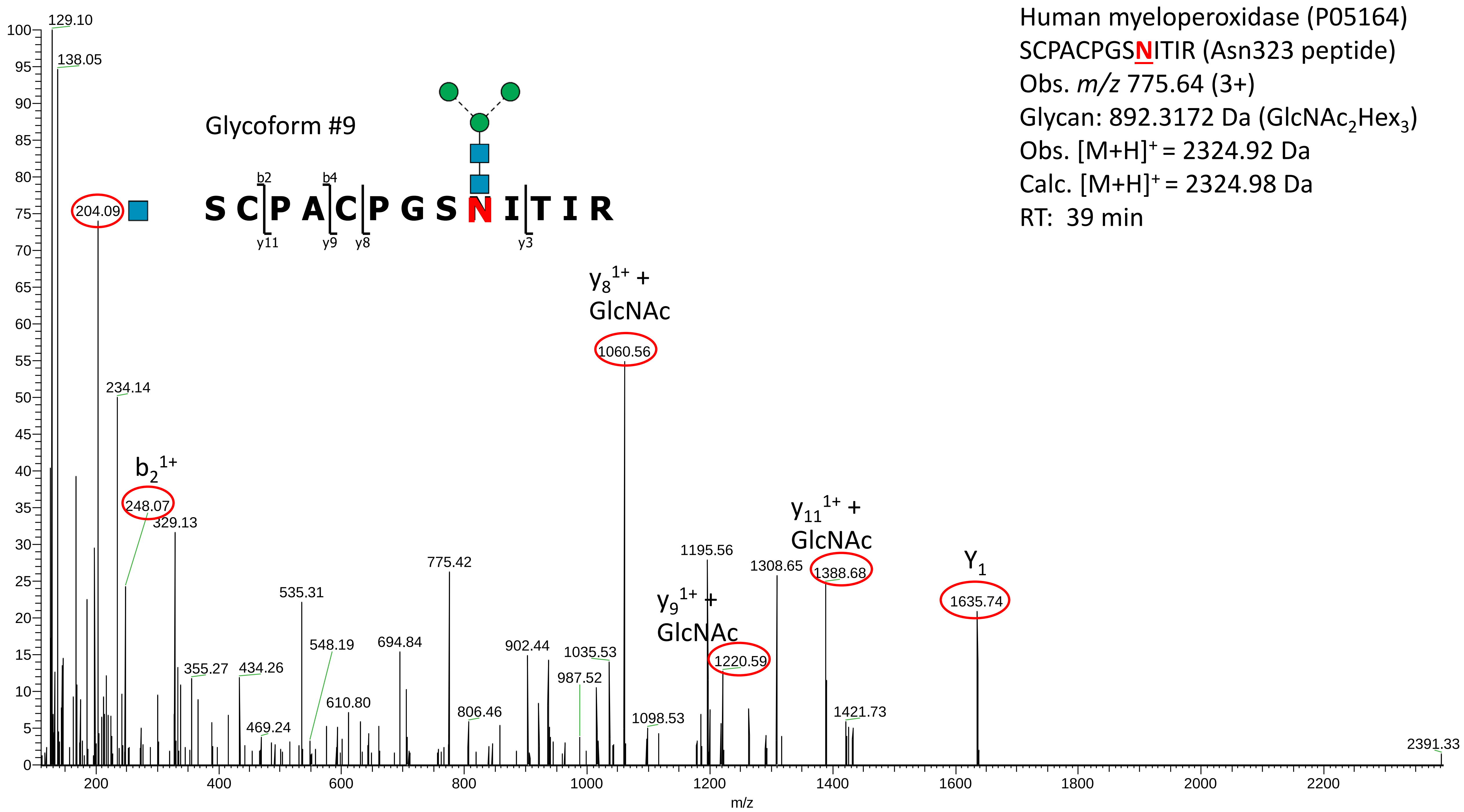
Human myeloperoxidase (P05164)
SCPACPGSNITIR (Asn323 peptide)
Obs. m/z 775.64 (3+)
Glycan: 892.3172 Da (GlcNAc2Hex3)
Obs. [M+H]+ = 2324.92 Da
Calc. [M+H]+ = 2324.98 Da
RT: 39 min
Glycoform #9
S C P A C P G S N I T I R
b2
b4
y11
y9
y8
y3
y8 1+ + GlcNAc
y9 1+ + GlcNAc
y11 1+ + GlcNAc
Y1
b2 1+
129.10
138.05
204.09
234.14
248.07
329.13
355.27
434.26
469.24
535.31
548.19
610.80
694.84
775.42
806.46
902.44
987.52
1035.53
1060.56
1098.53
1195.56
1220.59
1308.65
1388.68
1421.73
1635.74
2391.33
m/z

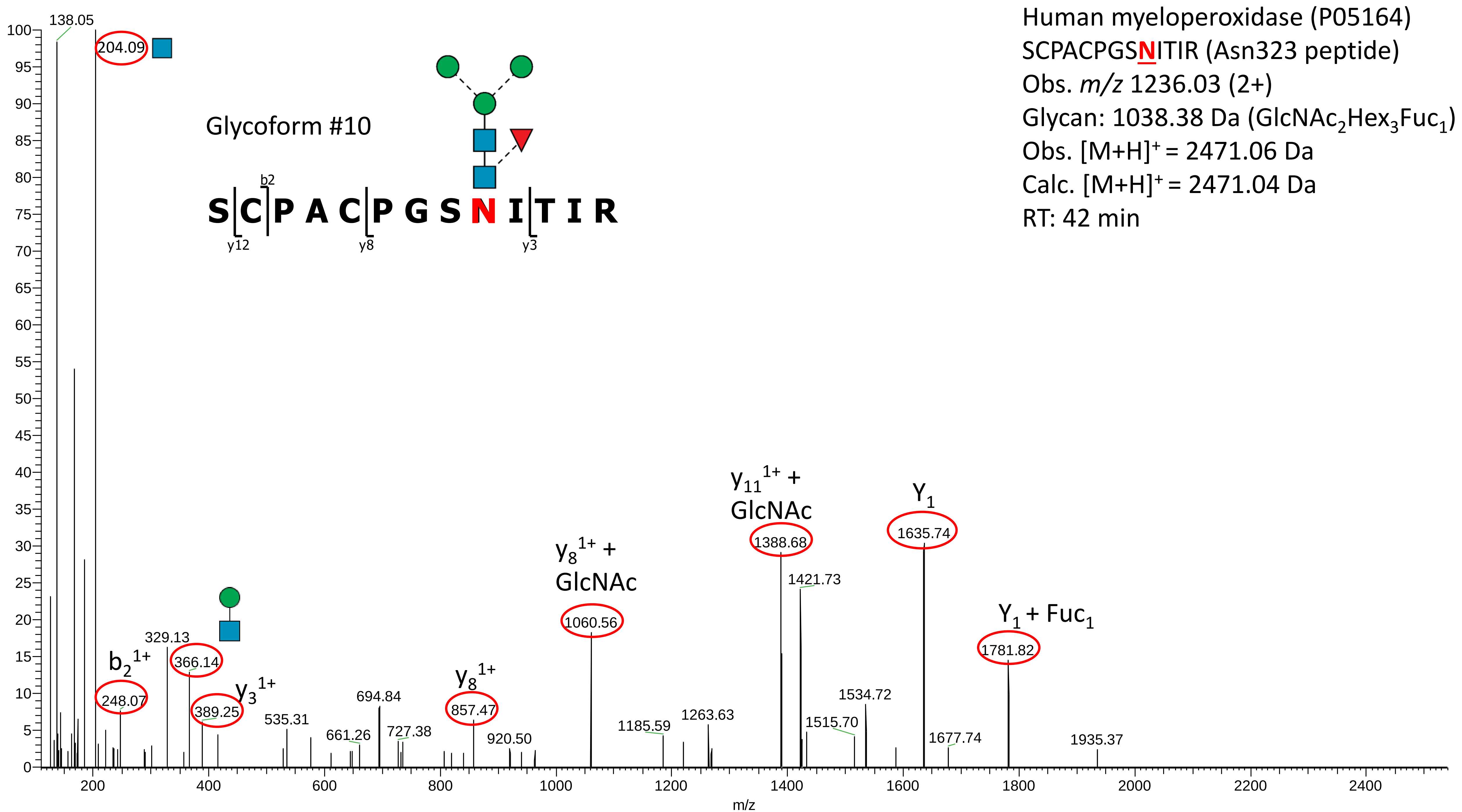

Human myeloperoxidase (P05164)
SCPACPGSNITIR (Asn323 peptide)
Obs. m/z 1236.03 (2+)
Glycan: 1038.38 Da (GlcNAc2Hex3Fuc1)
Obs. [M+H]+ = 2471.06 Da
Calc. [M+H]+ = 2471.04 Da
RT: 42 min
Glycoform #10
S C P A C P G S N I T I R
b2
y12
y8
y3
138.05
204.09
248.07
b2 1+
329.13
366.14
389.25
y3 1+
535.31
661.26
694.84
727.38
857.47
y8 1+
920.50
1060.56
y8 1+ + GlcNAc
1185.59
1263.63
1388.68
y11 1+ + GlcNAc
1421.73
1515.70
1534.72
1635.74
Y1
1677.74
1781.82
Y1 + Fuc1
1935.37
m/z

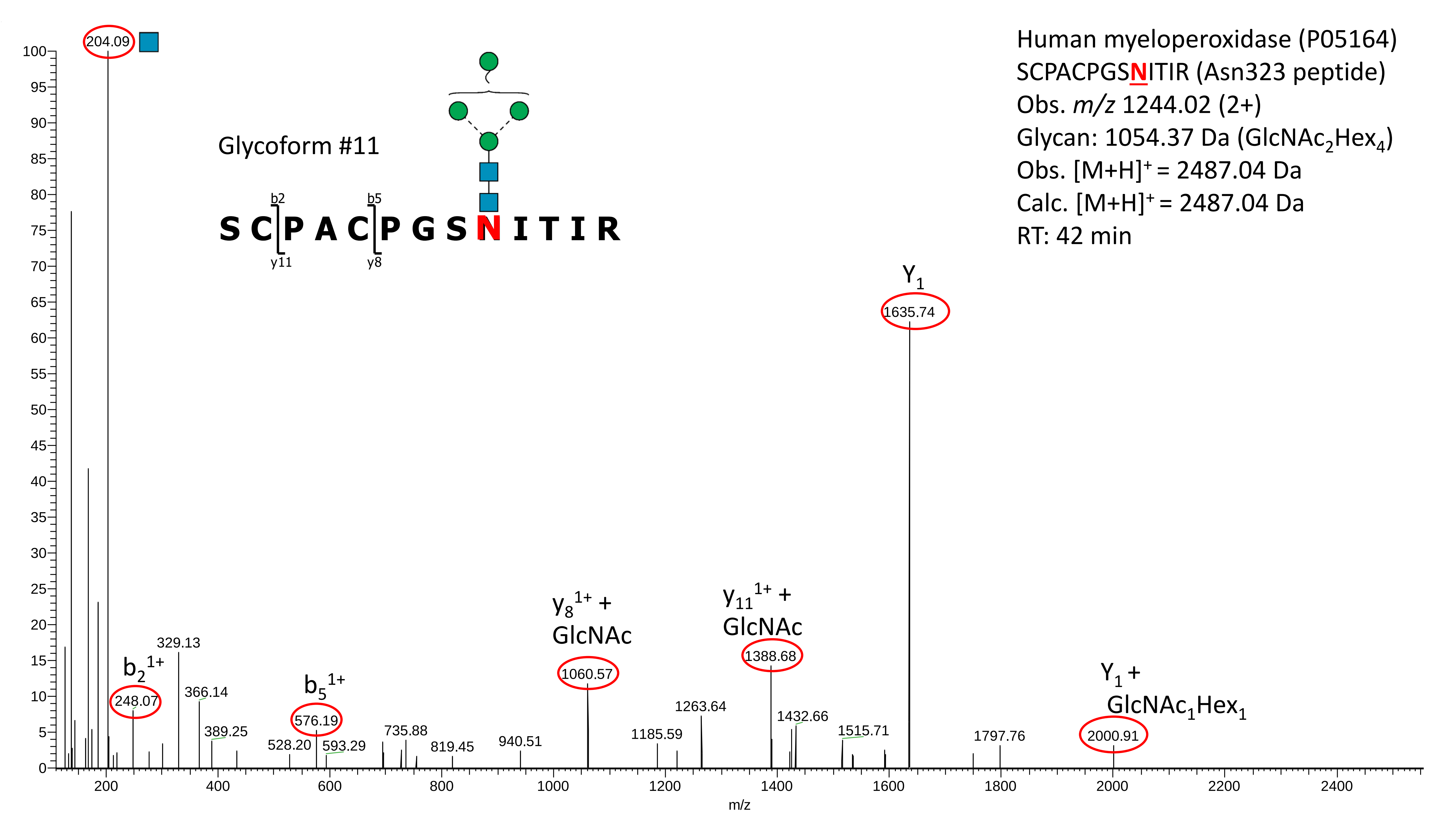

Human myeloperoxidase (P05164)
SCPACPGSNITIR (Asn323 peptide)
Obs. m/z 1244.02 (2+)
Glycan: 1054.37 Da (GlcNAc2Hex4)
Obs. [M+H]+ = 2487.04 Da
Calc. [M+H]+ = 2487.04 Da
RT: 42 min
Glycoform #11
S C P A C P G S N I T I R
b2
b5
y11
y8
204.09
Y1
1635.74
b2 1+
248.07
329.13
366.14
389.25
b5 1+
576.19
528.20
593.29
735.88
819.45
940.51
y8 1+ + GlcNAc
1060.57
1185.59
1263.64
y11 1+ + GlcNAc
1388.68
1432.66
1515.71
1797.76
Y1 + GlcNAc1Hex1
2000.91
m/z

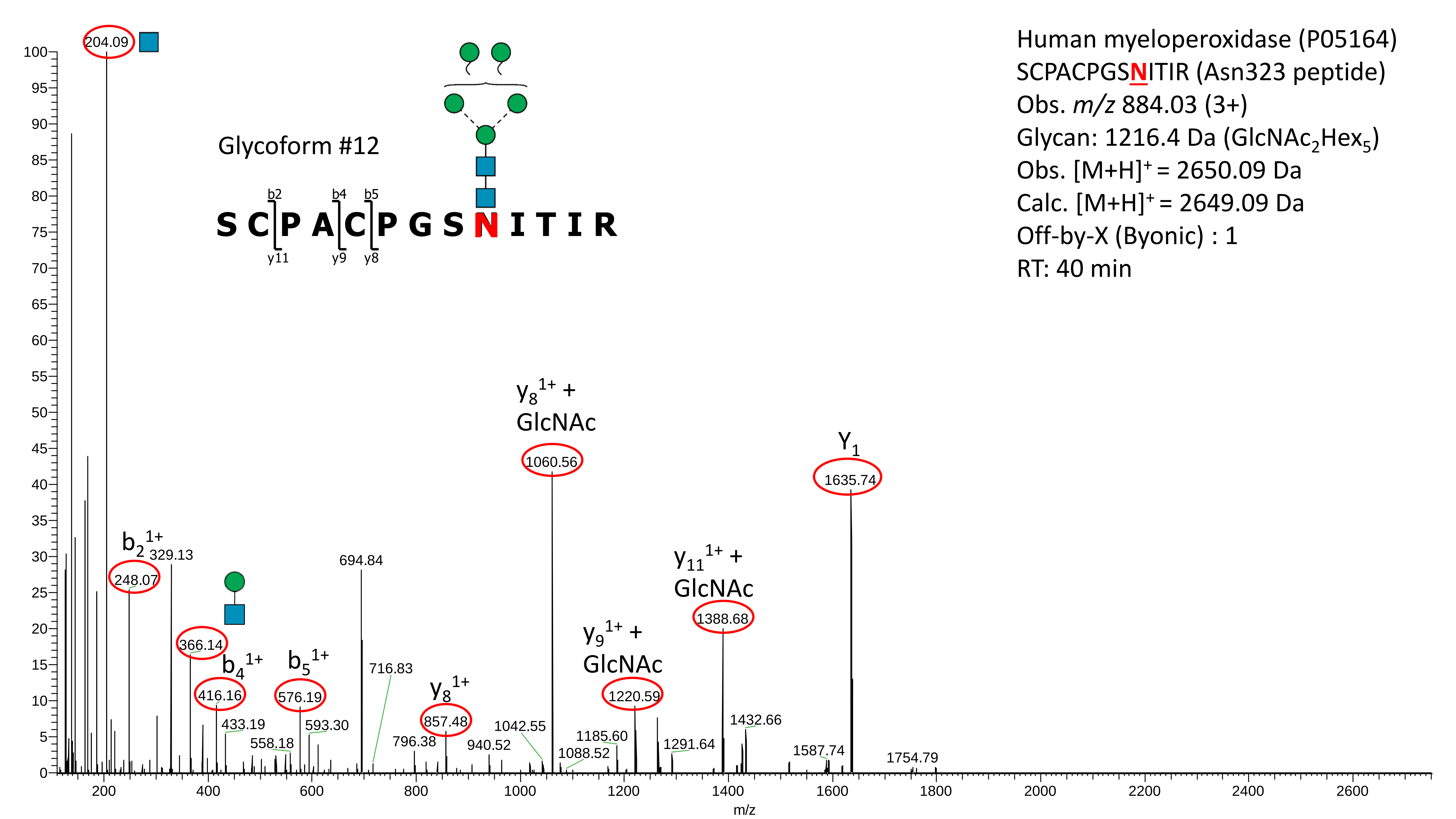
Human myeloperoxidase (P05164)
SCPACPGSNITIR (Asn323 peptide)
Obs. m/z 884.03 (3+)
Glycan: 1216.4 Da (GlcNAc2Hex5)
Obs. [M+H]+ = 2650.09 Da
Calc. [M+H]+ = 2649.09 Da
Off-by-X (Byonic) : 1
RT: 40 min
Glycoform #12
S C P A C P G S N I T I R
b2
b4
b5
y11
y9
y8
204.09
b2 1+
248.07
329.13
366.14
b4 1+
416.16
433.19
b5 1+
576.19
558.18
593.30
694.84
716.83
796.38
y8 1+
857.48
940.52
1042.55
y8 1+ + GlcNAc
1060.56
1088.52
1185.60
y9 1+ + GlcNAc
1220.59
1291.64
y11 1+ + GlcNAc
1388.68
1432.66
1587.74
Y1
1635.74
1754.79
m/z

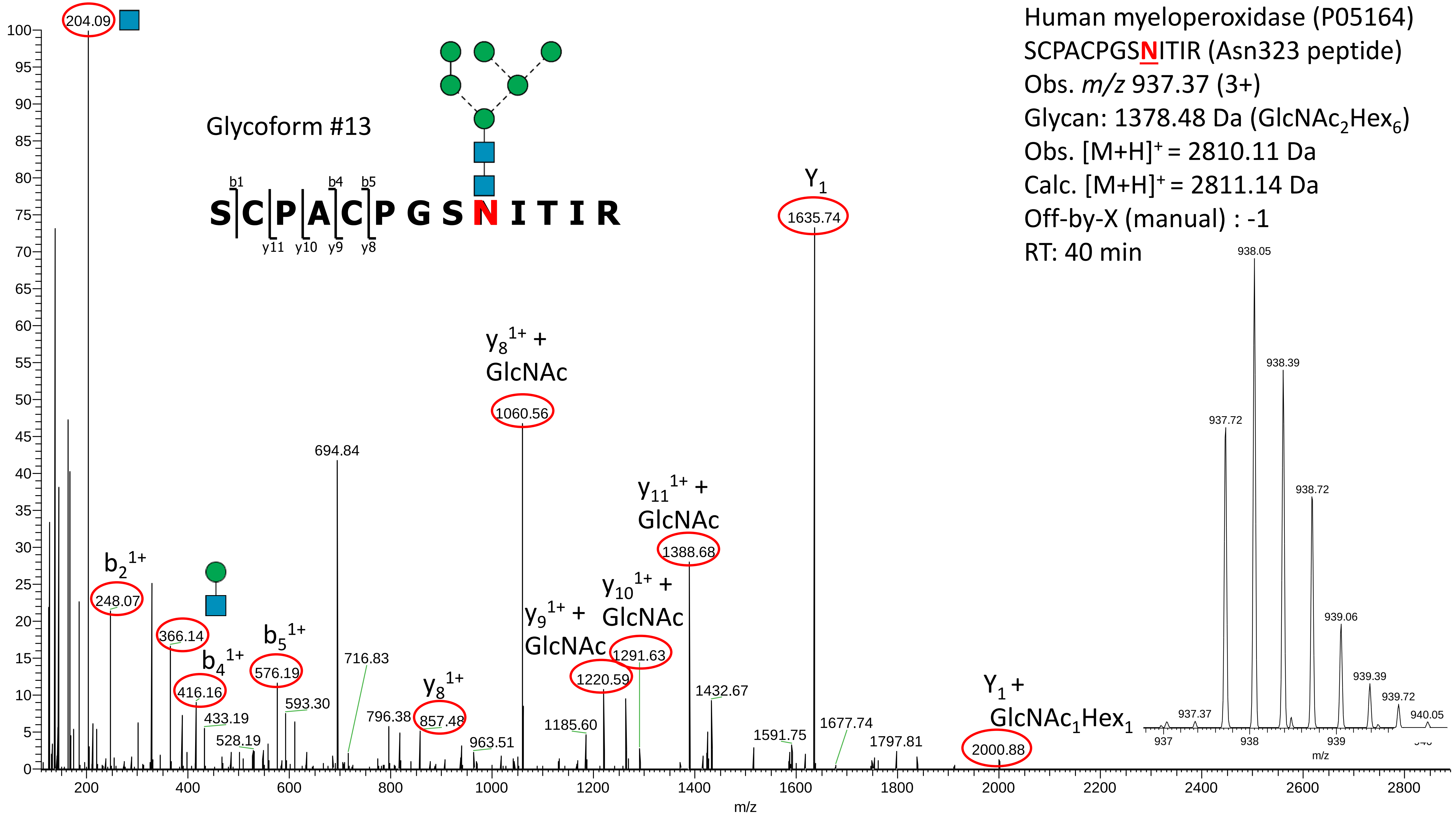

Human myeloperoxidase (P05164)
SCPACPGSNITIR (Asn323 peptide)
Obs. m/z 937.37 (3+)
Glycan: 1378.48 Da (GlcNAc2Hex6)
Obs. [M+H]+ = 2810.11 Da
Calc. [M+H]+ = 2811.14 Da
Off-by-X (manual) : -1
RT: 40 min
Glycoform #13
S C P A C P G S N I T I R
b1 b4 b5
y11 y10 y9 y8
204.09
b2 1+
248.07
366.14
b4 1+
416.16
433.19
528.19
b5 1+
576.19
593.30
694.84
716.83
796.38
y8 1+
857.48
963.51
y8 1+ + GlcNAc
1060.56
1185.60
y9 1+ + GlcNAc
1220.59
y10 1+ + GlcNAc
1291.63
y11 1+ + GlcNAc
1388.68
1432.67
1591.75
Y1
1635.74
1677.74
1797.81
Y1 + GlcNAc1Hex1
2000.88
937.37
937.72
938.05
938.39
938.72
939.06
939.39
939.72
940.05
m/z

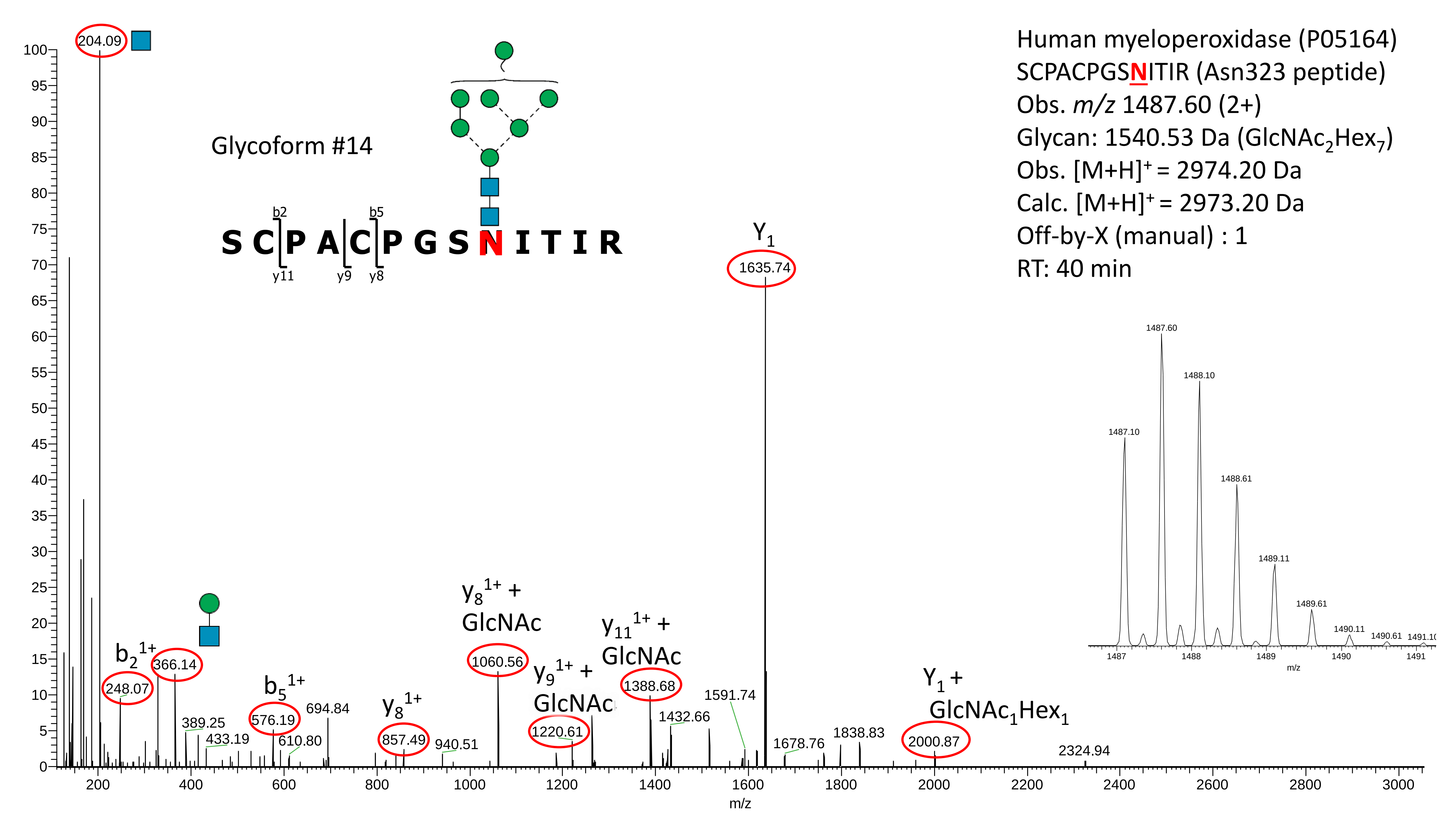

Human myeloperoxidase (P05164)
SCPACPGSNITIR (Asn323 peptide)
Obs. m/z 1487.60 (2+)
Glycan: 1540.53 Da (GlcNAc2Hex7)
Obs. [M+H]+ = 2974.20 Da
Calc. [M+H]+ = 2973.20 Da
Off-by-X (manual) : 1
RT: 40 min
Glycoform #14
S C P A C P G S N I T I R
b2
b5
y11
y9
y8
204.09
Y1
1635.74
b2 1+
248.07
366.14
389.25
433.19
b5 1+
576.19
610.80
694.84
y8 1+
857.49
940.51
y8 1+ + GlcNAc
1060.56
y9 1+ + GlcNAc
1220.61
y11 1+ + GlcNAc
1388.68
1432.66
1591.74
1678.76
1838.83
Y1 + GlcNAc1Hex1
2000.87
2324.94
1487.10
1487.60
1488.10
1488.61
1489.11
1489.61
1490.11
1490.61
m/z

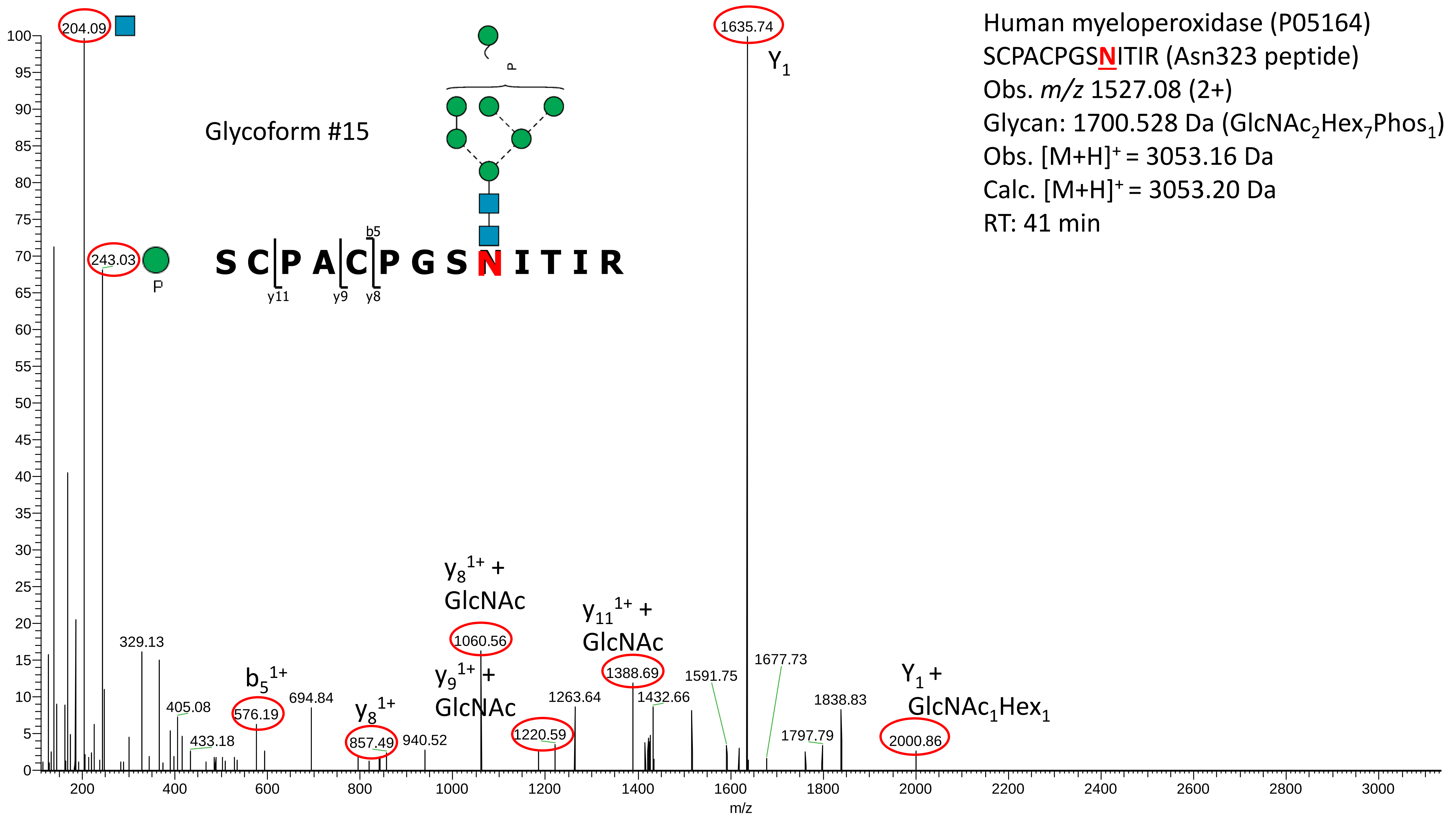

Human myeloperoxidase (P05164)
SCPACPGSNITIR (Asn323 peptide)
Obs. $m/z$ 1527.08 (2+)
Glycan: 1700.528 Da ($GlcNAc_2Hex_7Phos_1$)
Obs. $[M+H]^+$ = 3053.16 Da
Calc. $[M+H]^+$ = 3053.20 Da
RT: 41 min
Glycoform #15
S C P A C P G S N I T I R
b5
y11
y9
y8
P
204.09
243.03
1635.74
$Y_1$
329.13
405.08
433.18
$b_5^{1+}$
576.19
694.84
$y_8^{1+}$
857.49
940.52
$y_9^{1+}$ + GlcNAc
$y_8^{1+}$ + GlcNAc
1060.56
1220.59
1263.64
$y_{11}^{1+}$ + GlcNAc
1388.69
1432.66
1591.75
1677.73
1797.79
1838.83
$Y_1$ + $GlcNAc_1Hex_1$
2000.86
m/z

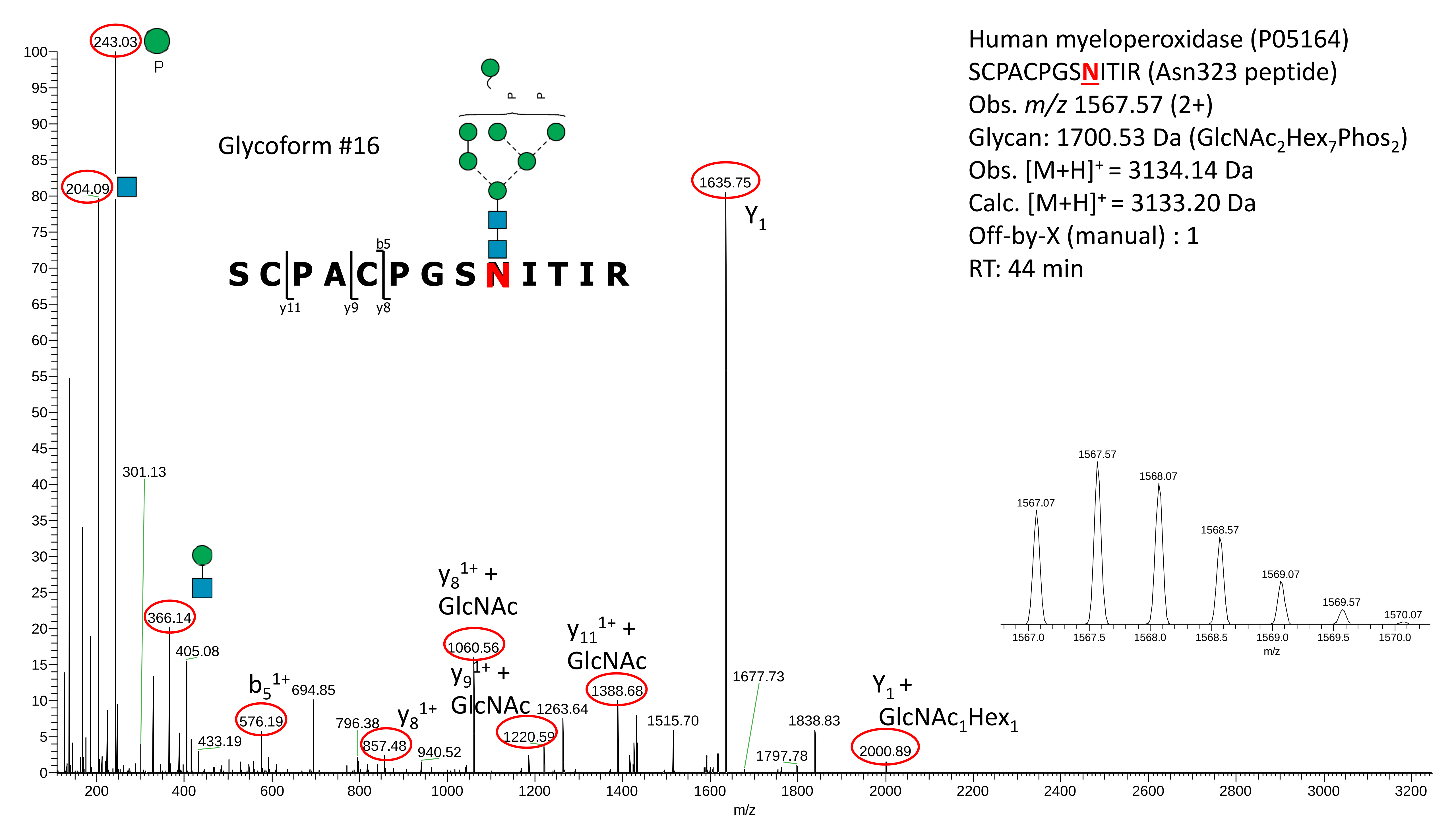

Human myeloperoxidase (P05164)
SCPACPGSNITIR (Asn323 peptide)
Obs. m/z 1567.57 (2+)
Glycan: 1700.53 Da (GlcNAc2Hex7Phos2)
Obs. [M+H]+ = 3134.14 Da
Calc. [M+H]+ = 3133.20 Da
Off-by-X (manual) : 1
RT: 44 min
Glycoform #16
S C P A C P G S N I T I R
b5
y11
y9
y8
P
243.03
204.09
301.13
366.14
405.08
433.19
b5 1+
576.19
694.85
796.38
y8 1+
857.48
940.52
y8 1+ + GlcNAc
1060.56
y9 1+ + GlcNAc
1220.59
1263.64
y11 1+ + GlcNAc
1388.68
1515.70
1635.75
Y1
1677.73
1797.78
1838.83
Y1 + GlcNAc1Hex1
2000.89
1567.07
1567.57
1568.07
1568.57
1569.07
1569.57
1570.07
m/z

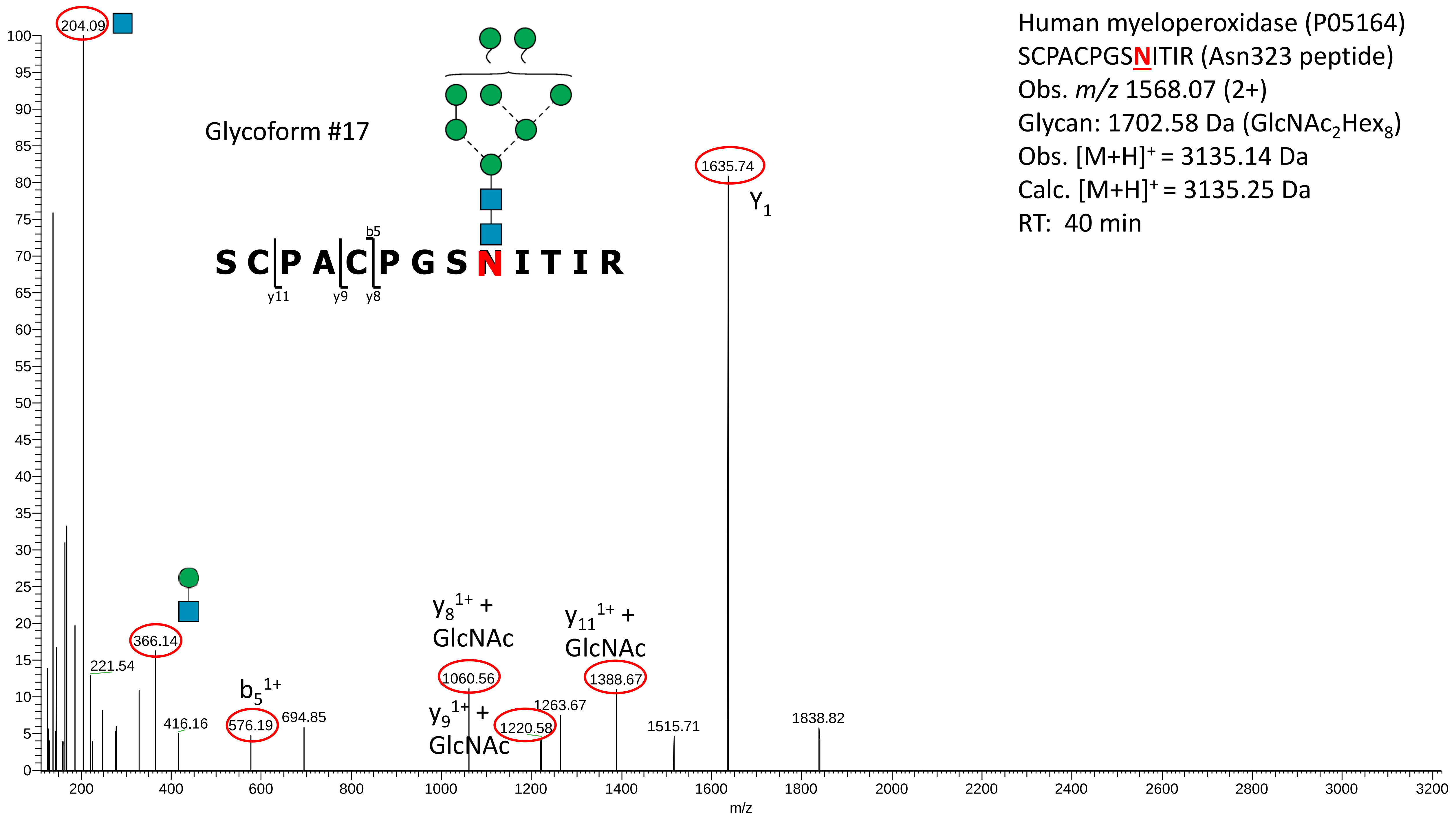
Human myeloperoxidase (P05164)
SCPACPGSNITIR (Asn323 peptide)
Obs. m/z 1568.07 (2+)
Glycan: 1702.58 Da (GlcNAc2Hex8)
Obs. [M+H]+ = 3135.14 Da
Calc. [M+H]+ = 3135.25 Da
RT: 40 min
Glycoform #17
S C P A C P G S N I T I R
b5
y11
y9
y8
204.09
1635.74
Y1
366.14
221.54
416.16
b5 1+
576.19
694.85
y8 1+ + GlcNAc
1060.56
y9 1+ + GlcNAc
1220.58
1263.67
y11 1+ + GlcNAc
1388.67
1515.71
1838.82
m/z

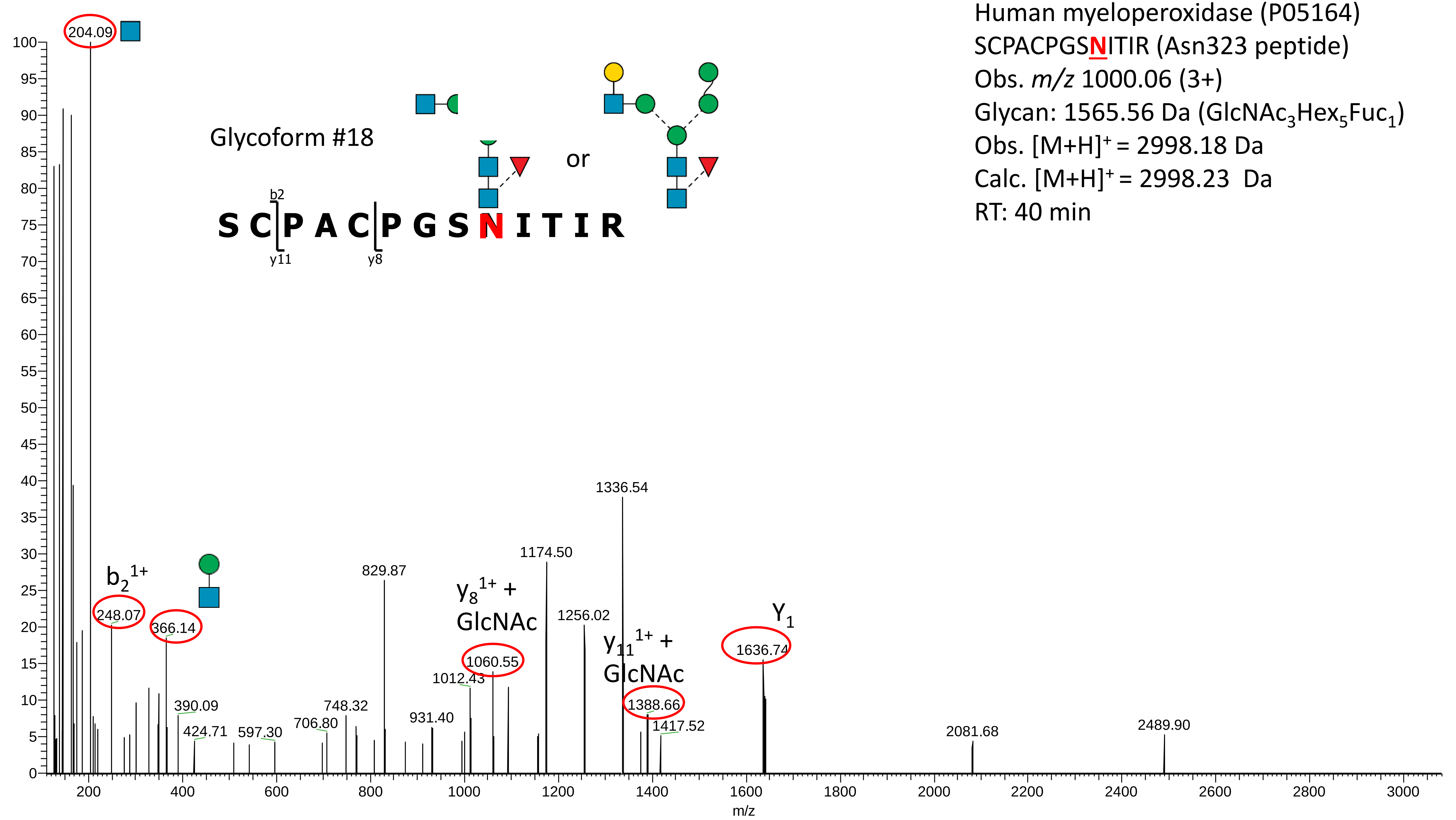

Human myeloperoxidase (P05164)
SCPACPGSNITIR (Asn323 peptide)
Obs. m/z 1000.06 (3+)
Glycan: 1565.56 Da (GlcNAc3Hex5Fuc1)
Obs. [M+H]+ = 2998.18 Da
Calc. [M+H]+ = 2998.23 Da
RT: 40 min
Glycoform #18
or
b2
S C P A C P G S N I T I R
y11
y8
204.09
b2 1+
248.07
366.14
390.09
424.71
597.30
706.80
748.32
829.87
931.40
1012.43
y8 1+ + GlcNAc
1060.55
1174.50
1256.02
1336.54
y11 1+ + GlcNAc
1388.66
1417.52
Y1
1636.74
2081.68
2489.90
m/z

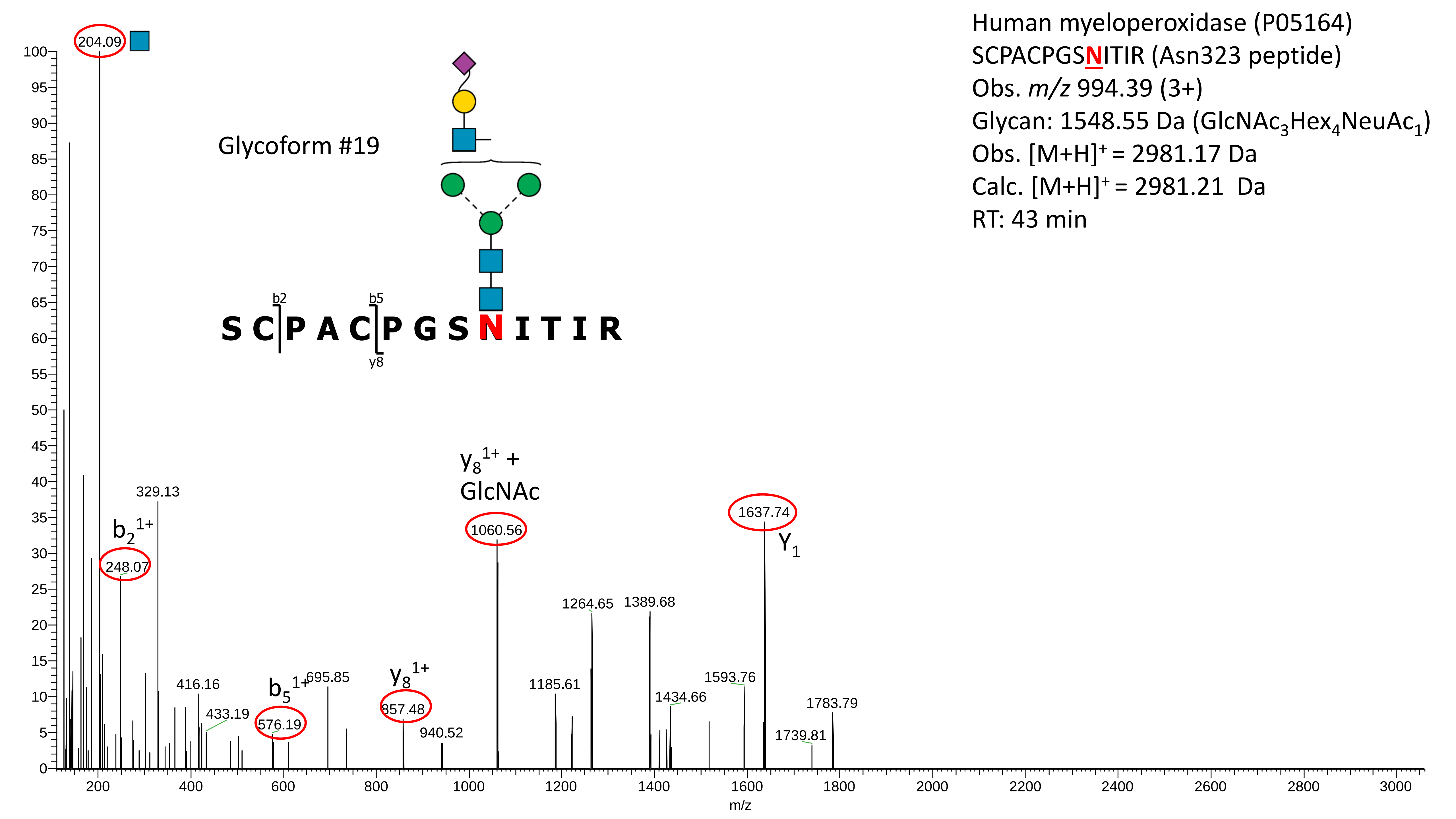

Human myeloperoxidase (P05164)
SCPACPGSNITIR (Asn323 peptide)
Obs. m/z 994.39 (3+)
Glycan: 1548.55 Da (GlcNAc3Hex4NeuAc1)
Obs. [M+H]+ = 2981.17 Da
Calc. [M+H]+ = 2981.21 Da
RT: 43 min
Glycoform #19
S C P A C P G S N I T I R
b2
b5
y8
204.09
329.13
b2 1+
248.07
416.16
433.19
b5 1+
576.19
695.85
y8 1+
857.48
940.52
y8 1+ + GlcNAc
1060.56
1185.61
1264.65
1389.68
1434.66
1593.76
1637.74
Y1
1739.81
1783.79
m/z

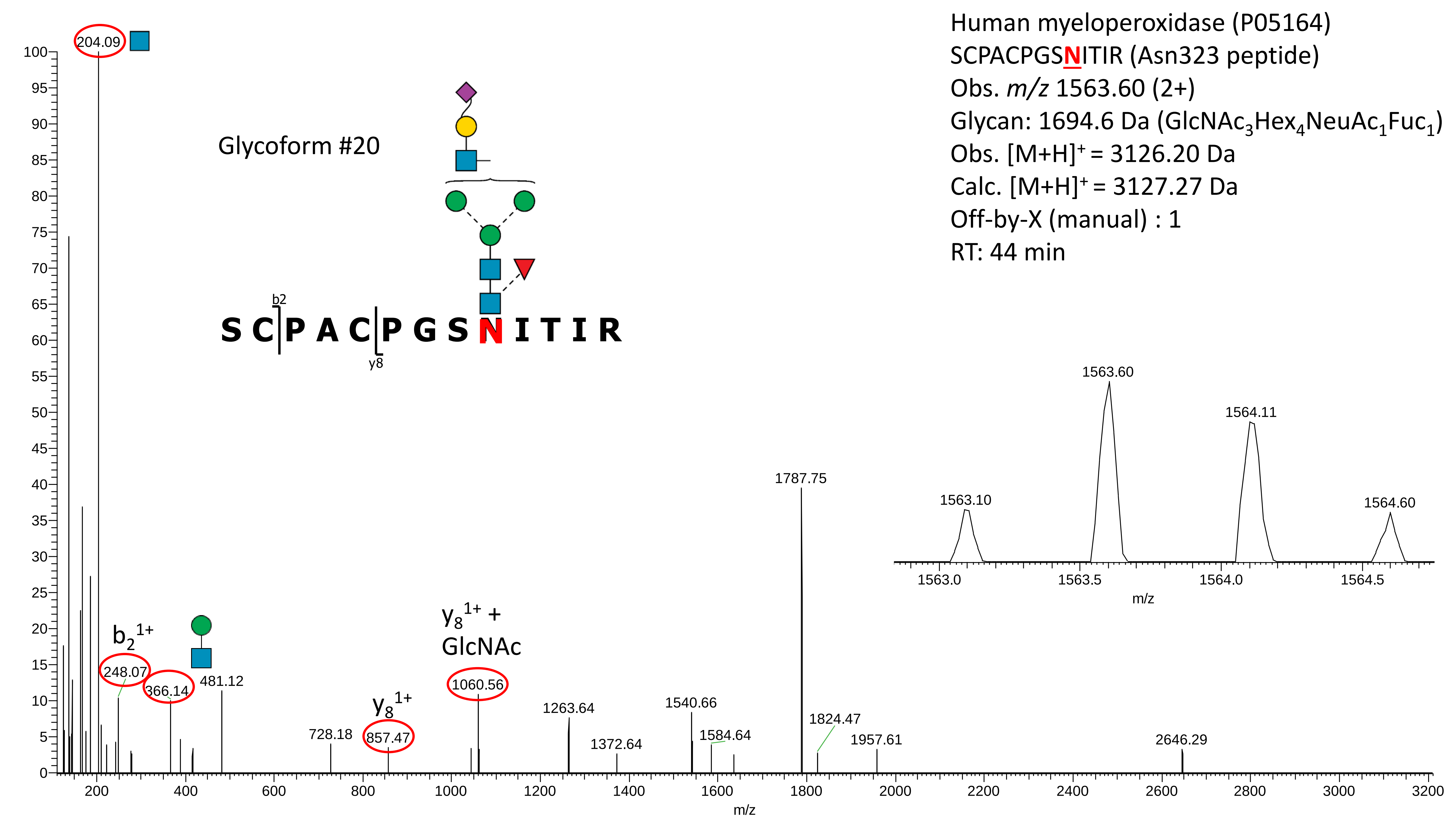

Human myeloperoxidase (P05164)
SCPACPGSNITIR (Asn323 peptide)
Obs. m/z 1563.60 (2+)
Glycan: 1694.6 Da (GlcNAc3Hex4NeuAc1Fuc1)
Obs. [M+H]+ = 3126.20 Da
Calc. [M+H]+ = 3127.27 Da
Off-by-X (manual) : 1
RT: 44 min
Glycoform #20
S C P A C P G S N I T I R
b2
y8
b2 1+
y8 1+
y8 1+ + GlcNAc
204.09
248.07
366.14
481.12
728.18
857.47
1060.56
1263.64
1372.64
1540.66
1584.64
1787.75
1824.47
1957.61
2646.29
1563.10
1563.60
1564.11
1564.60
m/z

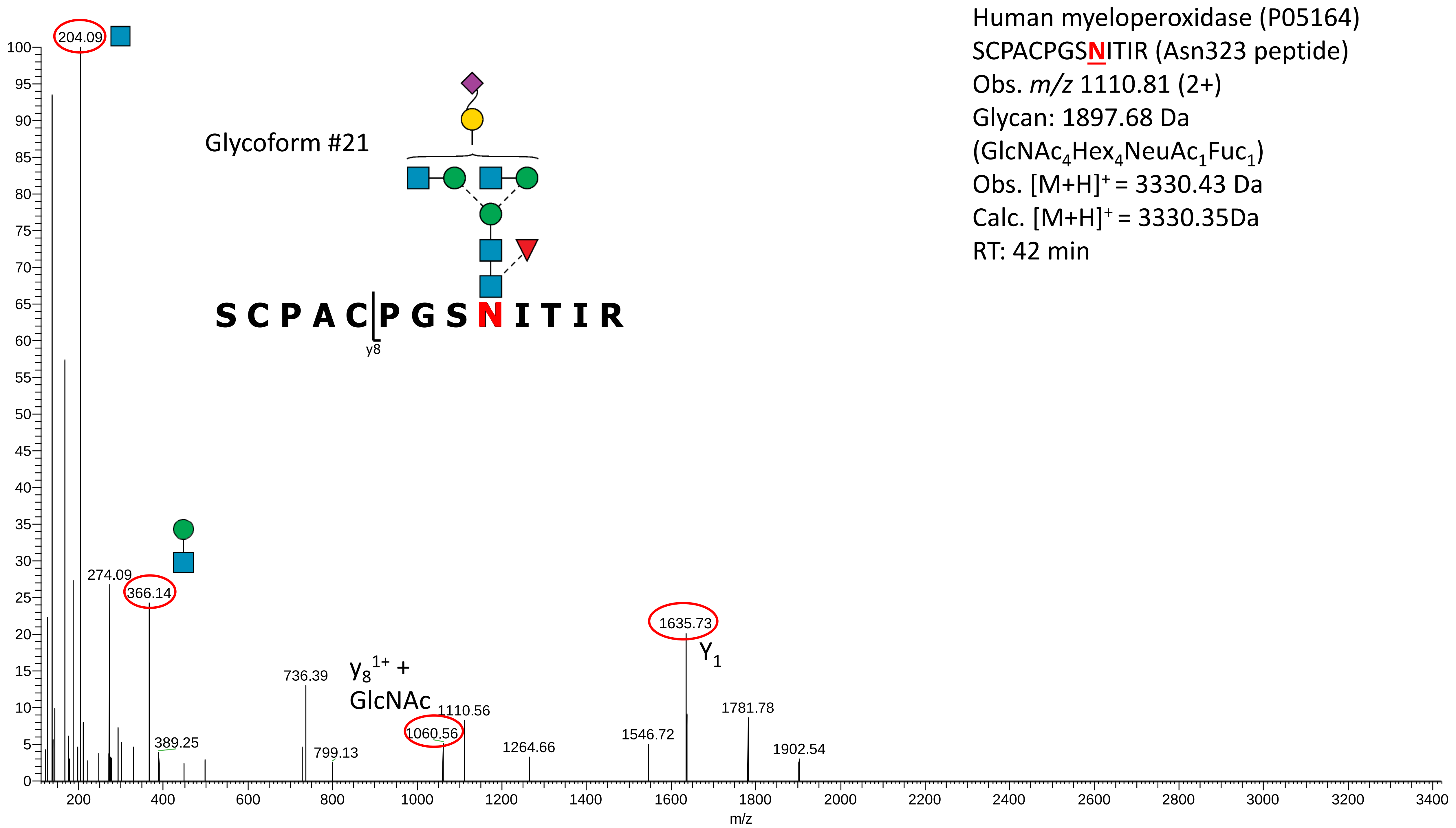

Human myeloperoxidase (P05164)
SCPACPGSNITIR (Asn323 peptide)
Obs. m/z 1110.81 (2+)
Glycan: 1897.68 Da
(GlcNAc4Hex4NeuAc1Fuc1)
Obs. [M+H]+ = 3330.43 Da
Calc. [M+H]+ = 3330.35Da
RT: 42 min
Glycoform #21
S C P A C P G S N I T I R
y8
204.09
274.09
366.14
389.25
736.39
799.13
y8 1+ + GlcNAc
1060.56
1110.56
1264.66
1546.72
1635.73
Y1
1781.78
1902.54
m/z

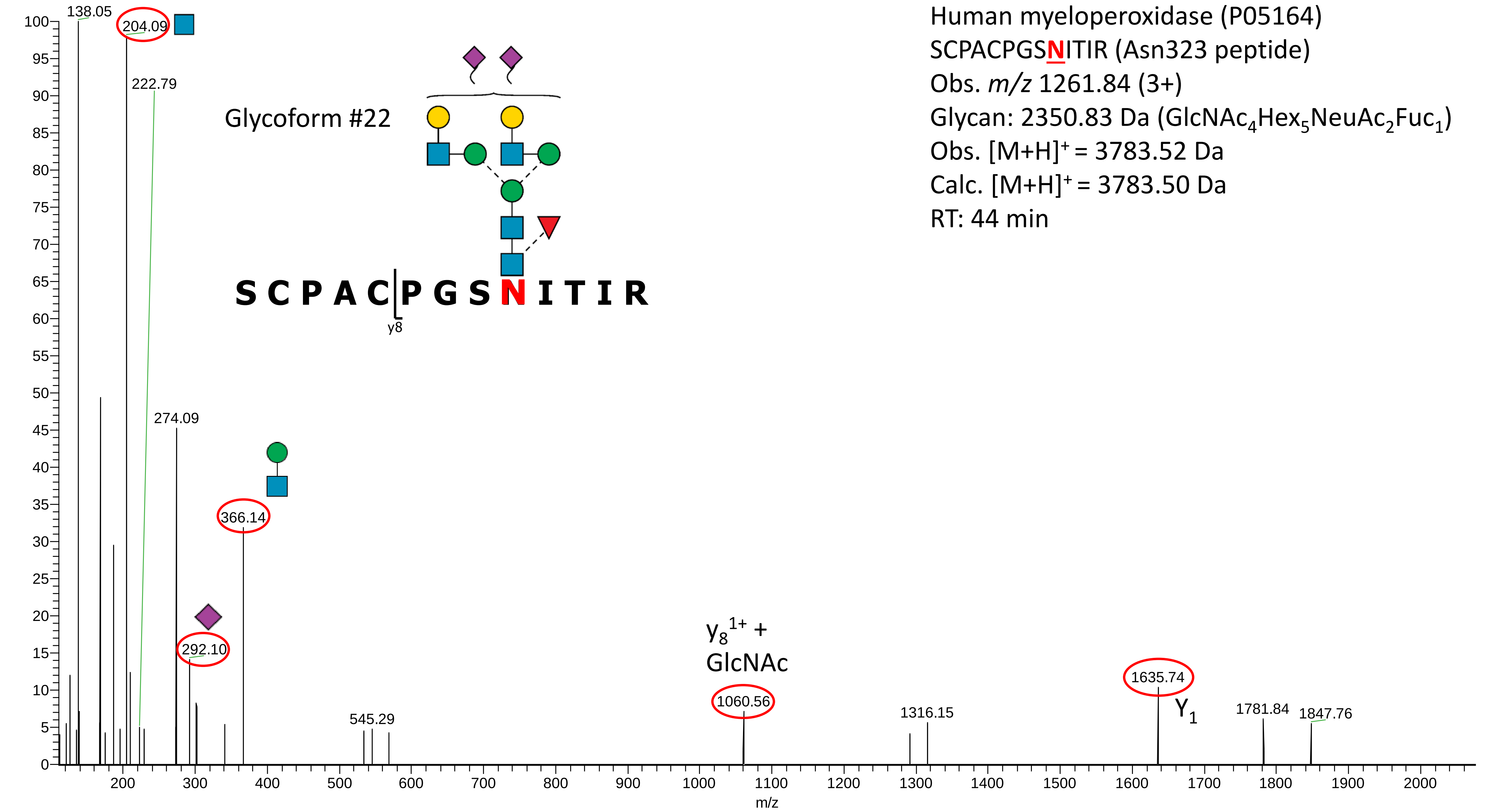
Human myeloperoxidase (P05164)
SCPACPGSNITIR (Asn323 peptide)
Obs. m/z 1261.84 (3+)
Glycan: 2350.83 Da (GlcNAc4Hex5NeuAc2Fuc1)
Obs. [M+H]+ = 3783.52 Da
Calc. [M+H]+ = 3783.50 Da
RT: 44 min
Glycoform #22
S C P A C P G S N I T I R
y8
138.05
204.09
222.79
274.09
366.14
292.10
545.29
y8 1+ + GlcNAc
1060.56
1316.15
1635.74
Y1
1781.84
1847.76
m/z

**Supplementary Data S3B**

Manually annotated reversed phase-LC-ESI-HCD-MS/MS (+) spectra of all glycosylated and non-glycosylated peptides containing Asn355 identified from the analysis of the unenriched peptide mixtures of nMPO.

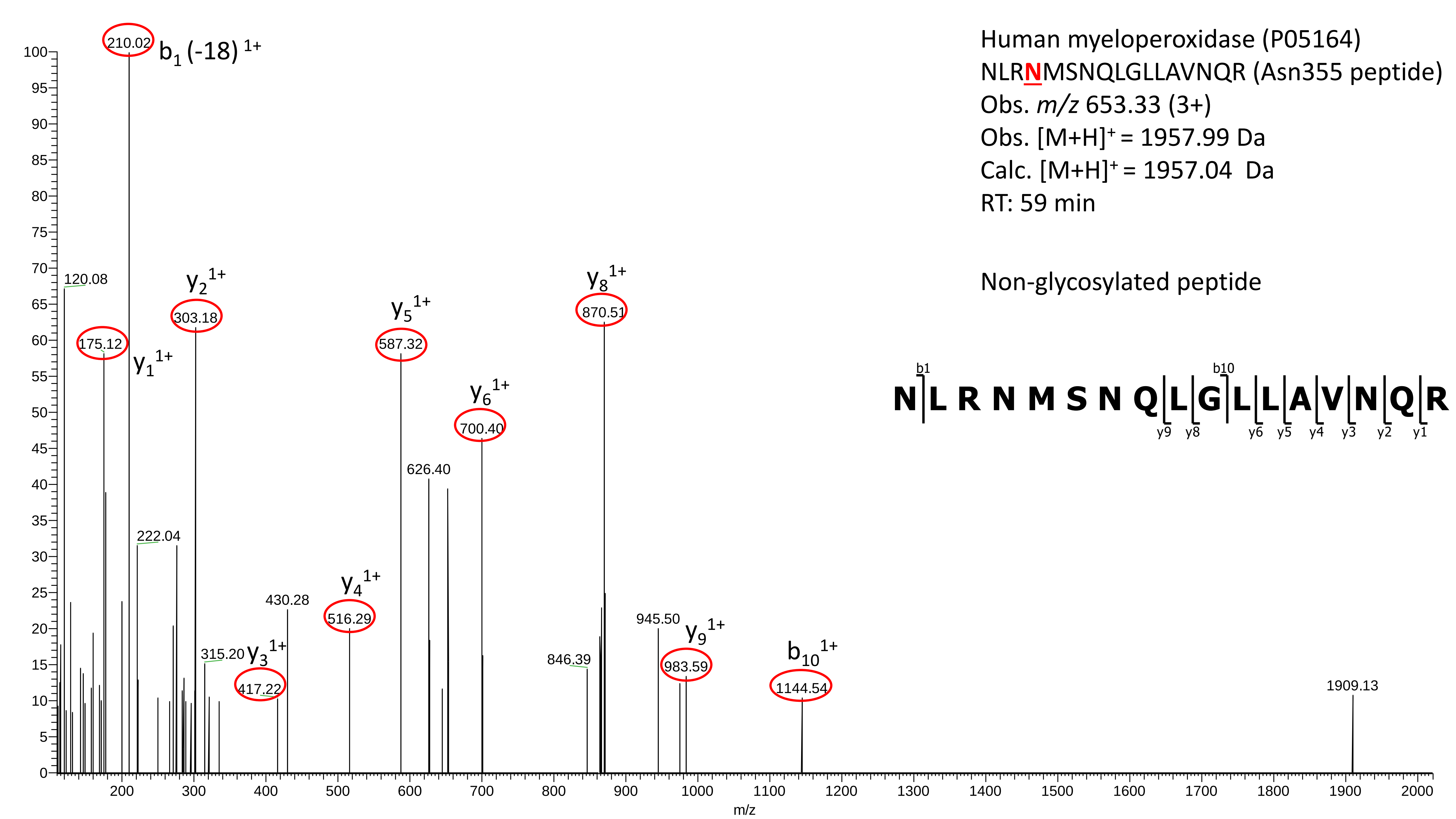
Human myeloperoxidase (P05164)
NLRNMSNQLGLLAVNQR (Asn355 peptide)
Obs. m/z 653.33 (3+)
Obs. [M+H]+ = 1957.99 Da
Calc. [M+H]+ = 1957.04 Da
RT: 59 min
Non-glycosylated peptide
N L R N M S N Q L G L L A V N Q R
b1 b10
y9 y8 y6 y5 y4 y3 y2 y1
210.02 b1 (-18) 1+
120.08
175.12 y1 1+
303.18 y2 1+
222.04
315.20
417.22 y3 1+
430.28
516.29 y4 1+
587.32 y5 1+
626.40
700.40 y6 1+
846.39
870.51 y8 1+
945.50
983.59 y9 1+
1144.54 b10 1+
1909.13
m/z

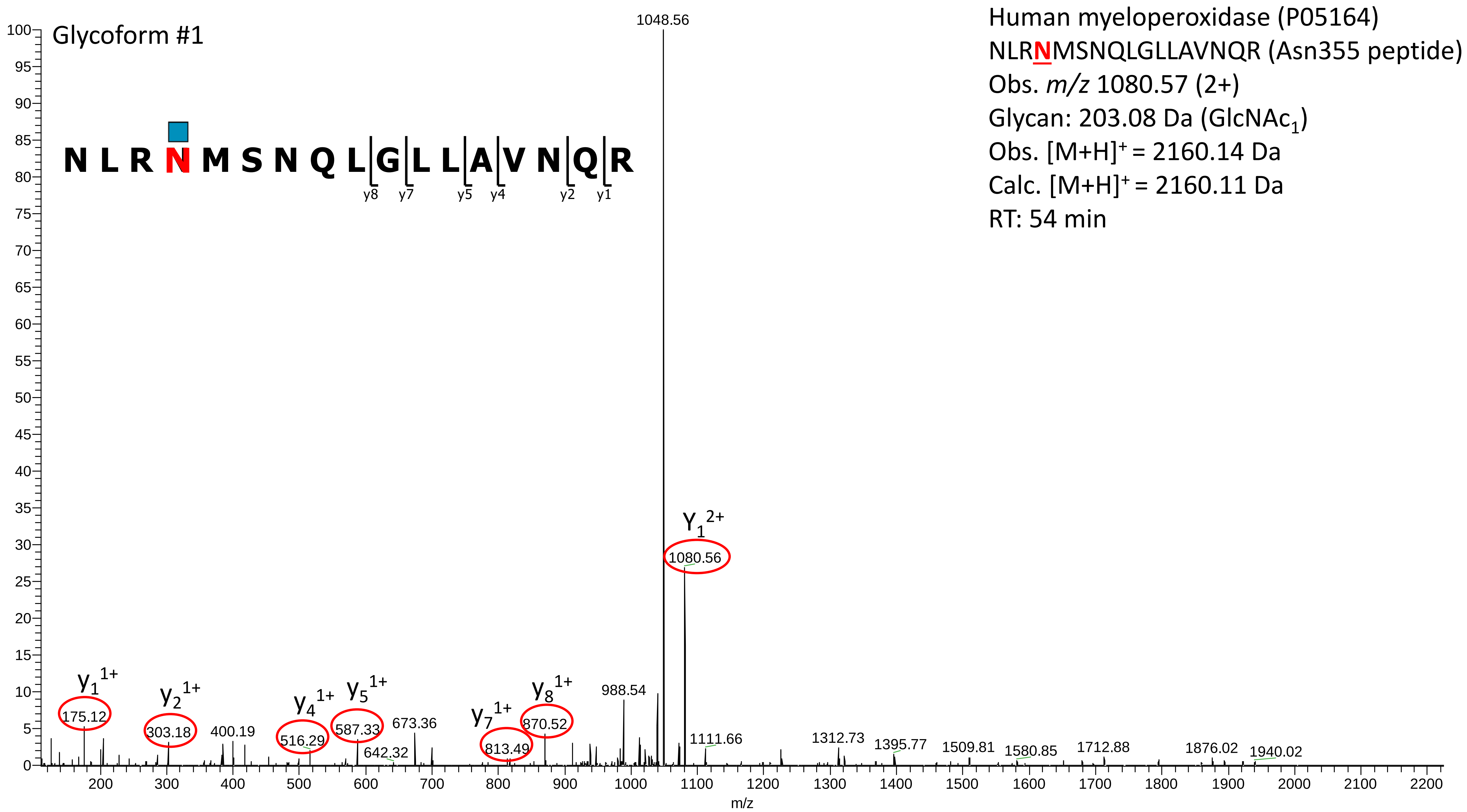
Glycoform #1
Human myeloperoxidase (P05164)
NLRNMSNQLGLLAVNQR (Asn355 peptide)
Obs. m/z 1080.57 (2+)
Glycan: 203.08 Da (GlcNAc1)
Obs. [M+H]+ = 2160.14 Da
Calc. [M+H]+ = 2160.11 Da
RT: 54 min
N L R N M S N Q L G L L A V N Q R
y8 y7 y5 y4 y2 y1
1048.56
Y1 2+
1080.56
y1 1+
175.12
y2 1+
303.18
400.19
y4 1+
516.29
y5 1+
587.33
642.32
673.36
y7 1+
813.49
y8 1+
870.52
988.54
1111.66
1312.73
1395.77
1509.81
1580.85
1712.88
1876.02
1940.02
m/z

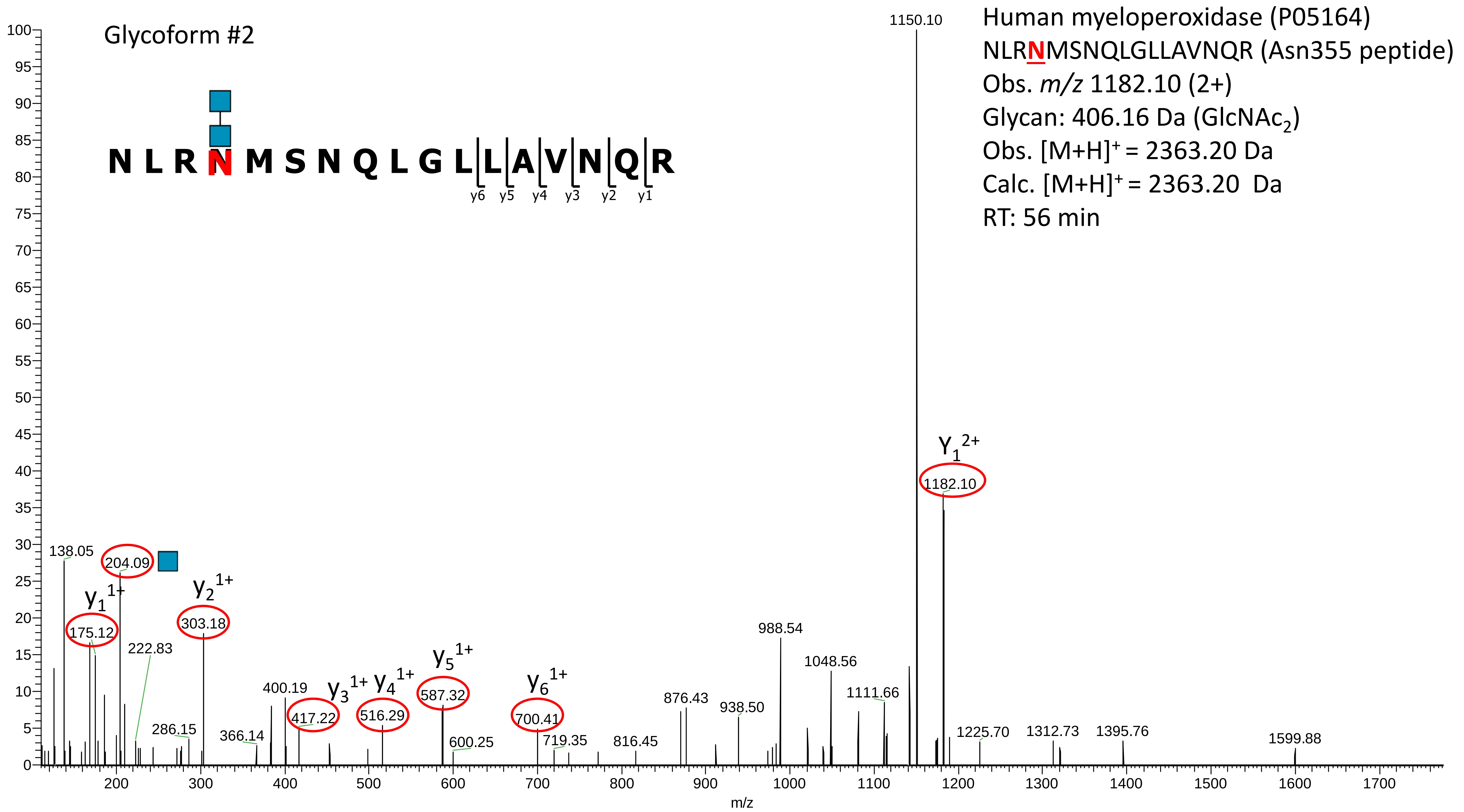
Glycoform #2
Human myeloperoxidase (P05164)
NLRNMSNQLGLLAVNQR (Asn355 peptide)
Obs. m/z 1182.10 (2+)
Glycan: 406.16 Da (GlcNAc2)
Obs. [M+H]+ = 2363.20 Da
Calc. [M+H]+ = 2363.20 Da
RT: 56 min
N L R N M S N Q L G L L A V N Q R
y6 y5 y4 y3 y2 y1
1150.10
Y1 2+
1182.10
138.05
204.09
y1 1+
175.12
y2 1+
303.18
222.83
286.15
366.14
400.19
y3 1+
417.22
y4 1+
516.29
y5 1+
587.32
600.25
y6 1+
700.41
719.35
816.45
876.43
938.50
988.54
1048.56
1111.66
1225.70
1312.73
1395.76
1599.88
m/z

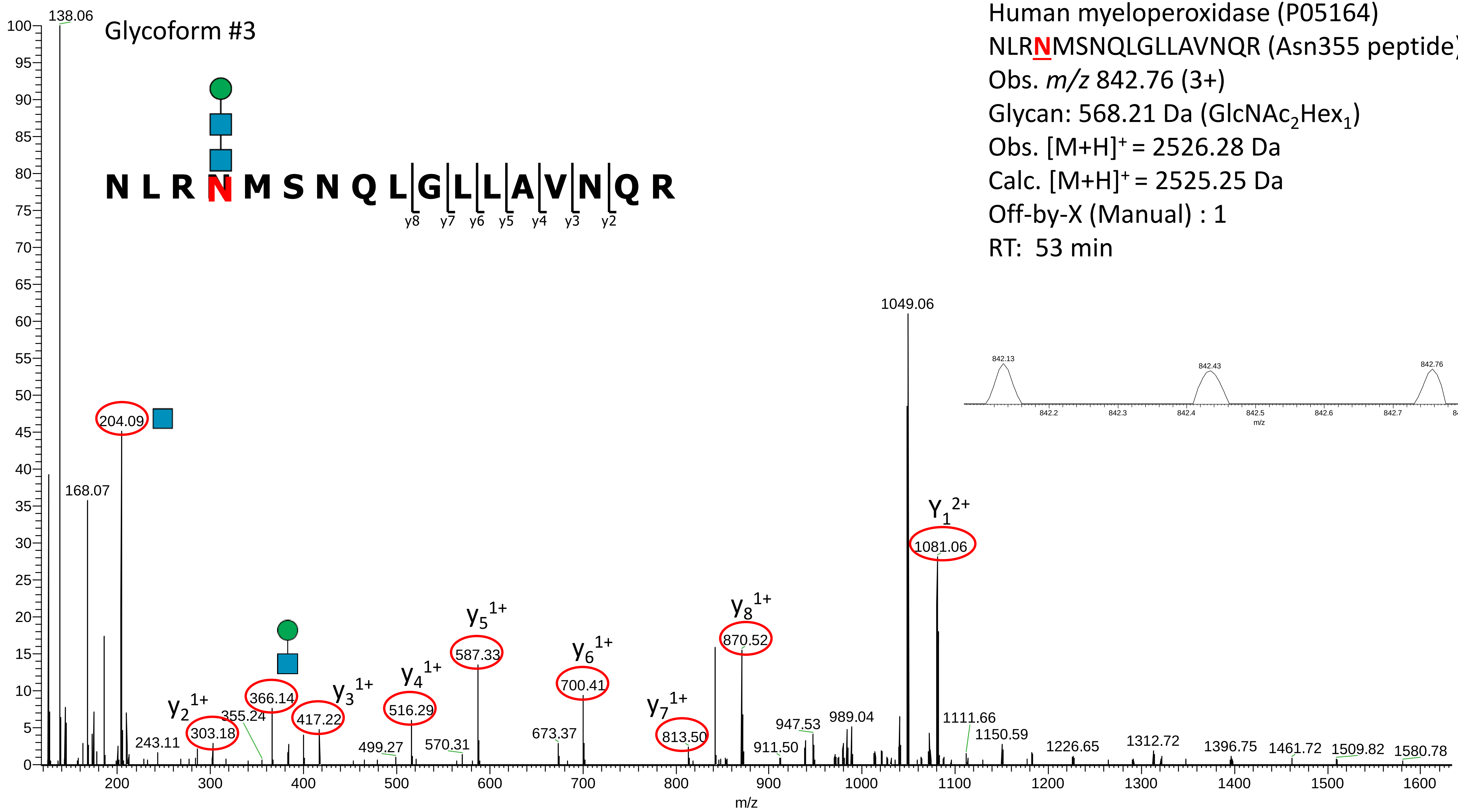


Human myeloperoxidase (P05164)

NLR**N**MSNQLGLLAVNQR (Asn355 peptide)

Obs. *m/z* 842.76 (3+)

Glycan: 568.21 Da ($GlcNAc_2Hex_1$)

Obs. $[M+H]^+$ = 2526.28 Da

Calc. $[M+H]^+$ = 2525.25 Da

Off-by-X (Manual) : 1

RT: 53 min

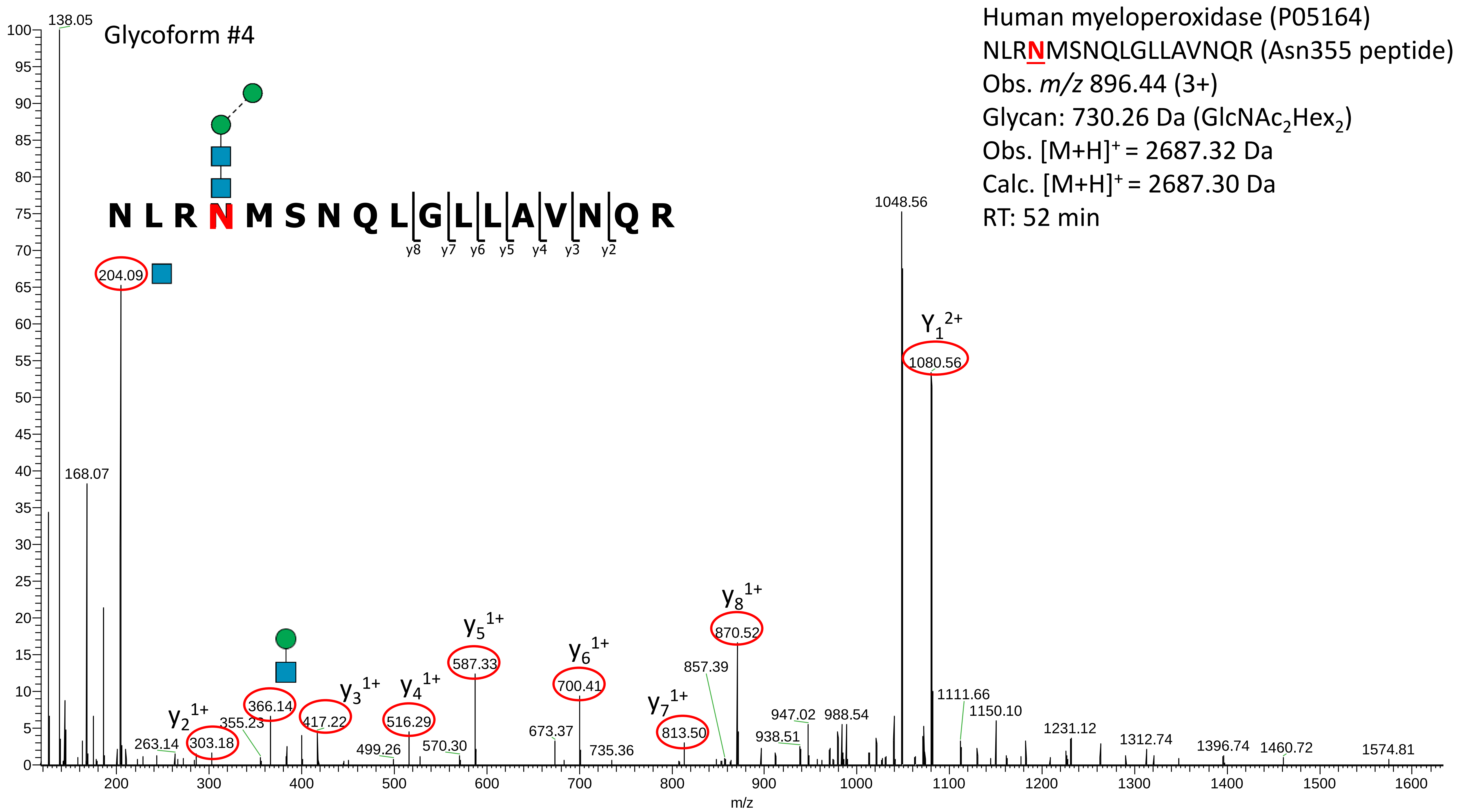
Glycoform #4
Human myeloperoxidase (P05164)
NLRNMSNQLGLLAVNQR (Asn355 peptide)
Obs. m/z 896.44 (3+)
Glycan: 730.26 Da (GlcNAc2Hex2)
Obs. [M+H]+ = 2687.32 Da
Calc. [M+H]+ = 2687.30 Da
RT: 52 min
N L R N M S N Q L G L L A V N Q R
y8 y7 y6 y5 y4 y3 y2
Y1 2+
y2 1+
y3 1+
y4 1+
y5 1+
y6 1+
y7 1+
y8 1+
138.05
168.07
204.09
263.14
303.18
355.23
366.14
417.22
499.26
516.29
570.30
587.33
673.37
700.41
735.36
813.50
857.39
870.52
938.51
947.02
988.54
1048.56
1080.56
1111.66
1150.10
1231.12
1312.74
1396.74
1460.72
1574.81
m/z

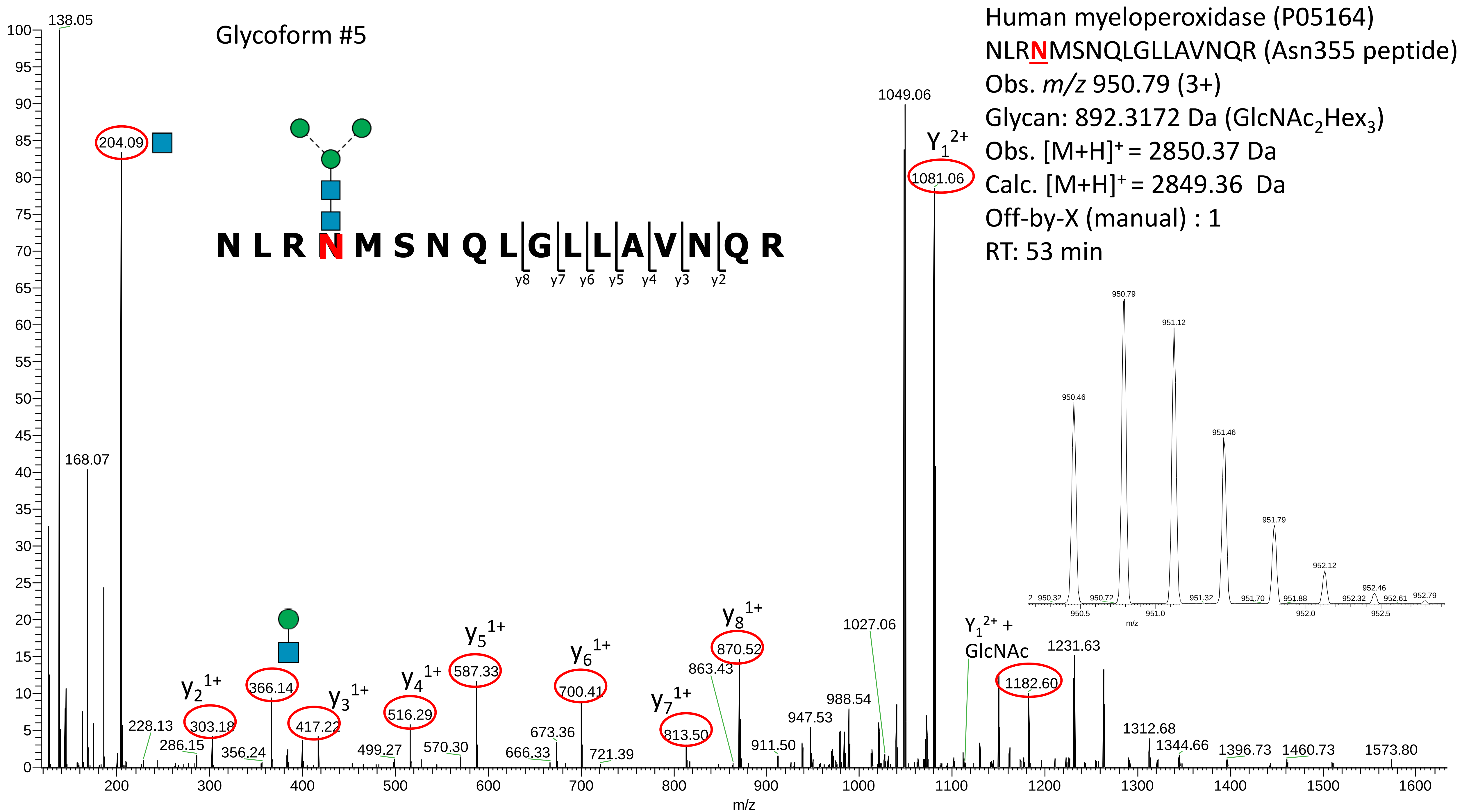

Glycoform #5
Human myeloperoxidase (P05164)
NLRNMSNQLGLLAVNQR (Asn355 peptide)
Obs. m/z 950.79 (3+)
Glycan: 892.3172 Da (GlcNAc2Hex3)
Obs. [M+H]+ = 2850.37 Da
Calc. [M+H]+ = 2849.36 Da
Off-by-X (manual) : 1
RT: 53 min
N L R N M S N Q L G L L A V N Q R
y8 y7 y6 y5 y4 y3 y2
138.05
168.07
204.09
228.13
286.15
y2 1+ 303.18
356.24
366.14
417.22
y3 1+
499.27
y4 1+ 516.29
570.30
y5 1+ 587.33
666.33
673.36
y6 1+ 700.41
721.39
y7 1+ 813.50
863.43
y8 1+ 870.52
911.50
947.53
988.54
1027.06
1049.06
Y1 2+ 1081.06
Y1 2+ + GlcNAc
1182.60
1231.63
1312.68
1344.66
1396.73
1460.73
1573.80
m/z
950.32
950.46
950.72
950.79
951.12
951.32
951.46
951.70
951.79
951.88
952.12
952.32
952.46
952.61
952.79
950.5
951.0
952.0
952.5

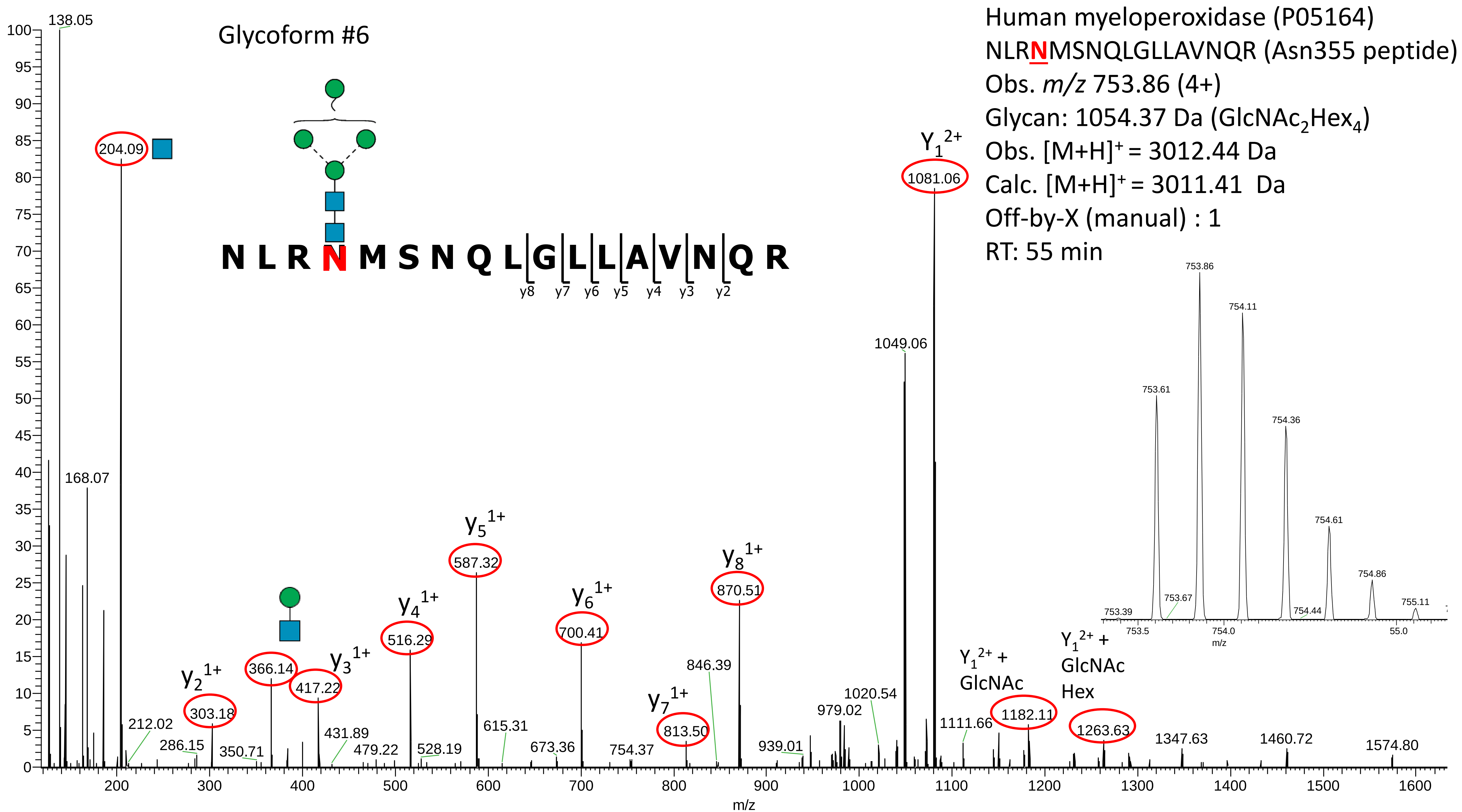

Glycoform #6
Human myeloperoxidase (P05164)
NLRNMSNQLGLLAVNQR (Asn355 peptide)
Obs. m/z 753.86 (4+)
Glycan: 1054.37 Da (GlcNAc2Hex4)
Obs. [M+H]+ = 3012.44 Da
Calc. [M+H]+ = 3011.41 Da
Off-by-X (manual) : 1
RT: 55 min
N L R N M S N Q L G L L A V N Q R
y8 y7 y6 y5 y4 y3 y2
138.05
204.09
168.07
212.02
286.15
y2 1+ 303.18
350.71
366.14
y3 1+ 417.22
431.89
479.22
y4 1+ 516.29
528.19
y5 1+ 587.32
615.31
673.36
y6 1+ 700.41
754.37
y7 1+ 813.50
846.39
y8 1+ 870.51
939.01
979.02
1020.54
1049.06
Y1 2+ 1081.06
1111.66
Y1 2+ + GlcNAc 1182.11
Y1 2+ + GlcNAc Hex 1263.63
1347.63
1460.72
1574.80
m/z
753.39
753.61
753.67
753.86
754.11
754.36
754.44
754.61
754.86
755.11
753.5
754.0
55.0

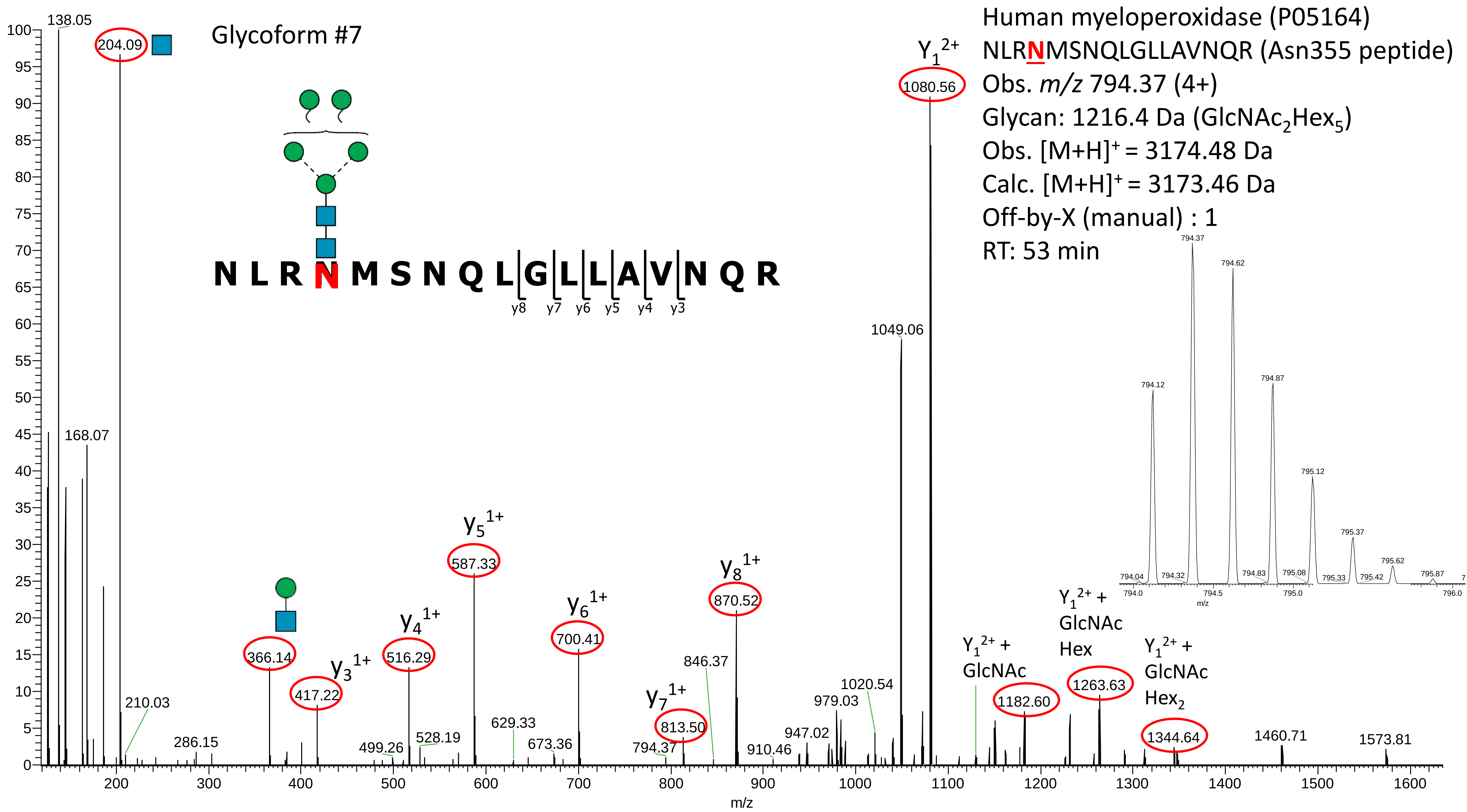
Glycoform #7
Human myeloperoxidase (P05164)
NLRNMSNQLGLLAVNQR (Asn355 peptide)
Obs. m/z 794.37 (4+)
Glycan: 1216.4 Da (GlcNAc2Hex5)
Obs. [M+H]+ = 3174.48 Da
Calc. [M+H]+ = 3173.46 Da
Off-by-X (manual) : 1
RT: 53 min
N L R N M S N Q L G L L A V N Q R
y8 y7 y6 y5 y4 y3
Y1 2+
1080.56
1049.06
138.05
204.09
168.07
210.03
286.15
366.14
417.22
y3 1+
y4 1+
516.29
y5 1+
587.33
499.26
528.19
629.33
673.36
y6 1+
700.41
y7 1+
813.50
794.37
846.37
y8 1+
870.52
910.46
947.02
979.03
1020.54
Y1 2+ + GlcNAc
1182.60
Y1 2+ + GlcNAc Hex
1263.63
Y1 2+ + GlcNAc Hex2
1344.64
1460.71
1573.81
m/z

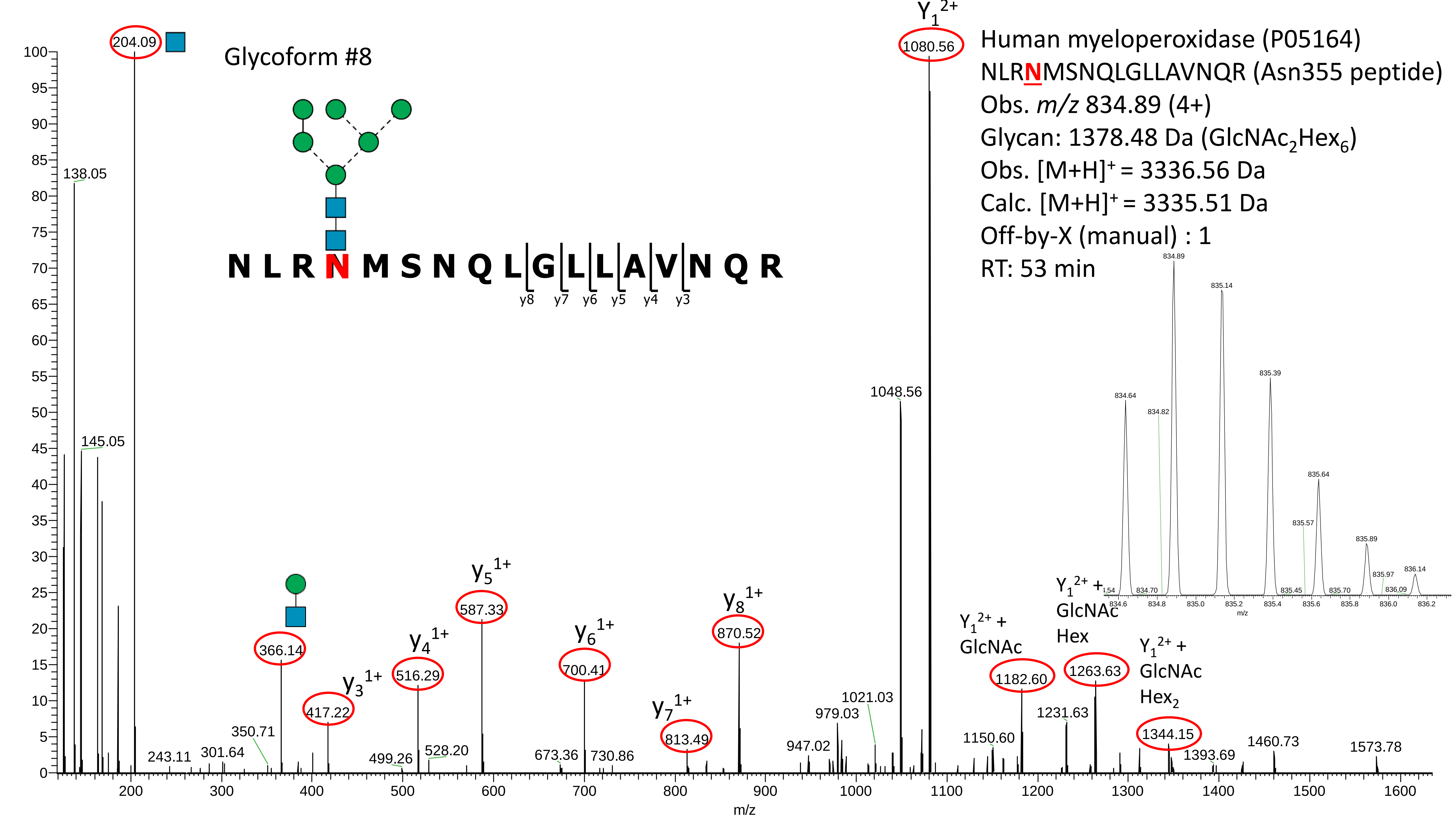

Glycoform #8
Human myeloperoxidase (P05164)
NLRNMSNQLGLLAVNQR (Asn355 peptide)
Obs. m/z 834.89 (4+)
Glycan: 1378.48 Da (GlcNAc2Hex6)
Obs. [M+H]+ = 3336.56 Da
Calc. [M+H]+ = 3335.51 Da
Off-by-X (manual) : 1
RT: 53 min
N L R N M S N Q L G L L A V N Q R
y8 y7 y6 y5 y4 y3
Y1 2+
1080.56
204.09
138.05
145.05
1048.56
366.14
417.22
y3 1+
516.29
y4 1+
587.33
y5 1+
700.41
y6 1+
813.49
y7 1+
870.52
y8 1+
243.11
301.64
350.71
499.26
528.20
673.36
730.86
947.02
979.03
1021.03
1150.60
Y1 2+ + GlcNAc
1182.60
1231.63
Y1 2+ + GlcNAc Hex
1263.63
Y1 2+ + GlcNAc Hex2
1344.15
1393.69
1460.73
1573.78
m/z
834.89
835.14
835.39
834.64
834.82
835.64
835.57
835.89
835.97
836.14
834.70
835.45
835.70
836.09
m/z

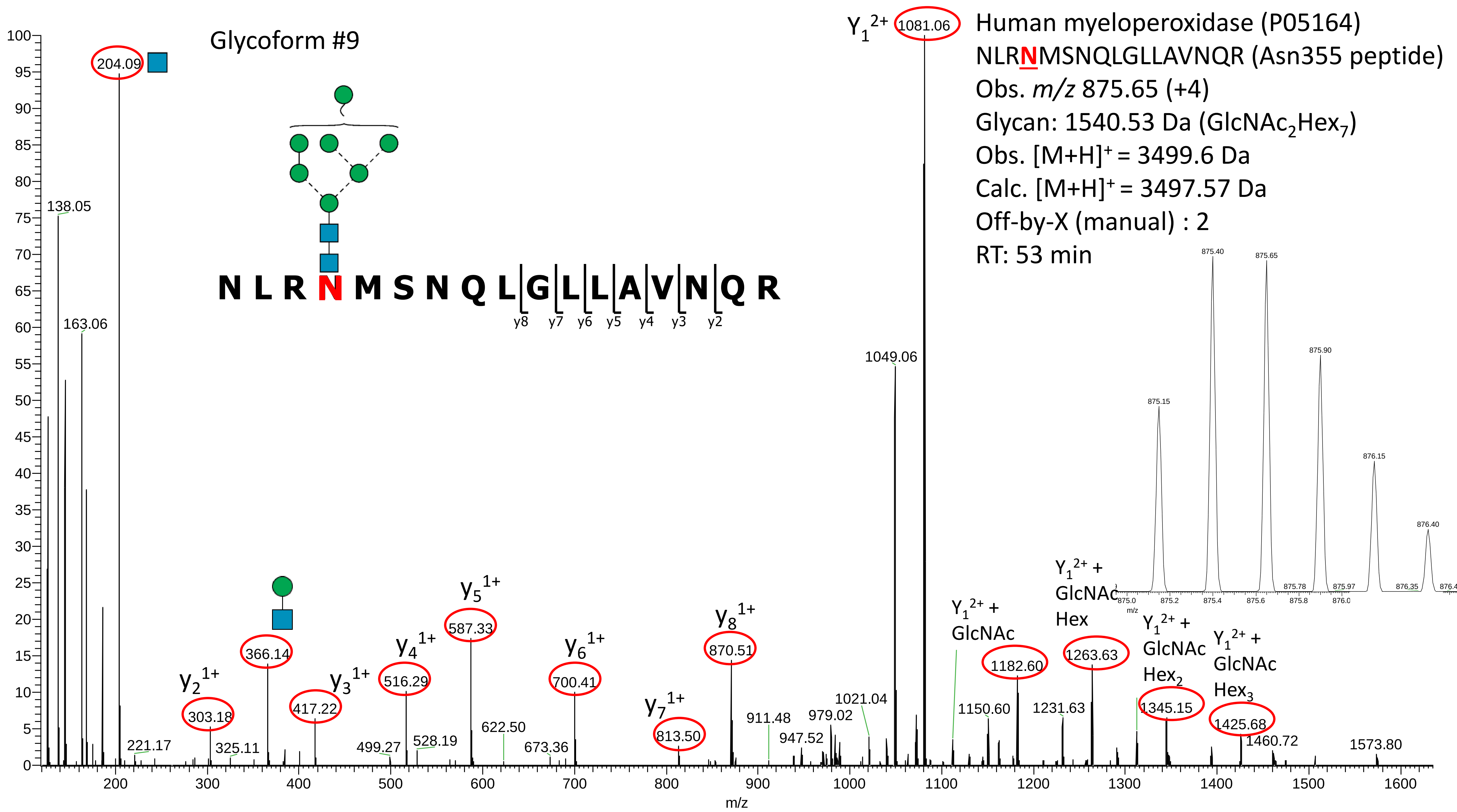

Glycoform #9
Human myeloperoxidase (P05164)
NLRNMSNQLGLLAVNQR (Asn355 peptide)
Obs. m/z 875.65 (+4)
Glycan: 1540.53 Da (GlcNAc2Hex7)
Obs. [M+H]+ = 3499.6 Da
Calc. [M+H]+ = 3497.57 Da
Off-by-X (manual) : 2
RT: 53 min
N L R N M S N Q L G L L A V N Q R
y8 y7 y6 y5 y4 y3 y2
Y1 2+ 1081.06
204.09
138.05
163.06
1049.06
y2 1+ 303.18
366.14
y3 1+ 417.22
y4 1+ 516.29
y5 1+ 587.33
y6 1+ 700.41
y7 1+ 813.50
y8 1+ 870.51
Y1 2+ + GlcNAc 1182.60
Y1 2+ + GlcNAc Hex 1263.63
Y1 2+ + GlcNAc Hex2 1345.15
Y1 2+ + GlcNAc Hex3 1425.68
221.17
325.11
499.27
528.19
622.50
673.36
911.48
947.52
979.02
1021.04
1150.60
1231.63
1460.72
1573.80
m/z
875.15
875.40
875.65
875.90
876.15
876.40

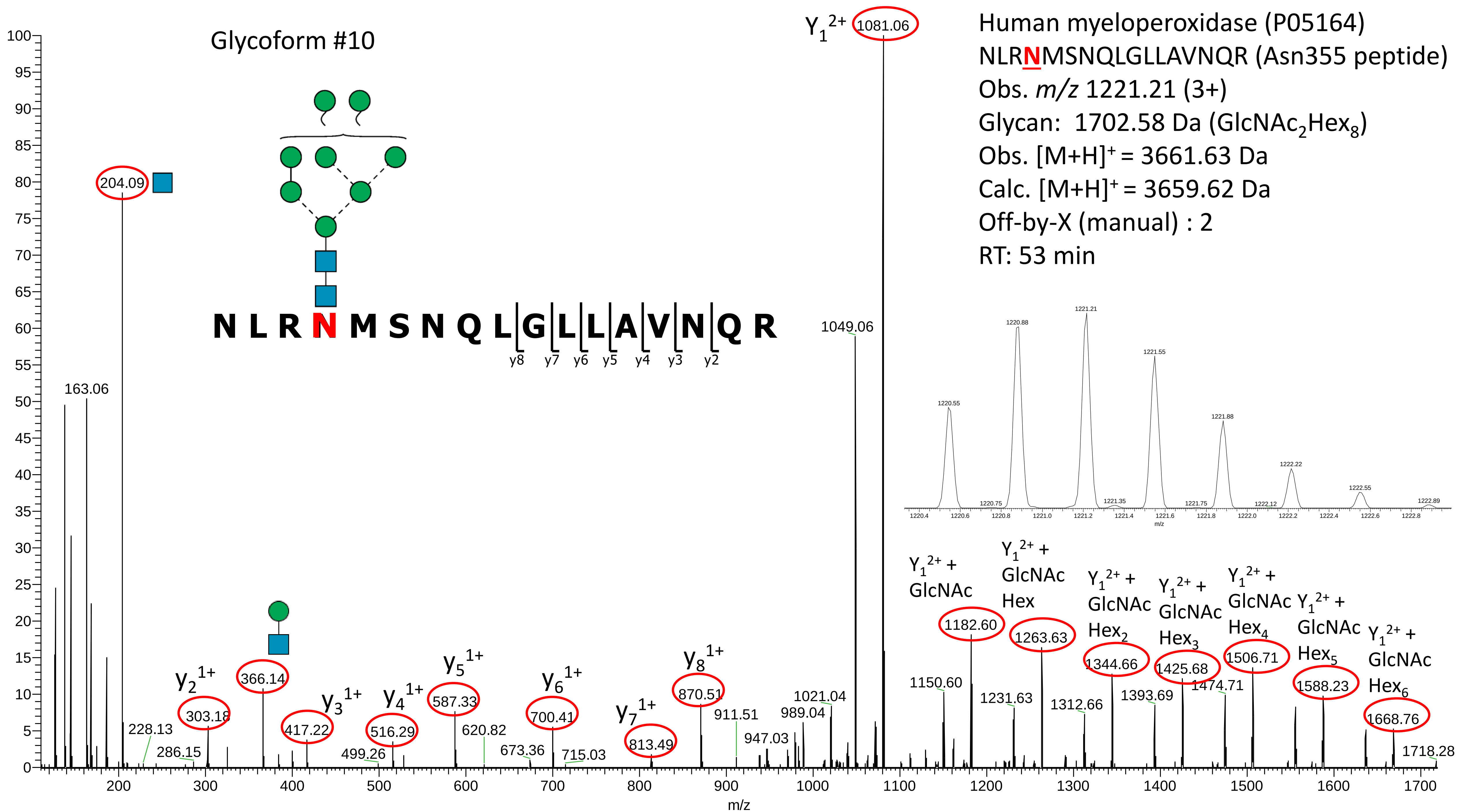

Glycoform #10
Human myeloperoxidase (P05164)
NLRNMSNQLGLLAVNQR (Asn355 peptide)
Obs. m/z 1221.21 (3+)
Glycan: 1702.58 Da (GlcNAc2Hex8)
Obs. [M+H]+ = 3661.63 Da
Calc. [M+H]+ = 3659.62 Da
Off-by-X (manual) : 2
RT: 53 min
N L R N M S N Q L G L L A V N Q R
y8 y7 y6 y5 y4 y3 y2
Y1 2+ 1081.06
1049.06
204.09
163.06
y2 1+ 303.18
366.14
y3 1+ 417.22
y4 1+ 516.29
y5 1+ 587.33
y6 1+ 700.41
y7 1+ 813.49
y8 1+ 870.51
228.13
286.15
499.26
620.82
673.36
715.03
911.51
947.03
989.04
1021.04
1150.60
Y1 2+ + GlcNAc 1182.60
Y1 2+ + GlcNAc Hex 1263.63
1231.63
Y1 2+ + GlcNAc Hex2 1344.66
1312.66
Y1 2+ + GlcNAc Hex3 1425.68
1393.69
Y1 2+ + GlcNAc Hex4 1506.71
1474.71
Y1 2+ + GlcNAc Hex5 1588.23
Y1 2+ + GlcNAc Hex6 1668.76
1718.28
1220.55
1220.75
1220.88
1221.21
1221.35
1221.55
1221.75
1221.88
1222.12
1222.22
1222.55
1222.89
m/z

**Supplementary Data S3C**

Manually annotated reversed phase-LC-ESI-HCD-MS/MS (+) spectra of all glycosylated and non-glycosylated peptides containing Asn391 identified from the analysis of unenriched peptide mixtures of nMPO.

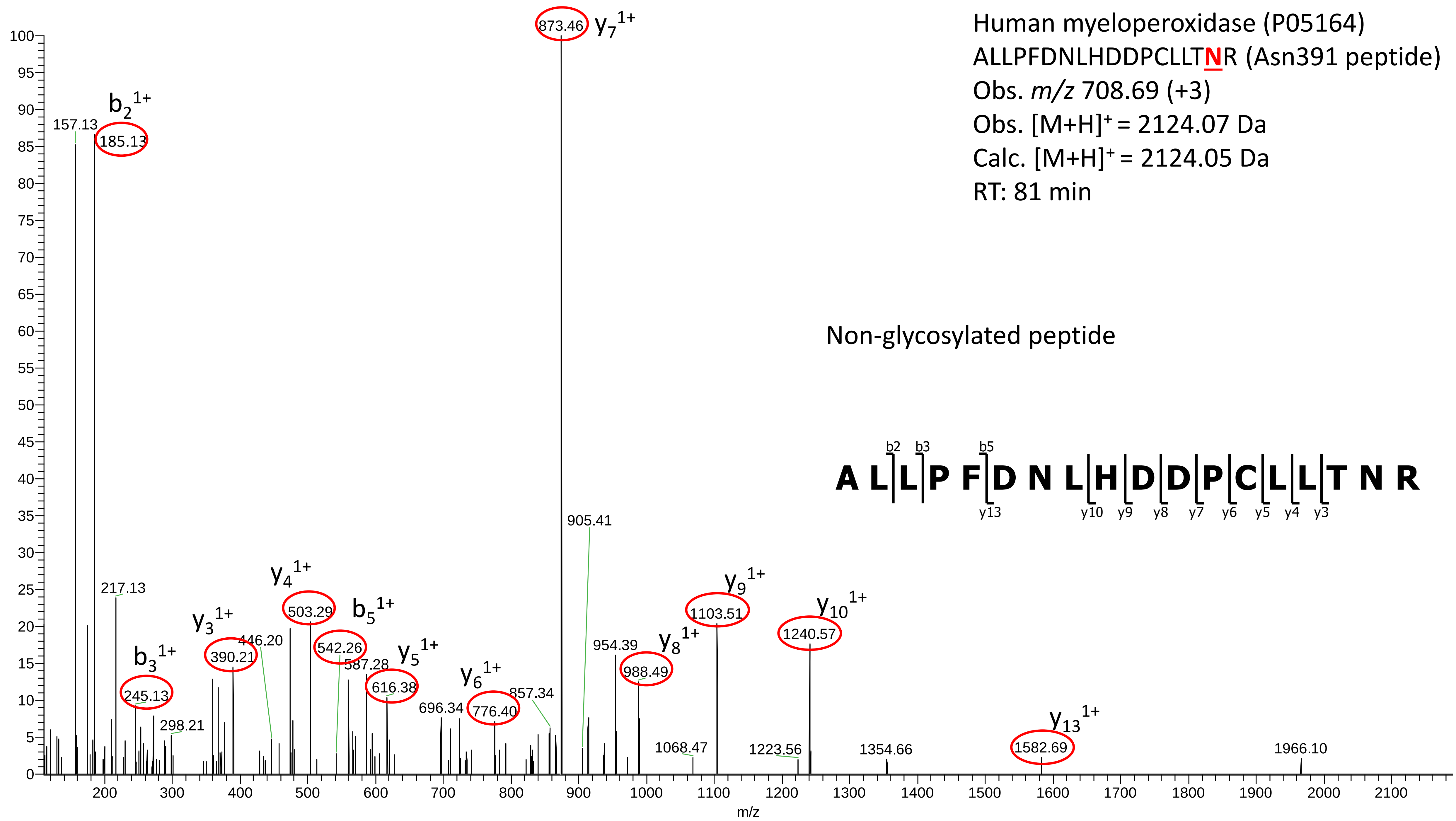
Human myeloperoxidase (P05164)
ALLPFDNLHDDPCLLTNR (Asn391 peptide)
Obs. m/z 708.69 (+3)
Obs. [M+H]+ = 2124.07 Da
Calc. [M+H]+ = 2124.05 Da
RT: 81 min
Non-glycosylated peptide
A L L P F D N L H D D P C L L T N R
b2 b3 b5
y13 y10 y9 y8 y7 y6 y5 y4 y3
873.46 y7 1+
b2 1+
157.13
185.13
217.13
b3 1+
245.13
298.21
y3 1+
390.21
446.20
y4 1+
503.29
b5 1+
542.26
587.28
y5 1+
616.38
696.34
y6 1+
776.40
857.34
905.41
954.39
y8 1+
988.49
1068.47
y9 1+
1103.51
1223.56
y10 1+
1240.57
1354.66
y13 1+
1582.69
1966.10
m/z

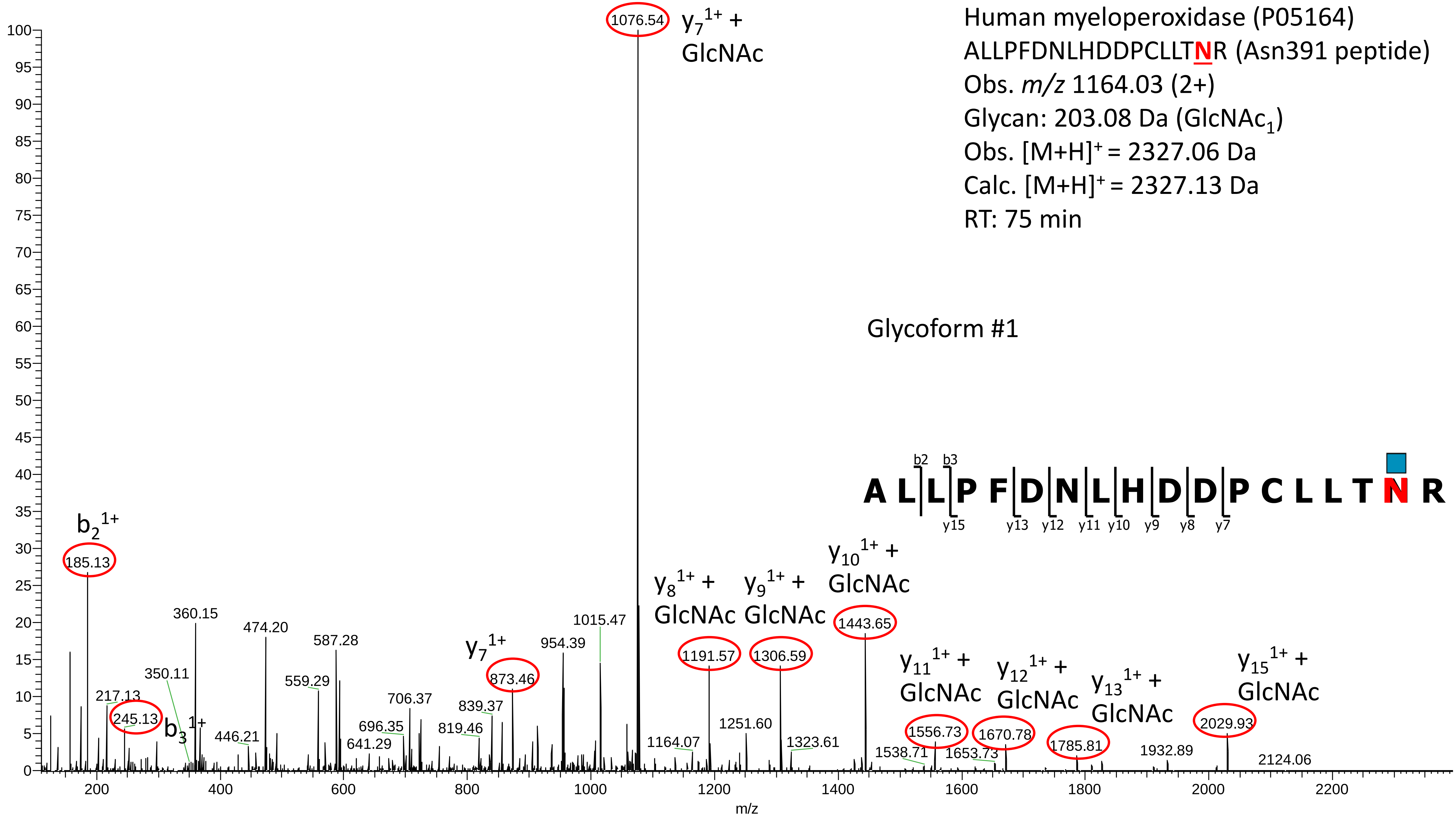

Human myeloperoxidase (P05164)
ALLPFDNLHDDPCLLTNR (Asn391 peptide)
Obs. m/z 1164.03 (2+)
Glycan: 203.08 Da (GlcNAc1)
Obs. [M+H]+ = 2327.06 Da
Calc. [M+H]+ = 2327.13 Da
RT: 75 min
Glycoform #1
A L L P F D N L H D D P C L L T N R
b2 b3
y15 y13 y12 y11 y10 y9 y8 y7
1076.54 y7 1+ + GlcNAc
b2 1+ 185.13
245.13 b3 1+
y7 1+ 873.46
y8 1+ + GlcNAc 1191.57
y9 1+ + GlcNAc 1306.59
y10 1+ + GlcNAc 1443.65
y11 1+ + GlcNAc 1556.73
y12 1+ + GlcNAc 1670.78
y13 1+ + GlcNAc 1785.81
y15 1+ + GlcNAc 2029.93
217.13
350.11
360.15
446.21
474.20
559.29
587.28
641.29
696.35
706.37
819.46
839.37
954.39
1015.47
1164.07
1251.60
1323.61
1538.71
1653.73
1932.89
2124.06
m/z

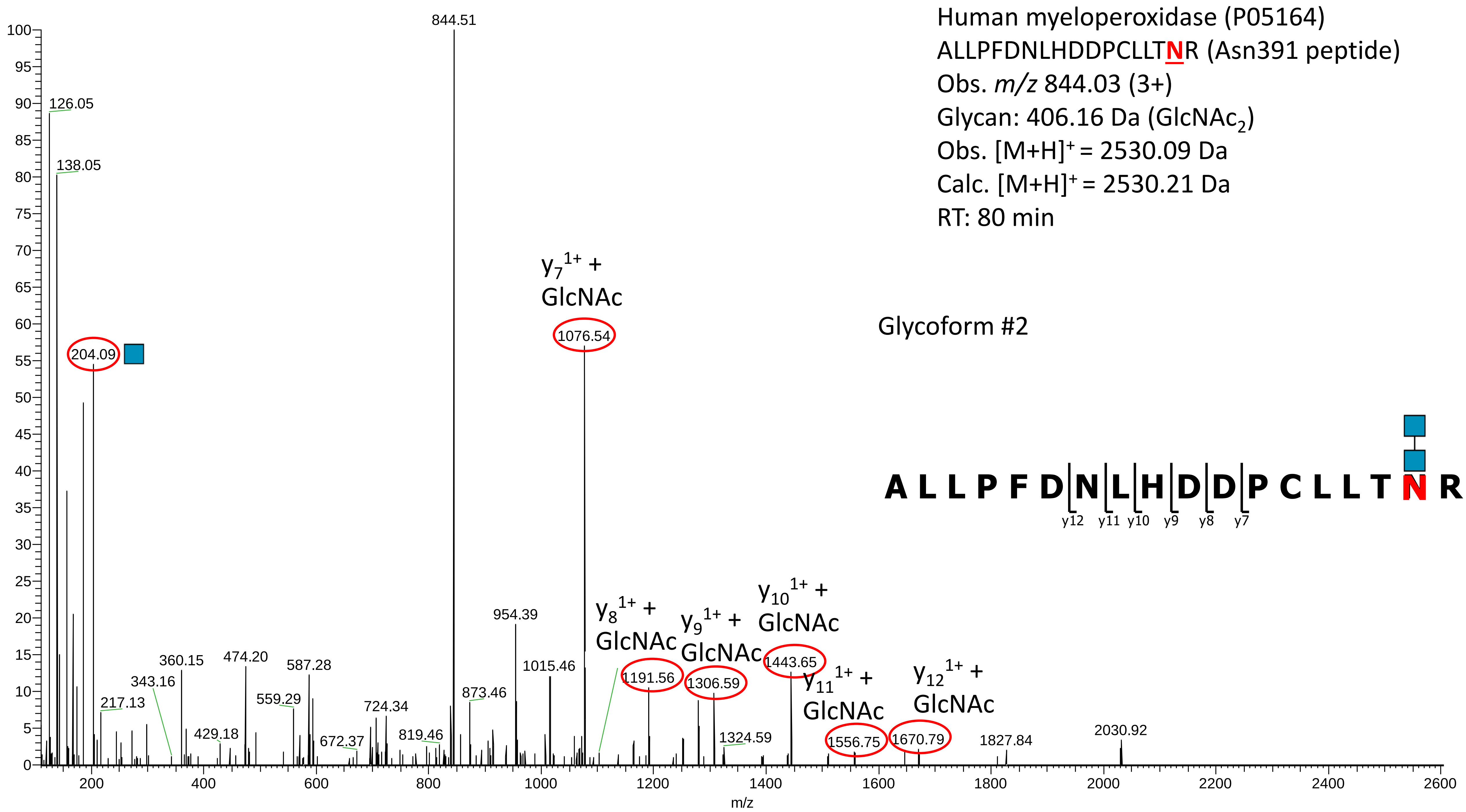
Human myeloperoxidase (P05164)
ALLPFDNLHDDPCLLTNR (Asn391 peptide)
Obs. m/z 844.03 (3+)
Glycan: 406.16 Da (GlcNAc2)
Obs. [M+H]+ = 2530.09 Da
Calc. [M+H]+ = 2530.21 Da
RT: 80 min
Glycoform #2
A L L P F D N L H D D P C L L T N R
y12 y11 y10 y9 y8 y7
y7 1+ + GlcNAc
y8 1+ + GlcNAc
y9 1+ + GlcNAc
y10 1+ + GlcNAc
y11 1+ + GlcNAc
y12 1+ + GlcNAc
844.51
126.05
138.05
204.09
1076.54
954.39
1015.46
873.46
819.46
724.34
672.37
587.28
559.29
474.20
429.18
360.15
343.16
217.13
1191.56
1306.59
1324.59
1443.65
1556.75
1670.79
1827.84
2030.92
m/z

Human myeloperoxidase (P05164)
ALLPFDNLHDDPCLLTNR (Asn391 peptide)
Obs. *m/z* 898.09 (3+)
Glycan: 568.21 Da ($GlcNAc_2Hex_1$)
Obs. $[M+H]^+$ = 2692.18 Da
Calc. $[M+H]^+$ = 2692.26 Da
RT: 77 min

Glycoform #3

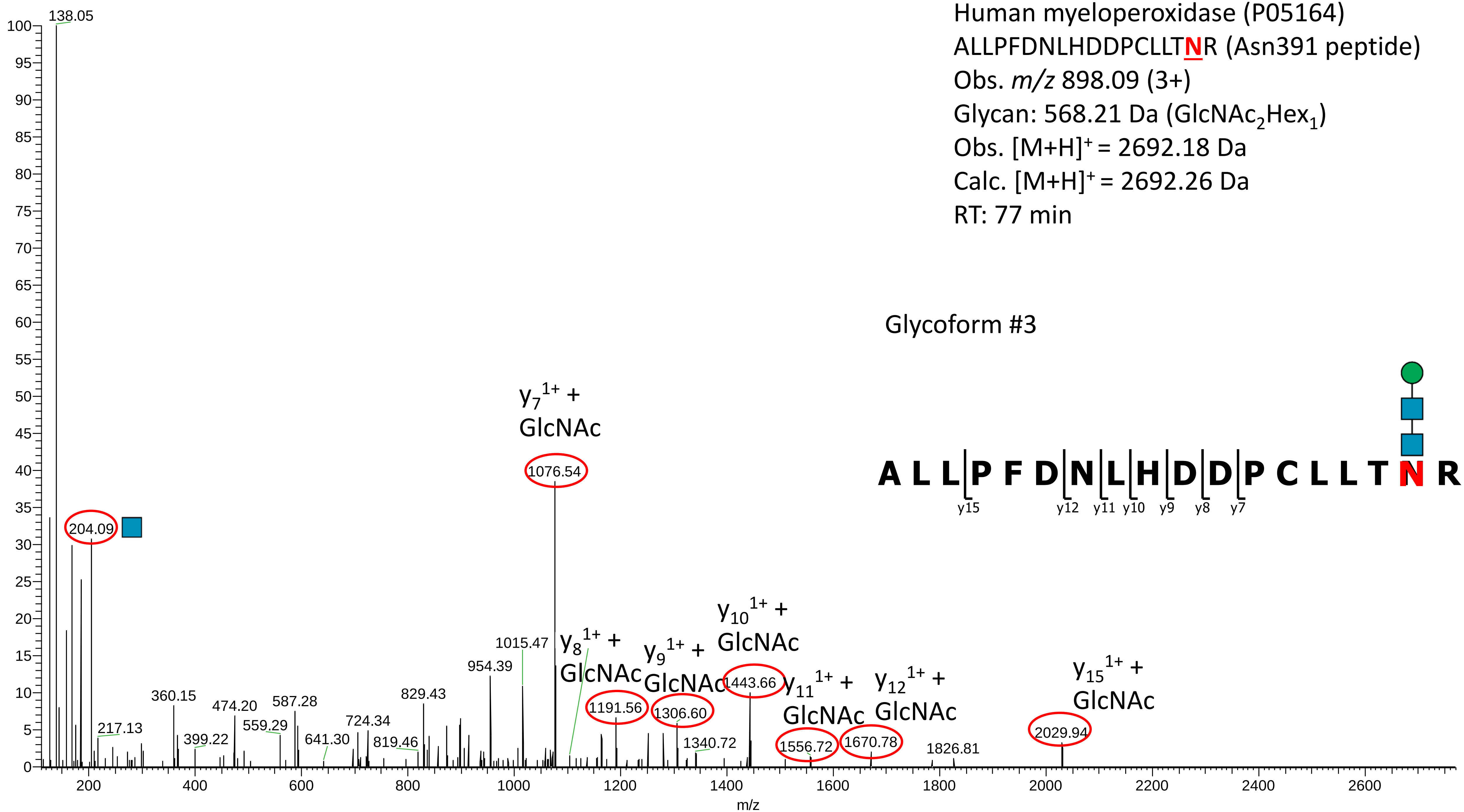

Human myeloperoxidase (P05164)

ALLPFDNLHDDPCLLT**N**R (Asn391 peptide)

Obs. *m/z* 952.45 (3+)

Glycan: 730.26 Da ($GlcNAc_2Hex_2$)

Obs. $[M+H]^+$ = 2855.35 Da

Calc. $[M+H]^+$ = 2854.31 Da

Off-by-X (manual) : 1

RT: 78 min

Glycoform #4

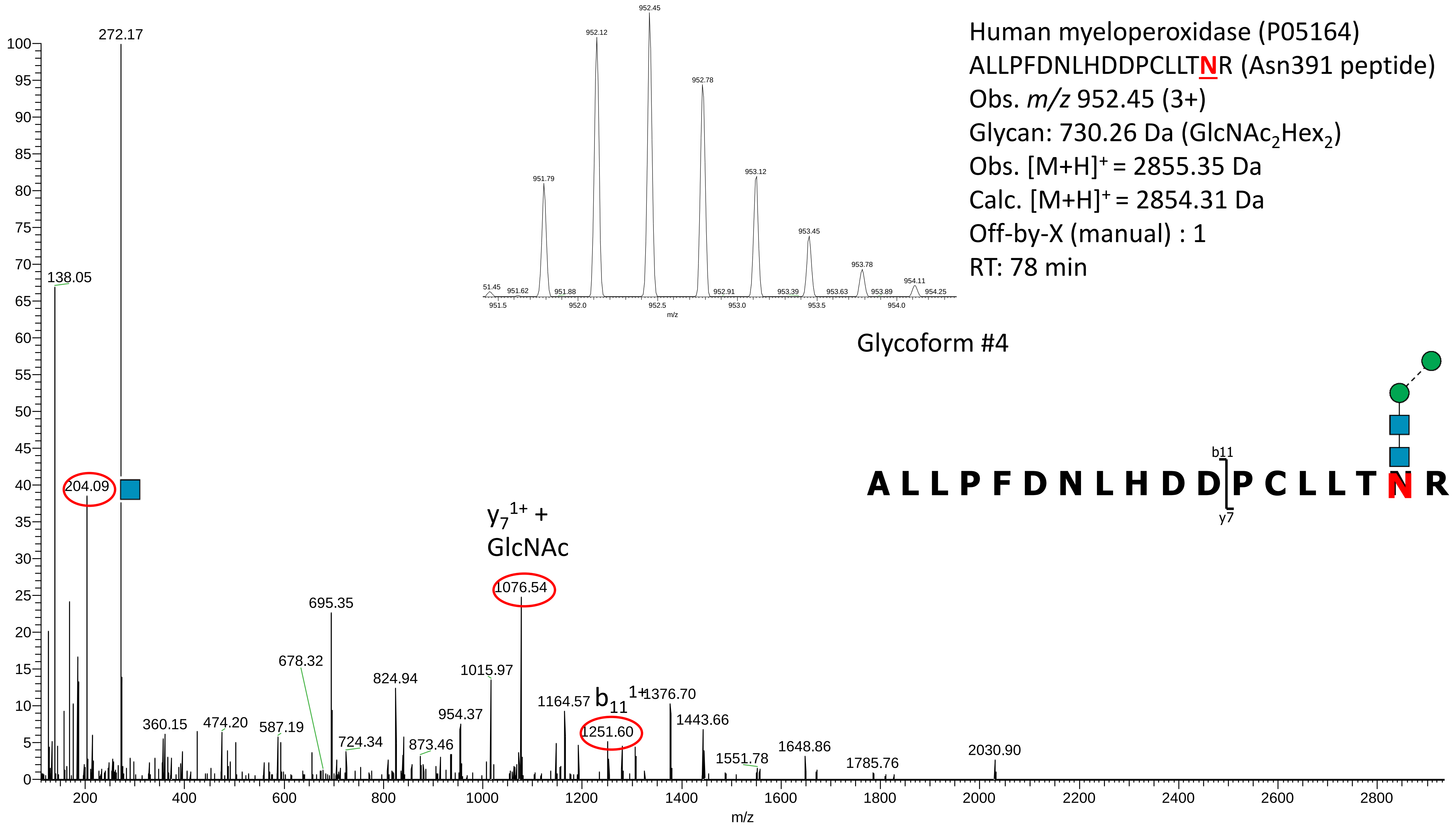

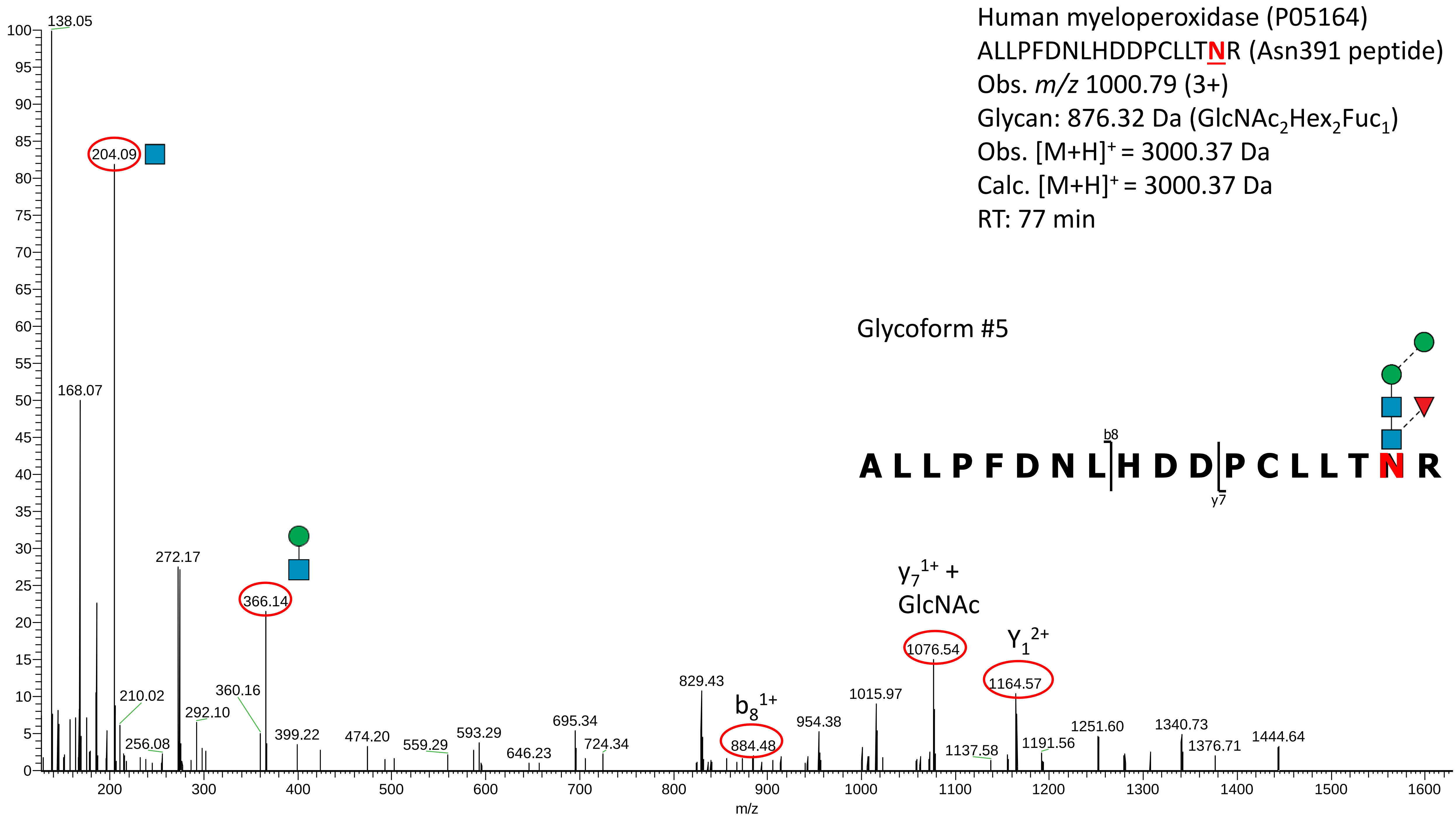
Human myeloperoxidase (P05164)
ALLPFDNLHDDPCLLTNR (Asn391 peptide)
Obs. m/z 1000.79 (3+)
Glycan: 876.32 Da (GlcNAc2Hex2Fuc1)
Obs. [M+H]+ = 3000.37 Da
Calc. [M+H]+ = 3000.37 Da
RT: 77 min
Glycoform #5
A L L P F D N L H D D P C L L T N R
b8
y7
y7 1+ + GlcNAc
Y1 2+
b8 1+
138.05
204.09
168.07
272.17
366.14
360.16
210.02
292.10
256.08
399.22
474.20
559.29
593.29
646.23
695.34
724.34
829.43
884.48
954.38
1015.97
1076.54
1137.58
1164.57
1191.56
1251.60
1340.73
1376.71
1444.64
m/z

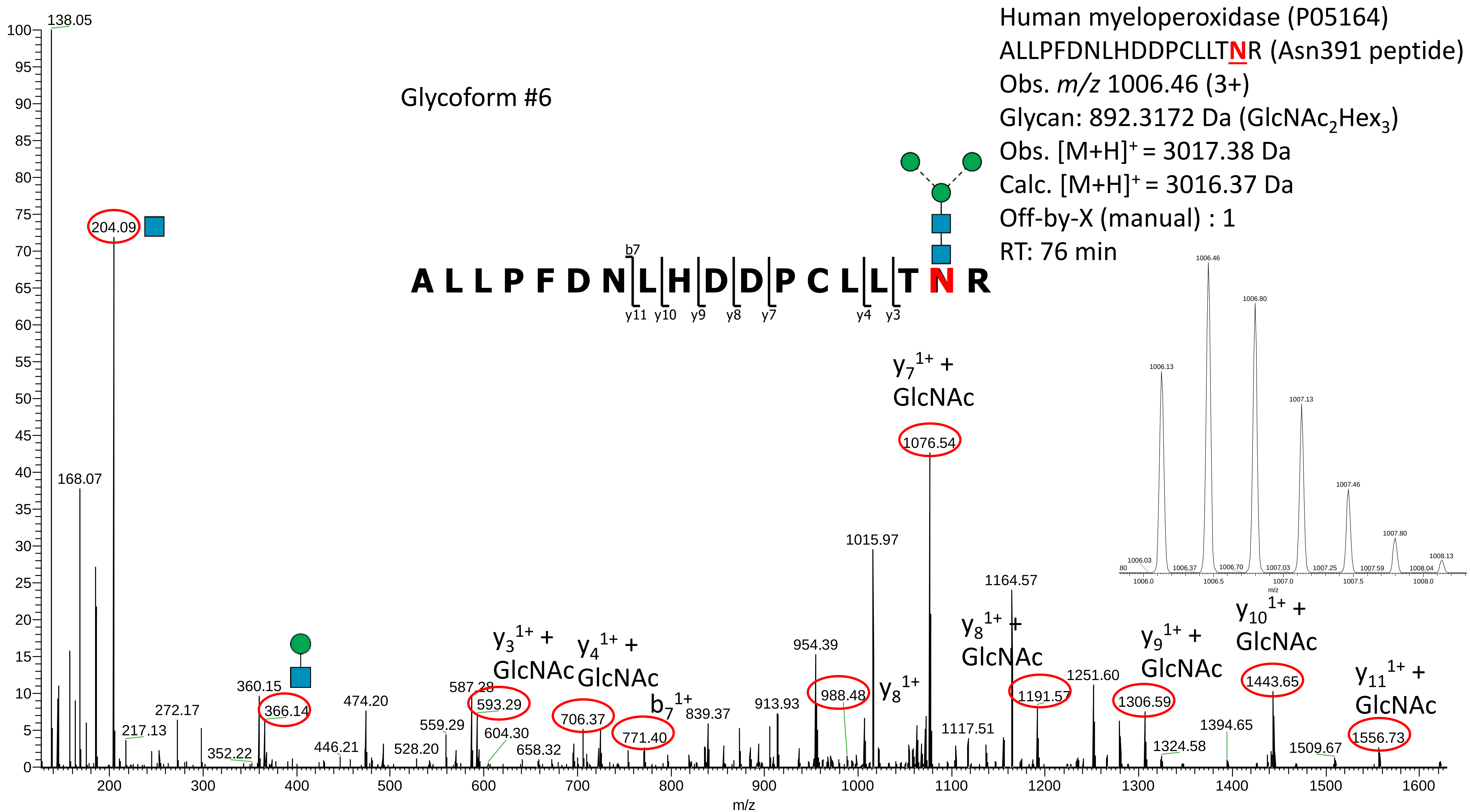

Human myeloperoxidase (P05164)
ALLPFDNLHDDPCLLTNR (Asn391 peptide)
Obs. m/z 1006.46 (3+)
Glycan: 892.3172 Da (GlcNAc2Hex3)
Obs. [M+H]+ = 3017.38 Da
Calc. [M+H]+ = 3016.37 Da
Off-by-X (manual) : 1
RT: 76 min
Glycoform #6
A L L P F D N L H D D P C L L T N R
b7
y11 y10 y9 y8 y7 y4 y3
y7 1+ + GlcNAc
y3 1+ + GlcNAc
y4 1+ + GlcNAc
b7 1+
y8 1+
y8 1+ + GlcNAc
y9 1+ + GlcNAc
y10 1+ + GlcNAc
y11 1+ + GlcNAc
138.05
168.07
204.09
217.13
272.17
352.22
360.15
366.14
446.21
474.20
528.20
559.29
587.28
593.29
604.30
658.32
706.37
771.40
839.37
913.93
954.39
988.48
1015.97
1076.54
1117.51
1164.57
1191.57
1251.60
1306.59
1324.58
1394.65
1443.65
1509.67
1556.73
m/z

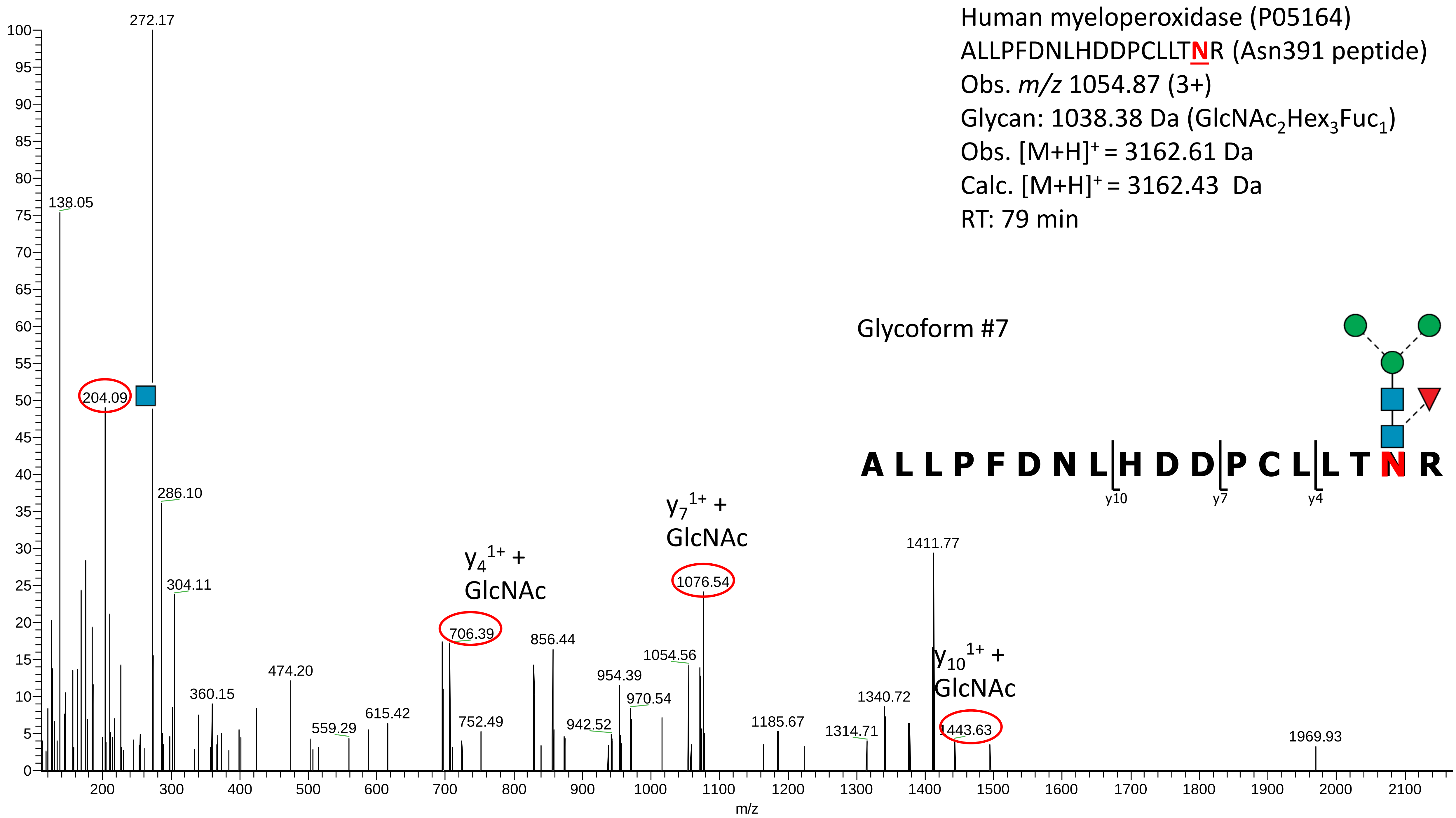
Human myeloperoxidase (P05164)
ALLPFDNLHDDPCLLTNR (Asn391 peptide)
Obs. m/z 1054.87 (3+)
Glycan: 1038.38 Da (GlcNAc2Hex3Fuc1)
Obs. [M+H]+ = 3162.61 Da
Calc. [M+H]+ = 3162.43 Da
RT: 79 min
Glycoform #7
A L L P F D N L H D D P C L L T N R
y10
y7
y4
272.17
138.05
204.09
286.10
304.11
360.15
474.20
559.29
615.42
706.39
y4 1+ + GlcNAc
752.49
856.44
942.52
954.39
970.54
1054.56
1076.54
y7 1+ + GlcNAc
1185.67
1314.71
1340.72
1411.77
1443.63
y10 1+ + GlcNAc
1969.93
m/z

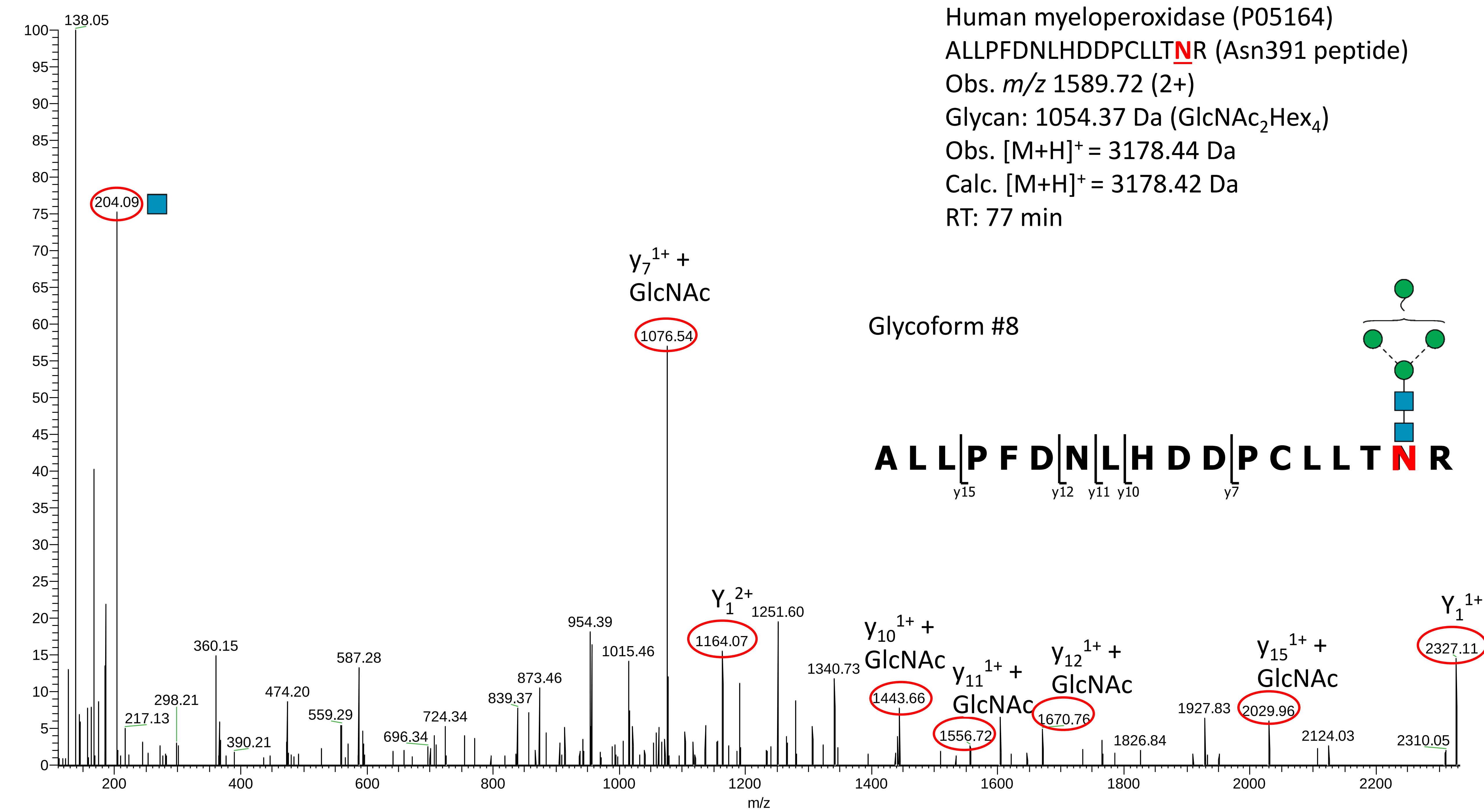
Human myeloperoxidase (P05164)
ALLPFDNLHDDPCLLTNR (Asn391 peptide)
Obs. m/z 1589.72 (2+)
Glycan: 1054.37 Da (GlcNAc2Hex4)
Obs. [M+H]+ = 3178.44 Da
Calc. [M+H]+ = 3178.42 Da
RT: 77 min
Glycoform #8
A L L P F D N L H D D P C L L T N R
y15
y12
y11
y10
y7
138.05
204.09
y7 1+ + GlcNAc
1076.54
Y1 2+
1164.07
1251.60
954.39
1015.46
360.15
587.28
474.20
298.21
217.13
390.21
559.29
696.34
724.34
839.37
873.46
1340.73
y10 1+ + GlcNAc
1443.66
y11 1+ + GlcNAc
1556.72
y12 1+ + GlcNAc
1670.76
1826.84
1927.83
y15 1+ + GlcNAc
2029.96
2124.03
2310.05
Y1 1+
2327.11
m/z

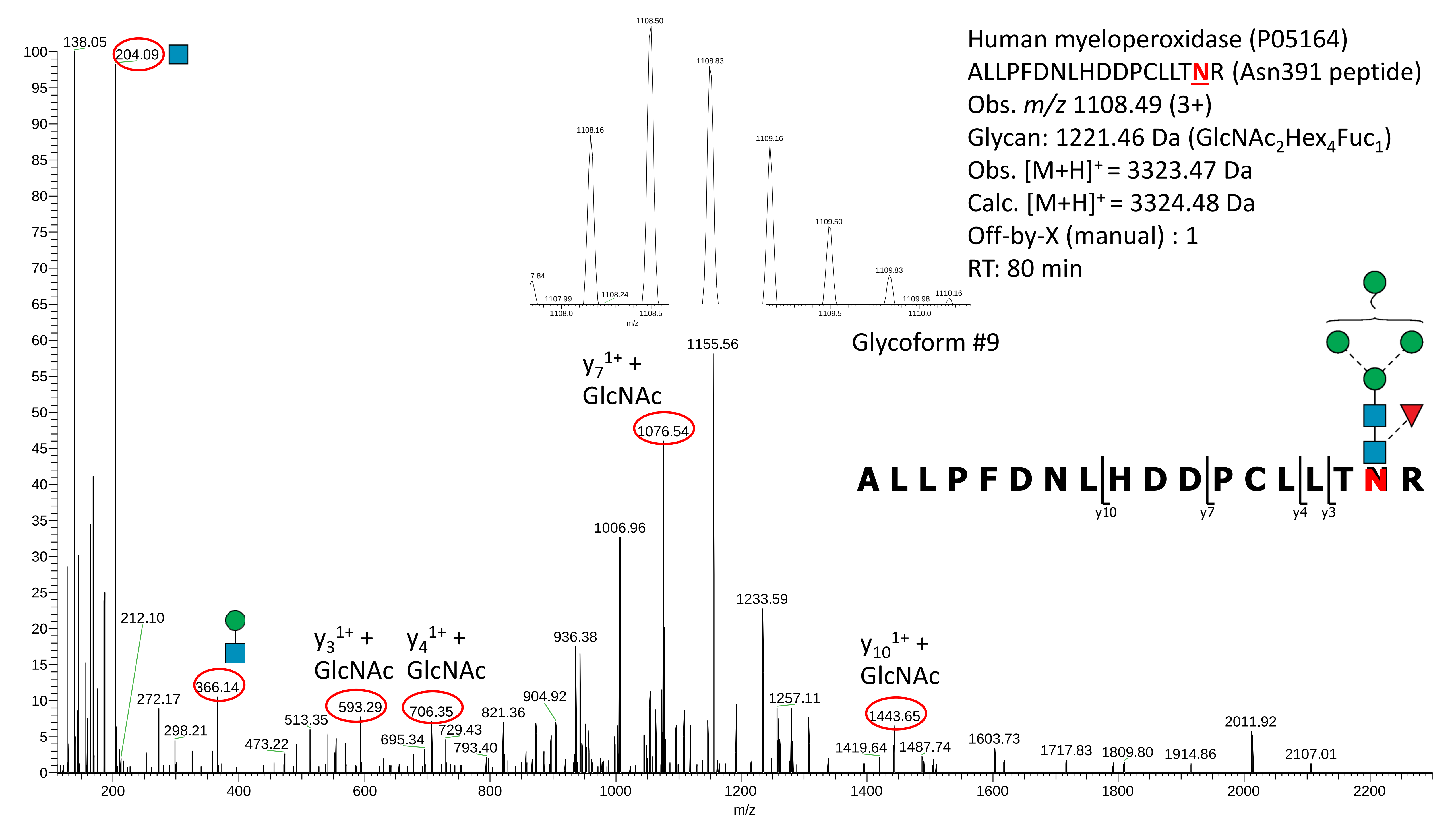

Human myeloperoxidase (P05164)
ALLPFDNLHDDPCLLTNR (Asn391 peptide)
Obs. m/z 1108.49 (3+)
Glycan: 1221.46 Da (GlcNAc2Hex4Fuc1)
Obs. [M+H]+ = 3323.47 Da
Calc. [M+H]+ = 3324.48 Da
Off-by-X (manual) : 1
RT: 80 min
Glycoform #9
A L L P F D N L H D D P C L L T N R
y10
y7
y4
y3
y7 1+ + GlcNAc
y3 1+ + GlcNAc
y4 1+ + GlcNAc
y10 1+ + GlcNAc
138.05
204.09
212.10
272.17
298.21
366.14
473.22
513.35
593.29
695.34
706.35
729.43
793.40
821.36
904.92
936.38
1006.96
1076.54
1155.56
1233.59
1257.11
1419.64
1443.65
1487.74
1603.73
1717.83
1809.80
1914.86
2011.92
2107.01
1108.50
1108.83
1108.16
1109.16
1109.50
1109.83
7.84
1107.99
1108.24
1109.98
1110.16
m/z

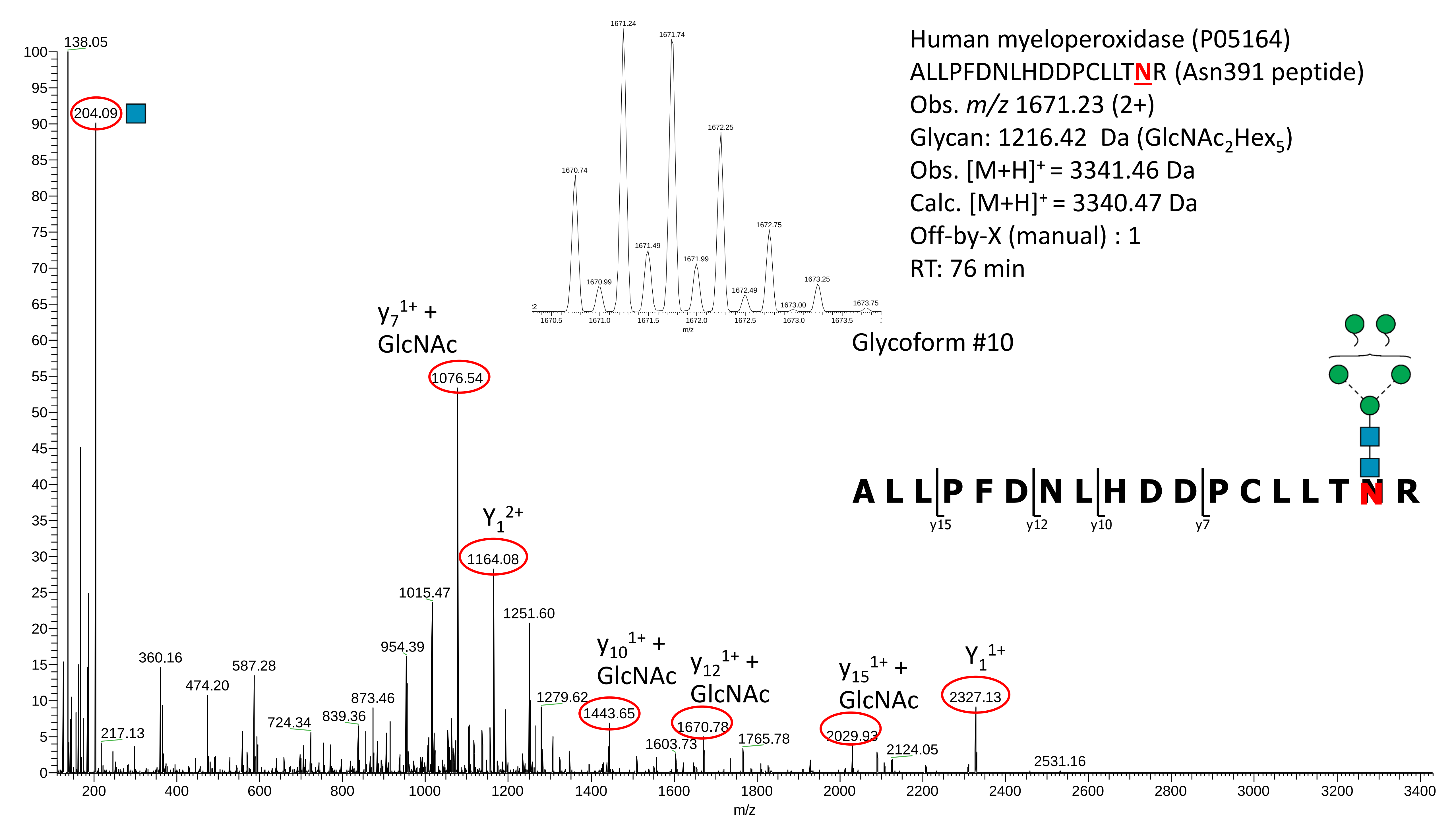

Human myeloperoxidase (P05164)
ALLPFDNLHDDPCLLTNR (Asn391 peptide)
Obs. m/z 1671.23 (2+)
Glycan: 1216.42 Da (GlcNAc2Hex5)
Obs. [M+H]+ = 3341.46 Da
Calc. [M+H]+ = 3340.47 Da
Off-by-X (manual) : 1
RT: 76 min
Glycoform #10
A L L P F D N L H D D P C L L T N R
y15
y12
y10
y7
138.05
204.09
217.13
360.16
474.20
587.28
724.34
839.36
873.46
954.39
1015.47
y7 1+ + GlcNAc
1076.54
Y1 2+
1164.08
1251.60
1279.62
y10 1+ + GlcNAc
1443.65
1603.73
y12 1+ + GlcNAc
1670.78
1765.78
y15 1+ + GlcNAc
2029.93
2124.05
Y1 1+
2327.13
2531.16
1670.74
1670.99
1671.24
1671.49
1671.74
1671.99
1672.25
1672.49
1672.75
1673.00
1673.25
1673.75
m/z

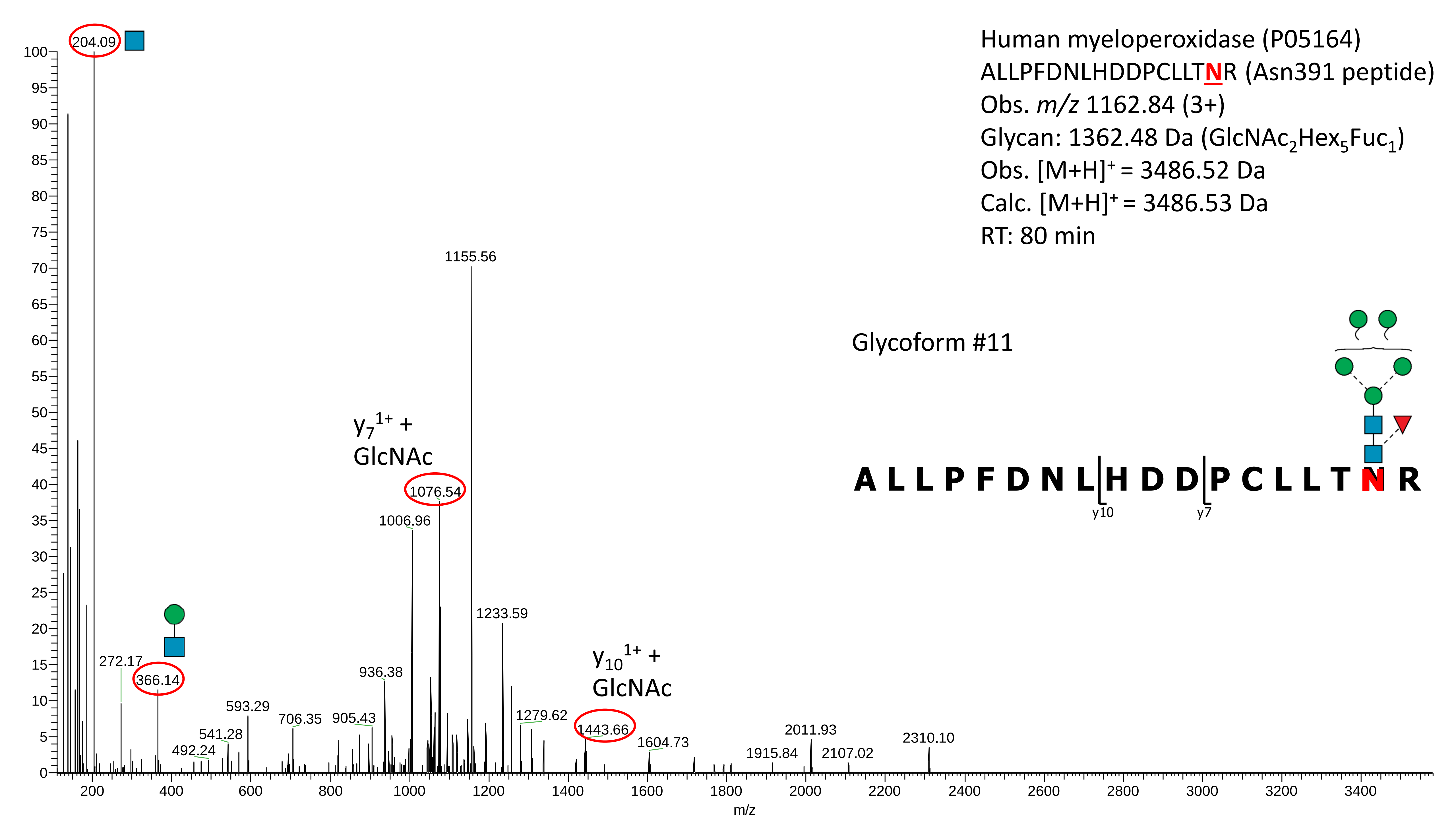

Human myeloperoxidase (P05164)
ALLPFDNLHDDPCLLTNR (Asn391 peptide)
Obs. m/z 1162.84 (3+)
Glycan: 1362.48 Da (GlcNAc2Hex5Fuc1)
Obs. [M+H]+ = 3486.52 Da
Calc. [M+H]+ = 3486.53 Da
RT: 80 min
Glycoform #11
A L L P F D N L H D D P C L L T N R
y10
y7
y7 1+ + GlcNAc
y10 1+ + GlcNAc
204.09
1155.56
1076.54
1006.96
1233.59
936.38
272.17
366.14
593.29
706.35
905.43
541.28
492.24
1279.62
1443.66
1604.73
2011.93
1915.84
2107.02
2310.10
m/z

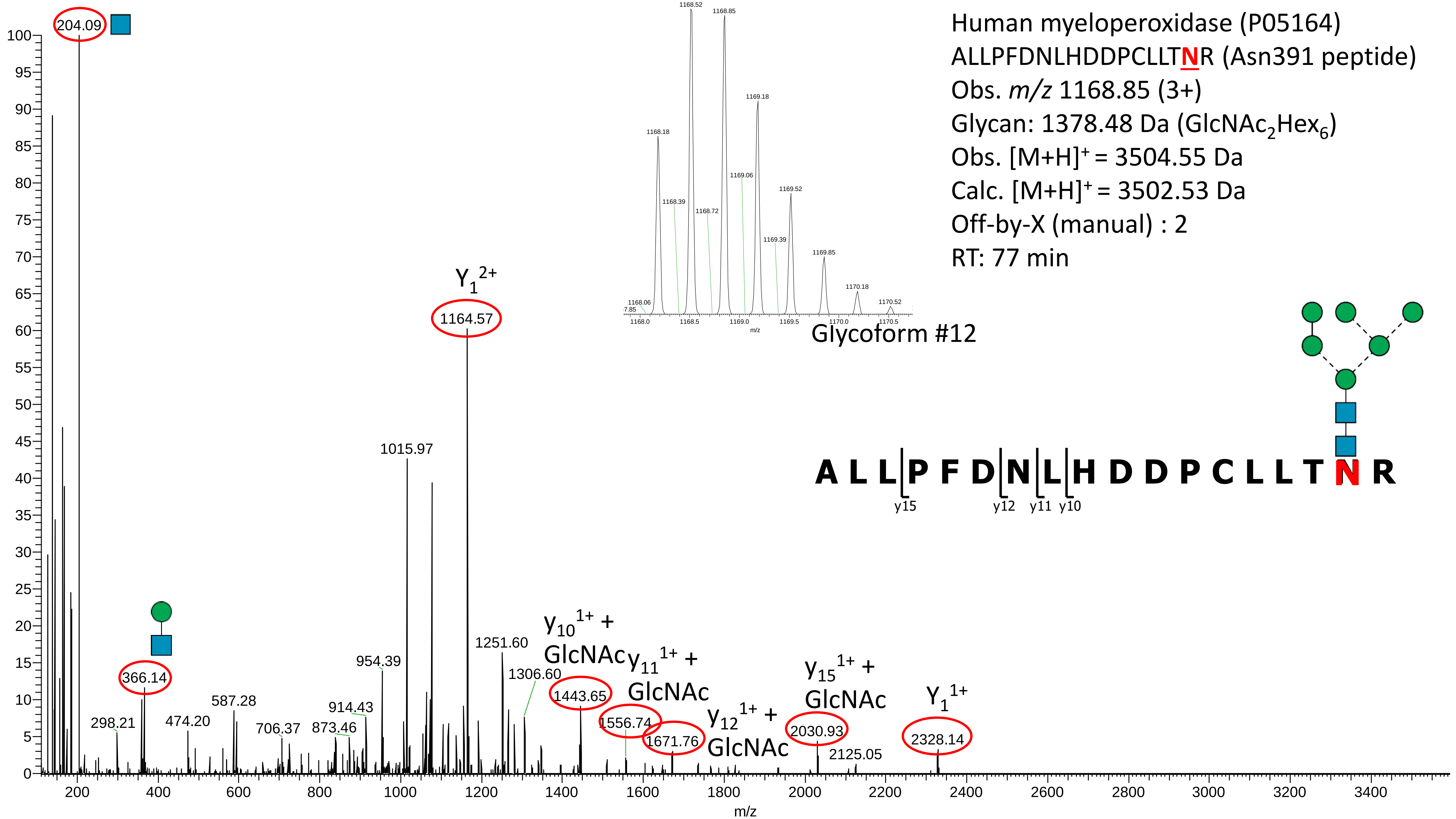
Human myeloperoxidase (P05164)
ALLPFDNLHDDPCLLTNR (Asn391 peptide)
Obs. m/z 1168.85 (3+)
Glycan: 1378.48 Da (GlcNAc2Hex6)
Obs. [M+H]+ = 3504.55 Da
Calc. [M+H]+ = 3502.53 Da
Off-by-X (manual) : 2
RT: 77 min
Glycoform #12
A L L P F D N L H D D P C L L T N R
y15 y12 y11 y10
204.09
Y1 2+
1164.57
1015.97
1251.60
1306.60
954.39
914.43
873.46
706.37
587.28
474.20
366.14
298.21
y10 1+ + GlcNAc
1443.65
y11 1+ + GlcNAc
1556.74
1671.76
y12 1+ + GlcNAc
y15 1+ + GlcNAc
2030.93
2125.05
Y1 1+
2328.14
m/z

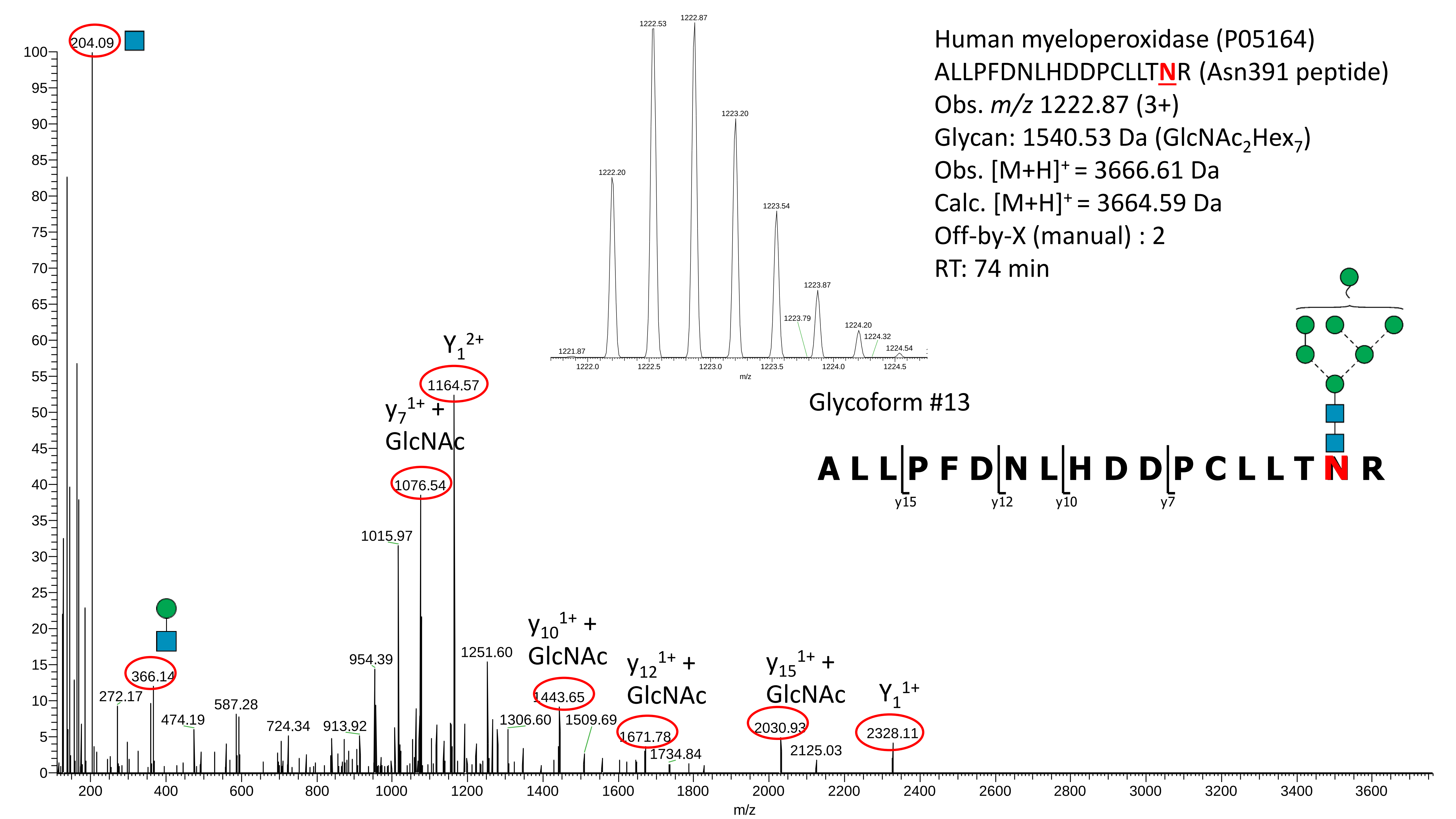

Human myeloperoxidase (P05164)
ALLPFDNLHDDPCLLTNR (Asn391 peptide)
Obs. m/z 1222.87 (3+)
Glycan: 1540.53 Da (GlcNAc2Hex7)
Obs. [M+H]+ = 3666.61 Da
Calc. [M+H]+ = 3664.59 Da
Off-by-X (manual) : 2
RT: 74 min
Glycoform #13
A L L P F D N L H D D P C L L T N R
y15
y12
y10
y7
204.09
Y1 2+
1164.57
y7 1+ + GlcNAc
1076.54
1015.97
954.39
1251.60
y10 1+ + GlcNAc
1443.65
y12 1+ + GlcNAc
1671.78
y15 1+ + GlcNAc
2030.93
Y1 1+
2328.11
366.14
272.17
474.19
587.28
724.34
913.92
1306.60
1509.69
1734.84
2125.03
1222.53
1222.87
1223.20
1222.20
1223.54
1223.87
1223.79
1224.20
1224.32
1224.54
1221.87
m/z

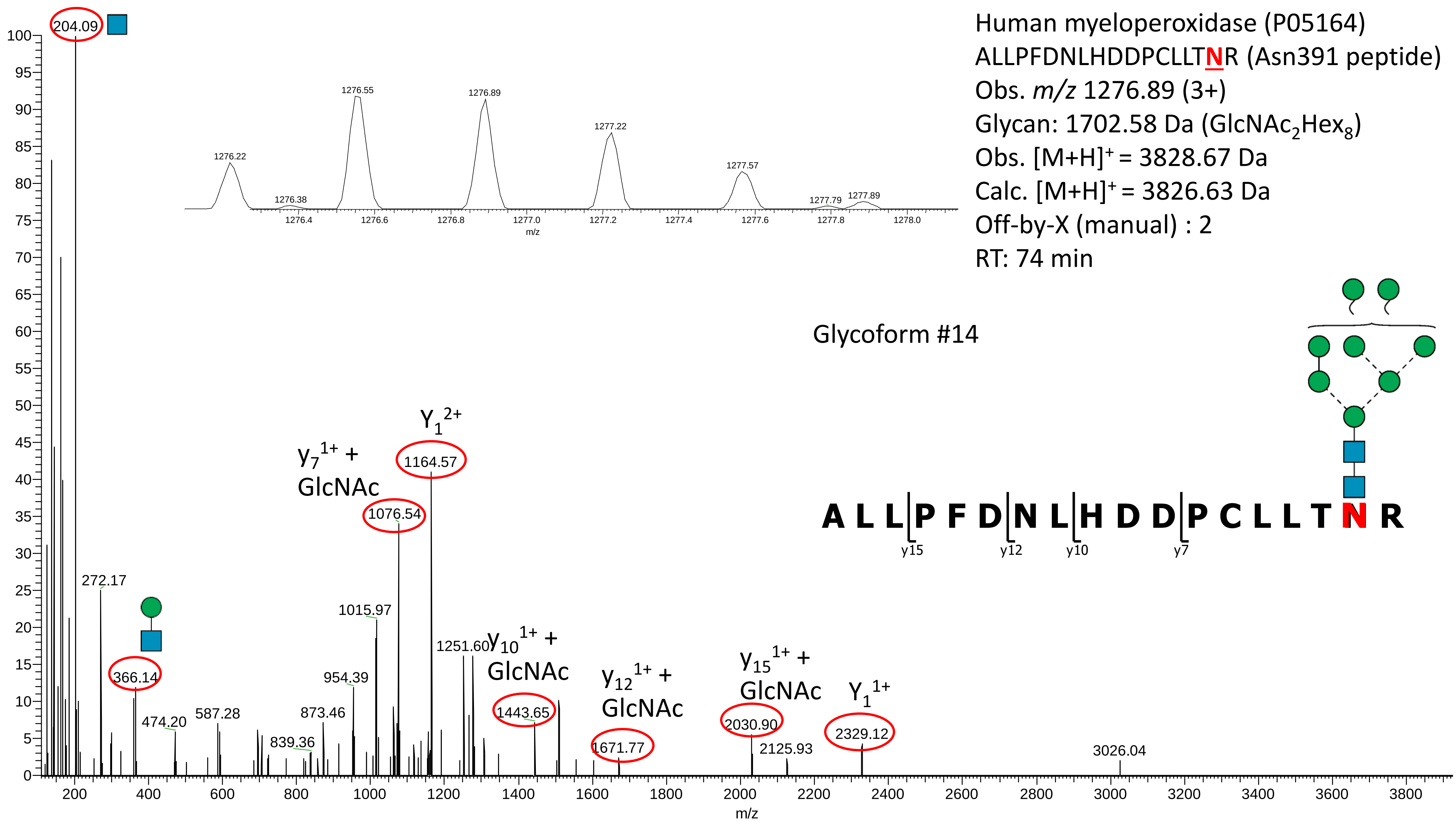
Human myeloperoxidase (P05164)
ALLPFDNLHDDPCLLTNR (Asn391 peptide)
Obs. m/z 1276.89 (3+)
Glycan: 1702.58 Da (GlcNAc2Hex8)
Obs. [M+H]+ = 3828.67 Da
Calc. [M+H]+ = 3826.63 Da
Off-by-X (manual) : 2
RT: 74 min
Glycoform #14
A L L P F D N L H D D P C L L T N R
y15
y12
y10
y7
204.09
272.17
366.14
474.20
587.28
839.36
873.46
954.39
1015.97
y7 1+ + GlcNAc
1076.54
Y1 2+
1164.57
1251.60
y10 1+ + GlcNAc
1443.65
y12 1+ + GlcNAc
1671.77
y15 1+ + GlcNAc
2030.90
2125.93
Y1 1+
2329.12
3026.04
1276.22
1276.38
1276.55
1276.89
1277.22
1277.57
1277.79
1277.89
m/z

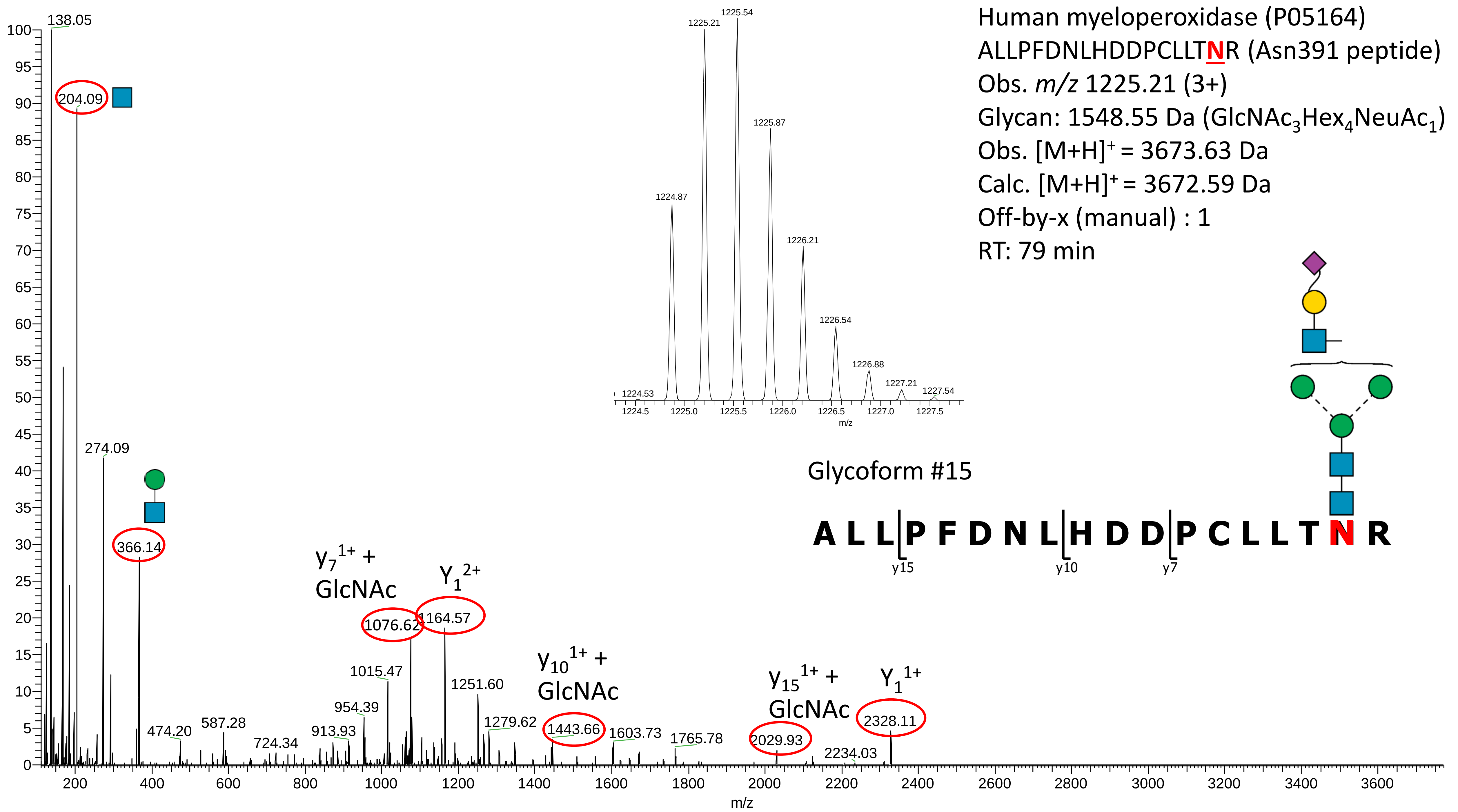

Human myeloperoxidase (P05164)
ALLPFDNLHDDPCLLTNR (Asn391 peptide)
Obs. m/z 1225.21 (3+)
Glycan: 1548.55 Da (GlcNAc3Hex4NeuAc1)
Obs. [M+H]+ = 3673.63 Da
Calc. [M+H]+ = 3672.59 Da
Off-by-x (manual) : 1
RT: 79 min
Glycoform #15
A L L P F D N L H D D P C L L T N R
y15
y10
y7
138.05
204.09
274.09
366.14
474.20
587.28
724.34
913.93
954.39
1015.47
y7 1+ + GlcNAc
1076.62
Y1 2+
1164.57
1251.60
1279.62
y10 1+ + GlcNAc
1443.66
1603.73
1765.78
y15 1+ + GlcNAc
2029.93
2234.03
Y1 1+
2328.11
1224.53
1224.87
1225.21
1225.54
1225.87
1226.21
1226.54
1226.88
1227.21
1227.54
m/z

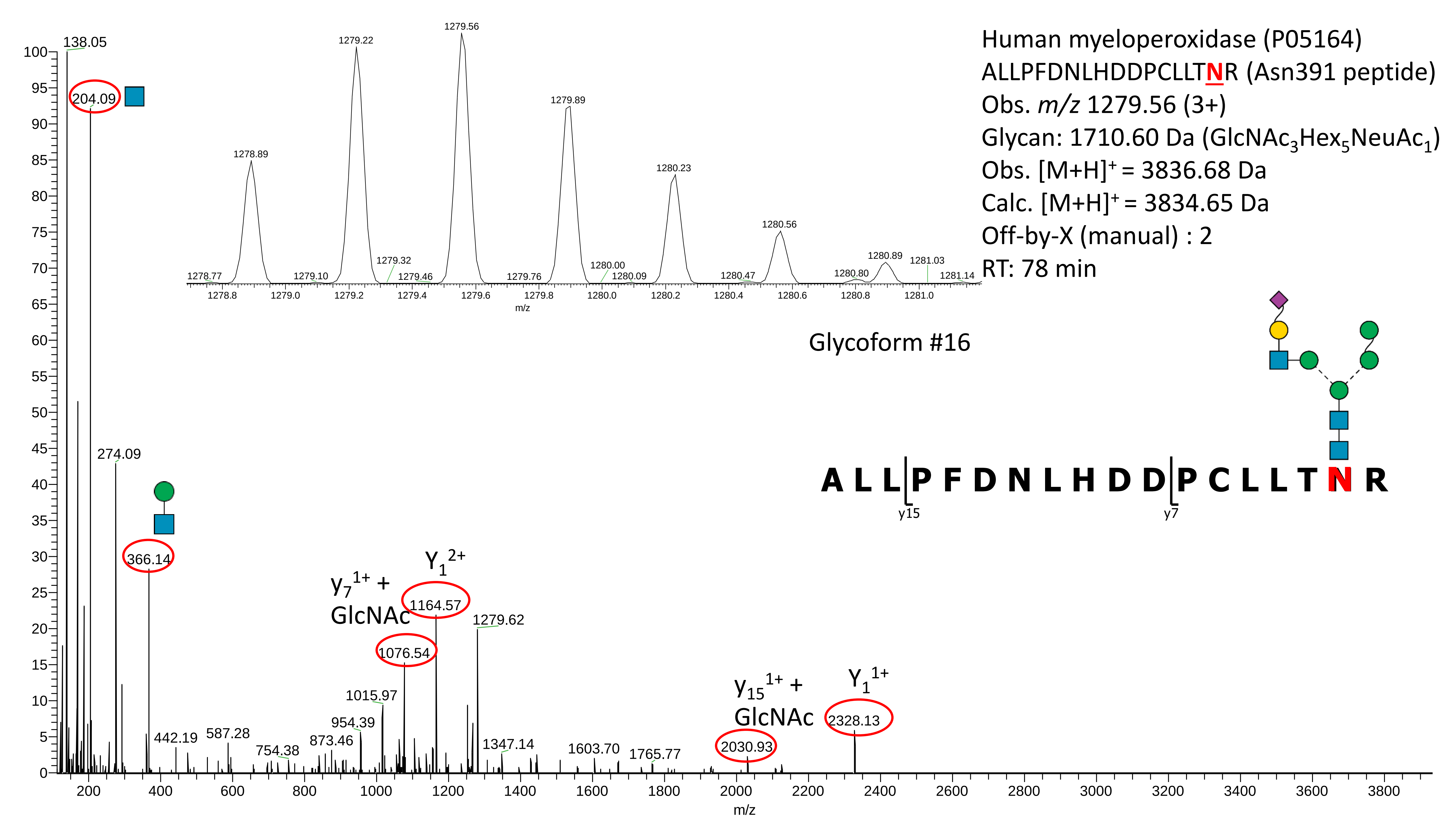

Human myeloperoxidase (P05164)
ALLPFDNLHDDPCLLTNR (Asn391 peptide)
Obs. m/z 1279.56 (3+)
Glycan: 1710.60 Da (GlcNAc3Hex5NeuAc1)
Obs. [M+H]+ = 3836.68 Da
Calc. [M+H]+ = 3834.65 Da
Off-by-X (manual) : 2
RT: 78 min
Glycoform #16
A L L P F D N L H D D P C L L T N R
y15
y7
138.05
204.09
274.09
366.14
442.19
587.28
754.38
873.46
954.39
1015.97
1076.54
1164.57
1279.62
1347.14
1603.70
1765.77
2030.93
2328.13
y7 1+ + GlcNAc
Y1 2+
y15 1+ + GlcNAc
Y1 1+
1278.77
1278.89
1279.10
1279.22
1279.32
1279.46
1279.56
1279.76
1279.89
1280.00
1280.09
1280.23
1280.47
1280.56
1280.80
1280.89
1281.03
1281.14
m/z

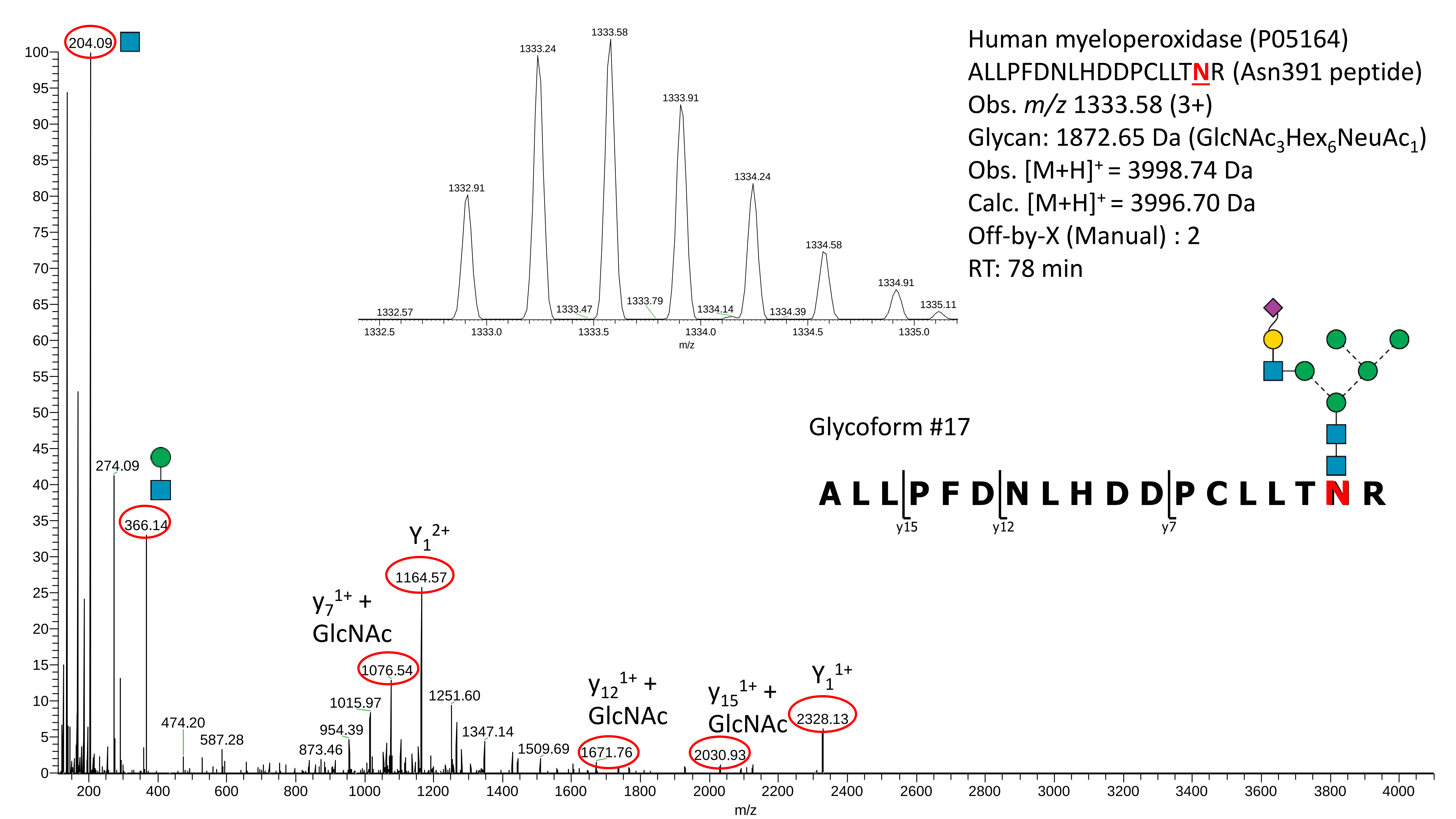
Human myeloperoxidase (P05164)
ALLPFDNLHDDPCLLTNR (Asn391 peptide)
Obs. m/z 1333.58 (3+)
Glycan: 1872.65 Da (GlcNAc3Hex6NeuAc1)
Obs. [M+H]+ = 3998.74 Da
Calc. [M+H]+ = 3996.70 Da
Off-by-X (Manual) : 2
RT: 78 min
Glycoform #17
A L L P F D N L H D D P C L L T N R
y15
y12
y7
204.09
274.09
366.14
474.20
587.28
873.46
954.39
1015.97
1076.54
1164.57
1251.60
1347.14
1509.69
1671.76
2030.93
2328.13
Y1 2+
y7 1+ + GlcNAc
y12 1+ + GlcNAc
y15 1+ + GlcNAc
Y1 1+
1332.57
1332.91
1333.24
1333.47
1333.58
1333.79
1333.91
1334.14
1334.24
1334.39
1334.58
1334.91
1335.11
m/z

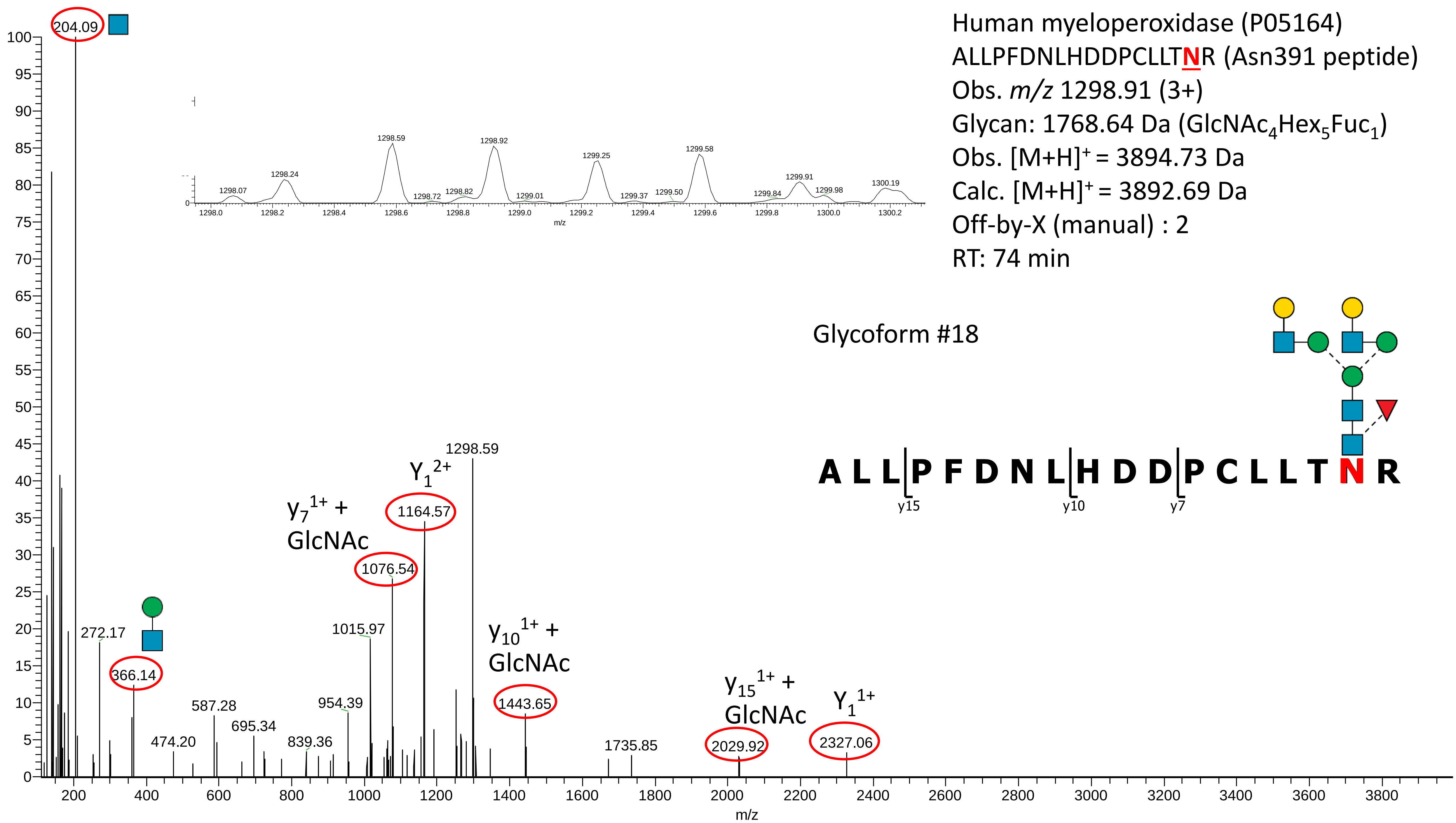

Human myeloperoxidase (P05164)
ALLPFDNLHDDPCLLTNR (Asn391 peptide)
Obs. m/z 1298.91 (3+)
Glycan: 1768.64 Da (GlcNAc4Hex5Fuc1)
Obs. [M+H]+ = 3894.73 Da
Calc. [M+H]+ = 3892.69 Da
Off-by-X (manual) : 2
RT: 74 min
Glycoform #18
A L L P F D N L H D D P C L L T N R
y15
y10
y7
204.09
272.17
366.14
474.20
587.28
695.34
839.36
954.39
1015.97
y7 1+ + GlcNAc
1076.54
Y1 2+
1164.57
1298.59
y10 1+ + GlcNAc
1443.65
1735.85
y15 1+ + GlcNAc
2029.92
Y1 1+
2327.06
1298.07
1298.24
1298.59
1298.72
1298.82
1298.92
1299.01
1299.25
1299.37
1299.50
1299.58
1299.84
1299.91
1299.98
1300.19
m/z

**Supplementary Data S3D**

Manually annotated reversed phase-LC-ESI-HCD-MS/MS (+) spectra of all glycosylated and non-glycosylated peptides containing Asn483 identified from the analysis of the unenriched peptide mixtures of nMPO.

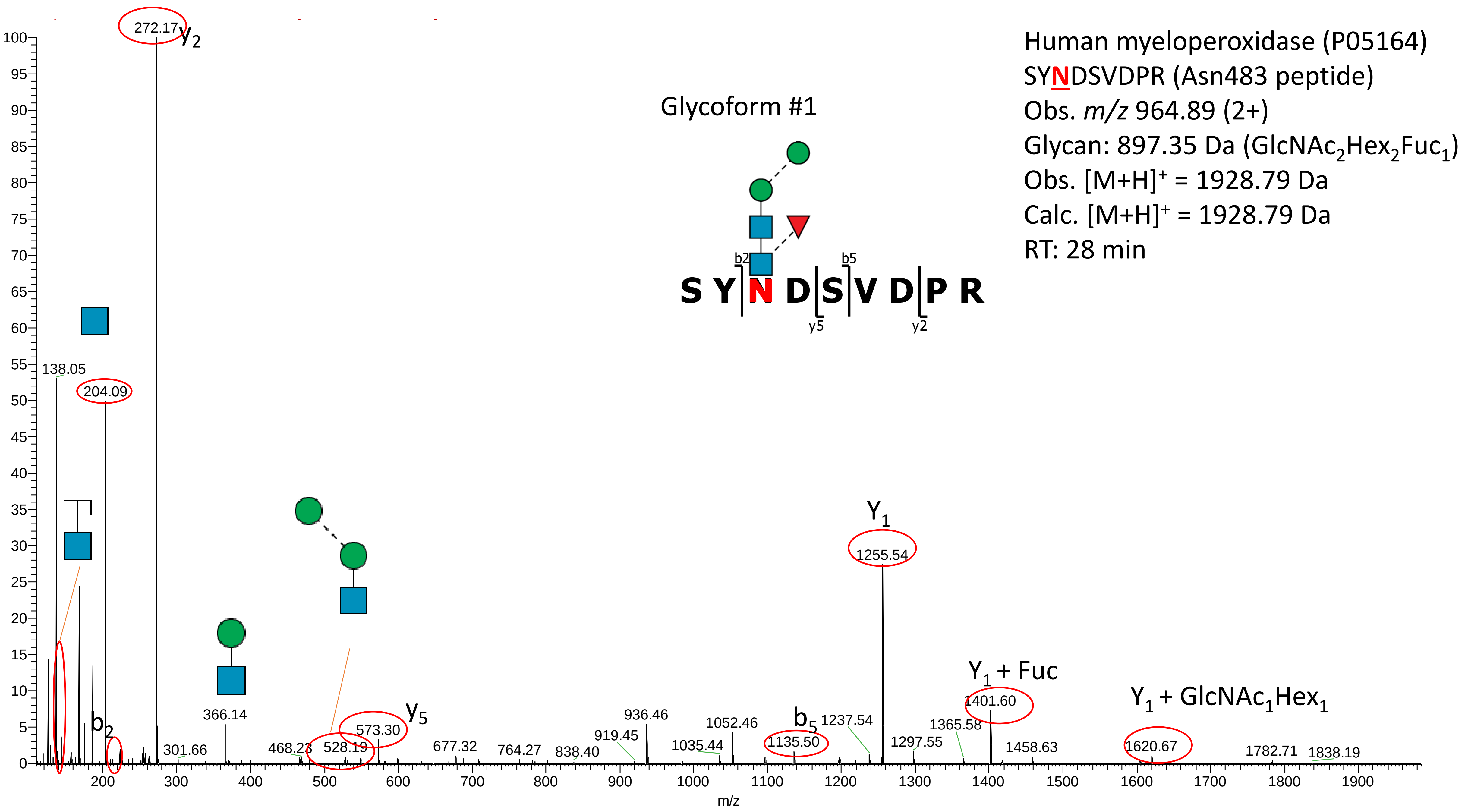

Human myeloperoxidase (P05164)
SYNDSVDPR (Asn483 peptide)
Obs. m/z 964.89 (2+)
Glycan: 897.35 Da (GlcNAc2Hex2Fuc1)
Obs. [M+H]+ = 1928.79 Da
Calc. [M+H]+ = 1928.79 Da
RT: 28 min
Glycoform #1
S Y N D S V D P R
b2
b5
y5
y2
272.17
y2
138.05
204.09
b2
301.66
366.14
468.23
528.19
573.30
y5
677.32
764.27
838.40
919.45
936.46
1035.44
1052.46
1135.50
b5
1237.54
Y1
1255.54
1297.55
1365.58
Y1 + Fuc
1401.60
1458.63
Y1 + GlcNAc1Hex1
1620.67
1782.71
1838.19
m/z

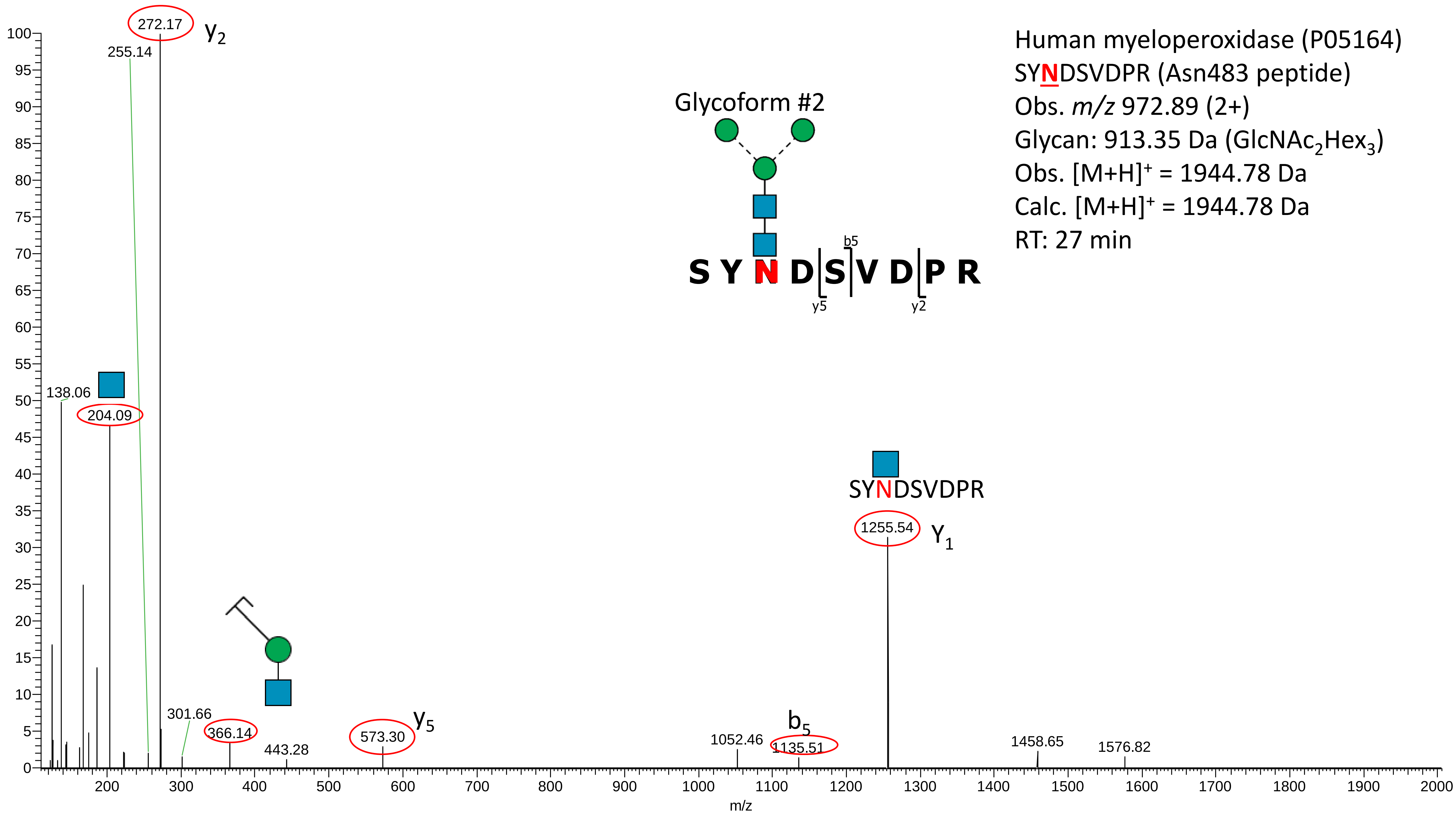

Human myeloperoxidase (P05164)
SYNDSVDPR (Asn483 peptide)
Obs. m/z 972.89 (2+)
Glycan: 913.35 Da (GlcNAc2Hex3)
Obs. [M+H]+ = 1944.78 Da
Calc. [M+H]+ = 1944.78 Da
RT: 27 min
Glycoform #2
S Y N D S V D P R
b5
y5
y2
272.17
y2
255.14
138.06
204.09
301.66
366.14
443.28
573.30
y5
1052.46
b5
1135.51
SYNDSVDPR
1255.54
Y1
1458.65
1576.82
m/z

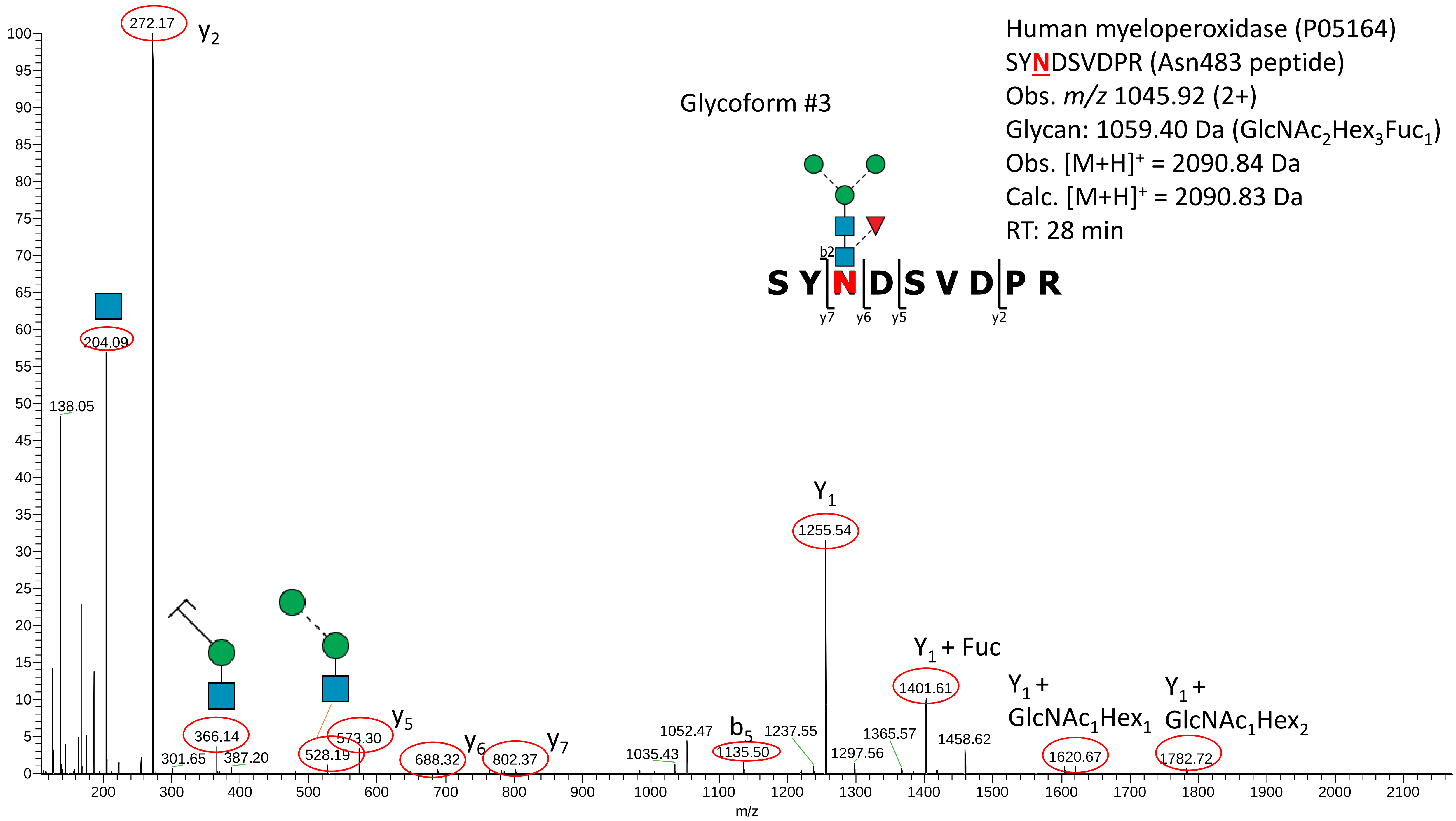

Human myeloperoxidase (P05164)
SYNDSVDPR (Asn483 peptide)
Obs. m/z 1045.92 (2+)
Glycan: 1059.40 Da (GlcNAc2Hex3Fuc1)
Obs. [M+H]+ = 2090.84 Da
Calc. [M+H]+ = 2090.83 Da
RT: 28 min
Glycoform #3
S Y N D S V D P R
b2
y7
y6
y5
y2
272.17
y2
204.09
138.05
366.14
301.65
387.20
528.19
573.30
y5
688.32
y6
802.37
y7
1035.43
1052.47
1135.50
b5
1237.55
Y1
1255.54
1297.56
1365.57
Y1 + Fuc
1401.61
1458.62
Y1 + GlcNAc1Hex1
1620.67
Y1 + GlcNAc1Hex2
1782.72
m/z

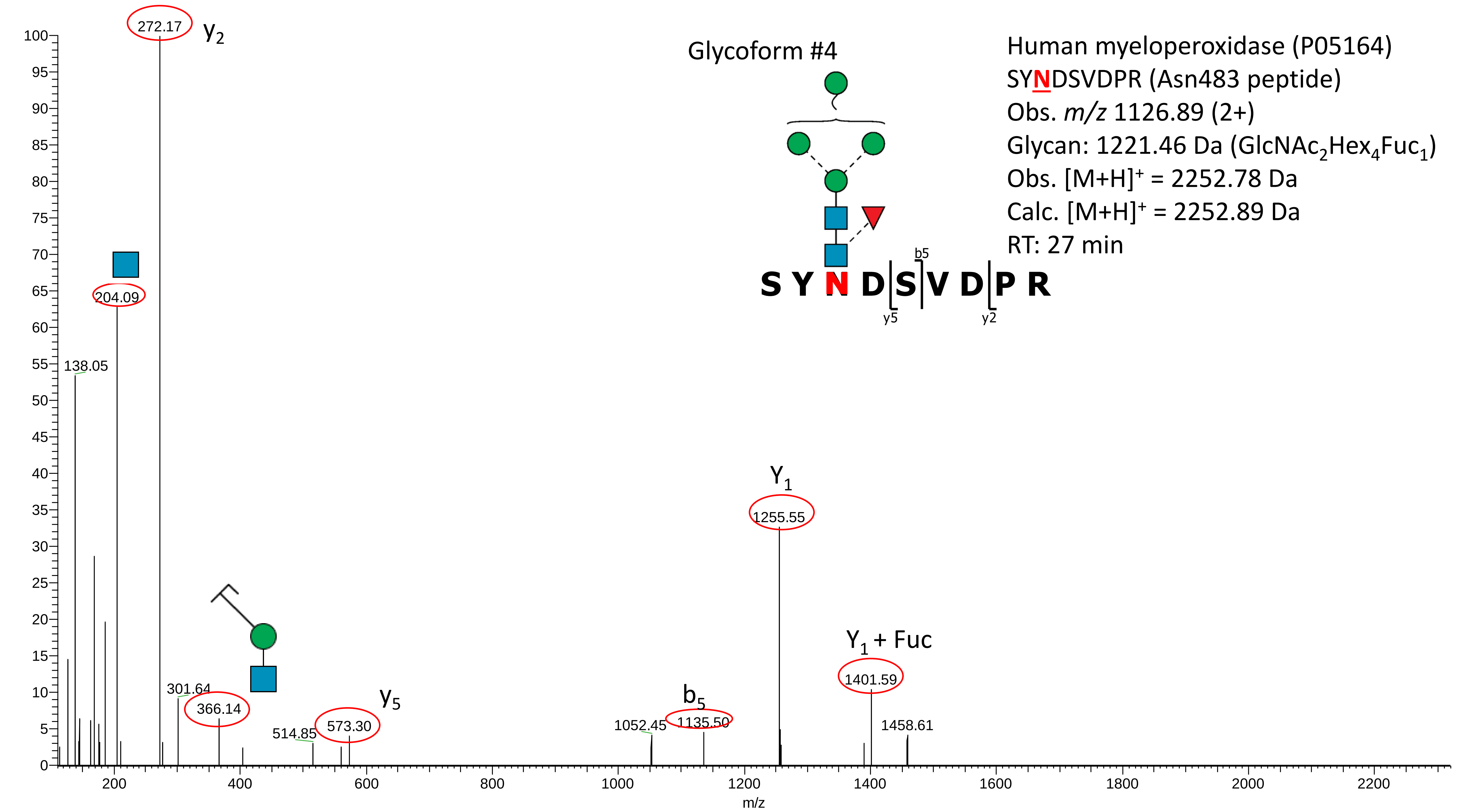
Human myeloperoxidase (P05164)
SYNDSVDPR (Asn483 peptide)
Obs. m/z 1126.89 (2+)
Glycan: 1221.46 Da (GlcNAc2Hex4Fuc1)
Obs. [M+H]+ = 2252.78 Da
Calc. [M+H]+ = 2252.89 Da
RT: 27 min
Glycoform #4
S Y N D S V D P R
b5
y5
y2
272.17
y2
204.09
138.05
301.64
366.14
514.85
573.30
y5
1052.45
1135.50
b5
Y1
1255.55
Y1 + Fuc
1401.59
1458.61
m/z

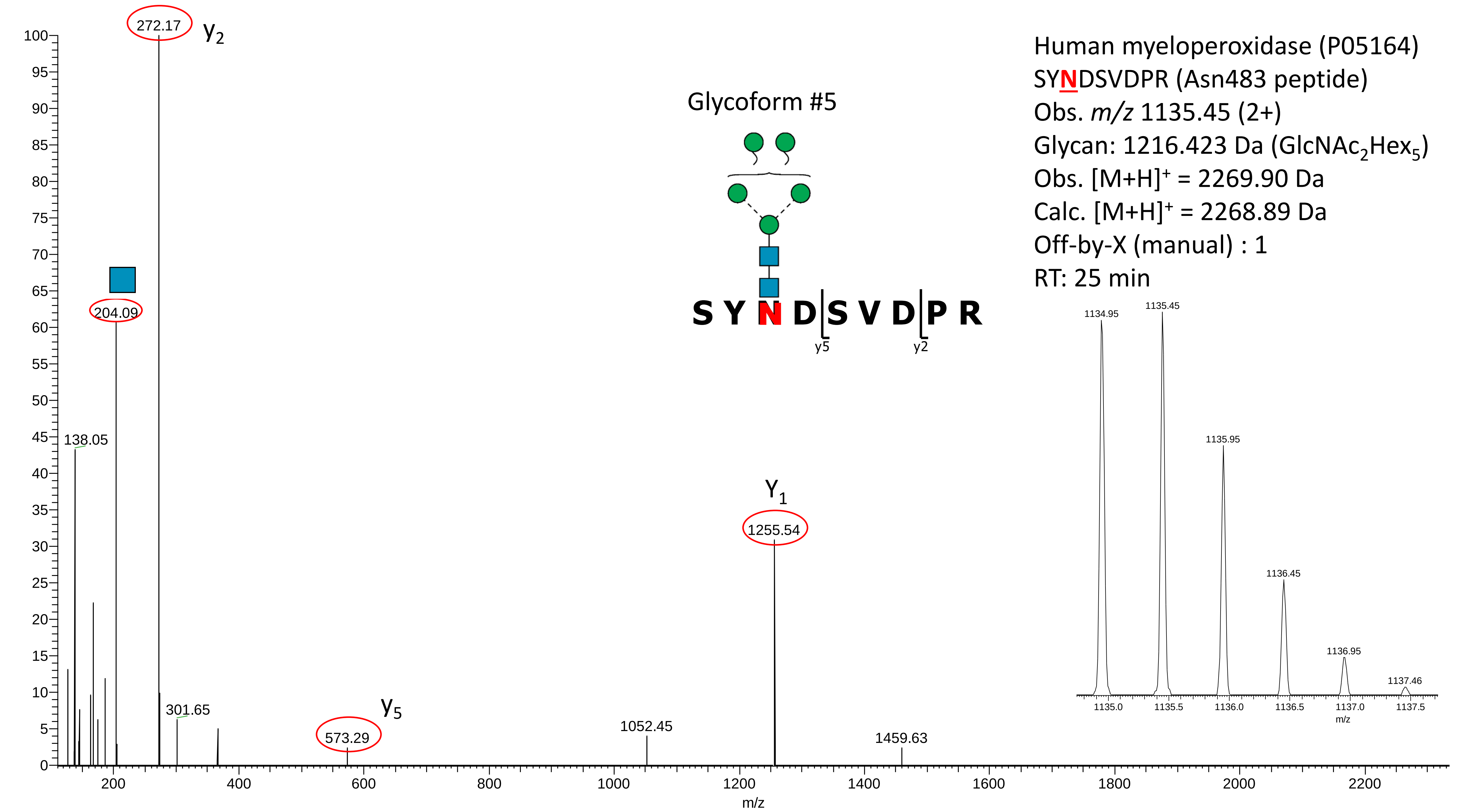
Human myeloperoxidase (P05164)
SYNDSVDPR (Asn483 peptide)
Obs. m/z 1135.45 (2+)
Glycan: 1216.423 Da (GlcNAc2Hex5)
Obs. [M+H]+ = 2269.90 Da
Calc. [M+H]+ = 2268.89 Da
Off-by-X (manual) : 1
RT: 25 min
Glycoform #5
S Y N D S V D P R
y5
y2
272.17
y2
204.09
138.05
301.65
573.29
y5
1052.45
Y1
1255.54
1459.63
1134.95
1135.45
1135.95
1136.45
1136.95
1137.46
1135.0
1135.5
1136.0
1136.5
1137.0
1137.5
m/z
m/z
0
5
10
15
20
25
30
35
40
45
50
55
60
65
70
75
80
85
90
95
100
200
400
600
800
1000
1200
1400
1600
1800
2000
2200

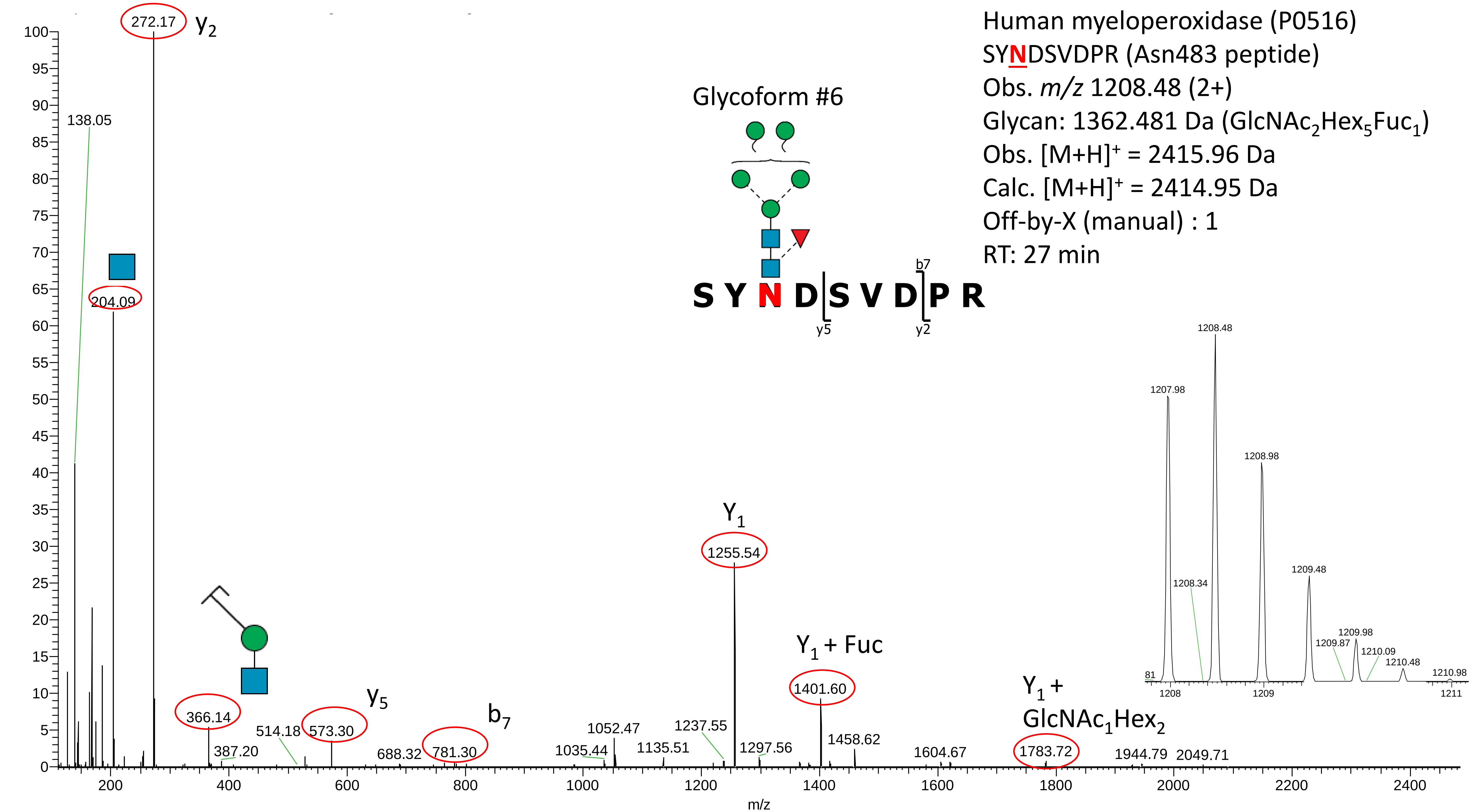
Human myeloperoxidase (P0516)
SYNDSVDPR (Asn483 peptide)
Obs. m/z 1208.48 (2+)
Glycan: 1362.481 Da (GlcNAc2Hex5Fuc1)
Obs. [M+H]+ = 2415.96 Da
Calc. [M+H]+ = 2414.95 Da
Off-by-X (manual) : 1
RT: 27 min
Glycoform #6
S Y N D S V D P R
b7
y5
y2
272.17
y2
138.05
204.09
366.14
387.20
514.18
573.30
y5
688.32
781.30
b7
1035.44
1052.47
1135.51
1237.55
Y1
1255.54
1297.56
Y1 + Fuc
1401.60
1458.62
1604.67
Y1 + GlcNAc1Hex2
1783.72
1944.79
2049.71
1207.98
1208.48
1208.34
1208.98
1209.48
1209.87
1209.98
1210.09
1210.48
1210.98
m/z

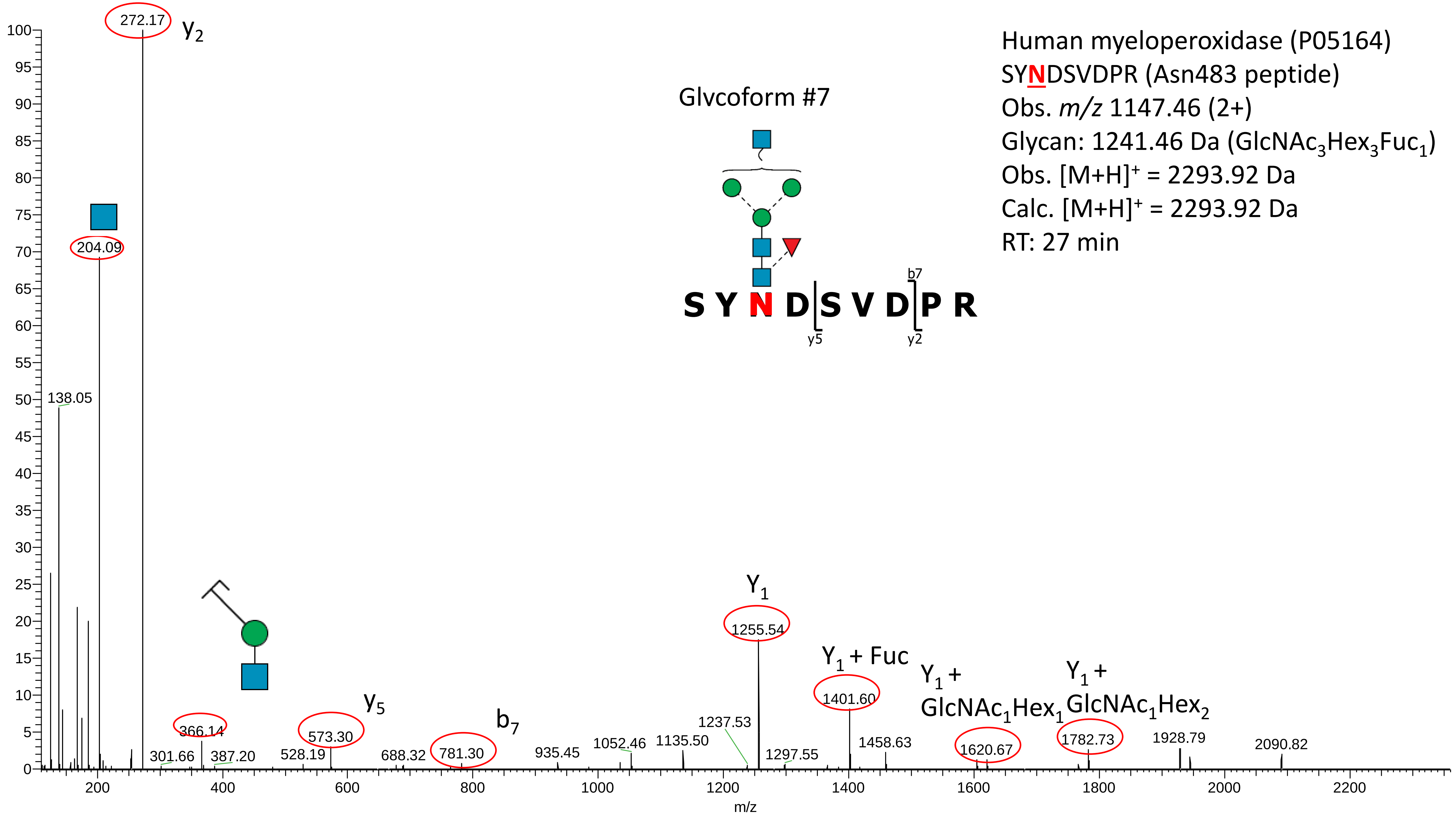
Human myeloperoxidase (P05164)
SYNDSVDPR (Asn483 peptide)
Obs. m/z 1147.46 (2+)
Glycan: 1241.46 Da (GlcNAc3Hex3Fuc1)
Obs. [M+H]+ = 2293.92 Da
Calc. [M+H]+ = 2293.92 Da
RT: 27 min
Glvcoform #7
S Y N D S V D P R
b7
y5
y2
272.17
y2
204.09
138.05
Y1
1255.54
Y1 + Fuc
1401.60
Y1 + GlcNAc1Hex1
1620.67
Y1 + GlcNAc1Hex2
1782.73
y5
573.30
b7
781.30
366.14
301.66
387.20
528.19
688.32
935.45
1052.46
1135.50
1237.53
1297.55
1458.63
1928.79
2090.82
m/z

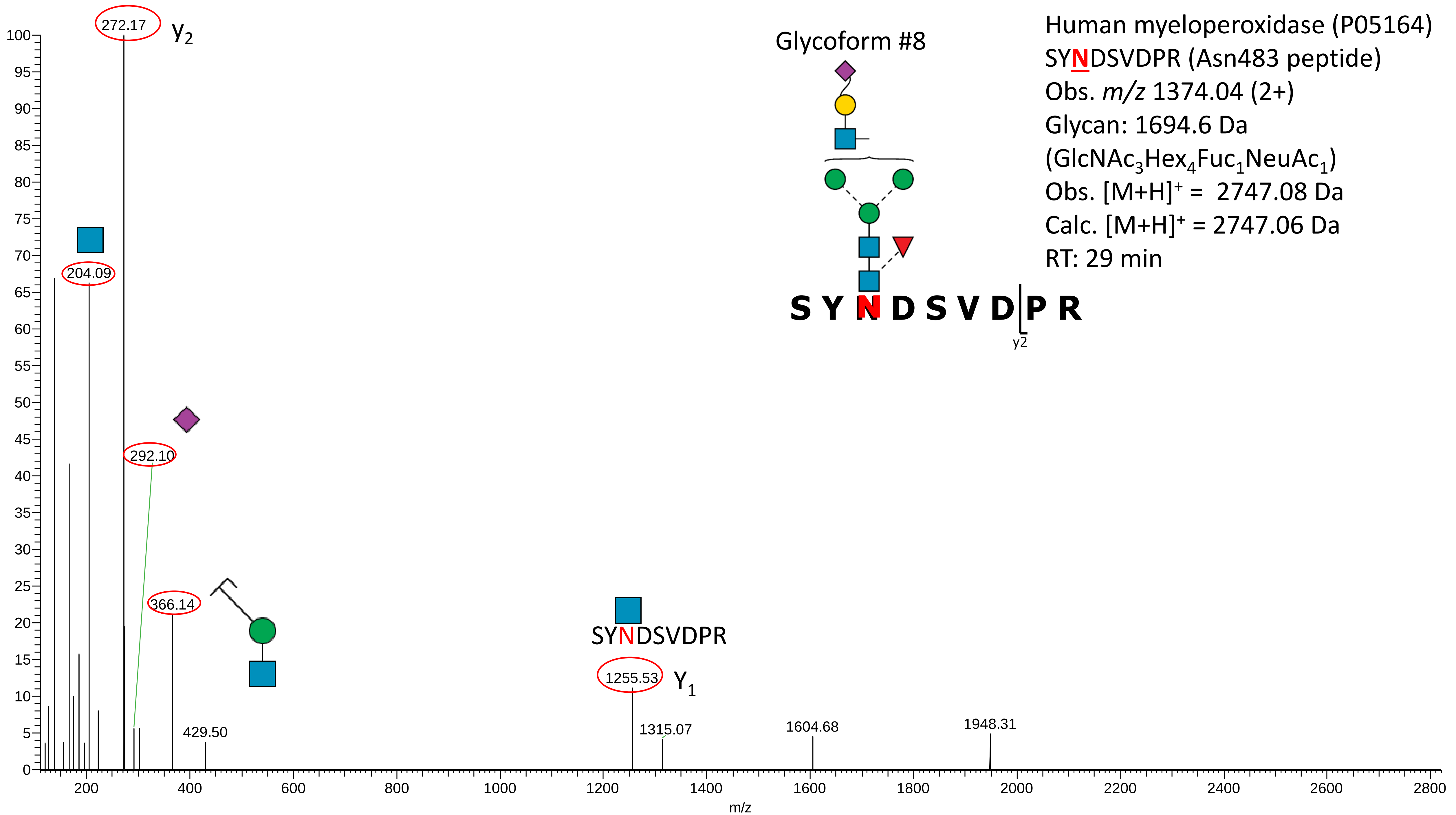

Human myeloperoxidase (P05164)
SYNDSVDPR (Asn483 peptide)
Obs. m/z 1374.04 (2+)
Glycan: 1694.6 Da
(GlcNAc3Hex4Fuc1NeuAc1)
Obs. [M+H]+ = 2747.08 Da
Calc. [M+H]+ = 2747.06 Da
RT: 29 min
Glycoform #8
S Y N D S V D P R
y2
272.17
y2
204.09
292.10
366.14
429.50
SYNDSVDPR
1255.53
Y1
1315.07
1604.68
1948.31
m/z

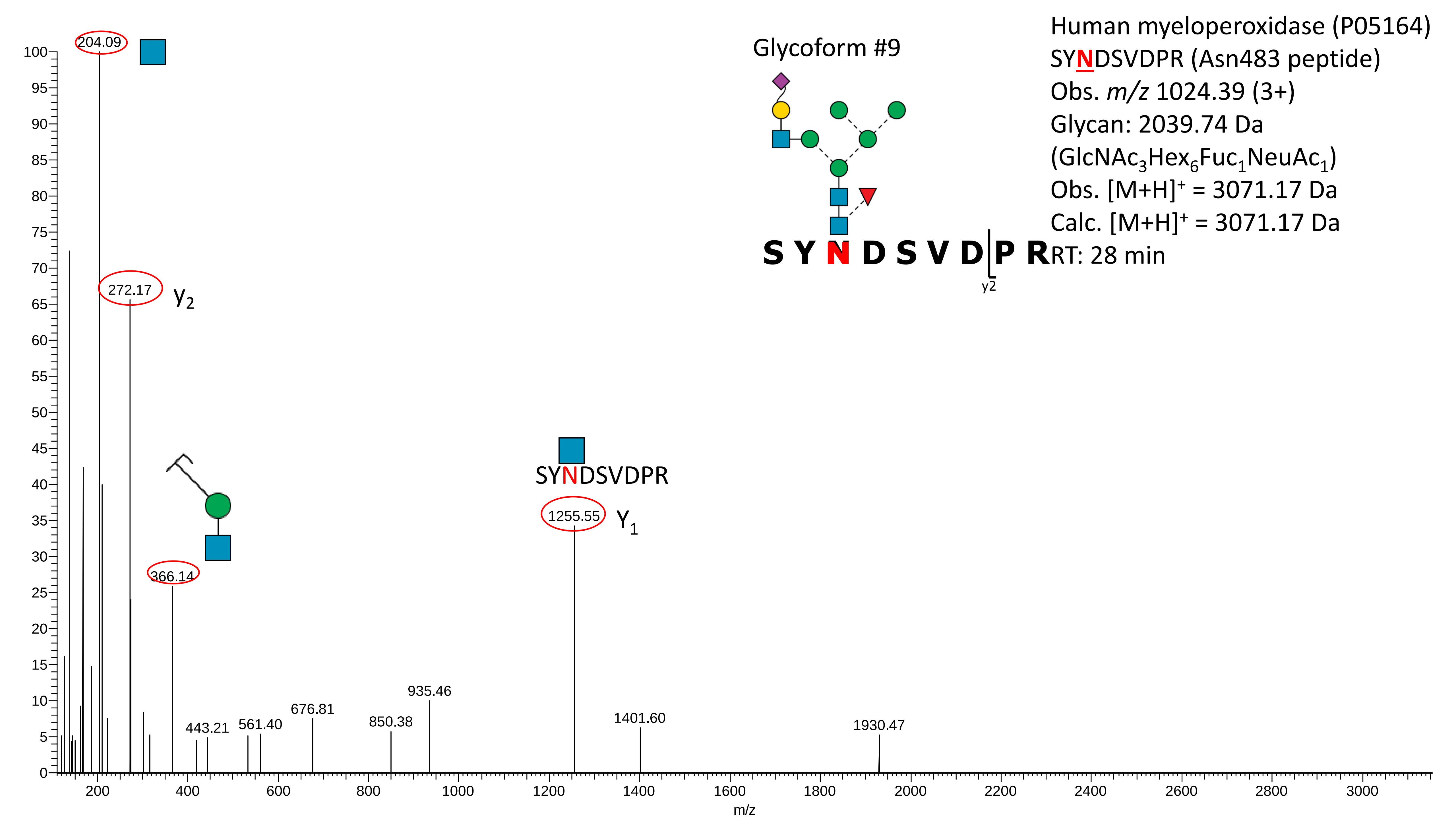

Human myeloperoxidase (P05164)
SYNDSVDPR (Asn483 peptide)
Obs. m/z 1024.39 (3+)
Glycan: 2039.74 Da
(GlcNAc3Hex6Fuc1NeuAc1)
Obs. [M+H]+ = 3071.17 Da
Calc. [M+H]+ = 3071.17 Da
RT: 28 min
Glycoform #9
S Y N D S V D P R
y2
204.09
272.17
y2
366.14
1255.55
Y1
SYNDSVDPR
443.21
561.40
676.81
850.38
935.46
1401.60
1930.47
m/z

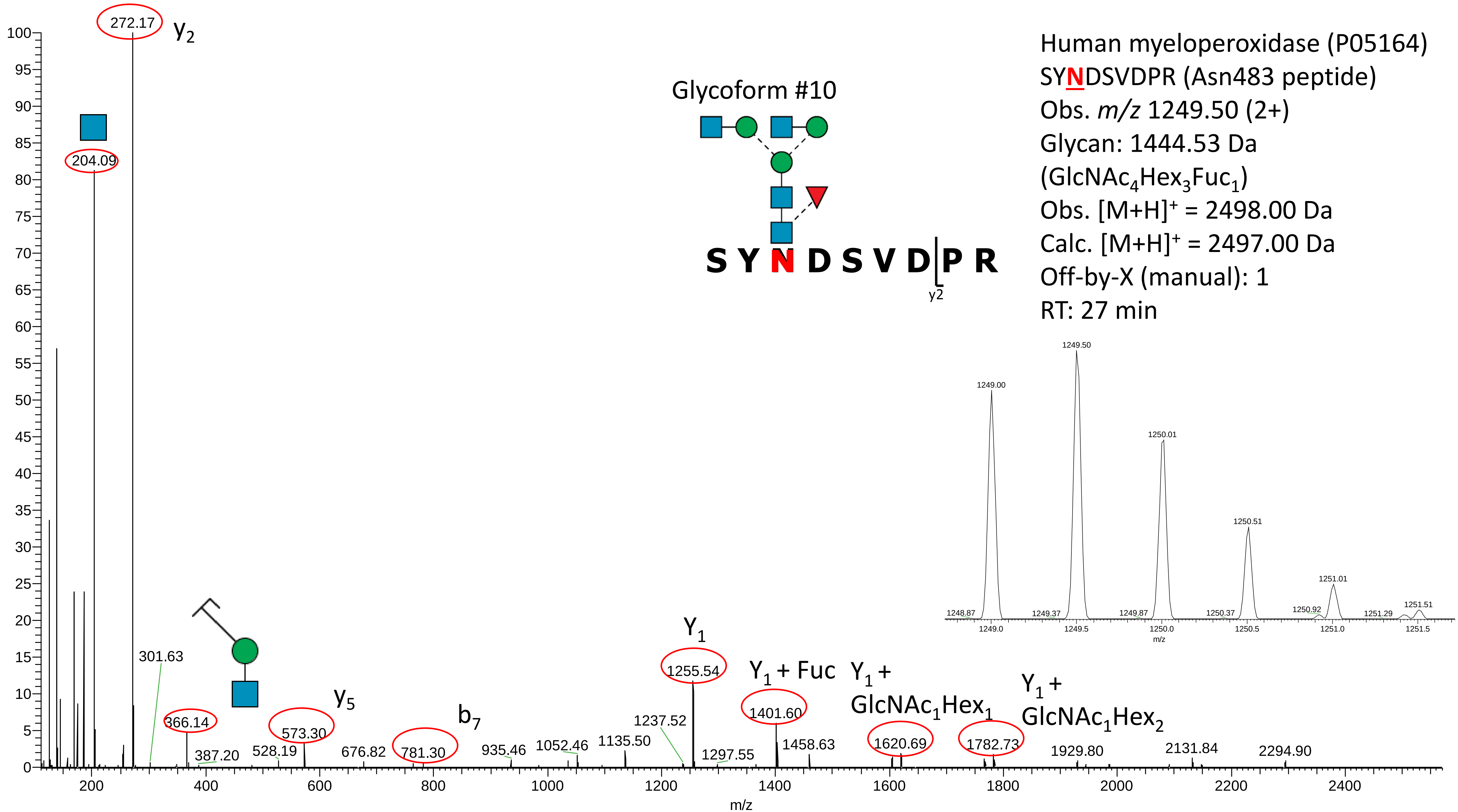

Human myeloperoxidase (P05164)
SYNDSVDPR (Asn483 peptide)
Obs. m/z 1249.50 (2+)
Glycan: 1444.53 Da
(GlcNAc4Hex3Fuc1)
Obs. [M+H]+ = 2498.00 Da
Calc. [M+H]+ = 2497.00 Da
Off-by-X (manual): 1
RT: 27 min
Glycoform #10
S Y N D S V D P R
y2
272.17
204.09
301.63
366.14
387.20
528.19
573.30
676.82
781.30
935.46
1052.46
1135.50
1237.52
1255.54
1297.55
1401.60
1458.63
1620.69
1782.73
1929.80
2131.84
2294.90
y5
b7
Y1
Y1 + Fuc
Y1 + GlcNAc1Hex1
Y1 + GlcNAc1Hex2
m/z
1249.00
1249.50
1250.01
1250.51
1251.01
1251.51
1248.87
1249.37
1249.87
1250.37
1250.92
1251.29

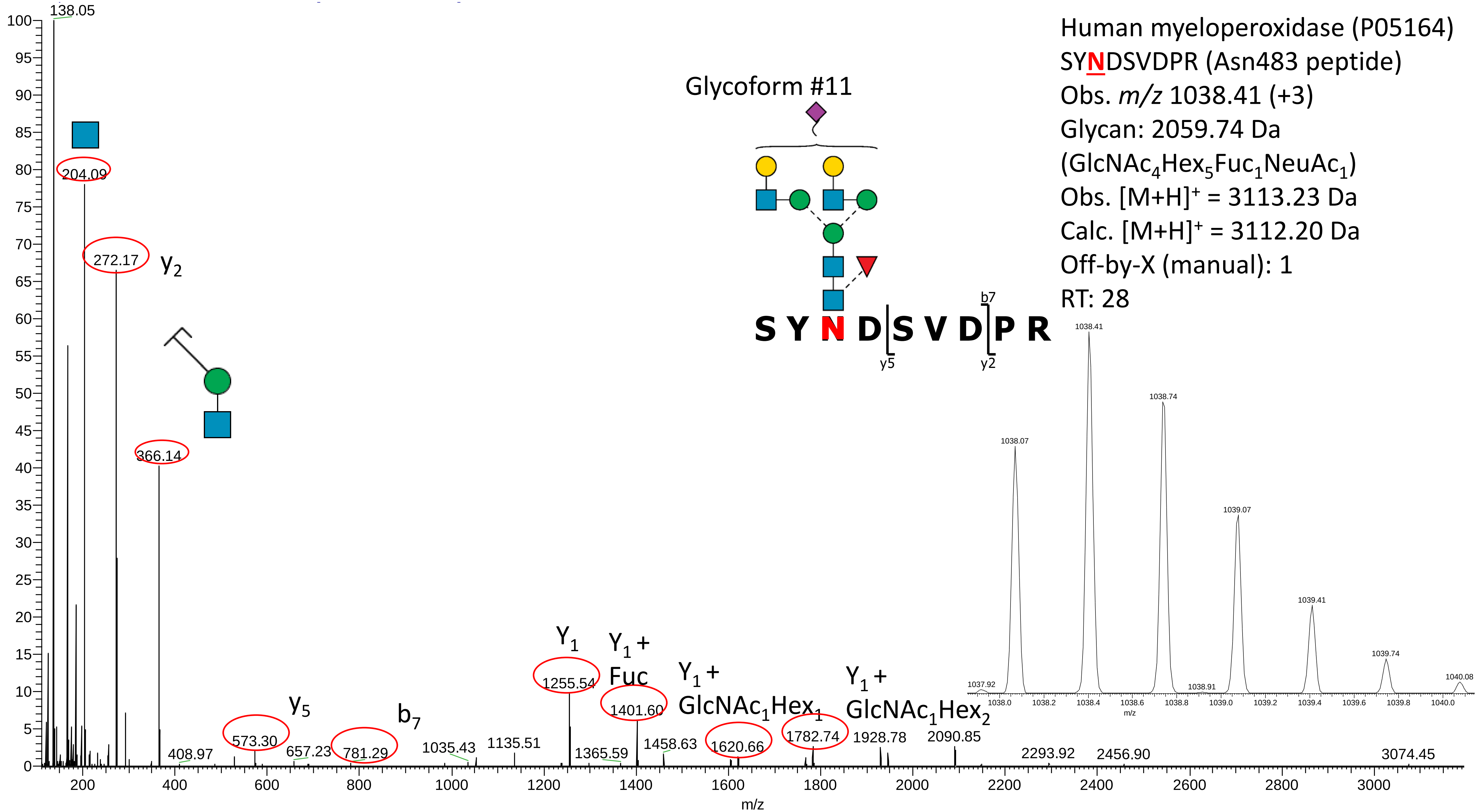
Human myeloperoxidase (P05164)
SYNDSVDPR (Asn483 peptide)
Obs. m/z 1038.41 (+3)
Glycan: 2059.74 Da
(GlcNAc4Hex5Fuc1NeuAc1)
Obs. [M+H]+ = 3113.23 Da
Calc. [M+H]+ = 3112.20 Da
Off-by-X (manual): 1
RT: 28
Glycoform #11
S Y N D S V D P R
b7
y5
y2
138.05
204.09
272.17
y2
366.14
408.97
573.30
y5
657.23
781.29
b7
1035.43
1135.51
Y1
1255.54
Y1 + Fuc
1401.60
1365.59
1458.63
Y1 + GlcNAc1Hex1
1620.66
1782.74
Y1 + GlcNAc1Hex2
1928.78
2090.85
2293.92
2456.90
3074.45
m/z
1037.92
1038.07
1038.41
1038.74
1038.91
1039.07
1039.41
1039.74
1040.08
m/z

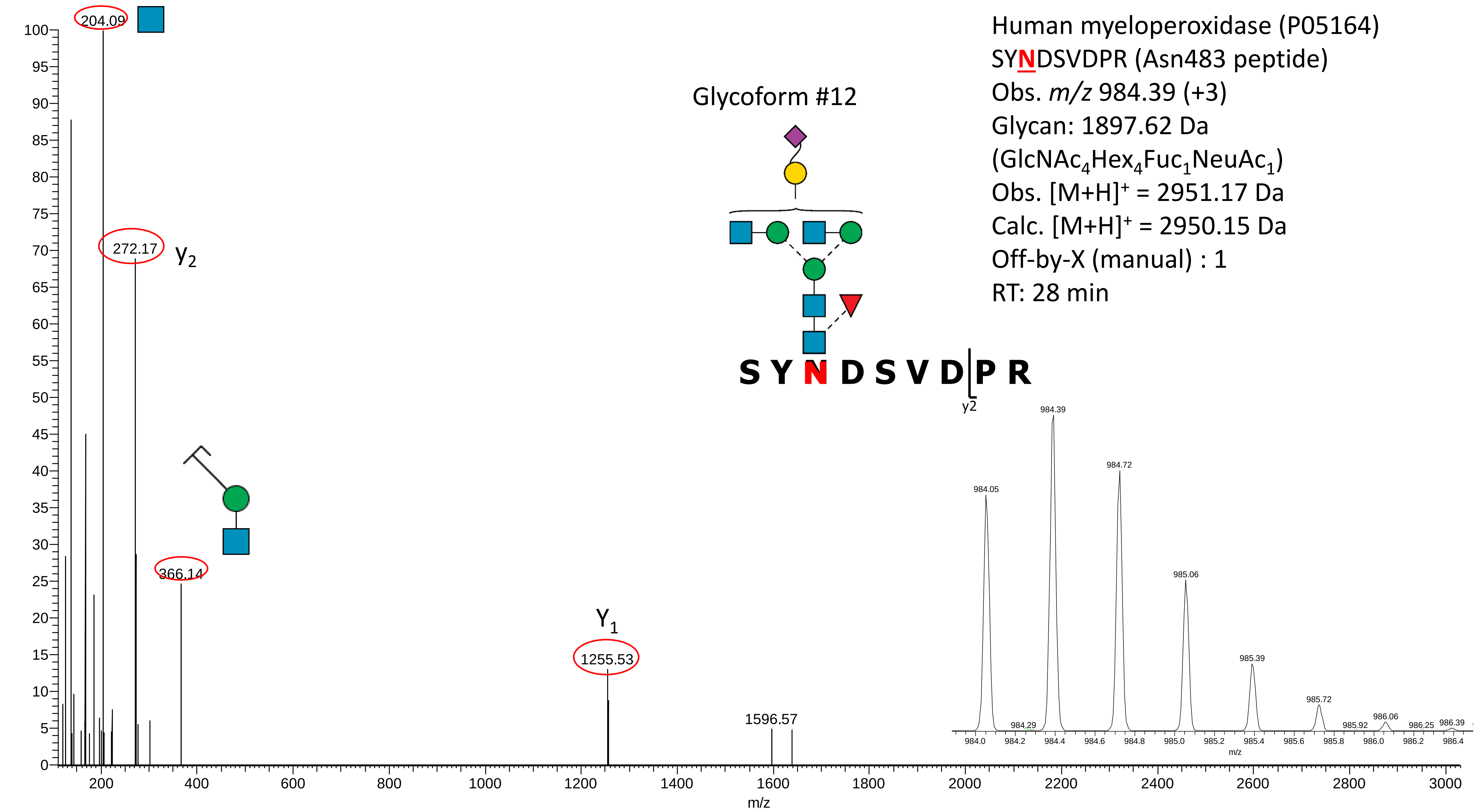
Human myeloperoxidase (P05164)
SYNDSVDPR (Asn483 peptide)
Obs. m/z 984.39 (+3)
Glycan: 1897.62 Da
(GlcNAc4Hex4Fuc1NeuAc1)
Obs. [M+H]+ = 2951.17 Da
Calc. [M+H]+ = 2950.15 Da
Off-by-X (manual) : 1
RT: 28 min
Glycoform #12
S Y N D S V D P R
y2
204.09
272.17
y2
366.14
Y1
1255.53
1596.57
984.05
984.29
984.39
984.72
985.06
985.39
985.72
985.92
986.06
986.25
986.39
m/z

**Supplementary Data S3E**

Manually annotated reversed phase-LC-ESI-HCD-MS/MS (+) spectra of all glycosylated and non-glycosylated peptides containing Asn729 identified from the analysis of unenriched peptide mixtures of nMPO.

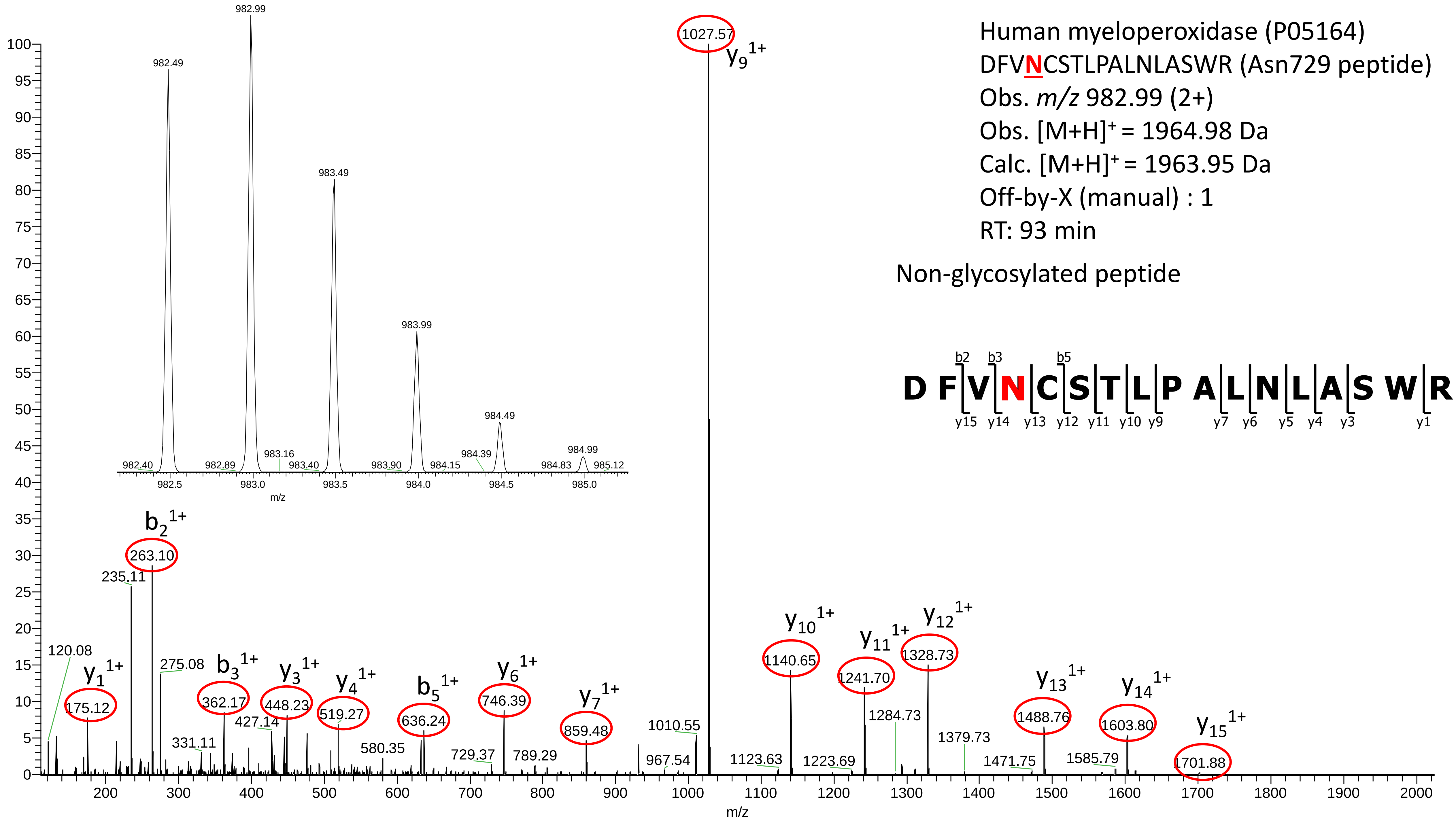

Human myeloperoxidase (P05164)
DFVNCSTLPALNLASWR (Asn729 peptide)
Obs. m/z 982.99 (2+)
Obs. [M+H]+ = 1964.98 Da
Calc. [M+H]+ = 1963.95 Da
Off-by-X (manual) : 1
RT: 93 min
Non-glycosylated peptide
D F V N C S T L P A L N L A S W R
b2 b3 b5
y15 y14 y13 y12 y11 y10 y9 y7 y6 y5 y4 y3 y1
1027.57 y9 1+
263.10 b2 1+
235.11
120.08
175.12 y1 1+
275.08
362.17 b3 1+
448.23 y3 1+
519.27 y4 1+
636.24 b5 1+
746.39 y6 1+
859.48 y7 1+
1140.65 y10 1+
1241.70 y11 1+
1328.73 y12 1+
1488.76 y13 1+
1603.80 y14 1+
1701.88 y15 1+
m/z

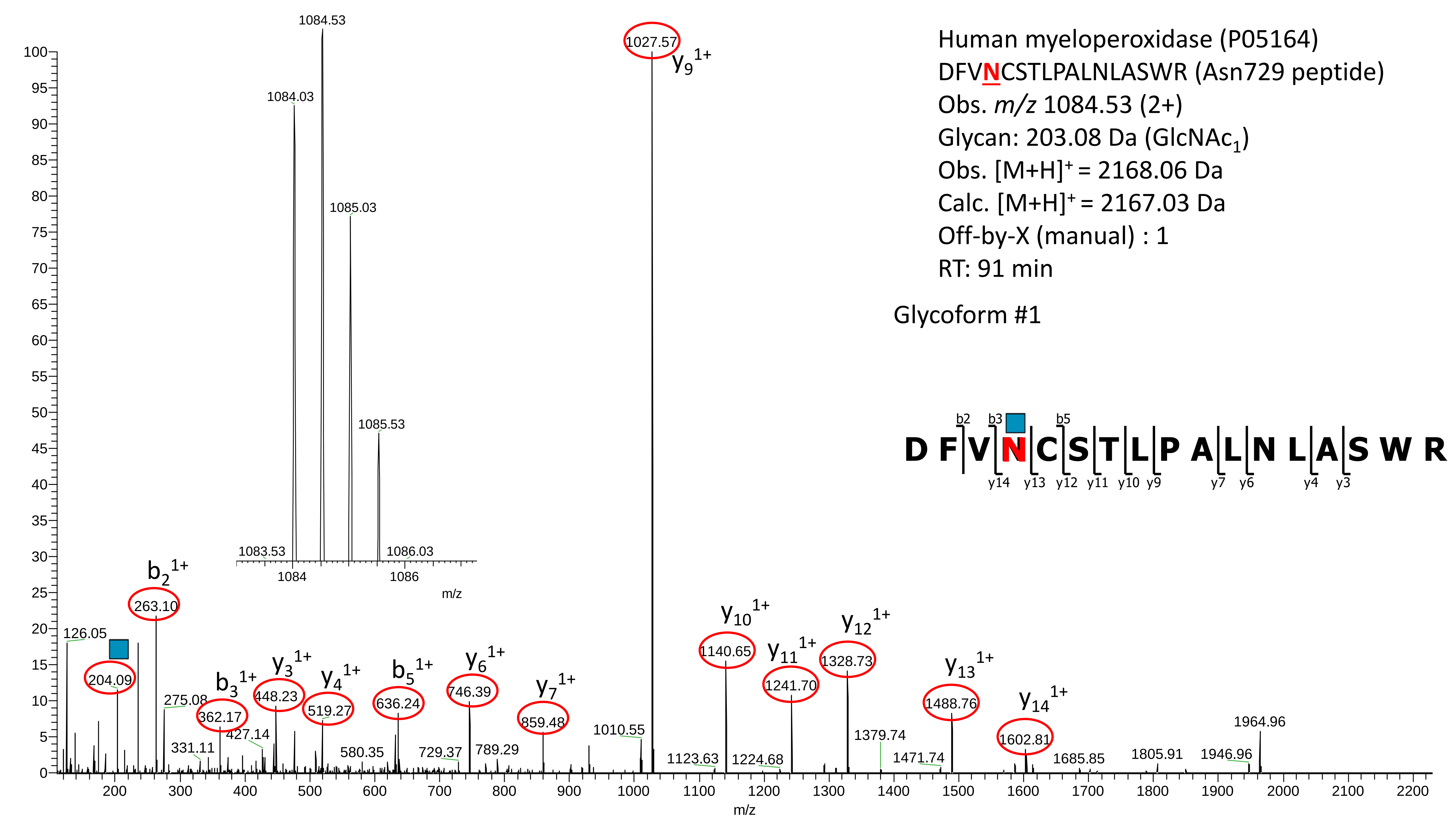
Human myeloperoxidase (P05164)
DFVNCSTLPALNLASWR (Asn729 peptide)
Obs. m/z 1084.53 (2+)
Glycan: 203.08 Da (GlcNAc1)
Obs. [M+H]+ = 2168.06 Da
Calc. [M+H]+ = 2167.03 Da
Off-by-X (manual) : 1
RT: 91 min
Glycoform #1
D F V N C S T L P A L N L A S W R
b2 b3 b5
y14 y13 y12 y11 y10 y9 y7 y6 y4 y3
1084.53
1084.03
1085.03
1085.53
1083.53
1086.03
1084
1086
m/z
1027.57
y9 1+
b2 1+
263.10
126.05
204.09
275.08
b3 1+
362.17
331.11
427.14
y3 1+
448.23
y4 1+
519.27
580.35
b5 1+
636.24
y6 1+
746.39
729.37
789.29
y7 1+
859.48
1010.55
y10 1+
1140.65
1123.63
y11 1+
1241.70
1224.68
y12 1+
1328.73
1379.74
1471.74
y13 1+
1488.76
y14 1+
1602.81
1685.85
1805.91
1946.96
1964.96
m/z

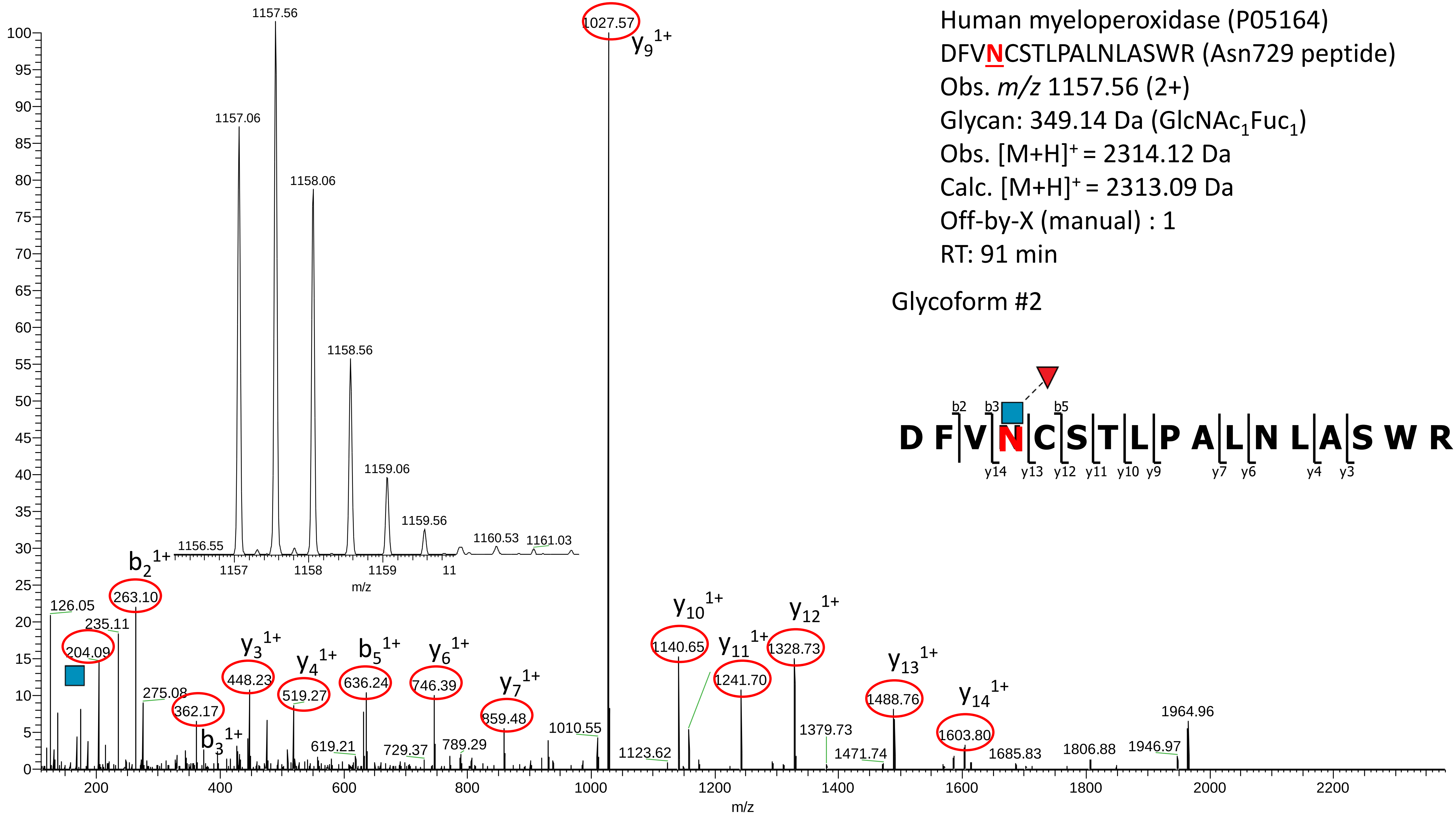

Human myeloperoxidase (P05164)
DFVNCSTLPALNLASWR (Asn729 peptide)
Obs. m/z 1157.56 (2+)
Glycan: 349.14 Da (GlcNAc1Fuc1)
Obs. [M+H]+ = 2314.12 Da
Calc. [M+H]+ = 2313.09 Da
Off-by-X (manual) : 1
RT: 91 min
Glycoform #2
D F V N C S T L P A L N L A S W R
b2 b3 b5
y14 y13 y12 y11 y10 y9 y7 y6 y4 y3
1157.56
1157.06
1158.06
1158.56
1159.06
1159.56
1156.55
1160.53
1161.03
1157
1158
1159
11
m/z
1027.57
y9 1+
b2 1+
263.10
126.05
235.11
204.09
275.08
362.17
b3 1+
y3 1+
448.23
y4 1+
519.27
619.21
b5 1+
636.24
729.37
y6 1+
746.39
789.29
y7 1+
859.48
1010.55
1123.62
y10 1+
1140.65
y11 1+
1241.70
y12 1+
1328.73
1379.73
1471.74
y13 1+
1488.76
y14 1+
1603.80
1685.83
1806.88
1946.97
1964.96
m/z

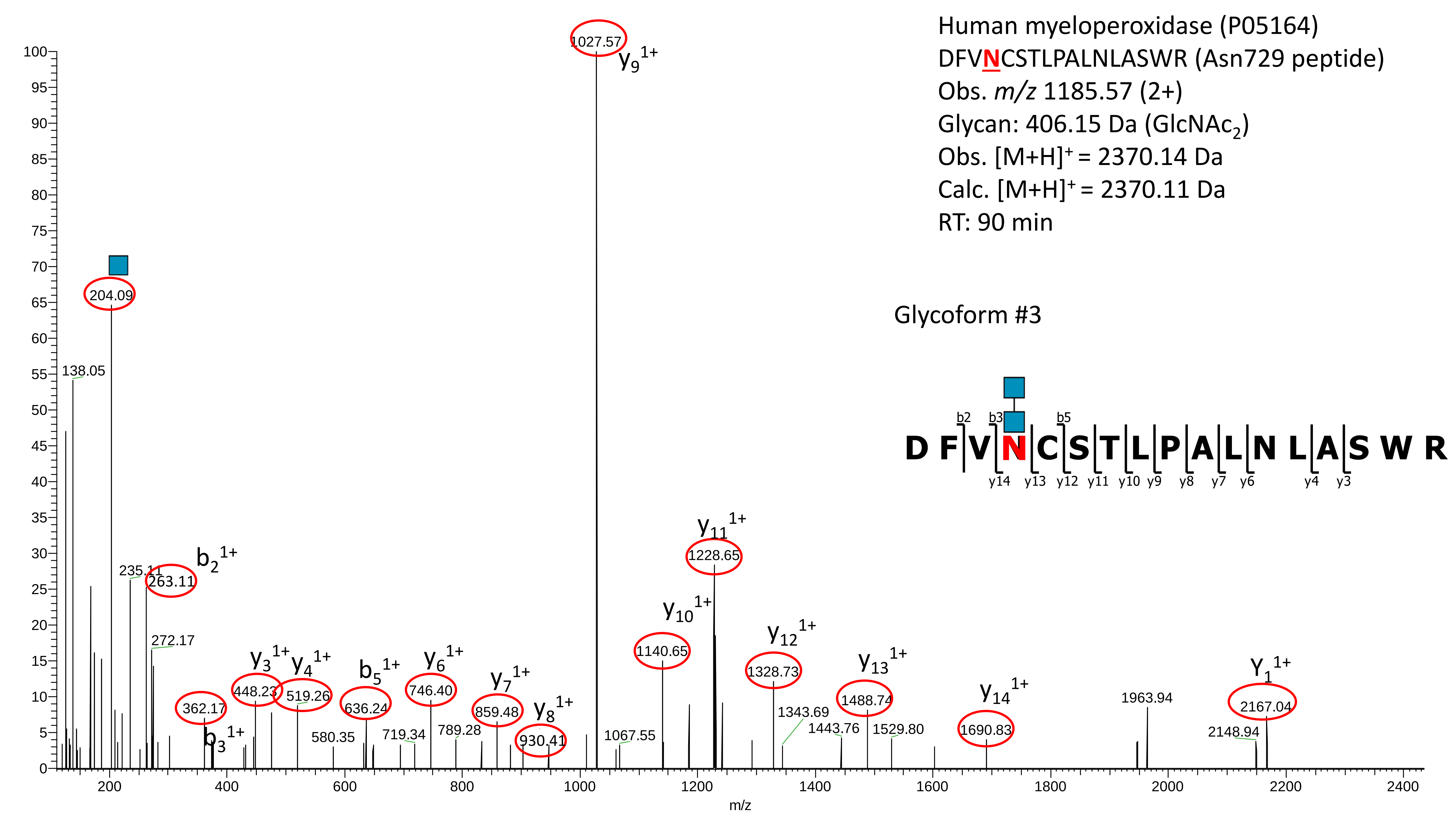

Human myeloperoxidase (P05164)
DFVNCSTLPALNLASWR (Asn729 peptide)
Obs. m/z 1185.57 (2+)
Glycan: 406.15 Da (GlcNAc2)
Obs. [M+H]+ = 2370.14 Da
Calc. [M+H]+ = 2370.11 Da
RT: 90 min
Glycoform #3
D F V N C S T L P A L N L A S W R
b2 b3 b5
y14 y13 y12 y11 y10 y9 y8 y7 y6 y4 y3
1027.57
y9 1+
204.09
138.05
235.11
263.11
b2 1+
272.17
362.17
b3 1+
448.23
y3 1+
519.26
y4 1+
580.35
636.24
b5 1+
719.34
746.40
y6 1+
789.28
859.48
y7 1+
930.41
y8 1+
1067.55
1140.65
y10 1+
1228.65
y11 1+
1328.73
y12 1+
1343.69
1443.76
1488.74
y13 1+
1529.80
1690.83
y14 1+
1963.94
2148.94
2167.04
Y1 1+
m/z

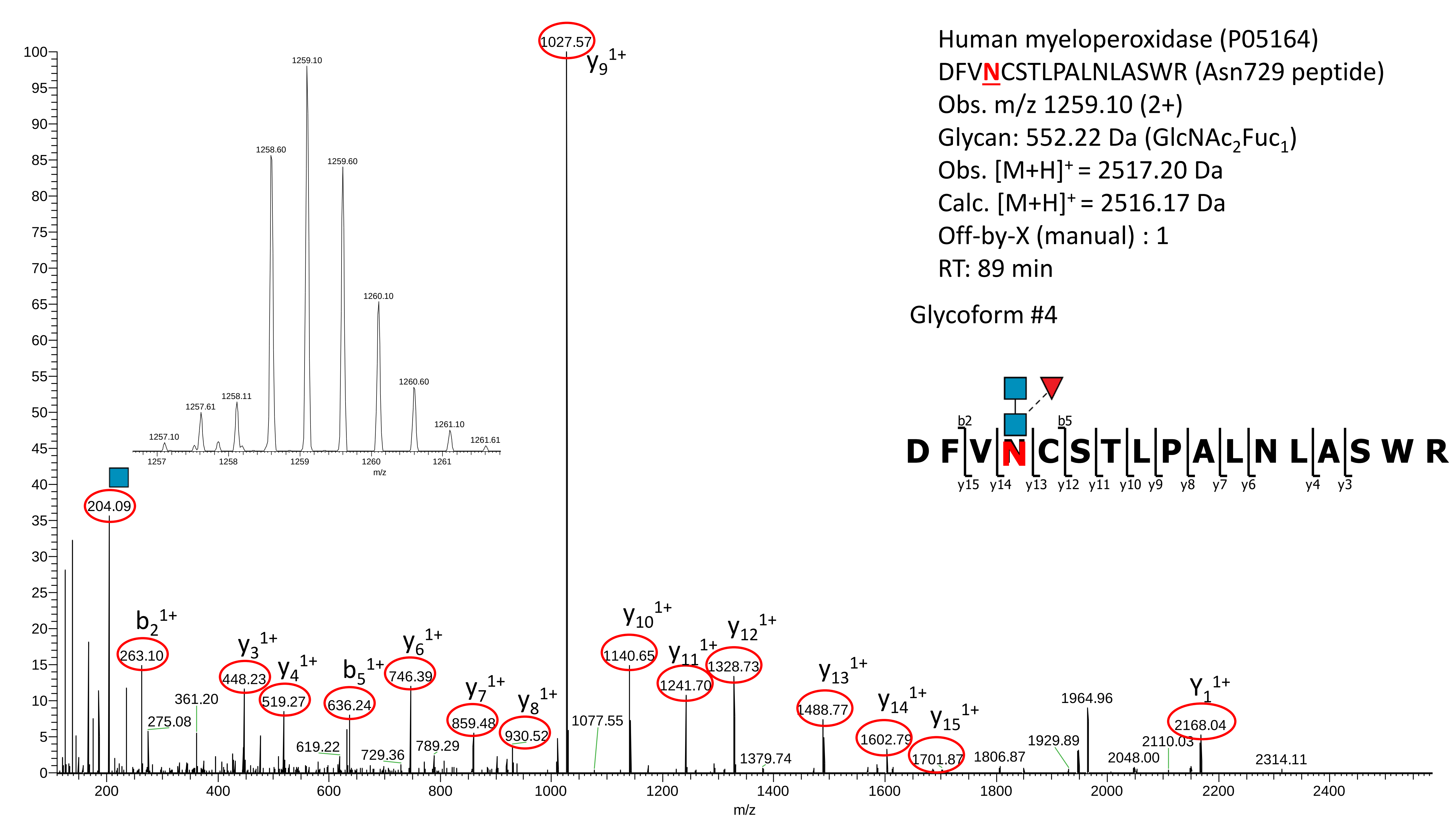

Human myeloperoxidase (P05164)
DFVNCSTLPALNLASWR (Asn729 peptide)
Obs. m/z 1259.10 (2+)
Glycan: 552.22 Da (GlcNAc2Fuc1)
Obs. [M+H]+ = 2517.20 Da
Calc. [M+H]+ = 2516.17 Da
Off-by-X (manual) : 1
RT: 89 min
Glycoform #4
D F V N C S T L P A L N L A S W R
b2 b5
y15 y14 y13 y12 y11 y10 y9 y8 y7 y6 y4 y3
1027.57 y9 1+
204.09
263.10 b2 1+
275.08
361.20
448.23 y3 1+
519.27 y4 1+
619.22
636.24 b5 1+
729.36
746.39 y6 1+
789.29
859.48 y7 1+
930.52 y8 1+
1077.55
1140.65 y10 1+
1241.70 y11 1+
1328.73 y12 1+
1379.74
1488.77 y13 1+
1602.79 y14 1+
1701.87 y15 1+
1806.87
1929.89
1964.96
2048.00
2110.03
2168.04 Y1 1+
2314.11
1257.10
1257.61
1258.11
1258.60
1259.10
1259.60
1260.10
1260.60
1261.10
1261.61
m/z

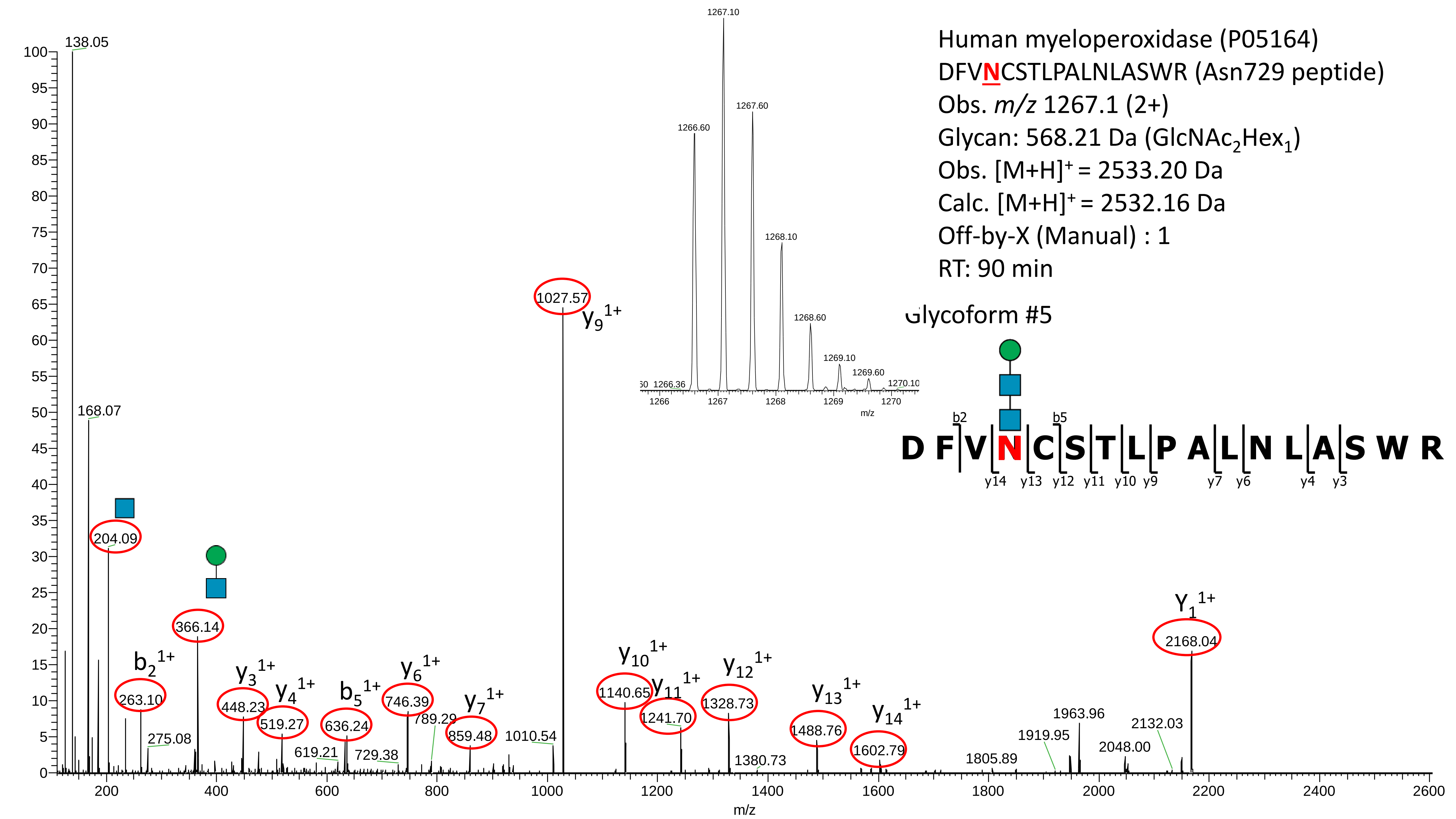
Human myeloperoxidase (P05164)
DFVNCSTLPALNLASWR (Asn729 peptide)
Obs. m/z 1267.1 (2+)
Glycan: 568.21 Da (GlcNAc2Hex1)
Obs. [M+H]+ = 2533.20 Da
Calc. [M+H]+ = 2532.16 Da
Off-by-X (Manual) : 1
RT: 90 min
Glycoform #5
D F V N C S T L P A L N L A S W R
b2 b5
y14 y13 y12 y11 y10 y9 y7 y6 y4 y3
138.05
168.07
204.09
366.14
263.10
275.08
448.23
519.27
619.21
636.24
729.38
746.39
789.29
859.48
1010.54
1027.57
1140.65
1241.70
1328.73
1380.73
1488.76
1602.79
1805.89
1919.95
1963.96
2048.00
2132.03
2168.04
b2 1+
y3 1+
y4 1+
b5 1+
y6 1+
y7 1+
y9 1+
y10 1+
y11 1+
y12 1+
y13 1+
y14 1+
Y1 1+
m/z

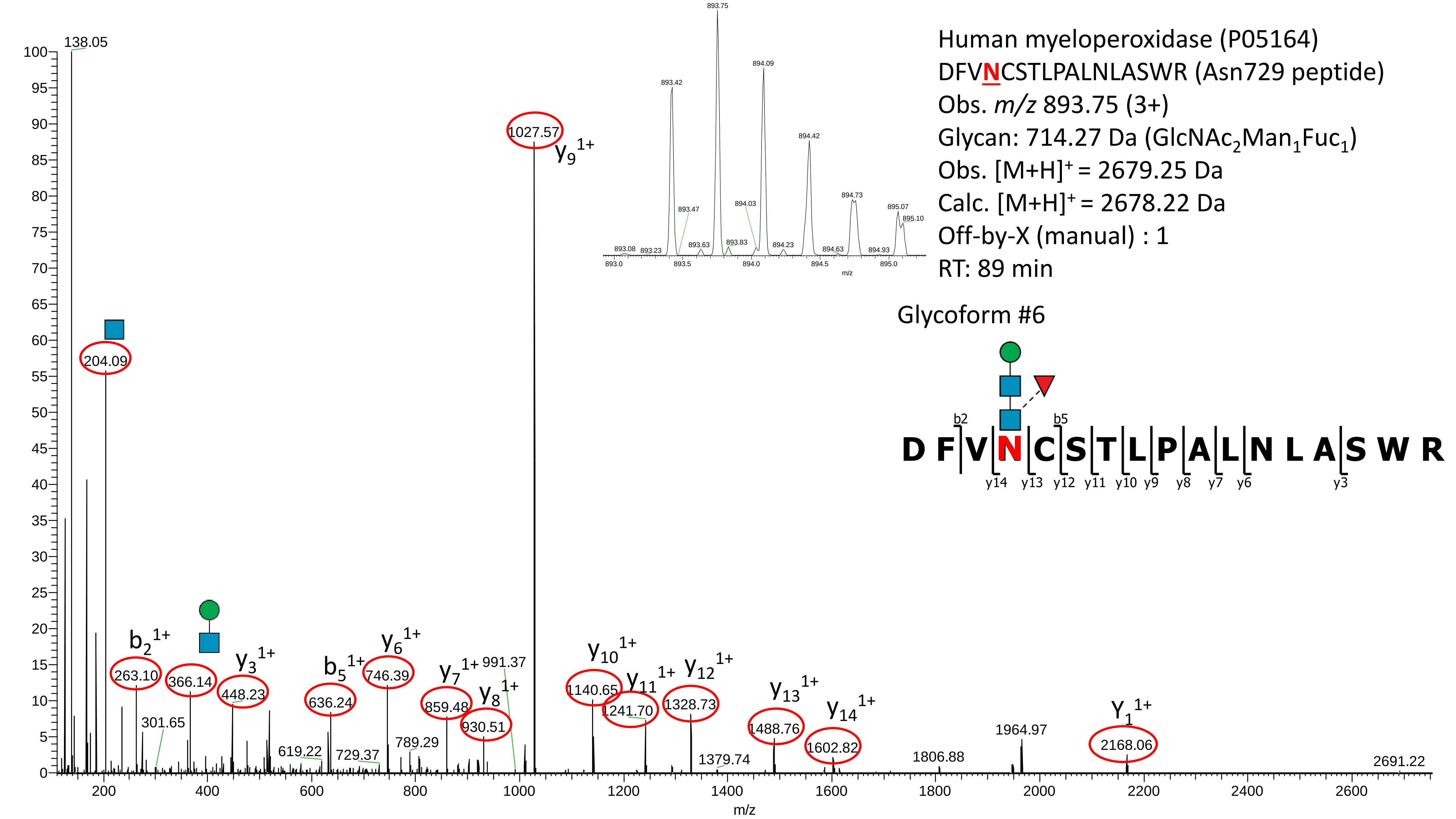
Human myeloperoxidase (P05164)
DFVNCSTLPALNLASWR (Asn729 peptide)
Obs. m/z 893.75 (3+)
Glycan: 714.27 Da (GlcNAc2Man1Fuc1)
Obs. [M+H]+ = 2679.25 Da
Calc. [M+H]+ = 2678.22 Da
Off-by-X (manual) : 1
RT: 89 min
Glycoform #6
D F V N C S T L P A L N L A S W R
b2
b5
y14 y13 y12 y11 y10 y9 y8 y7 y6 y3
893.75
893.42
894.09
894.42
894.73
895.07
895.10
893.47
894.03
893.08
893.23
893.63
893.83
894.23
894.63
894.93
893.0
893.5
894.0
894.5
895.0
m/z
138.05
204.09
1027.57
y9 1+
b2 1+
263.10
366.14
y3 1+
448.23
301.65
b5 1+
636.24
y6 1+
746.39
619.22
729.37
789.29
y7 1+
859.48
y8 1+
930.51
991.37
y10 1+
1140.65
y11 1+
1241.70
y12 1+
1328.73
1379.74
y13 1+
1488.76
y14 1+
1602.82
1806.88
1964.97
Y1 1+
2168.06
2691.22
100
95
90
85
80
75
70
65
60
55
50
45
40
35
30
25
20
15
10
5
0
200
400
600
800
1000
1200
1400
1600
1800
2000
2200
2400
2600
m/z

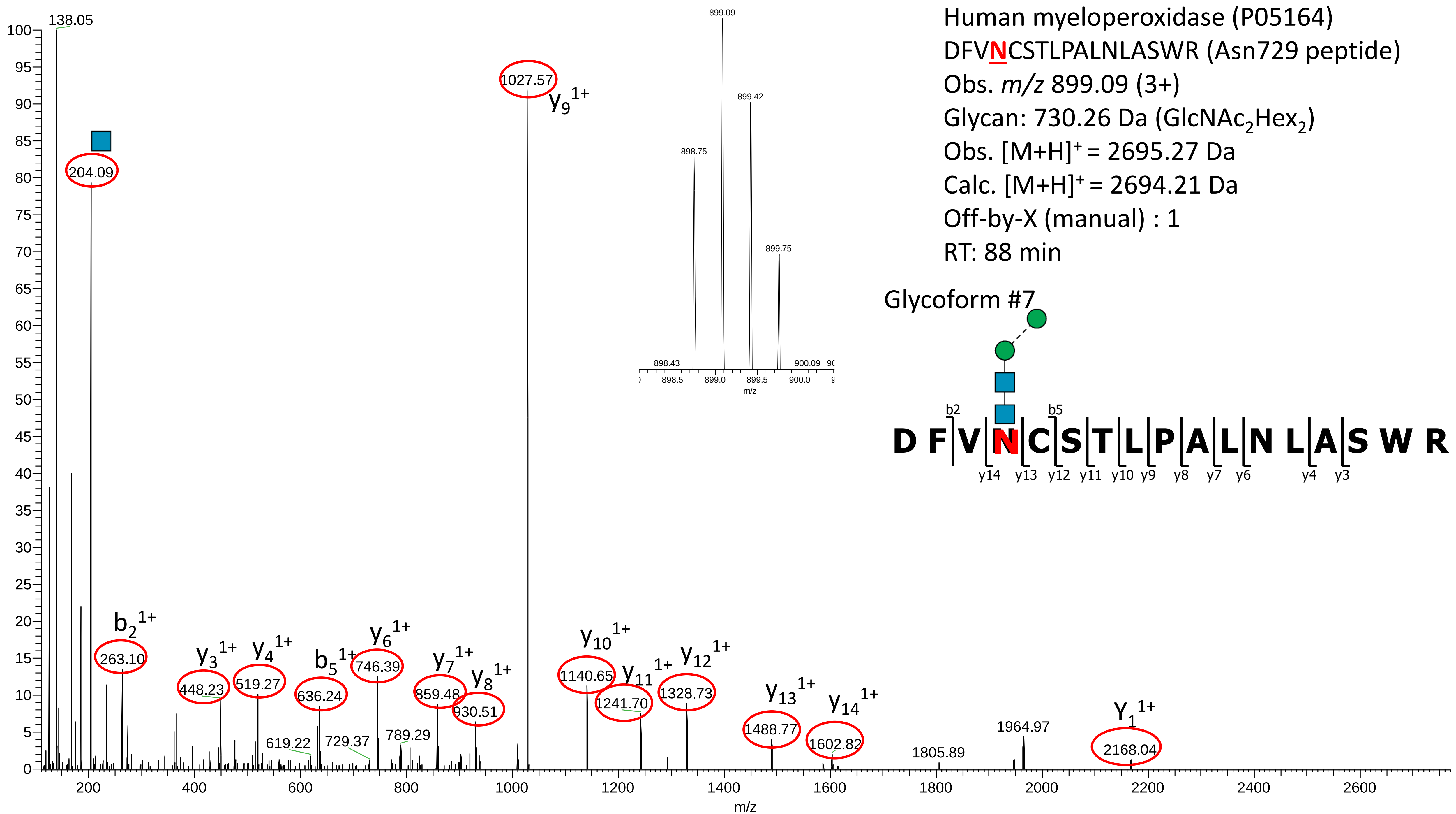
Human myeloperoxidase (P05164)
DFVNCSTLPALNLASWR (Asn729 peptide)
Obs. m/z 899.09 (3+)
Glycan: 730.26 Da (GlcNAc2Hex2)
Obs. [M+H]+ = 2695.27 Da
Calc. [M+H]+ = 2694.21 Da
Off-by-X (manual) : 1
RT: 88 min
Glycoform #7
D F V N C S T L P A L N L A S W R
b2
b5
y14 y13 y12 y11 y10 y9 y8 y7 y6 y4 y3
138.05
204.09
1027.57
y9 1+
b2 1+
263.10
y3 1+
448.23
y4 1+
519.27
b5 1+
636.24
619.22
729.37
y6 1+
746.39
789.29
y7 1+
859.48
y8 1+
930.51
y10 1+
1140.65
y11 1+
1241.70
y12 1+
1328.73
y13 1+
1488.77
y14 1+
1602.82
1805.89
1964.97
Y1 1+
2168.04
898.75
899.09
899.42
899.75
898.43
900.09
m/z

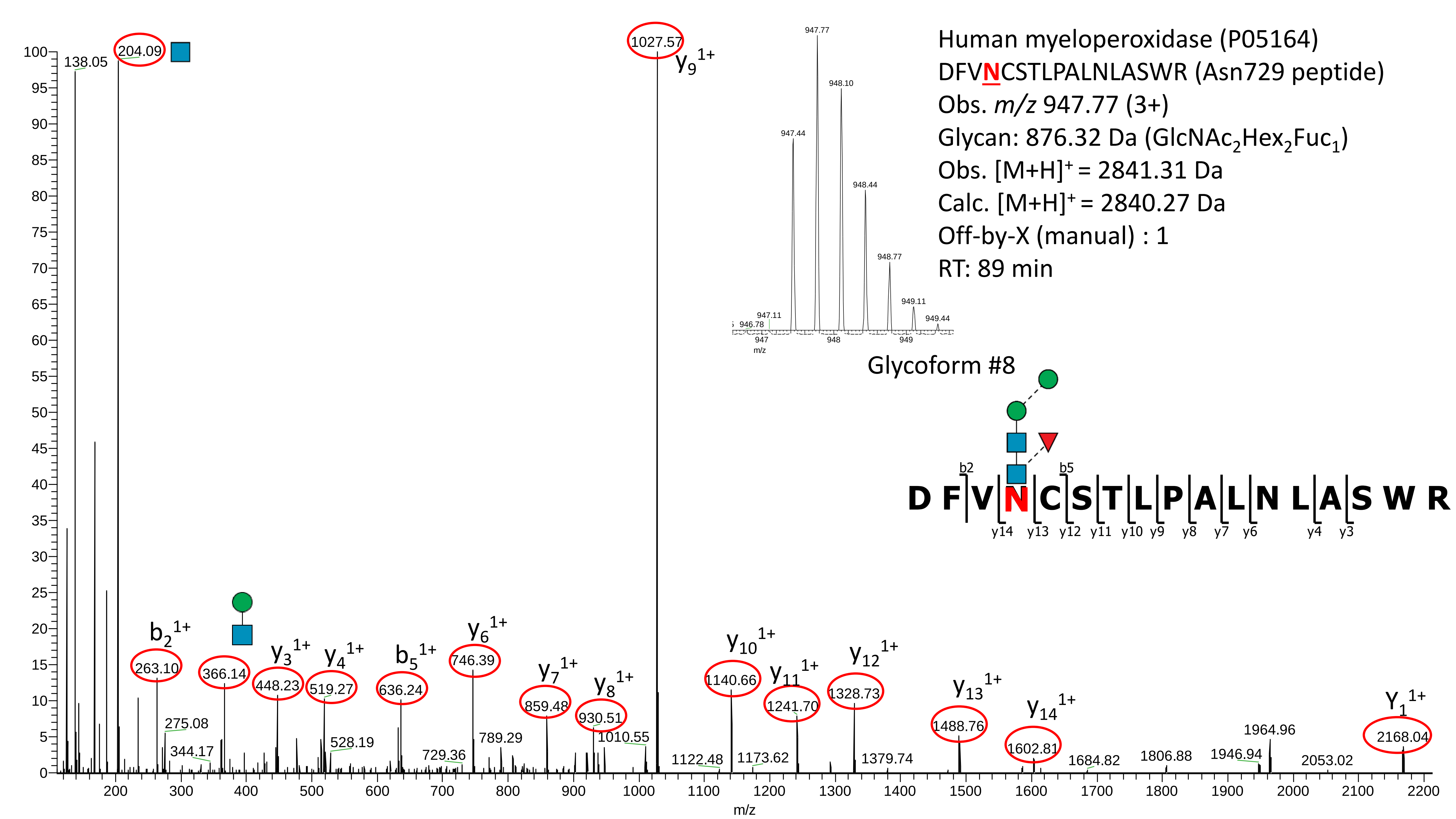
Human myeloperoxidase (P05164)
DFVNCSTLPALNLASWR (Asn729 peptide)
Obs. m/z 947.77 (3+)
Glycan: 876.32 Da (GlcNAc2Hex2Fuc1)
Obs. [M+H]+ = 2841.31 Da
Calc. [M+H]+ = 2840.27 Da
Off-by-X (manual) : 1
RT: 89 min
Glycoform #8
D F V N C S T L P A L N L A S W R
b2
b5
y14 y13 y12 y11 y10 y9 y8 y7 y6 y4 y3
947.44
947.77
948.10
948.44
948.77
949.11
949.44
946.78
947.11
947
948
949
m/z
138.05
204.09
1027.57
y9 1+
b2 1+
263.10
275.08
344.17
366.14
y3 1+
448.23
y4 1+
519.27
528.19
b5 1+
636.24
y6 1+
746.39
729.36
789.29
y7 1+
859.48
y8 1+
930.51
1010.55
y10 1+
1140.66
1122.48
y11 1+
1241.70
1173.62
y12 1+
1328.73
1379.74
y13 1+
1488.76
y14 1+
1602.81
1684.82
1806.88
1946.94
1964.96
2053.02
Y1 1+
2168.04
m/z

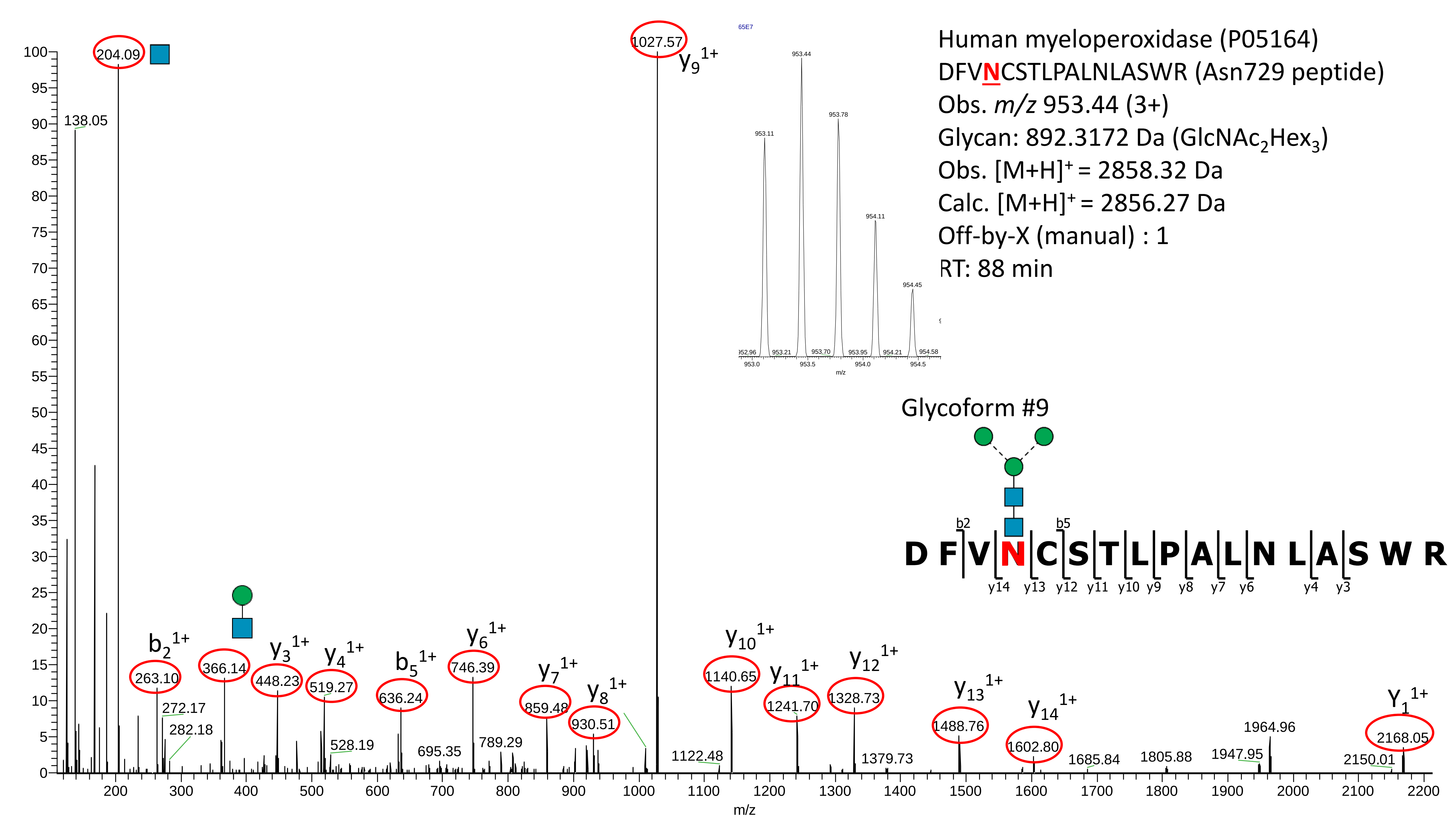

Human myeloperoxidase (P05164)
DFVNCSTLPALNLASWR (Asn729 peptide)
Obs. m/z 953.44 (3+)
Glycan: 892.3172 Da (GlcNAc2Hex3)
Obs. [M+H]+ = 2858.32 Da
Calc. [M+H]+ = 2856.27 Da
Off-by-X (manual) : 1
RT: 88 min
Glycoform #9
D F V N C S T L P A L N L A S W R
b2
b5
y14 y13 y12 y11 y10 y9 y8 y7 y6 y4 y3
953.11
953.44
953.78
954.11
954.45
m/z
204.09
138.05
1027.57
y9 1+
b2 1+
263.10
272.17
282.18
366.14
y3 1+
448.23
y4 1+
519.27
528.19
b5 1+
636.24
695.35
y6 1+
746.39
789.29
y7 1+
859.48
y8 1+
930.51
1122.48
y10 1+
1140.65
y11 1+
1241.70
y12 1+
1328.73
1379.73
y13 1+
1488.76
y14 1+
1602.80
1685.84
1805.88
1947.95
1964.96
2150.01
Y1 1+
2168.05
m/z

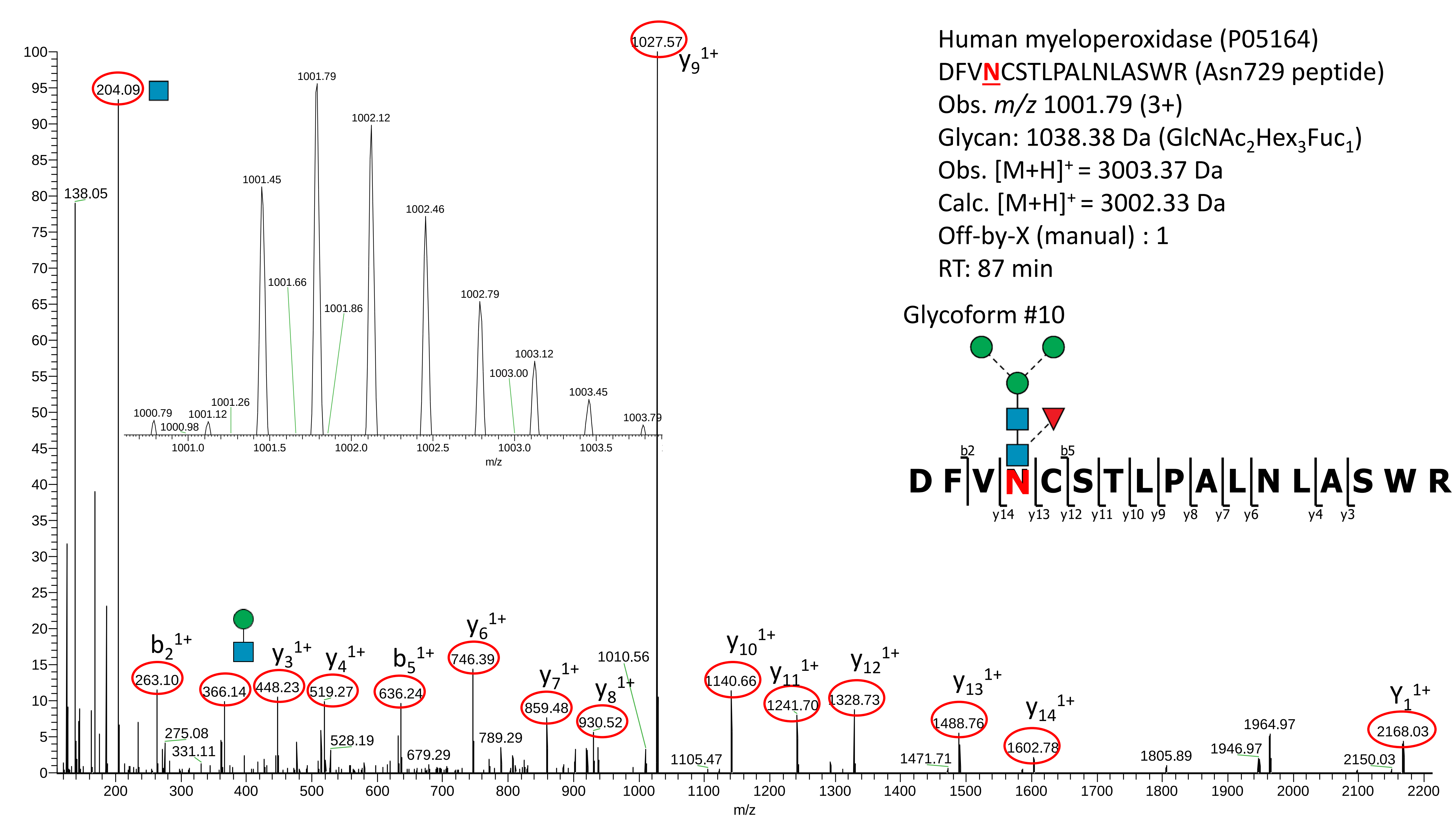

Human myeloperoxidase (P05164)
DFVNCSTLPALNLASWR (Asn729 peptide)
Obs. m/z 1001.79 (3+)
Glycan: 1038.38 Da (GlcNAc2Hex3Fuc1)
Obs. [M+H]+ = 3003.37 Da
Calc. [M+H]+ = 3002.33 Da
Off-by-X (manual) : 1
RT: 87 min
Glycoform #10
D F V N C S T L P A L N L A S W R
b2
b5
y14 y13 y12 y11 y10 y9 y8 y7 y6 y4 y3
1027.57
y9 1+
204.09
138.05
1001.79
1002.12
1001.45
1002.46
1001.66
1001.86
1002.79
1003.12
1003.00
1003.45
1003.79
1000.79
1001.12
1001.26
1000.98
b2 1+
263.10
275.08
331.11
366.14
y3 1+
448.23
y4 1+
519.27
528.19
b5 1+
636.24
679.29
y6 1+
746.39
789.29
y7 1+
859.48
y8 1+
930.52
1010.56
1105.47
y10 1+
1140.66
y11 1+
1241.70
y12 1+
1328.73
1471.71
y13 1+
1488.76
y14 1+
1602.78
1805.89
1946.97
1964.97
2150.03
Y1 1+
2168.03
m/z

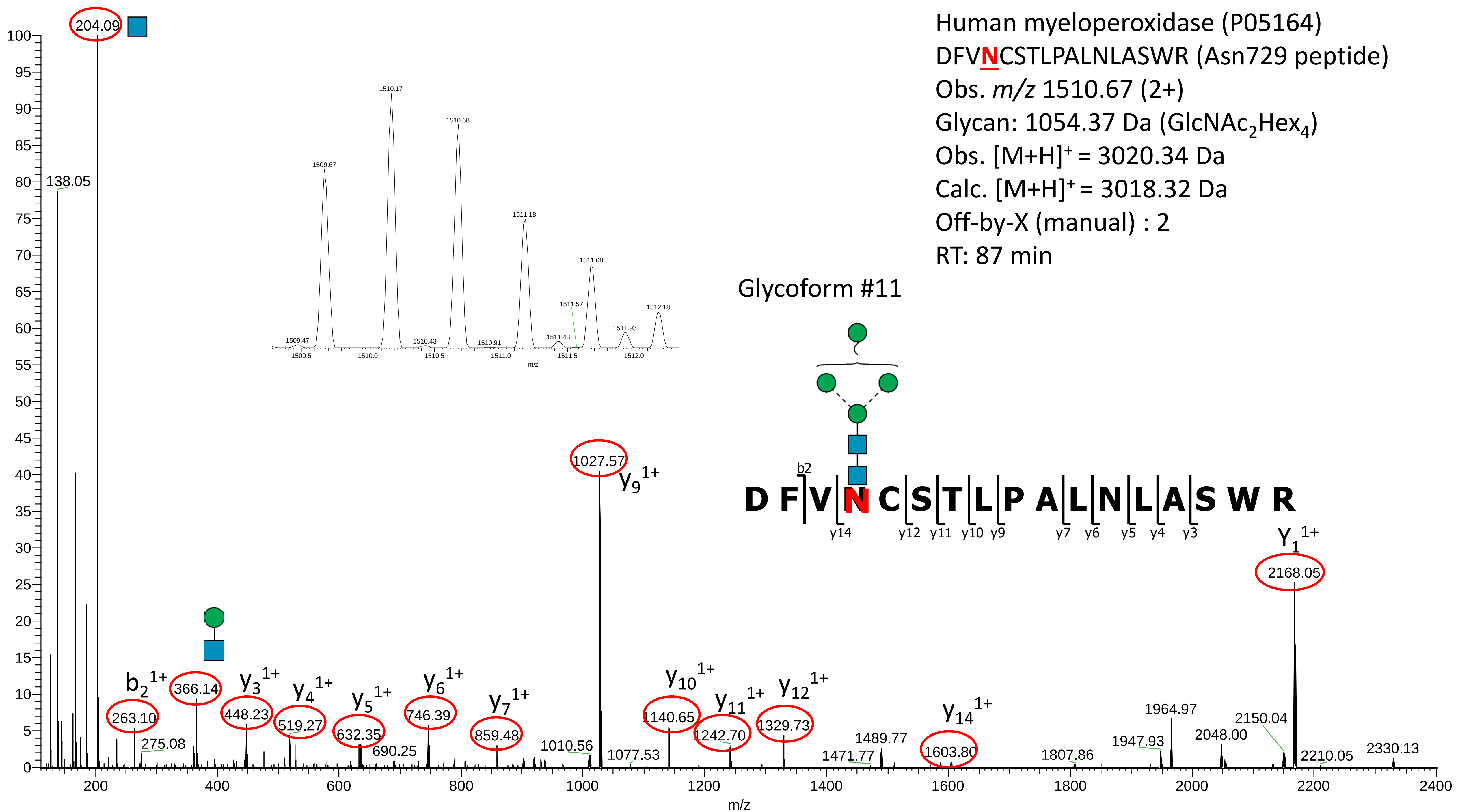

Human myeloperoxidase (P05164)
DFVNCSTLPALNLASWR (Asn729 peptide)
Obs. m/z 1510.67 (2+)
Glycan: 1054.37 Da (GlcNAc2Hex4)
Obs. [M+H]+ = 3020.34 Da
Calc. [M+H]+ = 3018.32 Da
Off-by-X (manual) : 2
RT: 87 min
Glycoform #11
D F V N C S T L P A L N L A S W R
b2
y14 y12 y11 y10 y9 y7 y6 y5 y4 y3
204.09
138.05
1027.57 y9 1+
2168.05 Y1 1+
b2 1+ 263.10
275.08
366.14
y3 1+ 448.23
y4 1+ 519.27
y5 1+ 632.35
690.25
y6 1+ 746.39
y7 1+ 859.48
1010.56
1077.53
y10 1+ 1140.65
y11 1+ 1242.70
y12 1+ 1329.73
1471.77
1489.77
y14 1+ 1603.80
1807.86
1947.93
1964.97
2048.00
2150.04
2210.05
2330.13
1509.47
1509.67
1510.17
1510.43
1510.68
1510.91
1511.18
1511.43
1511.57
1511.68
1511.93
1512.18
m/z

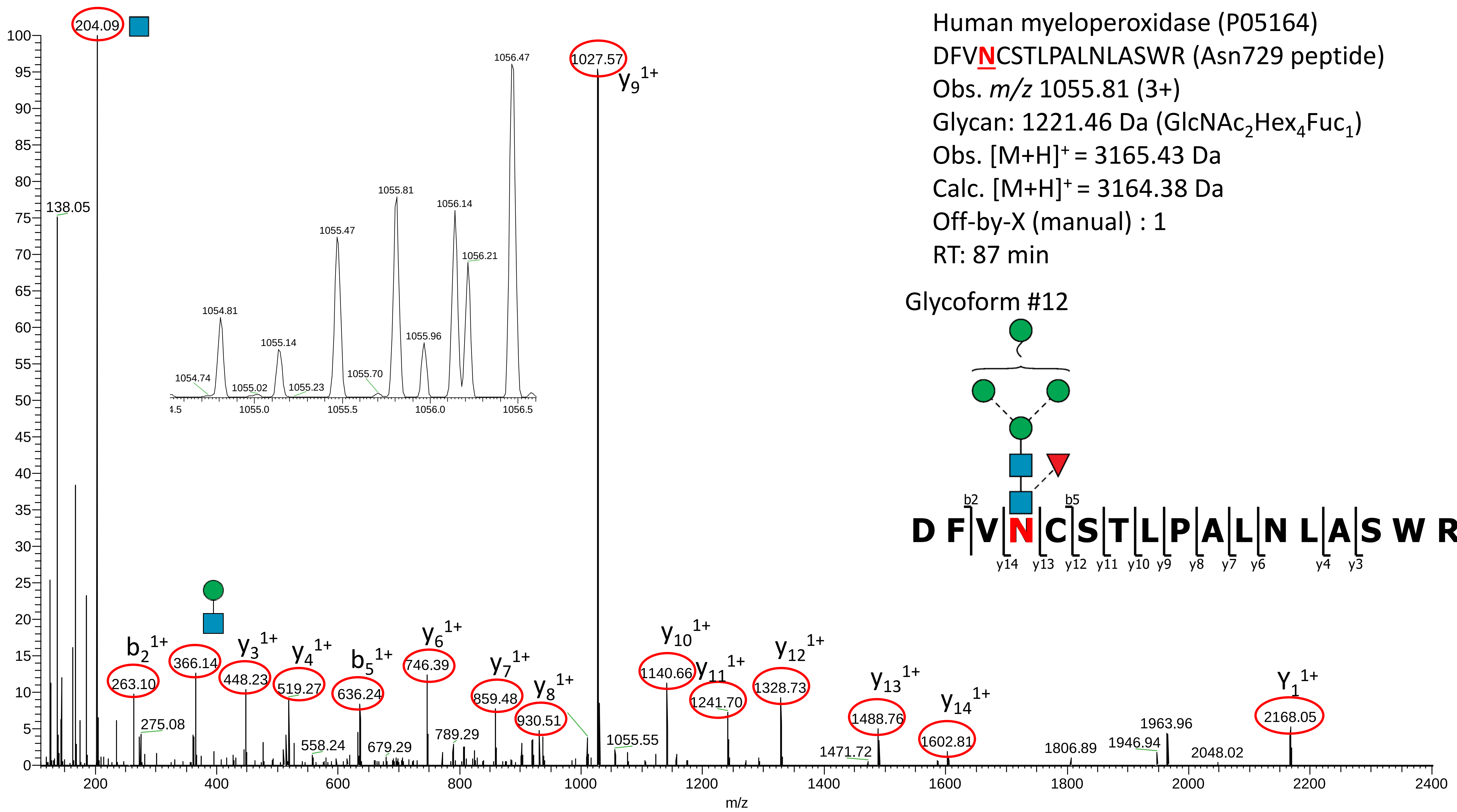
Human myeloperoxidase (P05164)
DFVNCSTLPALNLASWR (Asn729 peptide)
Obs. m/z 1055.81 (3+)
Glycan: 1221.46 Da (GlcNAc2Hex4Fuc1)
Obs. [M+H]+ = 3165.43 Da
Calc. [M+H]+ = 3164.38 Da
Off-by-X (manual) : 1
RT: 87 min
Glycoform #12
D F V N C S T L P A L N L A S W R
b2
b5
y14 y13 y12 y11 y10 y9 y8 y7 y6 y4 y3
204.09
138.05
1027.57
y9 1+
b2 1+
263.10
275.08
366.14
y3 1+
448.23
y4 1+
519.27
558.24
b5 1+
636.24
679.29
y6 1+
746.39
789.29
y7 1+
859.48
y8 1+
930.51
1055.55
y10 1+
1140.66
y11 1+
1241.70
y12 1+
1328.73
1471.72
y13 1+
1488.76
y14 1+
1602.81
1806.89
1946.94
1963.96
2048.02
Y1 1+
2168.05
1054.74
1054.81
1055.02
1055.14
1055.23
1055.47
1055.70
1055.81
1055.96
1056.14
1056.21
1056.47
m/z

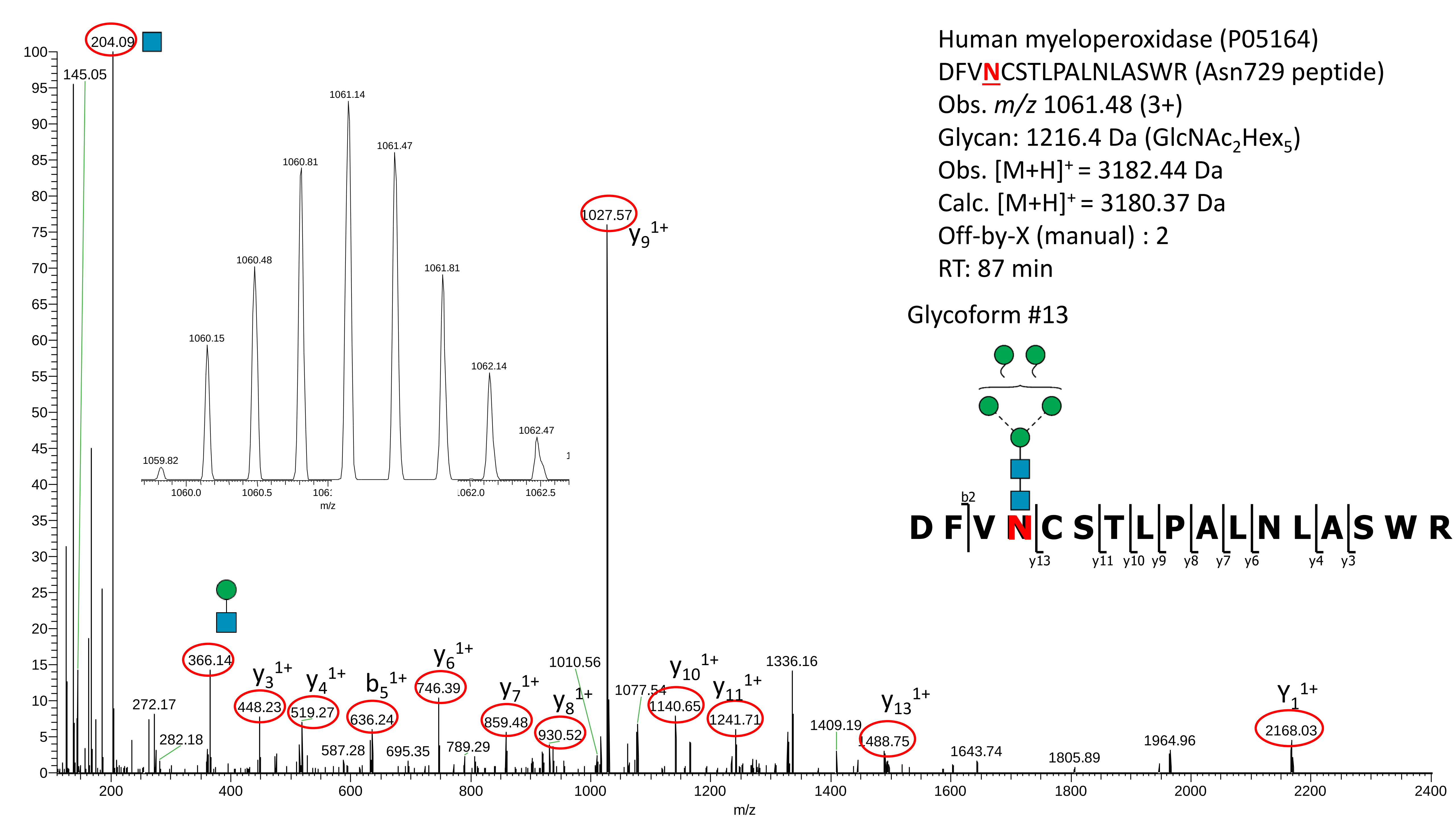

Human myeloperoxidase (P05164)
DFVNCSTLPALNLASWR (Asn729 peptide)
Obs. m/z 1061.48 (3+)
Glycan: 1216.4 Da (GlcNAc2Hex5)
Obs. [M+H]+ = 3182.44 Da
Calc. [M+H]+ = 3180.37 Da
Off-by-X (manual) : 2
RT: 87 min
Glycoform #13
D F V N C S T L P A L N L A S W R
b2
y13 y11 y10 y9 y8 y7 y6 y4 y3
204.09
145.05
1027.57
y9 1+
366.14
272.17
282.18
y3 1+ 448.23
y4 1+ 519.27
b5 1+ 636.24
587.28
695.35
y6 1+ 746.39
789.29
y7 1+ 859.48
y8 1+ 930.52
1010.56
1077.54
y10 1+ 1140.65
y11 1+ 1241.71
1336.16
1409.19
y13 1+ 1488.75
1643.74
1805.89
1964.96
Y1 1+ 2168.03
1059.82
1060.15
1060.48
1060.81
1061.14
1061.47
1061.81
1062.14
1062.47
m/z

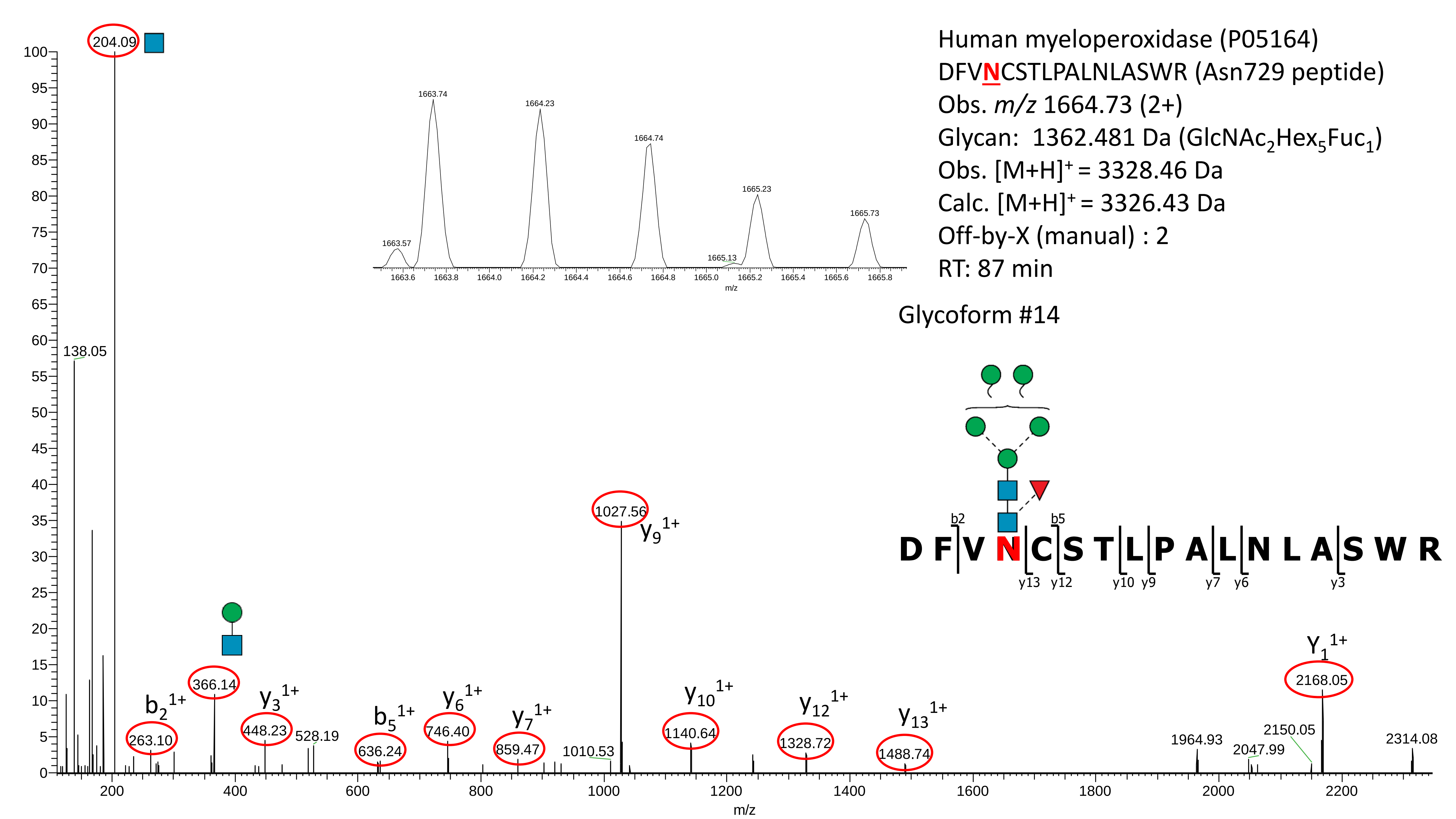

Human myeloperoxidase (P05164)
DFVNCSTLPALNLASWR (Asn729 peptide)
Obs. m/z 1664.73 (2+)
Glycan: 1362.481 Da (GlcNAc2Hex5Fuc1)
Obs. [M+H]+ = 3328.46 Da
Calc. [M+H]+ = 3326.43 Da
Off-by-X (manual) : 2
RT: 87 min
Glycoform #14
D F V N C S T L P A L N L A S W R
b2
b5
y13
y12
y10
y9
y7
y6
y3
1663.57
1663.74
1664.23
1664.74
1665.13
1665.23
1665.73
m/z
204.09
138.05
263.10
b2 1+
366.14
448.23
y3 1+
528.19
636.24
b5 1+
746.40
y6 1+
859.47
y7 1+
1010.53
1027.56
y9 1+
1140.64
y10 1+
1328.72
y12 1+
1488.74
y13 1+
1964.93
2047.99
2150.05
2168.05
Y1 1+
2314.08
m/z

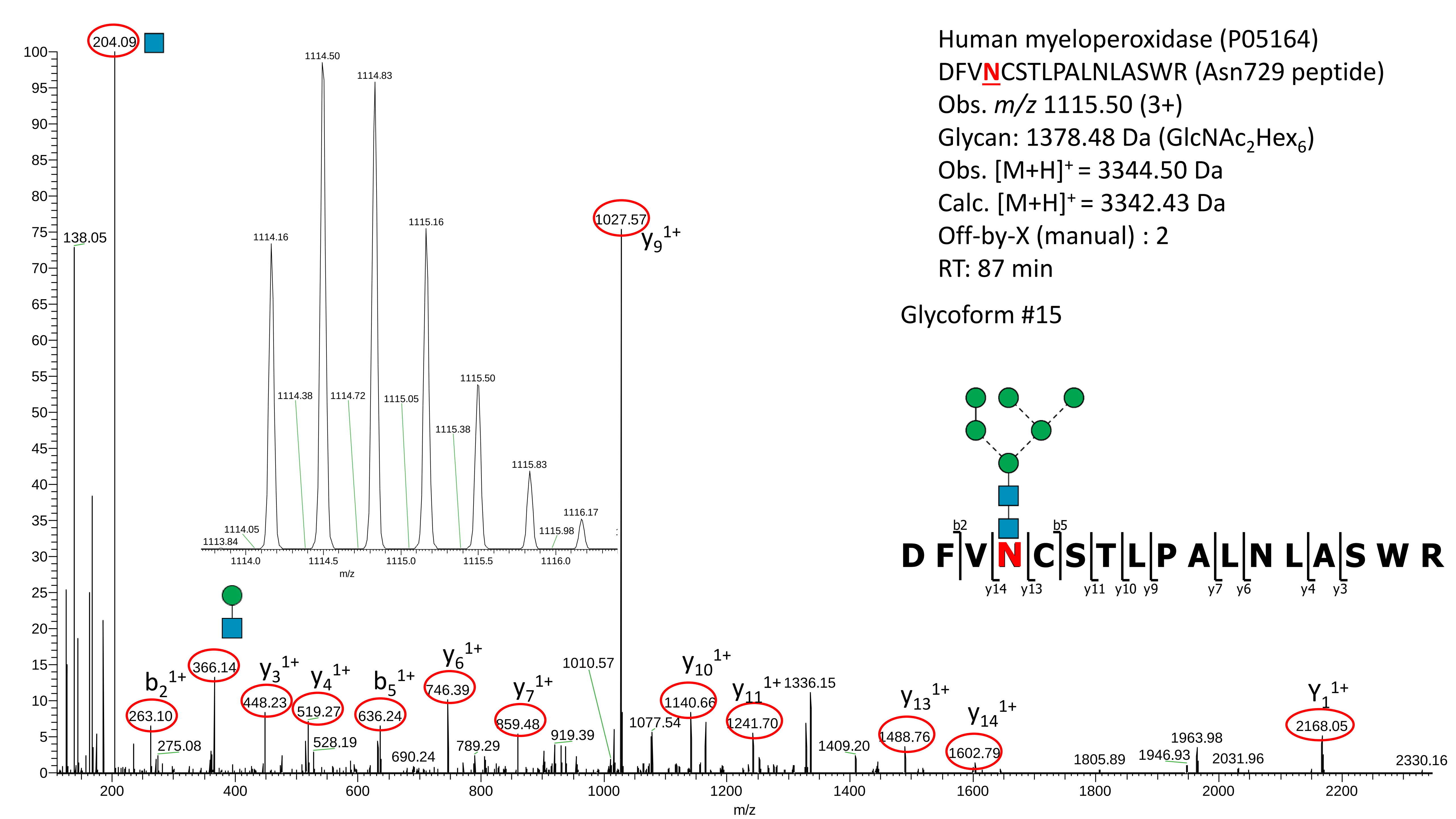

Human myeloperoxidase (P05164)
DFVNCSTLPALNLASWR (Asn729 peptide)
Obs. m/z 1115.50 (3+)
Glycan: 1378.48 Da (GlcNAc2Hex6)
Obs. [M+H]+ = 3344.50 Da
Calc. [M+H]+ = 3342.43 Da
Off-by-X (manual) : 2
RT: 87 min
Glycoform #15
D F V N C S T L P A L N L A S W R
b2 b5
y14 y13 y11 y10 y9 y7 y6 y4 y3
204.09
138.05
1027.57 y9 1+
b2 1+ 263.10
275.08
366.14
y3 1+ 448.23
y4 1+ 519.27
528.19
b5 1+ 636.24
690.24
y6 1+ 746.39
789.29
y7 1+ 859.48
919.39
1010.57
1077.54
y10 1+ 1140.66
y11 1+ 1241.70
1336.15
1409.20
y13 1+ 1488.76
y14 1+ 1602.79
1805.89
1946.93
1963.98
2031.96
Y1 1+ 2168.05
2330.16
m/z
1113.84
1114.05
1114.16
1114.38
1114.50
1114.72
1114.83
1115.05
1115.16
1115.38
1115.50
1115.83
1115.98
1116.17

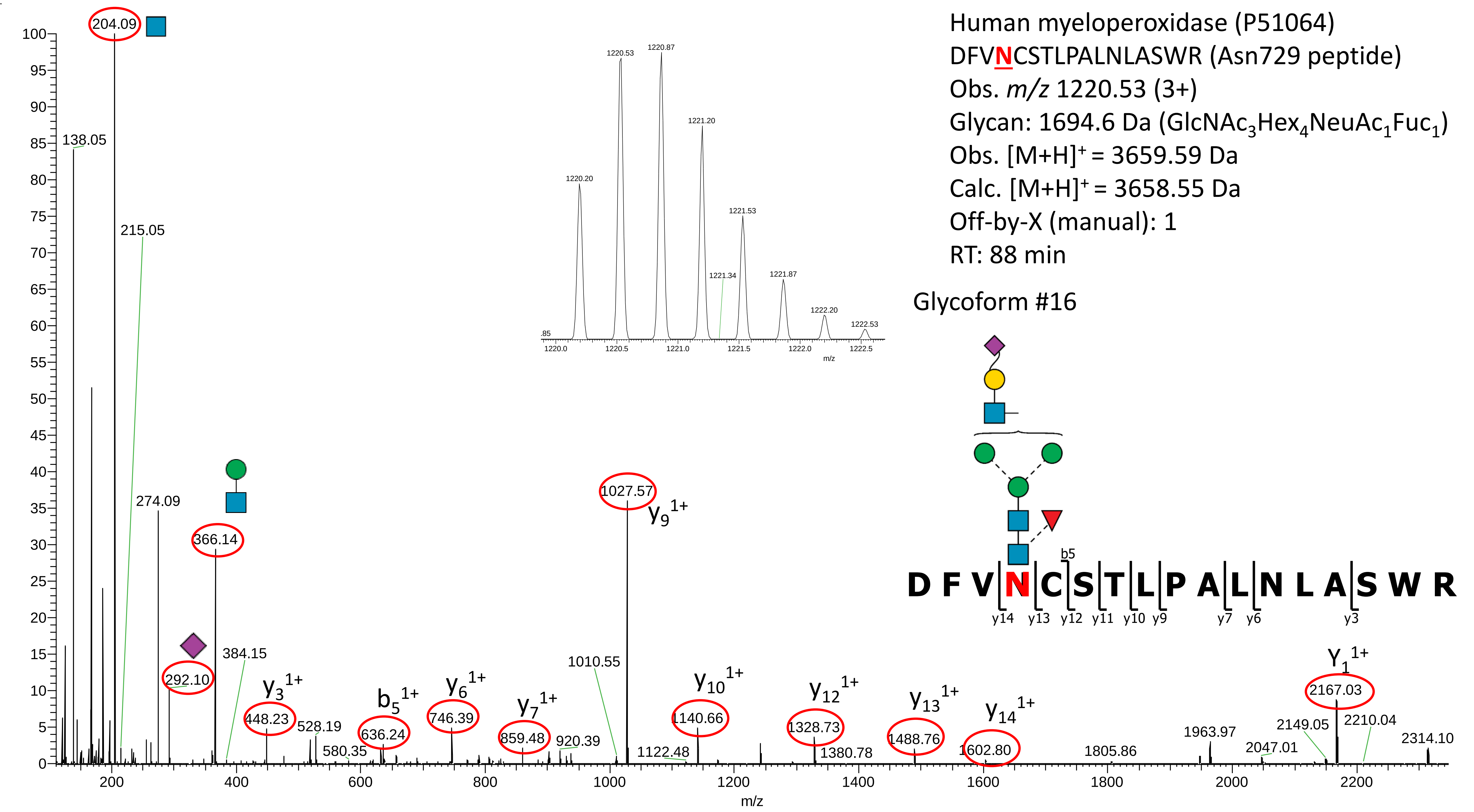

Human myeloperoxidase (P51064)
DFVNCSTLPALNLASWR (Asn729 peptide)
Obs. m/z 1220.53 (3+)
Glycan: 1694.6 Da (GlcNAc3Hex4NeuAc1Fuc1)
Obs. [M+H]+ = 3659.59 Da
Calc. [M+H]+ = 3658.55 Da
Off-by-X (manual): 1
RT: 88 min
Glycoform #16
D F V N C S T L P A L N L A S W R
b5
y14 y13 y12 y11 y10 y9 y7 y6 y3
204.09
138.05
215.05
274.09
366.14
292.10
384.15
448.23
528.19
580.35
636.24
746.39
859.48
920.39
1010.55
1027.57
1122.48
1140.66
1328.73
1380.78
1488.76
1602.80
1805.86
1963.97
2047.01
2149.05
2167.03
2210.04
2314.10
y9 1+
y3 1+
b5 1+
y6 1+
y7 1+
y10 1+
y12 1+
y13 1+
y14 1+
Y1 1+
m/z
1220.20
1220.53
1220.87
1221.20
1221.34
1221.53
1221.87
1222.20
1222.53
